# Biophytometallurgy: biomining metals from plant resources

**DOI:** 10.64898/2026.08.27.747594

**Authors:** Daniel A. Dailey, Edmaritz Hernández-Pagán, Stella Bailey, Nicolas E. Raffaele, Sofia T. Bavaresco, Abbey Piatt-Price, Juliana S. A. Carneiro, Rachel N. Austin, Colleen J. Doherty, Scott Banta

## Abstract

The physicochemical controls governing metal acquisition, release, and redistribution across biological interfaces remain poorly understood. Biophytometallurgy—the microbially assisted release and recovery of plant-associated metals—was used to probe the directionality of the mechanisms controlling nickel (Ni) and rare earth element (REE) release from plants of the genus *Phytolacca* during solid–liquid extraction. Ni, an established phytomining target, was included as a comparator for chemical extraction. Bulk characterization did not support a detectable dominant crystalline REE-phosphate-like compartment in hydroponically enriched shoots. Dissolution and rebinding experiments instead revealed chemically accessible nickel and REE compartments, the latter of which had behaviors consistent with apparent equilibrium-like partitioning under mildly acidic conditions. During sulfur biooxidation, *Acidithiobacillus ferrooxidans* promoted REE release while providing a competing cell-associated REE sink. Consequently, aqueous REE concentrations reflected net redistribution among the separable plant, solution, and microbial phases instead of dissolution alone. These results establish a framework for studying metal partitioning across complex and coupled biological systems and support a route for aqueous REE recovery from plants without thermochemical conversion to ash.

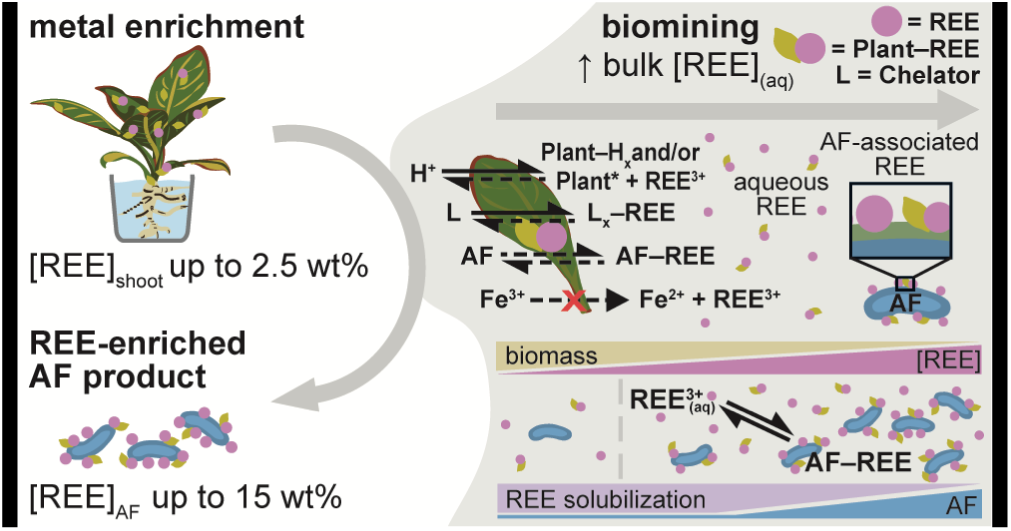

## Introduction

Hyperaccumulating plant species have a unique ability to concentrate metal(loid)s in their tissues.^1–4^ The idea of using hyperaccumulating plants to concentrate metals as crops (i.e., phytomining) is at least 30 years old.^5–7^ Only recently have efforts been made to commercialize this strategy for nickel (Ni).^8^ Using plants to pluck metals from the environment could form the basis of a sustainable mining practice.^7,9^ Rare earth elements (REEs) include scandium, yttrium, and the lanthanoids.^10^ Extending phytomining to REEs is interesting because conventional REE mining and refining can burden both the environment^11,12^ and the societies near regions that host these ore deposits.^9,13,14^ The use of REE hyperaccumulating species^15^ could possibly reduce the cost of metal production by pre-concentrating some dilute and/or unconventional non-ore feedstocks,^9,16^ such as contaminated soils,^17^, acid mine drainage, and possibly even the ocean.^18^

However, the inherently limited carbon efficiency of thermochemical methods to generate concentrated ash, or bio-ore,^19,20^ before subsequent metal extraction processes reduces the sustainability of the overall phytomining process.^9,21^ As an alternative, solid–liquid extraction can dissolve, or leach, target metals into solution while leaving the plant in a solid form (i.e., phytometallurgy).^22,23^ This approach could create opportunities to valorize the metal-depleted biomass.^9^ For example, this residue could be used as a feedstock for biochemical conversion of lignin into valuable bioproducts.^24^ Perhaps this non-ash residue could even serve as fertilizer for the phytomining process itself. Yet, studies reported that the cost and secondary pollution of chemicals were prohibitive for REE phytometallurgy.^25,26^ Few have considered the approach since.^23,27^

The “optimal” metal recovery strategy depends on the nature and strength of metal binding within plant tissues. Since REE binding environments remain incompletely understood,^28^ the advantages of solid– liquid extraction may not have been fully appreciated yet. In fern-type REE hyperaccumulators, current models suggest REEs are associated with silicon/pectin-, phosphate-, and monazite-like mineral-associated compartments (i.e., phytomineralization) within the apoplast and cell wall.^29–31^ In some of these ferns, an REE transporter has been identified which prefers the economically less significant light REE (LREE) uptake.^32^ These models suggest at least part of the plant-associated REE pool may be immobilized in relatively strong, poorly reversible, or even refractory mineral-like environments.^31,33^ Though the mechanisms governing metal release in these systems are not understood, refractory REE compartments^31,33,34^ may have increased reagent demand during solid–liquid extraction in previous studies.^25,26^

The working model for REE accumulation in *Phytolacca* species is mechanistically distinct, which could be favorable for phytometallurgy. Pectin-rich cell walls and phosphate-associated apoplastic sites likely still serve as key binding and detoxification sinks.^35^ Phosphate-associated retention has also been implicated in the root apoplast.^36^ REE mobilization and translocation have been linked to organic acids, such as citrate, oxalate, and related carboxylates.^36–38^ These processes can lead to an economically favorable tendency to enrich heavy REEs (HREEs) within the aerial tissues (i.e., shoots).^37–39^

Therefore, a portion of the REEs in *Phytolacca* may exist in comparatively labile or exchangeable ligand-bound compartments. If REE acquisition and storage in *Phytolacca* are controlled by dative bonds,^35^ we hypothesized REE could be released by shifting equilibria under mild conditions. We also thought this strategy could be relevant for Ni release because its association with cell-wall sites and endogenous carboxylates has been established in other metal-accumulating plants.^40,41^ Protonation of these binding sites can weaken metal retention,^42^ while adding ligands could favor release by stabilizing dissolved metal complexes for metal targets.^26,43^

If plant-associated REEs exchange reversibly with the aqueous phase, removal of dissolved REE ions by a competing sink could favor further release from the plant by the law of mass action.^42^ An acidophilic chemolithoautotroph like *Acidithiobacillus ferrooxidans* can generate protons by its oxidation of reduced inorganic sulfur (S^0^) compounds under aerobic conditions.^44^ It can also tolerate REEs^45^ and can putatively bind them under acidic conditions,^46^ possibly resulting in a bacterial phase that is more enriched in REEs than the plant precursor. Thus, this organism could be used as a living dual-function material suited to investigate the reversibility of REE release.

In this study, hydroponically enriched *Phytolacca* tissues were harvested, dried, and used as an experimental model to examine the nature of metal–*Phytolacca* interactions under defined conditions. We then used *A. ferrooxidans* to extract, recover, and bio-concentrate REEs from the harvested plants via solid–liquid extraction without the direct conversion of plant tissue to carbon emissions (i.e., biophytometallurgy).

## Methods

### Materials and characterization

Ultrapure H_2_O was used in all experiments unless otherwise noted. Chemicals, materials, and equipment; bacterial strains; and media formulations are provided in Tables S1–S3, respectively.

*A. ferrooxidans* were used to demonstrate the independent and integrated REE bioleaching and biorecovery processes, and *Escherichia coli* DH5*α* were used as a neutrophilic comparator for REE binding (Texts S1–S4; Figures S1–S2).

*Phytolacca acinosa* and *Phytolacca americana* tissues were obtained from plants grown in soil and separate hydroponically grown plants subjected to Ni and mixed REE treatment (Text S5). Dried tissues and calcined ash residues were characterized to determine whether metal enrichment altered the plant matrix. Characterization included elemental analysis (Texts S6–S7; Equations S1–S2), thermogravimetric analysis (TGA) (Text S8; Equations S3–S4), calcination (Text S9), X-ray diffraction (XRD) (Text S10; Equation S5), Fourier-transform infrared spectroscopy (FTIR) (Text S11; Equations S6–S14), and scanning electron microscopy with energy dispersive X-ray spectroscopy (SEM-EDS) (Text S12). Water washing tests were used to test whether REEs were associated with detectable mineral-like phases after thermal treatment (Text S13; Equations S15–S22). BET surface area analysis was also performed on select *P. americana* shoot tissues, *A. ferrooxidans*, and S^0^ formulations (Text S14).

### Solid–liquid extraction of target metals from *Phytolacca*

Solid–liquid extraction was used to determine whether acid- and ligand-assisted leaching could solubilize Ni and REEs from enriched *Phytolacca* tissues under incubation conditions relevant to acidophilic bioleaching (Text S15; Equations S23–S34).

We encourage readers to consider the limitations of this solubilization calculation methodology (Text S15).

### REE repartitioning to (un)leached *Phytolacca*

REE repartitioning tests were used to evaluate whether prior extraction depleted the apparent REE-binding capacity of dried *P. americana* shoots. Selected Ce/Dy-enriched *P. americana* residues from the electrolyte, acid, and chelator extraction experiments were separated from the supernatant after centrifugation (4,695 x g, 7 min), washed with H_2_O (5 mL) before being centrifuged again, and dried (30 °C, 2 d). In addition to unleached *P. americana* shoots, these residues were re-exposed to a standardized Ce^3+^/Dy^3+^ reload solution (50 µM each, pH 4.5, 0.1 M KNO_3_, 30 °C, 1 d, 150 rpm). The reload experiment compared the moles of REEs released during the preceding extraction with the moles of REEs removed from the reload solution during the subsequent rebinding step (*R*_reload_). The final corrected REE loading after reloading (*q*_after,corr_) was compared with that of non-leached Ce/Dy-enriched *P. americana* shoots to determine whether prior leaching or REE-enrichment changed the endpoint REE-binding capacity of the remaining solids. Selected untreated, post-leached, and post-reloaded solids were analyzed by FTIR. Equations used to calculate the operational reload efficiency and REE adsorption capacity are provided in Equations S35–S45.

### Microbial REE binding

REE binding by *A. ferrooxidans* and *E. coli* DH5*α* was measured to determine whether these bacteria could concentrate soluble REEs under acidic conditions relevant to selected H_2_SO_4_ plant leachates generated herein (Text S16; Equations S46–S63).

*A. ferrooxidans* subjected to pretreatments intended to disrupt the outer membrane or metabolic capabilities were also used for some REE binding tests (Text S17; Equation S64). In an attempt to improve the REE binding capacity of *A. ferrooxidans*, growth curves were collected for cultures of F2S medium with and without 15 µM Tb_2_(SO_4_)_3_ supplementation (Text S18).

### Solution properties of *Phytolacca* leachates

Some 0.22 µm-filtered H_2_SO_4_ and H_2_O leachates were characterized to determine whether exposure to plant tissues altered solution chemistry in ways that could affect metal speciation or bacterial REE binding. The pH, solution potential (ORP), ferrous iron (Fe^2+^) concentration, total dissolved metals, and SYBR Green fluorescence signal were measured (Text S15). For some samples (n = 2), the buffering capacity was determined by NaOH–HCl titrations of the filtrates diluted in H_2_O (Text S19; Equations S65–S76). To assess whether dissolved metals were associated with higher-molecular-weight plant-derived solution components, these filtrates were separated using 3 kDa regenerated-cellulose centrifugal ultrafiltration devices (Text S20; Equation S55, Equations S77–S84).

### Biomining Phytolacca using A. ferrooxidans

*A. ferrooxidans* tolerance to *P. americana* shoots was first evaluated in Fe^2+^- and S^0^-containing media (Text S21). Integrated bioleaching–biorecovery experiments were performed to test whether sulfur oxidation by *A. ferrooxidans* could release REEs from *Phytolacca* shoots while partitioning released REEs onto the bacteria, followed by recovery of separable coproducts (Text S18, Text S21; Equation S32, Equations S85– S86, Equations S87–S94).

Separate abiotic adsorption experiments with brown and yellow S^0^ were performed using Ce^3+^ or Dy^3+^ to assess apparent REE partitioning to sulfur (Text S22; Equations S32–S34, Equations S95–S101).

### Statistical analysis

Statistical analyses were selected according to the experimental design and raw or transformed response variable for each dataset as indicated in the referenced Supplementary Tables. Unless otherwise noted, data are reported and plotted as mean *±* standard deviation, with biological replicates overlaid (n *≥* 3). Statistical significance was evaluated at *α* = 0.05 unless otherwise noted.

## Results

### *Phytolacca* was hydroponically enriched in REEs and Ni

To test this proof-of-concept, *P. acinosa* and *P. americana* were grown hydroponically (Figure 1a) and supplemented with Ni- or mixed REE-chloride salts to induce accumulation of the supplemented metal^47^ (Table S4; Figure 1b). In *P. acinosa*, REE supplementation increased REE concentrations in roots and shoots by about two orders of magnitude relative to non-supplemented controls, with no resolved root–shoot difference within either supplementation condition (Table S5). REE supplementation did not significantly increase shoot HREE concentrations relative to matched roots (Table S6). After log_10_ transformation, shoots had higher HREE/LREE molar ratios than roots (Table S7); only the shoot ratios were greater than unity (Table S8). The Ni concentrations in *P. acinosa* shoots exceeded matched root concentrations at both 25 and 50 mM Ni supplementation, though neither shoots nor roots showed a statistically resolved difference between these supplementation levels after multiplicity adjustment (Table S9). Analysis of *P. americana* further showed that metal supplementation generated enriched *Phytolacca* shoot biomass. Ce/Dy treatment increased REE concentrations in *P. americana* shoots relative to hydroponic non-supplemented controls, while hydroponic and soil-grown non-supplemented controls showed no resolved difference (Table S10). REE-supplemented *P. americana* shoots also had HREE/LREE molar ratios greater than unity (Table S10). The REE-supplemented *P. americana* shoots did not look notably different than the controls (Figures S3–S4), though the intensity of the Dy and Ce emission spectra detected on the surface was qualitatively higher than the controls (Figures S5–S6). The *P. americana* shoots had higher specific surface area than hydroponic controls after Ce/Dy treatment (Table S11), though these samples were ground with different equipment to avoid metal contamination.

**Figure 1:**
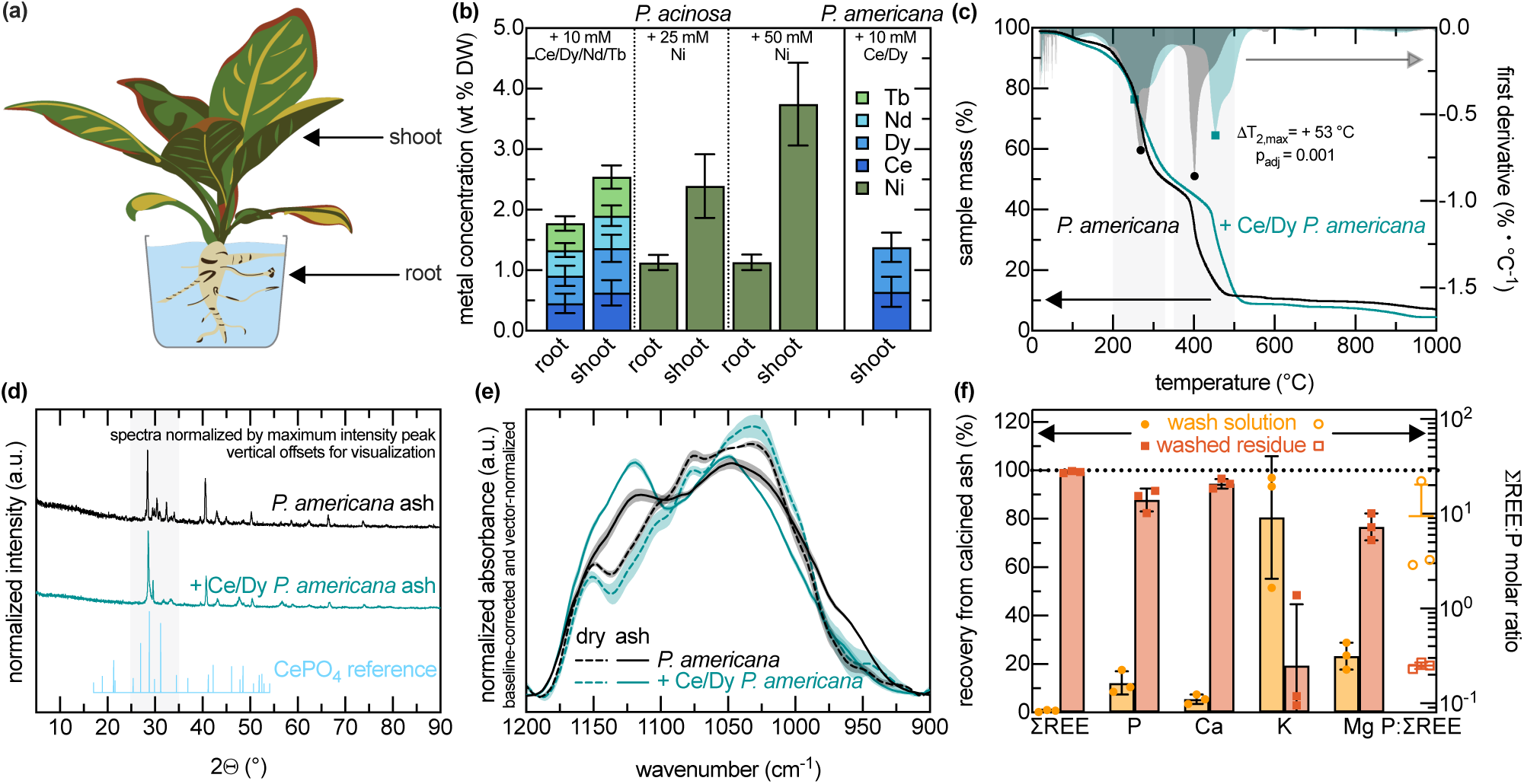
REE-enriched *Phytolacca* shoots did not exhibit characteristics of a dominant crystalline REE-phosphate phase. (a) Schematic of harvested *Phytolacca* tissues analyzed. (b) Elemental concentrations of Ce, Dy, Nd, Tb, and/or Ni in *Phytolacca* tissues after the hydroponic supplementation conditions indicated on the panel (n = 3–9). Metal concentrations reported on dry mass basis (DW). (c) Representative TGA (lines, left y-axis) and DTG (shaded profiles, right y-axis) profiles of non-supplemented and Ce/Dy-enriched *P. americana* shoots (n = 1). (d) XRD patterns of calcined ash residues (n = 1) compared with the calculated powder diffraction pattern for monazite-(Ce), generated from ICSD-79746 (ICSD release 2026.1).^51,52^ (e) Region-specific baseline-corrected and vector-normalized FTIR spectra of dried *P. americana* shoots without (black) and with Ce/Dy-enrichment (green) before (dashed lines) and after calcination (solid lines) under the indicated supplementation conditions. (f) Elemental release after water-washing of calcined ash residues from Ce/Dy-enriched *P. americana* shoots.

TGA showed that metal-supplemented *P. americana* shoots were more resistant to thermal decomposition of organic matter than controls (Figure 1c). Consistent with mass-loss traces and derivative thermogravimetry (DTG) metrics (Table S12), REE-supplemented *P. americana* shoots decomposed at higher characteristic temperatures, particularly in the higher-temperature derivative feature, than hydroponic controls (Table S13). Greater thermal recalcitrance was also qualitatively observed in REE-treated *P. acinosa* roots and shoots, dose-dependently in Ni-enriched *P. acinosa* shoots, and in soil-grown *P. americana* shoots relative to hydroponic non-supplemented controls (Table S12; Figure S7). The post-TGA residues were enriched in inorganic constituents relative to the starting tissues, with Ca, K, Mg, P, Fe, Ni, and REEs concentrations typically one order of magnitude higher (Table S14).

Because TGA indicated that most organic combustion was complete by 600 *^◦^*C while some inorganic residue remained, calcination at 600 *^◦^*C was used to generate REE-bearing ash to test for behavior consistent with a thermally transformed REE-phosphate-like phase.^31^ Similar to the TGA residues, calcined ashes were enriched in inorganic constituents relative to the dried tissues after 600 *^◦^*C treatment (Table S15). However, REE/P molar ratios did not indicate selective REE enrichment relative to P after 600 *^◦^*C calcination. Post-TGA residues of REE-supplemented *P. americana* had lower REE/P molar ratios than both dried tissues and residues calcined at 600 *^◦^*C (Table S16). XRD patterns did not show distinct CePO_4_-like reflections in *P. americana* relative to non-REE-supplemented controls before or after calcination (Figure 1d, Figure S8).^48^ After region-specific baseline correction and vector normalization, REE-supplemented *P. americana* shoots, especially after calcination, showed broad absorbance in the PO_4_^3−^/P–O/C–O stretching envelope (Figure 1e),^49,50^ which was also observed in hydroponic and soil-grown controls (Figure S9).

Although XRD and FTIR did not support a resolved monazite-like REE phosphate phase, water-washing tests were also performed on the ash to indirectly probe for behaviors of a refractory REE-P phase.^31^ For REE-enriched *P. americana* ash, REE release to the wash solution was not higher than P release (Figure 1f; Table S17). However, REE release was lower than K and Mg release and not resolved from Ca release, whereas P release did not differ from the other measured major inorganic elements (Table S17). REE-enriched *P. americana* ash did not consistently show resolved major-element release behavior relative to ash from other tissue types, including soil- and hydroponically grown *P. americana* and hydroponically grown *P. acinosa* controls (Figure S10).

Together, these results show that *Phytolacca* shoots were enriched in REEs but do not support crystalline phosphate-like geological REE mineralization as the dominant REE compartment, even after calcination.

### Solid–liquid extraction dissolved REEs and Ni from *Phytolacca*

After generating metal-enriched biomass, solid–liquid extraction was used to investigate metal release from enriched *Phytolacca* source materials under bioleaching-relevant temperature and agitation conditions. Shoot tissues were prioritized due to faster and greater REE solubilization than from matched *P. acinosa* roots in preliminary tests using Na_2_EDTA leaching (Figure S11). H_2_O-only controls and several H_2_SO_4_, citric acid, and Na_2_EDTA extraction conditions reached apparent target-metal solubilization plateaus within 7 d for the Ni- and Ce/Dy-enriched *P. americana* shoots (Figure S12); subsequent extraction tests were therefore compared at this endpoint. Extended leaching using H_2_O or H_2_SO_4_ under static benchtop conditions did not measurably increase the final solubilized Ni fraction from *P. acinosa* but increased REE solubilization from enriched *P. americana* (Figures S12–S13).

Across *P. americana* shoot extractions, REE solubilization increased above the H_2_O-labile baseline with increasing reactant-to-REE molar ratio (Figure 2a). For H_2_SO_4_, REE solubilization followed a single log-linear relationship with nominal H_2_SO_4_-to-REE ratio whether the ratio was varied by acid concentration or biomass loading (Table S18). At equimolar reactant concentrations ranging from 10*^−^*^2^ − 10^−1^, K_2_SO_4_ and KNO_3_ solubilized much less REEs than H_2_SO_4_ (Table S19; Figure S14a). Although acid-specific behaviors were detected across the mineral and organic acids tested at matched pH (Figure S14b), the pooled acid-leaching data were consistent with a shared pH-dependent trend (Table S20). There were significant and insignificant increases in the extent and Dy/Ce solubilization selectivity across the conditions with SO_4_^2−^ (Table S21).

**Figure 2:**
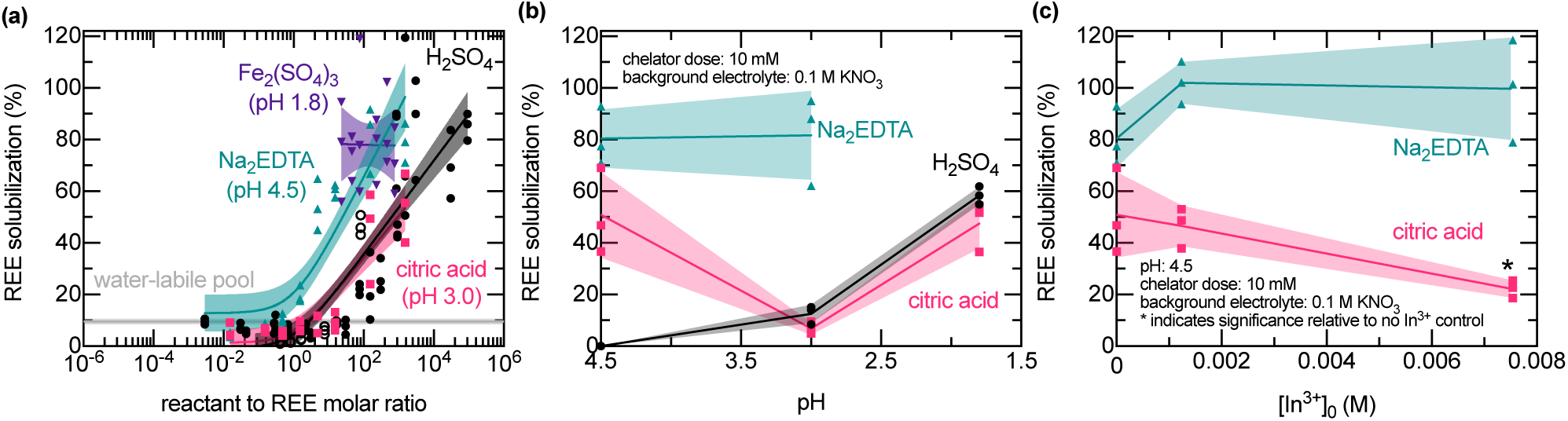
pH and ligand availability modulated acid- and chelator-assisted REE solubilization from *P. americana* shoots. (a) REE solubilization above the H_2_O-labile compartment as a function of nominal reactant:REE molar ratio for tested reactants. Solid lines show fitted logarithmic models [*Y* = *a·*log(*X* +1)+*b*] and shaded regions show 95% confidence intervals (CIs), with biological replicates overlaid (n = 3–6). The horizontal gray line and shaded region show H_2_O-only controls (n = 3). (b) REE solubilization at fixed reactant dose, biomass loading, and background electrolyte across matched-pH H_2_SO_4_, citric acid, and Na_2_EDTA conditions. (c) REE solubilization from fixed-dose citric acid and Na_2_EDTA solutions pre-loaded with increasing In^3+^ concentrations.

Next, the feasibility of using Fe^3+^ to enhance REE release beyond the matched-acidity baseline was investigated (Figure 2a). Although Ce/Dy-enriched *P. americana* shoots may have produced higher Fe^2+^ concentrations than matched no-tissue controls after incubation under identical conditions (Figure S15), increasing Fe_2_(SO_4_)_3_ supplementation up to 0.05 M did not concomitantly increase REE solubilization relative to the matched-pH control (Table S22).

At fixed pH, increasing citric acid and Na_2_EDTA concentrations increased REE solubilization within the tested ranges (Figure 2a). To decouple acidity and chelation, fixed-dose chelator tests were performed under matched-pH conditions with KNO_3_ background electrolyte to reduce ionic-strength differences across reagent conditions (Figure 2b). Under these conditions, matched-pH comparisons with the H_2_SO_4_ baseline showed that citric acid increased REE solubilization only at pH 4.5, whereas Na_2_EDTA increased REE solubilization at pH 3.0 and 4.5 (Table S23).

To test whether chelator-assisted release depended on unoccupied ligand availability, fixed-dose chelator solutions were pre-loaded with In^3+^ before extraction (Figure 2c). Increasing In^3+^ dose suppressed citrate-assisted REE solubilization at the highest tested In^3+^ loading but did not suppress Na_2_EDTA-assisted release, yielding a significant chelator-by-In^3+^ dose interaction (Table S24); dissolved In concentrations changed minimally across these conditions (Figure S16).

Because total REE solubilization does not indicate whether individual REEs were released proportionally, Dy and Ce solubilization were compared across the acid- and chelator-assisted *P. americana* shoot leaching tests. Without KNO_3_ background electrolyte, Dy solubilization generally exceeded Ce solubilization (Figure S17). Multiple linear regression of log_2_-transformed, feed-normalized HREE/LREE molar selectivity identified higher transformed selectivity for Na_2_EDTA and lower transformed selectivity for HNO_3_ than for H_2_SO_4_ without KNO_3_, while adding KNO_3_ decreased transformed selectivity within H_2_SO_4_ (Table S25).

For the *P. acinosa* shoots, increasing H_2_SO_4_ and Na_2_EDTA concentrations at pH 4.5 increased Ni solubilization, and whereas citric acid at pH 3.0 provided minimal benefit above the pH-matched baseline (Figure S18a). Although biomass loading was not varied, Ni solubilization from shoots supplemented with different Ni concentrations followed a single log-linear relationship with nominal reactant-to-Ni ratio across the tested datasets (Table S26).

Together, these results showed that acid- and chelator-assisted extraction could dissolve metals from *Phytolacca* shoot tissues under bioleaching-relevant incubation conditions, but the extent varied across the *Phytolacca* species, target metal, and reactant chemistries examined.

### REE repartitioning from *P. americana* shoots may reflect equilibrium-like behavior

The preceding extraction experiments reinforced that acids and chelators could solubilize metals from plants.^25,26^ For conditions which reached apparent target-metal solubilization plateaus by the 7 d endpoint (Figure S12), condition-specific (surface-area-normalized) *K_d,i,app_*for the dissolution reaction only began to significantly decrease at higher reactant to target metal ratios (Figure S19). However, these data did not determine whether REE solubilization reflected irreversible *Phytolacca* disruption or a reversible shift in *Phytolacca*–solution partitioning.

To assess whether REE release occurred with a concomitant disruption of the plant tissue, apparent double-stranded DNA (dsDNA) signals were measured before and after leaching. Endpoint H_2_SO_4_ leachates contained little measurable SYBR Green signal relative to the initial leaching baseline under the tested conditions (Figure S20). In contrast, intentionally disrupted *P. americana* shoots produced elevated fluorescence relative to no-tissue controls (Figure S21), and autoclaved and non-autoclaved shoots incubated in sterile H_2_O generated time-dependent fluorescence increases relative to no-shoot controls (Figure S22).

Further tests explored whether REE release could be achieved under aqueous conditions without strong acidification, chelator addition, or matrix disruption. Ce/Dy-enriched *P. americana* shoots were incubated at pH 4.5 under a common 0.1 M KNO_3_ background electrolyte while added electrolyte composition was varied (Figure 3a). Under these conditions, SO_4_^2−^-containing electrolytes transiently increased REE solubilization relative to the NO_3_^−^-only condition, whereas release profiles approached apparent plateaus over the tested incubation period (Table S27). These electrolyte-supplemented conditions increased the extent of REE solubilization when separately performed for 7 d (Figure S23). The H_2_SO_4_ + KNO_3_ condition also increased SYBR Green signal relative to both final no-biomass controls and the initial reactant solution before biomass addition; this increase was not observed for H_2_SO_4_ alone under non-identical test conditions, was restricted to select pH 4.5 leaching conditions, and was not observed across all acidic or chelator-assisted extractions (Figure S20).

**Figure 3:**
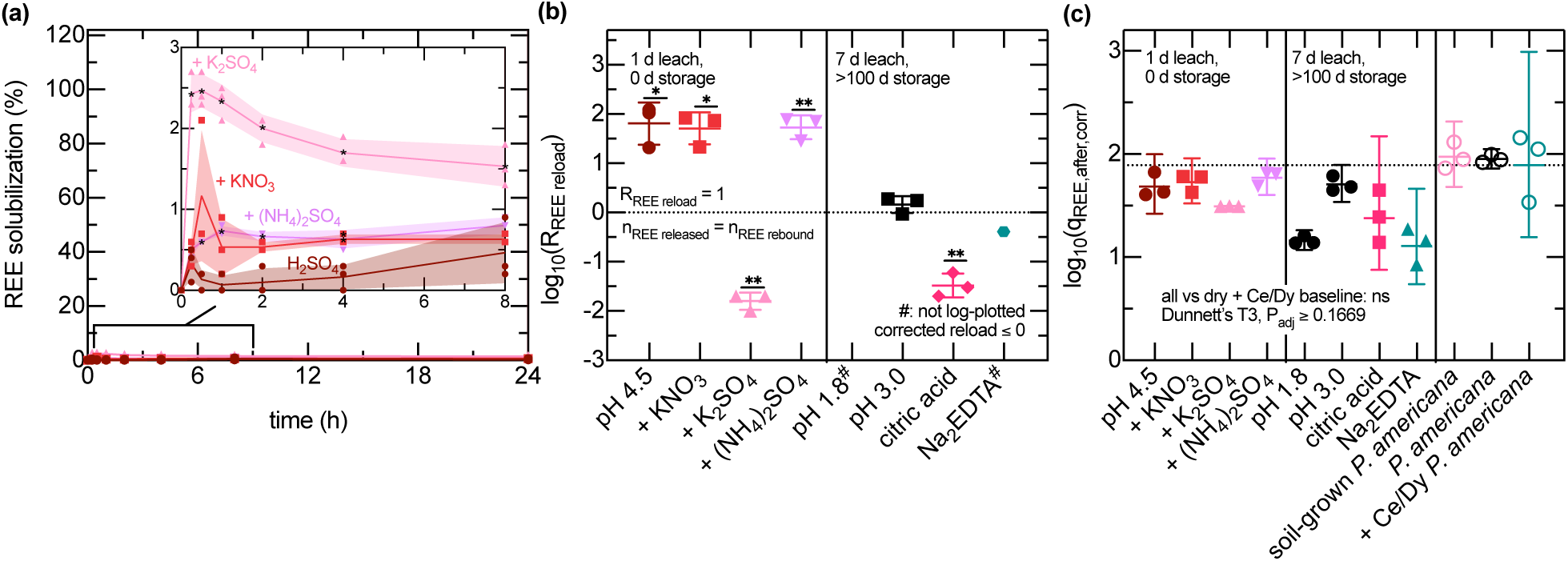
REE release and rebinding by *P. americana* are extraction-pathway dependent. (a) Time-course measurements of REE solubilization from dried Ce/Dy-enriched *P. americana* shoots at pH 4.5 under a common 0.1 M KNO_3_ background electrolyte. Treatments were the H_2_SO_4_ pH 4.5 baseline or the same baseline supplemented with K_2_SO_4_ (0.1 M), (NH_4_)_2_SO_4_ (0.1 M), or additional KNO_3_ (0.2 M). Asterisks indicate significance relative to the H_2_SO_4_ baseline at the corresponding time point. (b) *R*_reload_ for post-leached solids after standardized Ce/Dy rebinding, plotted as log_10_(*R*_reload_). The vertical divider separates residues reloaded immediately after 1 d extraction from residues stored under static benchtop conditions for a while after the 7 d extraction before rebinding. Asterisks indicate significant deviation from log_10_(*R*_reload_) = 0. (c) *q*_after,corr_ after standardized Ce/Dy rebinding, plotted after log_10_ transformation. The dotted line indicates the dry Ce/Dy-enriched *P. americana* baseline.

The post-leached *P. americana* tissues from selected tests were recovered and re-exposed to a soluble Ce/Dy solution to assess whether the plant-associated REE pool exhibited reversible partitioning between plant solids and bulk solution. For the 1 d background-electrolyte supplemented leaching conditions, *R*_reload_ depended on the prior extraction chemistry (Figure 3b). The K_2_SO_4_-leached tissues had lower *R*_reload_ than the matched H_2_SO_4_ baseline, whereas the KNO_3_ and (NH_4_)_2_SO_4_ conditions were not statistically resolved from the baseline (Table S28). For the tissues from 7 d leaching conditions, which differed from the 1 d set in leaching duration, storage time before reloading, and reactant chemistry, *R*_reload_ was not resolved from unity after H_2_SO_4_-leaching at pH 3.0, whereas the tissues leached at pH 1.8 produced non-positive *R*_reload_ values that could not be plotted (Figure 3b). The citric acid- and Na_2_EDTA-leached conditions were consistent with incomplete reload, though the latter was interpreted descriptively because only one *R*_reload_ value was positive (Table S28). Similar outcomes were observed when some of these conditions were performed in parallel under a common 7 d duration (Figure S24a–f).

Additionally, *q*_after,corr_ was used to assess whether prior leaching depleted the apparent REE-binding capacity of the plant tissues (Figure 3c). Across the tested post-leached conditions, the final REE loading of post-leached REE-enriched *P. americana* shoots was not statistically resolved from corresponding non-leached controls, although median normalized values were below unity for all post-leached conditions (Table S29).

With *n* = 3 per condition, the exact two-tailed Wilcoxon tests could not yield *P <* 0.05, so these results do not establish preservation of endpoint REE-binding capacity. Similar trends were observed across consecutive binding, unbinding, and rebinding cycles for the tests performed in parallel under a common 7 d duration (Figure S24g).

FTIR was used to assess whether the reloaded tissues exhibited comparable spectral features to the non-leached Ce/Dy-enriched *P. americana* shoots. FTIR spectra of untreated, Ce/Dy-enriched, post-leached, and post-reloaded *P. americana* residues showed conserved features across the O–H, C–H, C=O/COO*^−^*/amide, and PO_4_^3–^ /P–O/C–O stretching regions (Figure S25). In particular, the major 1200–900 cm*^−^*^1^ PO_4_^3–^ /P– O/C–O envelope was retained after H_2_O, pH 3.0 H_2_SO_4_, (NH_4_)_2_SO_4_, and K_2_SO_4_ treatments, indicating that REE release and reload behavior were not accompanied by selective depletion of a dominant FTIR-detectable phosphate-/carbohydrate-associated phase. Residues leached under higher H_2_SO_4_ concentrations could not be analyzed equivalently, precluding assessment of whether more acidic extraction conditions produced stronger matrix alteration than the conditions shown in Figure 3c. Attempts to lyophilize shoots leached with 0.1–3 M H_2_SO_4_ produced viscous purple extracts (Figure S26). After 100-fold dilution in H_2_O, the extracts exhibited broad UV absorbance that was absent from H_2_O and H_2_SO_4_ controls and proportional to the acidity of the leaching solution (Figure S27).

Together, these results showed that REE depletion and rebinding were dependent on solution chemistry beyond pH, and the pattern of similarities in the REE-binding capacity and vibrational features between the post-leached tissues and the non-leached Ce/Dy-enriched *P. americana* was consistent with apparent equilibrium-like REE partitioning.

### *A. ferrooxidans* could concentrate soluble REEs relative to enriched *Phytolacca* under acidic conditions

Having explored the metal release and repartitioning dynamics of *Phytolacca*, the feasibility of using *A. ferrooxidans* to bind and concentrate REEs from acidic solutions was evaluated. Initial tests were performed with an initial *A. ferrooxidans* loading of OD_600_ = 1.0. Time-course measurements reached an apparent REE binding maximum for both an equimolar Ce_2_/Dy_2_/Nd_2_/Tb_2_-(SO_4_)_3_ simulant and an isohydric *P. acinosa* H_2_SO_4_ leachate in about 2 h at pH 1.8 (Figure S28a); subsequent binding tests were therefore performed for 2 h or longer. The recovery of soluble REEs in these solutions was low (Figure S28b), and the apparent HREE/LREE binding selectivity was qualitatively different across the two conditions at the endpoint (Figure S28e).

Additional tests indicated that REE binding capacity by *A. ferrooxidans* was inversely proportional to the initial cell density loaded in equimolar Ce_2_/Dy_2_/Nd_2_/Tb_2_-(SO_4_)_3_ simulants (Figure S29) and Tb_2_(SO_4_)_3_-only simulants (Figure S30). Subsequent tests were therefore mostly performed using OD_600_ = 0.1.

Binding isotherms in the Tb_2_(SO_4_)_3_ simulant showed that, at high bulk Tb^3+^ concentrations, *A. ferrooxidans* achieved a higher cell-associated Tb concentration than the REE concentration measured in Ce/Dy-enriched *P. americana* shoots (Figure 4a). This inferred Tb binding capacity and its inversely proportional relationship with initial *A. ferrooxidans* cell density agreed qualitatively but not quantitatively with measured cell-associated Tb in recovered cell pellets (Figure 4b). At the highest Tb_2_(SO_4_)_3_ concentration tested in Figure 4, the fraction of inferred prewash Tb retained after washing significantly increased between 2 and 24 h, while no significant difference in inferred binding capacity was detected (Table S30; Figure S31). After 24 h of binding under the same conditions, no significant difference was detected in the fraction of inferred prewash Tb retained after a single 5 min EDTA wash compared with the standard two-stage EDTA washing protocol (Table S31; Figure S32). For abiotic controls performed in parallel, added phosphate promoted Tb depletion from solution after 24 h (Figure S33a), and the EDTA wash protocol quantitatively redissolved this depleted Tb (Figure S33b). Cells with and without Tb exposure during incubation looked similar after EDTA or wash washing (Figure S34), and the intensity of the Tb emission spectra of cells incubated with Tb after EDTA washing (Figure S35) was qualitatively similar to controls which had never been exposed to Tb (Figure S36). The intensity of the Tb emission spectra of cells incubated with Tb was qualitatively lower than after EDTA washing (Figure S37) than water washing (Figure S38).

**Figure 4:**
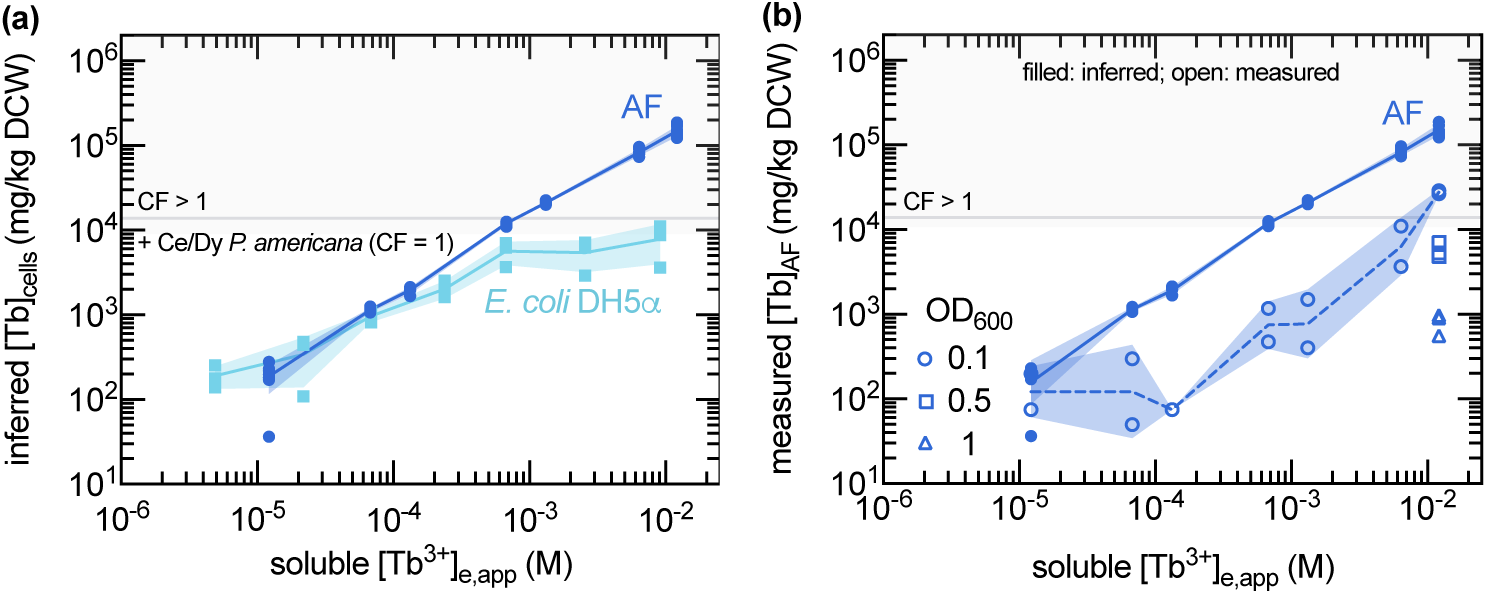
*A. ferrooxidans* concentrated Tb more strongly than *E. coli* DH5*α* under acidic binding conditions. (a) Inferred Tb associated with *A. ferrooxidans* (n = 8) and *E. coli* DH5*α* (n = 3) as a function of soluble Tb^3+^ in AFM1 BS at pH 1.8 (OD_600_ = 0.1, 30 °C, 150 rpm, 2 h). (b) Inferred and measured Tb associated with *A. ferrooxidans* for the data plotted in (a) and some other tests (*n* = 2 samples prepared from 4 combined biological replicates because of sample mass limitations. The gray horizontal reference line indicates the Tb-equivalent loading corresponding to a concentration factor (CF) of 1 relative to the Ce/Dy-enriched *P. americana* shoots.

Under non-identical conditions, *A. ferrooxidans* retained high Tb-association and citrate-dissociation Tb capacities across three sorption–elution cycles (Figure S39a), with no significant effects of cycle, recovery step, or their interaction on log_10_-transformed inferred Tb values (Table S32). Heat-pretreated *A. ferrooxidans* had higher inferred Tb binding capacity and citrate-eluted Tb capacity relative to the untreated controls across three sorption–elution cycles (Table S33; Figure S39b-c). UV-C-pretreated *A. ferrooxidans* had insignificantly higher inferred Tb binding capacity than controls, and the citrate-eluted Tb capacity was significantly higher during the last two elution cycles (Table S33; Figure S39b-c). In separate experiments, these heat and UV-C pretreatments increased fluorescence intensity from indicators which are responsive to cell permeability and membrane-potential polarization, but not lipid-derived free radicals (Table S34; Figure S40a-c). Both pretreatments also reduced the apparent rate of Fe^2+^ and brown S^0^ biooxidation (Figure S40d-e) and endpoint planktonic cell density measurements (Figure S40f).

*A. ferrooxidans* had high REE binding capacity under the tested acidic conditions. For the Tb_2_(SO_4_)_3_ simulant at pH 1.8, the Tb binding capacity by *E. coli* DH5*α* was lower than that of *A. ferrooxidans* (Figure 4a). *E. coli* DH5*α* achieved a higher maximum inferred Tb recovery than *A. ferrooxidans* (Figure S41a), though the condition-specific *K*_d,Tb,app_(Figure S41b–c) and apparent lower-bound selectivity for Tb over impurities estimate (Figure S41d) became lower than that for *A. ferrooxidans* as the apparent equilibrium concentration of soluble Tb increased. Under different, less acidic conditions, Tb binding capacity by *E. coli* DH5*α* was similar to that of *A. ferrooxidans* under acidic conditions at high bulk concentrations and higher at lower Tb^3+^ concentrations (Figure S42).

To test whether prior Tb^3+^ exposure could improve inferred REE binding capacity by *A. ferrooxidans*,^46^ F2S growth medium was supplemented with 15 µM Tb_2_(SO_4_)_3_, matching the soluble REE concentration achieved from Ce/Dy/Nd/Tb-enriched *P. acinosa* shoots after H_2_SO_4_ leaching at pH 1.8. Tb^3+^ supplementation in F2S growth medium did not inhibit Fe^2+^ biooxidation or planktonic cell growth by *A. ferrooxidans* (Figure S43a–b). *A. ferrooxidans* grown in Tb-supplemented F2S medium produced higher Tb concentrations in recovered cell pellets than baseline F2S controls; however, this growth-associated Tb carryover was lower than the value measured after exposure to 12 mM Tb^3+^ (Table S35; Figure S43c). Tb_2_(SO_4_)_3_ supplementation in F2S growth medium did not increase REE binding capacity by *A. ferrooxidans* in Tb_2_(SO_4_)_3_ or equimolar Ce_2_/Dy_2_/Nd_2_/Tb_2_-(SO_4_)_3_ simulants at pH 1.8, though cells grown under this condition notably showed REE binding as low as pH 1.0 (Table S36; Figure S43d).

Collectively, these simulant-solution binding experiments showed that *A. ferrooxidans* could bind and sometimes concentrate REEs in acidic leachates relative to REE concentrations in Ce/Dy-enriched *P. americana* shoots. However, the tests did not determine whether metals released from plant tissues would remain equally available in plant-derived leachates. This distinction was important because adding *Phytolacca* shoots altered solution chemistry relative to initial conditions and no-biomass controls during H_2_O and H_2_SO_4_ leaching, including apparent metal availability (Figure 2a–b, Figure S18), pH and ORP (Tables S37– S39; Figure S22a–b, Figure S44), buffering capacity (Figures S45–S46), apparent dsDNA signal measured by SYBR Green fluorescence (Figure S20, Figure S22d), and SO_4_^2−^ concentration (Figure S22c). To assess whether leachates contained plant-derived solution components that could alter metal speciation, some 0.22 µm-filtered H_2_O and H_2_SO_4_ leachates were fractionated by 3 kDa centrifugal ultrafiltration before additional solution-property measurements. Across the ultrafiltration tests, mass retention above the 3 kDa cutoff generally remained near the expected 50% concentration-factor value (Table S40; Figure S47a), whereas the UV-active plant-derived material proxy (Table S41; Figure S47b) and Ni or REEs (Table S42; Figure S47c) did not always partition similarly. Simple and additive regression models suggested Ni retention from *P. acinosa* leachates was largely associated with mass retention, while AUC retention provided some explanatory value in the combined model; adding pH provided a minor benefit but was an insignificant predictor when mass retention and AUC retention were both included (Table S43). In contrast, REE retention from REE-enriched *P. americana* leachates was mostly associated with leachate pH; pH was a significant predictor in individual and additive models including mass and/or AUC retention, whereas mass and AUC retention were insignificant after pH was included (Table S44).

Although these solution-phase measurements did not identify specific metal-binding species, they showed that exposure to *Phytolacca* shoots altered solution properties that could change metal partitioning from the solution to *A. ferrooxidans* relative to REE-SO_4_ simulants. Therefore, further tests probed whether leachate matrix chemistry measurably altered REE binding with *A. ferrooxidans* or whether association was better explained by bulk loading variables. REE binding from an H_2_SO_4_–*P. acinosa* leachate occurred on a similar time scale as binding from an equimolar Ce_2_/Dy_2_/Nd_2_/Tb_2_-(SO_4_)_3_ simulant with a higher REE concentration under isohydric conditions (Figure S28a). To assess differences in binding capacity, endpoint binding data were analyzed using class-adjusted log–log multiple linear regression with solution class, initial REE concentration, *A. ferrooxidans* cell density, and their interaction as predictors. Across these tests, REE binding capacity was primarily associated with initial soluble REE concentration and *A. ferrooxidans* cell-density loading, whereas solution-class terms had a comparatively smaller but resolved offset relative to the Tb^3+^-only reference and OD_600_ contributed only a small negative association; plant-derived leachates did not show resolved matrix-specific suppression of REE binding relative to Tb_2_(SO_4_)_3_ or equimolar Ce_2_/Dy_2_/Nd_2_/Tb_2_-(SO_4_)_3_ simulants under the tested acidic conditions (Table S45).

Together, these results suggested that *Phytolacca* leachates contained unidentified plant-derived components that altered bulk solution properties and metal partitioning, but did not always prevent *A. ferrooxidans* from binding soluble REEs under acidic conditions.

### *A. ferrooxidans* simultaneously dissolved and recovered REEs from *Phytolacca*

After demonstrating that H_2_SO_4_ could release REEs from *Phytolacca* tissues and that *A. ferrooxidans* could concentrate REEs from acidic solutions relative to enriched *P. americana* shoots, these functions were tested in an integrated biological extraction and concentration process. Fe-based medium was not used beyond the tolerance screening because it showed strong inhibition at relevant *P. americana* tissue loadings (Figure S48), introduced a large Fe impurity pool, and Fe^3+^ did not improve REE dissolution from *P. americana* (Table S22).

REE solubilization from Ce/Dy-enriched *P. americana* shoots was tested under conditions that varied S^0^ dosage, initial *A. ferrooxidans* density, and *P. americana* shoot loading (Figure S49). At fixed inoculum density and *P. americana* loading, increasing S^0^ dosage increased REE solubilization relative to the no-S^0^ control until an apparent saturation level was reached (Figure S49a), consistent with greater S^0^ biooxidation by *A. ferrooxidans*, as indicated by lower pH, more oxidizing conditions, and higher conductivity relative to the no-S^0^ control (Figure S49f,k,p). At 1% (w/v) shoot loading, increasing inoculum density did not monotonically increase REE solubilization (Figure S49b), even though higher inoculum density generated stronger acidification and conductivity changes consistent with greater S^0^ biooxidation (Figure S49g,q). At the lower inoculum density of OD_600_ = 0.01, REE solubilization was greatest at intermediate shoot loading rather than increasing monotonically with *P. americana* loading (Figure S49c). In contrast, at the higher inoculum density of OD_600_ = 0.1, REE solubilization decreased as *P. americana* loading increased (Figure S49d), despite time-course measurements indicating rapid and sustained S^0^ biooxidation characteristics (Figure S49i,n,s). At fixed *P. americana* loading of 1% (w/v), REE solubilization from cultures which were not inoculated with *A. ferrooxidans* increased as the initial pH decreased in the presence and absence of S^0^ (Figure S49e). These abiotic pH, solution potential, conductivity, and apparent planktonic OD_600_ time-course measurements (Figure S49j,o,t,y) were not consistent with the corresponding data from controls that lacked the *P. americana* shoots (Figure S50).

REE solubilization across these bioleaching experiments was then evaluated against bulk pH to examine whether treatment-dependent differences reflected additional controls on the bioleaching outcomes. When time-course data were replotted as REE solubilization versus pH (Figure 5a), REE solubilization generally increased as pH decreased, similar to proton-dependent release observed during chemical leaching (Figure 2a, Figure S14). The time-dependent, surface-area-normalized distribution ratios also exhibited condition-dependent relationships with the reactant-to-REE molar ratio (Figure 5b), which remained after removing the coupled mathematical dependency on plant loading (Figure 5c). Endpoint regression analysis supported this trend, as lower endpoint pH was significantly associated with greater REE solubilization in a pH-only model; adding treatment identity substantially increased model explanatory power, and treatment identity remained significant after accounting for endpoint pH, whereas pH was insignificant once treatment identity was included (Table S46).

**Figure 5:**
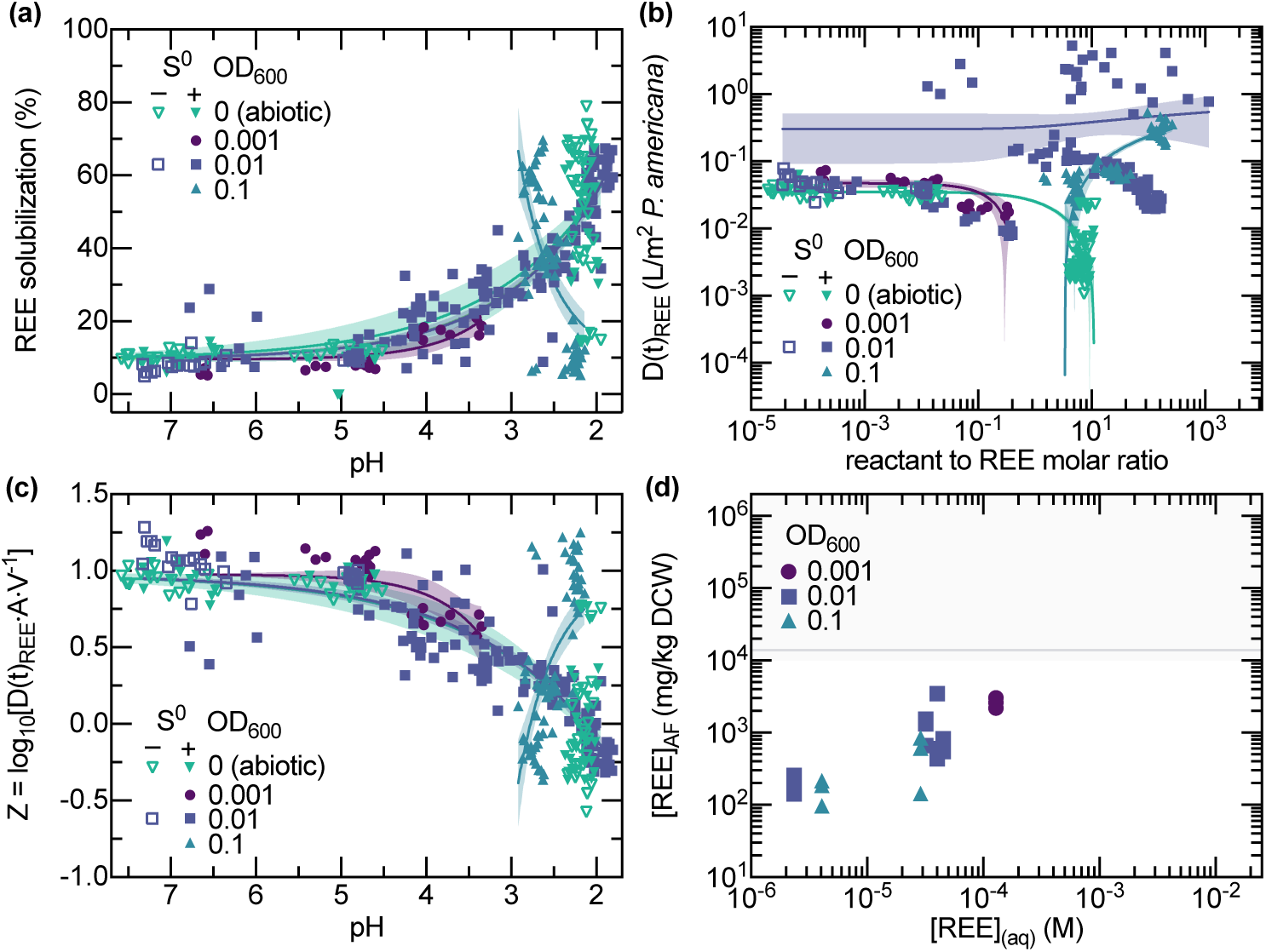
Sulfur biooxidation by *A. ferrooxidans* released REEs from *P. americana*, but REE solubilization reflected coupled reaction and partitioning behavior. (a) Combined time-course REE solubilization data from experiments varying S^0^ dosage, initial inoculum density, and *P. americana* shoot loading, plotted against measured pH from data series shown in Figure S49. Curves were fitted using REE solubilization = REE solubilization_0_ + *A ·* 10*^−k·^*^pH^, and the baseline REE solubilization was constrained to the water-labile fraction of 9.54%. (b) Time-dependent solid–liquid distribution ratios of REEs versus reactant-to-REE molar ratio. Solid lines show fitted logarithmic models [*D*(*t*)_REE_ = *a ·* log(*X* + 1) + *b*] and shaded bands show 95% CIs, where *X* is the estimated reactant-to-REE molar ratio (Equations S102–S104). (c) Logarithmic ratio of nonsolubilized to solubilized REE inventory versus measured pH, which removes the algebraic dependency of plant loading from the solid–liquid distribution ratios. *Z* = 0 corresponds to equal nonsolubilized and solubilized inventories. *Z <* 0 indicates more REEs were solubilized, and *Z >* 0 more REEs were nonsolubilized. Solid lines show fitted logarithmic models [*Z* = 0.977 + *A ·* 10*^−k·^*^pH^] and shaded bands show 95% CIs. (d) Endpoint REE concentrations of recovered *A. ferrooxidans* plotted against the bulk aqueous REE concentration for the bioleaching cultures. The gray horizontal reference line indicates the Tb-equivalent loading corresponding to a concentration factor (CF) of 1 relative to the Ce/Dy-enriched *P. americana* shoots. Colors and symbol shapes distinguish initial inoculum densities. Open and filled symbols indicate the absence and presence of elemental sulfur, respectively.

Mixed-effects analysis of the full time-course data further supported that pH did not fully explain the treatment-dependent REE solubilization behavior. REE solubilization, pH, solution potential, and conductivity each showed time, treatment, and time-by-treatment effects, indicating that endpoint differences were not simply final-time-point offsets (Table S47). In secondary reduced endpoint models, which were interpreted descriptively because treatment variables were coupled across the experimental design, initial inoculum density was negatively associated with endpoint REE solubilization when modeled with endpoint pH, whereas sulfur dosage and log_10_-transformed *P. americana* loading were not significant in their respective pH-adjusted models (Table S46).

The matched inoculum-density series showed that inoculum condition contributed to treatment-dependent trajectories. At fixed S^0^ and *P. americana* loading, mixed-effects analysis showed time, inoculum-group, and time-by-inoculum-group effects for REE solubilization, pH, ORP, and conductivity (Table S48). Similarly, cumulative pH decrease did not explain REE release AUC, whereas adding categorical inoculum group produced a significant model for replicate-level summary analyses (Table S49). Inoculum group also explained variation in REE-versus-pH slope and maximum REE solubilization, whereas minimum pH alone did not. Because inoculum group was collinear with pH-decrease AUC and minimum pH in the combined models, these analyses suggested that inoculum-dependent REE release coincided with inoculum-dependent acidification and solution-chemistry trajectories rather than supporting a pH-independent inoculum effect.

Since the removal of aqueous REE attributed to the brown S^0^ seemed lower than the decrease in REE solubilization observed in the bioleaching experiments (Figure S51), subsequent tests evaluated whether lower REE solubilization at higher inoculum density reflected partitioning of soluble REEs from the bulk solution to *A. ferrooxidans*. Across the arbitrary subset of bioleaching conditions where the *A. ferrooxidans*-associated REE concentrations were measured, these measurements generally increased with bulk solution REE concentration and decreased with increasing inoculum density (Figure 5d). Inclusion of these *A. ferrooxidans*-associated REE concentrations improved the closure of the mass balances for the underlying bioleaching tests (Figure S52). In the absence of *P. americana*, the addition of 100 µM Ce^3+^, Dy^3+^, and some other cations to the growth medium with a lower initial pH of 2.0 increased the rate and extent of planktonic *A. ferrooxidans* growth and net acidification relative to the baseline controls (Figure S53). In separate cultures which contained *P. americana*, similarly, the presence of soluble Ce and Dy reduced the plant-associated delay in acidification when solubilized from the Ce/Dy-enriched *P. americana* or when added to the growth media with non-supplemented control shoots at concentrations achieved during bioleaching experiments with (Figure S54a–c). The concomitant decrease in the concentration of aqueous Ce and Dy for the conditions with a higher initial inoculum density was not observed in the corresponding abiotic or lower initial inoculum density controls (Figure S54d–i).

A two-stage centrifugation workflow was then evaluated using analogous *P. acinosa* bioleaching cultures to test whether these phases could be operationally separated. This approach separated cultures into REE-depleted *P. acinosa* solids, putatively REE-enriched *A. ferrooxidans*, and aqueous bioleachate co-product streams (Figure S55).

Together, these results support a model in which REE solubilization during bioleaching reflects a balance between REE release from *Phytolacca* tissues and partitioning of soluble REEs to the *A. ferrooxidans* phase.

## Discussion

Biophytometallurgy is an aqueous strategy to recover target metals from metal-enriched plant biomass without direct thermochemical conversion to ash. In this concept, the plant provides an initial biological concentration step during phytomining, and then microorganisms generate reactant(s) that promote metal release via solid–liquid extraction during subsequent biophytometallurgical processing of the harvested plant. This study used biophytometallurgy primarily within a Biology-as-a-Tool (BaaT) framework to examine whether equilibrium shifting was feasible during solid—liquid extraction to indirectly provide insight into the nature of metal—plant partitioning. We hypothesized that if REE release from the plant was governed by mass action, and if the extent of reaction was limited by the product concentration within this deliberately constrained model system, then the microorganism could act as a living dual-function material that promotes the removal of REEs from the plant by removing soluble REEs from the aqueous phase.

*Phytolacca* was selected as a hypothetical phytomining system because the genus includes fast-growing, high-biomass species^53^ and species which can accumulate and fractionate REEs.^39^ The genus includes species native to all continents except Antarctica, Europe, and Oceania.^54^ Given Phytolacca’s unsuitability as a food crop for humans, using species from this genus for phytomining offers the advantage of preventing the in-corporation of phytomined metals into human diets.^55^ The properties of the tested *Phytolacca* materials and subsequent results should not be interpreted as direct proxies for field performance or hyperaccumulation classification.^56^ The hydroponic cultivation system and associated growth conditions used in this study (Text S5) should be considered when interpreting plant physiology (Table S12), and metal concentrations and tissue-partitioning (Table S4). The aggressive growth and potential for dispersal of *Phytolacca* during harvesting and handling^57^ should also be considered prior to field demonstrations. Ni, a more-established phytomining target, was included as a comparator to contextualize the reactant demands of some of the tested solid–liquid extraction approaches. However, comparisons between Ni- and REE-enriched materials also reflect differences in enrichment conditions, including phosphate omission during REE enrichment and, for some experiments, plant species (Text S5). We were curious to see if *Phytolacca* would permit a high maximum solid loadings during bioleaching^58^ if hydration-induced softening of plant tissues^59^ reduced localized contact stresses during particle collisions. This assumes drying-induced hornification would not notably restrict reswelling.^60^

*A. ferrooxidans* was selected because it can oxidize inorganic S^0^ to release protons,^44^ and it has a high acid and REE tolerance.^45^ The putatively high REE-binding capacity^46^ also made it an attractive candidate for recovering dissolved REEs into a microbial phase under acidic conditions, possibly up to a higher metal grade than the plant precursor. *A. ferrooxidans* were not expected to utilize the plant biomass as an energy or carbon source.^44^ In fact, plant-derived low-molecular-weight organic acids were expected to inhibit its metabolism of supplemented energy sources.^61,62^ Growth-permissive conditions were feasible at higher initial cell-to-plant biomass ratios (Figure S54). Ce and Dy mixtures added to or solubilized from *P. americana* also appeared to reduce the apparent inhibition of the bacteria from the plant (Figure S54).^63^ However, plant-loading-dependent changes in biooxidation and acidification (Figure S48) did not resolve or decouple the contributions of chemical inhibition, biomass-associated buffering, and physical constraints, and a physical upper limit on solids loading was not established.

The feasibility of this biophytometallurgy concept depends on two coupled design constraints. First, the plant must contain a chemically accessible compartment(s) within the overall metal pool. Second, the microorganism must remain sufficiently active in the plant-supplemented acidic or ligand-assisted solution to contribute to both metal release and recovery. Four process design questions were raised, and efforts made to answer them, in this project.

The first question asked was whether *Phytolacca* behaved like a refractory REE-phosphate mineral phase^34,64^ found within Earth’s crust. Asking whether this plant was a rock from a mineralogy perspective seemed useful for curating the tested (bio)phytometallurgy reactant and concentration conditions, and answering this question from a cutoff grade perspective would have been important during technoeconomic assessment if our goal was to target higher technology readiness levels. Elemental analysis indicated that the Ce/Dy-enriched *P. americana* shoots were more concentrated in REEs than the average rock (Table S4),^65^ but did not provide an answer as to whether the plant behaved like a mineral. Given the elemental compositions measured, the entire REE pool in the tested *P. americana* shoots could not be bound in REE-orthophosphate compartments with an ideal 1:1 molar stoichiometry (Table S4; Figure S5). Some portion, however, could have been, and REE orthophosphate (nanocrystalline-)phytomineralization has been speculated to occur in *Phytolacca*.^31^ If monazite phytomineralization were the predominant REE compartment in the plant, and if it behaved like crystalline geological monazite, the (presented) biophytometallurgical concept would be rendered pointless. Under this scenario, organic acids may dissolve more REEs than mineral acids. However, a biologically mediated solid–liquid extraction process would be slow,^33^ and likely require more demanding thermal and chemical conditions^66,67^ than could be generated or tolerated by heterotrophs or chemolithoautotrophs. The *Phytolacca* shoots did not look (Figure S56), diffract (Figure 1d, Figure S8),^51,52^ vibrate (Figure 1e, Figure S9),^49,50^ or dissolve (Figure 2, Figure 3a, Figure S12a–c, Figure S14a–c, Figure S18b, Figure S23a–b)^31,34,64,68^ as would be expected from a predominant crystalline geological monazite-like REE compartment, before or after calcination. It is possible that these *Phytolacca* shoots contained a (nanocrystalline-)phytomineralized monazite-like REE-phase, which might not be detectable by these bulk characterization techniques.^31^ Additionally, the short-term hydroponic exposure to high dissolved REE concentrations under phosphate-depleted conditions used to generate these *Phytolacca* shoots (Text S5) could have biased the metal pools toward exchangeable forms and limited the formation of phytomineralization, which might require longer exposure times or greater phosphate availability. When plant-mediated REE mineralization were discovered in some fern species, pinna were digested using dilute acetic acid at ambient temperature while stirring, and the dissolved REEs from this assay were assigned as phosphate-bound fraction.^31^ It is possible^69–71^ these results imply that the plant-formed (nanocrystalline-)monazite was more dissoluble by an organic acid under ambient temperature conditions^31^ that would be expected for crystalline geological monazite.^72^ This led us to speculate that even if (nanocrystalline-)phytomineralization were the predominant REE compartment in these *Phytolacca* materials, we could examine the reactions that govern metal release under lower reactant concentration and temperature conditions than had been used in earlier studies of REE phytometallurgy.

The second question asked under what conditions were target metal compartments chemically accessible in *Phytolacca*. Under the tested conditions, Fe^3+^ did not appear to enhance REE dissolution from the Ce/Dy-enriched *Phytolacca* shoots through redoxolysis (Table S22). A very minor portion of the REE pool was released through cation exchange (Figure 3a), but a limited set of shared dissolution characteristics with ion-adsorption clays^73^ may have broader geochemical implications if this behavior was observed using field-grown hyperaccumulator biomass (Table S20; Figure S17).^74,75^ A small fraction of the REEs appeared weakly retained or exchangeable under mild temperature, mixing, and acidity;^34,64,68^ the Ni-enriched *P. acinosa* materials released a larger fraction of their target-metal inventory in H_2_O, but their apparent Ni solubilization subsequently decreased during extended storage (Figures S12–S13). In contrast, acid promoted REE solubilization from a larger REE compartment (Figure S14), and the extent did not seem grossly higher than what was reported in previous studies, some of which utilized more environmentally representative biomass (Texts S23–S24; Figure S57). REE and Ni solubilization increased with nominal H_2_SO_4_-to-targetmetal ratios across the tested H_2_SO_4_ concentrations regardless of whether metal loading, acid concentration, or biomass loading was varied (Table S18, Table S26). The Ni comparison therefore extended the assessment of reactant dosage to plant materials with different target-metal grades. These behaviors are expected for proton-dependent processes governed by mass action.^42^ Additional results were consistent with REE solubilization being driven by a proton-dependent process under mildly acidic conditions, which was putatively ligand protonation (Figure 2b, Figure 3b–c, Figure S20, Figure S25, Figure S47). Chelation promoted Ni and REE solubilization from *Phytolacca* beyond the effects of acidity (Figure 2b, Figure S18). The mechanism, which could have been ligand competition (Figure 3b–c, Figure S20, Figure S25),^72,76^ depended on ligand identity, availability, and concentration (Figure 2c).^25,26^ The overall extraction and rebinding data are consistent with REE dissolution being governed by apparent reversible equilibria under mildly acidic conditions. The rebinding experiments cannot distinguish whether the REEs returned to the original binding site (Figure 3b–c, Figure S23, Figure S24) and so these data do not resolve the microscopic mechanism governing the reversibly exchanging REE compartment, nor the size of this fraction, within the overall REE pool.

The third question raised was whether *A. ferrooxidans* could be used to concentrate, or biobeneficiate, REEs dissolved from plants. The microbial binding capacity for REEs increased with bulk concentration and decreased with cell loading (Figure 4, Figure S29a–b). These trends are expected for biosorption.^42^ The evidence provided of intracellular accumulation (Text S25; Equations S105–S122) could indicate that REE uptake by the bacteria^77,78^ contributed to the observed metal partitioning (Figures S53–S54). The physiological basis for the ability of *A. ferrooxidans* to bind and putatively uptake soluble REEs under acidic conditions is unexpected and unknown, and warrants further study. *A. ferrooxidans* concentrated soluble REEs from acidic simulants relative to the REE-enriched *Phytolacca* material, without a requirement for genetic modification (Figure 4).^46^ In contrast to earlier work from the Banta lab,^46^ Tb_2_(SO_4_)_3_-supplementation at the onset of growth neither inhibited the growth nor the metabolism of *A. ferrooxidans* (Figure S43a–b).^45^ The previous claim of inhibition was likely confounded by the use TbCl_3_^46^ as there was minimal difference in the apparent inhibition of Fe^2+^ biooxidation attributed to TbCl_3_ and NaCl across matched Cl^−^ concentrations ranging from 10*^−^*^8^ *−* 10*^−^*^1^ M (unpublished, Banta lab). However, there were no obvious reason why the addition of Tb^3+^ through TbCl_3_ (data not shown) or Tb_2_(SO_4_)_3_ (Figure S43) to the F2S medium at the onset of growth did not increase the inferred REE binding capacity under the reported experimental conditions.^46^ The *Phytolacca* changed some properties of the leaching solution (Figure 2a–b; Tables S37–S39; Figure S18, Figure S20, Figure S22, Figures S45–S47) which made it unclear if *A. ferrooxidans* would maintain the ability to bind and/or uptake metals under such conditions. Ni retention during ultrafiltration largely tracked retained liquid mass, whereas REE retention was more strongly associated with leachate pH (Tables S43–S44; Figure S47). The different operational partitioning behavior of Ni and REE in these solutions is consistent with the presence of unidentified colloidal species^79^ and soluble complexants,^80,81^ respectively. Through separate binding experiments, it was shown that the presence of these putative complexants did not inherently eliminate the ability of *A. ferrooxidans* to bind REEs (Figure S28a).

The final question asked was whether bioleaching and biorecovery could be coupled in a simultaneous release–repartitioning process. No common monotonic trend of REE solubilization as a function of pH was observed across the bioleaching experiments (Figure 5a), consistent with *A. ferrooxidans* acting as both an acid-generating source and a competing REE-associated sink (Figure S54). Although full mass balance closure was not routinely measured or achieved (Figure S52a–b), the measured aqueous REE concentration putatively reflects the net outcome of release from *Phytolacca*-associated compartments, accumulation in bulk bioleachate, and repartitioning across the separable *Phytolacca* and *A. ferrooxidans* solid phases (Figure 5d, Figure S55, Figure S52c). Consequently, unlike conventional hydrometallurgy, in which extraction is typically defined by the amount of soluble target species within the aqueous phase, techno-economic assessment of scaled biophytometallurgical operations would require accounting for REE recovery and repartitioning among the bioleachate, microbial concentrate, and residual plant solids.

In sum, a BaaT framework was used to explore the dynamics controlling metal redistribution across interfaces within a multi-phase biophytometallurgy system to test speculative hypotheses regarding plant biochemistry. Granted, adding a living component to this phytometallurgy system introduced additional binding environments, reactions, and biological constraints and complicated interpretation of the data. Perhaps performing a phase-resolved mass balance after solid–liquid extraction in the presence and absence of a chelating ion exchange resin would offer a more definitive way to assess the reversibility of the metal-*Phytolacca* interactions. Nonetheless, these efforts demonstrated that biophytometallurgy could provide a route to recover metals from plant resources that could leave the plant biomass in a solid form, which could possibly improve the sustainability^7^ and economics^9^ of the phytometallurgy and phytomining approaches. From a unit operation perspective, (bio)phytometallurgy, that is, solid–liquid extraction, seems interoperable with established^82^ and emerging^83^ (bio)hydrometallurgical capabilities. Since we made no attempts to do so, we expect there are vast opportunities for others to improve upon the efficiency and/or design of this biophytometallurgical proof-of-concept. Beyond extending biophytometallurgy to mine and refine other metals from other plants, future studies could consider whether this framework could useful for examining the mechanisms and consequences of metal partitioning across other biological interfaces. For example, the Banta lab is examining why the partitioning of (antimicrobial-)cations that do not serve as energy substrates for *A. ferrooxidans*^44^ to the (brown) S^0^ interface (Figure S51), and perhaps to or within *A. ferrooxidans* (Text S25), could increase the growth and metabolism more than an equimolar addition of Fe^2+^, which can serve as an energy substrate (Figure S53).

### Associated content

Protocol for routine cell culturing (Text S1); Protocol for cell harvesting and washing (Text S2); Protocol for cell density normalization and quantification (Text S3); Protocol for media preparation (Text S4); Protocol for *Phytolacca* cultivation (Text S5); Protocol for digestion of solid samples (Text S6); Protocol for elemental analysis (Text S7); Protocol for thermogravimetric analysis (Text S8); Protocol for calcination at 600 *^◦^*C (Text S9); Protocol for X-ray diffraction on dried and calcined samples (Text S10); Protocol for Fourier-transform infrared (FTIR) spectroscopy on dried and calcined samples (Text S11); Protocol for scanning electron microscopy with energy-dispersive X-ray spectroscopy on plant and bacterial samples (Text S12); Protocol for inorganic residue washing experiments (Text S13); Protocol for BET surface area analysis of solid samples (Text S14); Protocol for solid–liquid extraction experiments (Text S15); Protocol for REE binding and elution experiments (Text S16); Protocol for membrane fluorescence and growth assays performed on pretreated *A. ferrooxidans* (Text S17); Protocol for *A. ferrooxidans* time-course culture measurements (Text S18); Protocol for NaOH–HCl buffering capacity titrations (Text S19); Protocol for centrifugal ultrafiltration experiments (Text S20); Protocols for screening, bioleaching, cell recovery, and coproduct separation workflows (Text S21); Protocol for S^0^ adsorption experiments (Text S22); Protocol for comparison of REE solubilization and plant elemental composition across studies (Text S23); Descriptions of conditions used for each study in the REE solubilization and plant elemental composition comparison (Text S24); Theoretical and experimental considerations for interpretation of cell-associated metal concentration data (Text S25); Equations for elemental concentrations of liquid and solid samples (Equations S1–S2); Equations for enrichment factors after thermal treatment of plant tissues (Equations S3–S4); Equation for XRD intensity normalization (Equation S5); Equations for FTIR baseline correction, normalization, and spectral-comparison calculations (Equations S6–S14); Equations for mass-balance, enrichment, and elemental-release calculations for water-washing experiments (Equations S15–S22); Equations for metal solubilization and reactant-to-metal-ratio calculations for solid–liquid extraction experiments (Equations S23– S29); Equations for REE group amounts and feed-normalized HREE/LREE solubilization selectivity (Equations S30–S31); Equations for apparent solid–liquid distribution coefficients, surface-area normalization, and selectivity (Equations S32–S34); Equations for REE reload-efficiency and corrected adsorption-capacity calculations (Equations S35–S45); Equations for bacterial REE association, dry-cell-weight normalization, wash-stage mass balances, and concentration-factor calculations (Equations S46–S57); Equations for geometric cell surface-area and dry-mass estimates (Equations S58–S63); Equation for background correction of membrane-associated fluorescence assays (Equation S64); Equations for buffering-capacity and integrated acid- and base-side contribution calculations (Equations S65–S76); Equations for mass, metal, and UV-metric recovery and partitioning calculations for centrifugal-ultrafiltration experiments (Equations S77–S84); Equations for logarithmic REE pool ratios and overall bioleaching mass-balance closure (Equations S85–S86); Equations for stage-specific and overall recovery calculations for plant residues, aqueous REEs, and *A. ferrooxidans* (Equations S87–S94); Equations for control correction and apparent REE adsorption calculations for (elemental) S^0^ (Equations S95–S101); Equations for ideal strong-monoprotic-acid and sulfuric-acid concentrations inferred from initial pH (Equations S102–S104); Equations for ionic activity corrections, EDTA protonation, conditional complexation, and free-metal concentrations (Equations S105–S114); Equations for illustrative first-order extraction kinetics and wash-duration sufficiency criteria (Equations S115– S117); Equations for Tb depletion, recovery, mass-balance closure, and recovered-fraction distribution in phosphate-challenge experiments (Equations S118–S122); Chemicals, materials, and equipment used in this study (Table S1); Bacterial strains used in this study (Table S2); Media formulations used in this study (Table S3); Replicate-level elemental concentrations and molar ratios in lyophilized *Phytolacca* tissues (Table S4); Two-way ANOVA and Holm–Bonferroni-adjusted comparisons of REE concentrations in *P. acinosa* roots and shoots after Ce/Dy/Nd/Tb supplementation (Table S5); Two-way ANOVA and Šídák-adjusted comparisons of HREE concentrations in *P. acinosa* roots and shoots after Ce/Dy/Nd/Tb supplementation (Table S6); Welch’s *t* test comparing HREE/LREE molar ratios between Ce/Dy/Nd/Tb-supplemented *P. acinosa* shoots and roots (Table S7); One-sample Wilcoxon signed-rank tests of HREE/LREE molar ratios in Ce/Dy/Nd/Tb-supplemented *P. acinosa* shoots and roots against unity (Table S8); Two-way ANOVA and Holm–Bonferroni-adjusted comparisons of Ni concentrations in *P. acinosa* roots and shoots after Ni supplementation (Table S9); Brown–Forsythe and Welch ANOVAs with Dunnett’s T3 comparisons of REE concentrations and a Wilcoxon signed-rank test of HREE/LREE molar ratios in *P. americana* shoots (Table S10); Replicate-level BET surface areas and BJH cumulative pore volumes of control and Ce/Dy-enriched *P. americana* shoots and brown and yellow S^0^ (Table S11); Thermal decomposition metrics extracted from TGA/DTG analysis of lyophilized *Phytolacca* tissues (Table S12); Two-way ANOVA and Šídák-adjusted comparisons of lower- and higher-temperature DTG peaks in Ce/Dy-supplemented and control *P. americana* shoots (Table S13); Elemental concentrations and enrichment factors in post-TGA *Phytolacca* residues (Table S14); Elemental concentrations and enrichment factors in *Phytolacca* residues after calcination at 600 *^◦^*C (Table S15); Two-way ANOVA and Tukey-adjusted comparisons of baseline-corrected TREE/P molar ratios by thermochemical treatment and plant material (Table S16); Paired *t* test and repeated-measures ANOVA of elemental release during washing of REE-enriched *P. americana* shoot ash (Table S17); Comparison of log-linear fits relating REE solubilization to nominal H_2_SO_4_-to-REE ratios varied through acid dosage or biomass loading (Table S18); Comparison of concentration-dependent log-linear fits of REE solubilization using equimolar dosages of H_2_SO_4_, K_2_SO_4_, and KNO_3_ (Table S19); Brown–Forsythe and Welch ANOVAs and extra sum-of-squares *F* tests of acid identity and pH-dependent REE solubilization (Table S20); Two-way ANOVA and Šídák-adjusted comparisons of acid and electrolyte effects on REE solubilization and selectivity (Table S21); Linear regression and extra sum-of-squares *F* test of the Fe_2_(SO_4_)_3_ concentration effect on REE solubilization (Table S22); Matched-pH statistical comparisons of chelator-mediated REE solubilization against H_2_SO_4_ controls (Table S23); Two-way ANOVA and Šídák-adjusted comparisons of chelator identity and In^3+^ pre-loading effects on REE solubilization (Table S24); Multiple linear regression of feed-normalized HREE/LREE molar selectivity by reagent, KNO_3_ addition, and their interaction (Table S25); Comparison of log-linear fits relating Ni solubilization from *P. acinosa* shoots to reactant-to-Ni ratios varied through Ni supplementation during plant cultivation (Table S26); Repeated-measures ANOVA and Dunnett comparisons of electrolyte-dependent REE solubilization from REE-enriched *P. americana* shoots (Table S27); One-sample *t* tests, Brown–Forsythe and Welch ANOVAs, and Dunnett’s T3 comparisons of REE reload efficiency in *P. americana* shoots (Table S28); Wilcoxon signed-rank tests of log_10_-transformed REE loading ratios after reloading leached *P. americana* residues relative to the non-leached REE-enriched reference (Table S29); Brown–Forsythe and Welch ANOVAs with Dunnett’s T3 comparisons of incubation-time effects on Tb retention and inferred binding capacity of *A. ferrooxidans* (Table S30); Brown–Forsythe and Welch ANOVAs with Dunnett’s T3 comparisons of Tb retention across EDTA washing protocols after 24 h of binding (Table S31); Two-way ANOVA of inferred Tb recovery by *A. ferrooxidans* across repeated adsorption–desorption cycles (Table S32); Šídák- and Tukey-adjusted comparisons of Tb binding, citrate elution, and recovery across *A. ferrooxidans* pretreatments and repeated adsorption–desorption cycles (Table S33); Brown–Forsythe and Welch ANOVAs with Dunnett’s T3 comparisons of membrane-associated fluorescence responses across *A. ferrooxidans* pretreatments (Table S34); One-way ANOVA and Šídák-adjusted comparisons of cell-associated Tb after *A. ferrooxidans* growth in Tb-supplemented medium and subsequent binding tests (Table S35); Two-way ANOVA and Šídák-adjusted comparisons of REE-binding capacity after *A. ferrooxidans* growth in F2S medium with or without Tb supplementation (Table S36); Two-way ANOVA of lixiviant, tissue condition, and their interaction for ΔpH and ΔORP during solid–liquid extraction of metals from *Phytolacca* shoots (Table S37); Dunnett-adjusted comparisons of ΔpH between biomass treatments and no-shoot controls within each lixiviant (Table S38); Dunnett-adjusted comparisons of ΔORP between biomass treatments and no-shoot controls within each lixiviant (Table S39); One-sample *t* tests of mass retention against 50% after solid–liquid extraction of metals from *Phytolacca* shoots (Table S40); One-sample *t* tests of UV absorbance retention against 50% in ultrafiltration fractions of *Phytolacca* shoot leachates (Table S41); One-sample *t* tests of Ni or REE retention against 50% after solid–liquid extraction of metals from *Phytolacca* shoots (Table S42); Regression models relating Ni retention during 3 kDa ultrafiltration of *P. acinosa* leachates to mass retention, UV absorbance retention, and pH (Table S43); Regression models relating REE retention during 3 kDa ultrafiltration of *P. americana* leachates to mass retention, UV absorbance retention, and pH (Table S44); Multiple linear regression of inferred REE-binding capacity by *A. ferrooxidans* across most binding tests (Table S45); Regression models relating endpoint REE solubilization in *P. americana* bioleaching experiments to experimental factors and measured solution properties (Table S46); REML mixed-effects analysis of time, treatment, and their interaction for REE solubilization, pH, ORP, and conductivity across 14 bioleaching and abiotic-control treatments (Table S47); REML mixed-effects analysis of time, inoculum group, and their interaction for REE solubilization, pH, ORP, and conductivity in the matched inoculum-density series (Table S48); Regression analysis of replicate-level bioleaching time-course summary metrics by inoculum group and pH metrics (Table S49); Calibration of SYBR Green fluorescence and optical density for estimating planktonic *A. ferrooxidans* cell density (Figure S1); Wet and dry cell mass of *A. ferrooxidans* and *E. coli* suspensions normalized to OD_600_ = 1.0 (Figure S2); SEM images of shoot tissue morphology in hydroponically grown, Ce/Dy-supplemented *P. americana* shoots (Figure S3); SEM images of shoot tissue morphology in hydroponically grown control *P. americana* shoots (Figure S4); SEM imaging and EDS mapping of a faceted structure associated with Ce/Dy-supplemented *P. americana* shoot tissue (Figure S5); SEM imaging and EDS mapping of a shoot tissue fragment from hydroponically grown control *P. americana* (Figure S6); TGA and DTG profiles of lyophilized *Phytolacca* tissues by tissue type, growth condition, and metal supplementation (Figure S7); Raw X-ray diffraction patterns of dried and 600 *^◦^*C-calcined *Phytolacca* shoots with a silicon-mount reference (Figure S8); Comparison of FTIR spectra for dried *Phytolacca* and ash by treatment condition after regional baseline correction and vector normalization (Figure S9); Deportment of Ca, K, Mg, and P during water washing of *Phytolacca* ash produced at 600 *^◦^*C (Figure S10); Time course of REE solubilization from *P. acinosa* tissues using 0.05 M Na_2_EDTA at pH 4.5 (Figure S11); Time course of REE and Ni solubilization from metal-enriched *Phytolacca* shoots during acid and EDTA extraction (Figure S12); Long-term REE and Ni solubilization from *Phytolacca* shoots following extended static storage, by pH and biomass loading (Figure S13); Effects of mineral acid identity, pH, and electrolyte addition on REE solubilization and Dy/Ce selectivity from Ce/Dy-enriched *P. americana* shoots (Figure S14); Apparent endpoint Fe^2+^ speciation as a function of initial Fe^3+^ concentrations incubated with and without Ce/Dyenriched *P. americana* shoots (Figure S15); Initial and final soluble In concentrations in citric acid and EDTA competition tests after incubation with and without *P. americana* shoots (Figure S16); Dy and Ce solubilization and feed-normalized HREE/LREE selectivity from *P. americana* shoots after solid–liquid extraction with and without KNO_3_ addition (Figure S17); Ni and REE solubilization from enriched *P. acinosa* shoots as a function of reagent dosage during acid and chelator extraction (Figure S18); Condition-specific apparent solid–liquid distribution coefficients and Dy/Ce selectivity during REE and Ni extraction from *Phytolacca* shoots as a function of reagent concentration (Figure S19); Endpoint SYBR Green fluorescence in Niand REE-enriched *Phytolacca* shoot leachates and matched no-biomass controls across extraction conditions (Figure S20); SYBR Green fluorescence of *P. americana* shoot lysates as a function of tissue mass after SDS and heat treatment (Figure S21); Time course of pH, ORP, dissolved sulfate, and SYBR Green fluorescence during water incubation of autoclaved and non-autoclaved *P. americana* shoots (Figure S22); REE solubilization time courses, apparent solid–liquid distribution coefficients, and HREE/LREE selectivity during acid, chelator, and electrolyte extraction of *P. americana* shoots (Figure S23); Ce and Dy uptake, release, and retention during sequential rebinding, unbinding, and rebinding of previously leached *P. americana* shoot residues (Figure S24); Baseline-corrected and vector-normalized FTIR spectra of *P. americana* shoots before extraction, after leaching, and after Ce/Dy reloading (Figure S25); Photographs of residues formed during attempted lyophilization of *P. americana* shoot leachates generated with 0.1 and 3 M H_2_SO_4_ (Figure S26); UV–visible absorbance spectra of diluted residues obtained by attempted lyophilization of H_2_SO_4_ leachates from *P. americana* shoots (Figure S27); Time-dependent inferred REE loading, grade– recovery relationships, and apparent partitioning by *A. ferrooxidans* in mixed-REE simulants and *P. acinosa* shoot leachate at pH 1.8 (Figure S28); Inferred REE binding, grade–recovery relationships, and apparent HREE/LREE selectivity of *A. ferrooxidans* as a function of initial mixed-REE concentrations and cell densities (Figure S29); Inferred Tb binding, grade–recovery relationships, and apparent solid–liquid distribution coefficients of *A. ferrooxidans* as a function of initial Tb concentrations and cell densities (Figure S30); Effect of binding duration on inferred Tb association with *A. ferrooxidans* and Tb distribution across sequential EDTA washes, water washes, and final pellets (Figure S31); Effect of EDTA washing duration and protocol on cell-associated Tb retention and recovery after 24 h binding by *A. ferrooxidans* (Figure S32); Tb distribution and recovery after cell-free incubation with or without excess phosphate and subsequent sequential washing (Figure S33); SEM images of *A. ferrooxidans* preparations after 24 h incubation with or without Tb and subsequent EDTA or water-only washing (Figure S34); SEM imaging and EDS mapping of Tb-exposed, EDTA-washed *A. ferrooxidans* biomass mounted on carbon tape (Figure S35); SEM imaging and EDS mapping of *A. ferrooxidans* biomass collected before Tb-binding experiments and mounted on carbon tape (Figure S36); SEM imaging and EDS mapping of Tb-exposed, EDTA-washed *A. ferrooxidans* biomass mounted on a silicon wafer (Figure S37); SEM imaging and EDS mapping of Tb-exposed, water-washed *A. ferrooxidans* biomass mounted on a silicon wafer (Figure S38); Inferred Tb binding, citrate elution, and recovery across three adsorption–desorption cycles and different *A. ferrooxidans* pretreatments (Figure S39); Membrane-associated fluorescence, (apparent) substrate oxidation, and growth of *A. ferrooxidans* following heat, UV-C, and Triton X-100 and sonication treatments (Figure S40); Tb grade–recovery relationships, apparent partitioning, and lower-bound selectivity estimates for *A. ferrooxidans* and *E. coli* DH5*α* at pH 1.8 (Figure S41); Inferred Tb loading, recovery, and apparent partitioning by *E. coli* DH5*α* as a function of initial Tb concentration and cell density at pH 6.0 (Figure S42); Fe^2+^ oxidation, growth, and subsequent REE association of *A. ferrooxidans* cultured with or without 30 *µ*M Tb^3+^ from Tb_2_(SO_4_)_3_ (Figure S43); Biomass-associated shifts in pH and ORP relative to matched no-shoot controls during water, acid, and EDTA extraction of *Phytolacca* shoots (Figure S44); Biomass-corrected acid- and base-side buffering profiles of *Phytolacca* shoot leachates as a function of initial pH and biomass loadings (Figure S45); Integrated buffering contributions and their distribution across pH intervals in Ni- and Ce/Dy-enriched *Phytolacca* shoot leachates (Figure S46); Mass, UV absorbance, and Ni or REE retention during 3 kDa centrifugal ultrafiltration of some *Phytolacca* shoot leachates (Figure S47); Endpoint Fe^2+^ oxidation and pH in *A. ferrooxidans* cultures grown with increasing *P. americana* shoot loading in Fe^2+^- or S^0^-containing media (Figure S48); Time course of REE solubilization, solution chemistry, and apparent cell densities during *P. americana* bioleaching at varied shoot loadings, S^0^ dosages, and *A. ferrooxidans* inoculum densities (Figure S49); Time course of pH, ORP, conductivity, and apparent cell densities in bioleaching controls without *P. americana* shoots (Figure S50); Inferred Ce and Dy loading, apparent partitioning, and Dy/Ce selectivity of brown and yellow S^0^ at pH 2.0 (Figure S51); Ce and Dy mass balances and cell-associated recovery in EDTA-washed *A. ferrooxidans* pellets after bioleaching of *P. americana* shoots for some experiments (Figure S52); Time course of planktonic *A. ferrooxidans* density and pH under 100 *µ*M cation supplementation in the absence of plant biomass (Figure S53); Time courses of pH, soluble Ce and Dy, and planktonic cell density across *P. americana* shoot-enrichment, dissolved-REE supplementation, and *A. ferrooxidans* inoculation conditions (Figure S54); Partitioning of plant solids, REEs, and *A. ferrooxidans* during two-stage centrifugation of *P. acinosa* bioleaching cultures (Figure S55); Photographic comparison of dried Ce/Dy-enriched *P. americana* shoots, ash produced at 600 *^◦^*C, and monazite-bearing ore (Figure S56); Literature comparison of REE solubilization, feed-normalized HREE/LREE selectivity, and REE and P composition in plant tissues and thermally treated residues (Figure S57).

## Acknowledgments

During investigation, D.A.D. used the newest available models of ChatGPT (OpenAI) to research and devise experimental controls and their limitations for all experiments, some of which were ultimately performed. ChatGPT was not used to generate experimental data or make final scientific conclusions. After the original draft was completed by D.A.D., E.H.-P., S.B., S.T.B., N.E.R., and A.P.-P., D.A.D. used ChatGPT to assist with language editing, organization, and concision; the identification of additional potentially relevant sources, many of which were not relevant or cited; interpretation of author-generated statistical outputs; feedback on author-generated visualizations for concision and accessibility; LaTeX formatting and troubleshooting for the manuscript and responses to reviewers; and to see the extent the models thought the claims made herein were supported by the data. The Reviewer 3 software (Reviewer3, Inc.) was also used to assess the strength of the support for the presented claims, which prompted three minor wording changes to emphasize the limitations to some data. All AI-assisted material was reviewed, revised, and independently verified by the authors against the underlying experimental data and primary literature. The authors take full responsibility for the accuracy, integrity, and content of the manuscript and Supporting Information.

D.A.D., S.T.B., and S.A.B. gratefully acknowledge funding through ARPA-E grant DE-AR0001340 from the US Department of Energy. D.A.D. was partially supported by the Bourbon County Educational Fund, the Tau Beta Pi Fellowship (Zimmerman #12), and the National Science Foundation Graduate Research Fellowship Program (DGE-2437839). E.H.-P. and C.J.D. gratefully acknowledge funding through a DARPA Young Investigator Award D19AP00026 from the Department of Defense and the S24AC00044-00 award from the Office of Surface Mining, Reclamation, and Enforcement. S.B. and R.N.A. gratefully acknowledge support from an NSF MRI award 2320054 which facilitated the ICP-MS measurements. Any opinions, findings, and conclusions or recommendations expressed in this material are those of the author(s) and do not necessarily reflect the views or policies of the National Science Foundation, the Department of Defense, or the U.S. Government.

The authors thank Natalie R. Ling for creation of non-data figures; William J. Wei for assistance with the XRD analysis; and Wei Wang, Sameera S. Abeyrathna, and Farid F. Khoury for their thoughtful discussion and feedback on our experiments.

## Data availability

Full data for this work beyond what is provided in the Supplementary information is available upon request.

