## Supporting Information v02 for "Biophytometallurgy: biomining metals from plant resources"

Number of pages: 132
Number of texts: 25
Number of equations: 122
Number of tables: 49
Number of figures: 57

### CONTENTS

**Text S1**. Protocol for routine cell culturing.
**Text S2**. Protocol for cell harvesting and washing.
**Text S3**. Protocol for cell density normalization and quantification.
**Text S4**. Protocol for media preparation.
**Text S5**. Protocol for *Phytolacca* cultivation.
**Text S6**. Protocol for digestion of solid samples.
**Text S7**. Protocol for elemental analysis.
**Text S8**. Protocol for thermogravimetric analysis.
**Text S9**. Protocol for calcination at 600 °C.
**Text S10**. Protocol for X-ray diffraction on dried and calcined samples.
**Text S11**. Protocol for Fourier-transform infrared (FTIR) spectroscopy on dried and calcined samples.
**Text S12**. Protocol for scanning electron microscopy with energy dispersive X-ray spectroscopy on plant and bacteria samples.
**Text S13**. Protocol for inorganic residue washing experiments.
**Text S14**. Protocol for BET surface area analysis for solid samples.
**Text S15**. Protocol for solid-liquid extraction experiments.
**Text S16**. Protocol for REE binding and elution experiments.
**Text S17**. Protocol for membrane fluorescence and growth assays performed on pretreated *A. ferrooxidans*.
**Text S18**. Protocol for *A. ferrooxidans* time-course culture measurements.
**Text S19**. Protocol for NaOH–HCl buffering capacity titrations.
**Text S20**. Protocol for centrifugal ultrafiltration experiments.
**Text S21**. Protocols for screening, bioleaching, cell recovery, and coproduct separation workflows.
**Text S22**. Protocol for $S^0$ adsorption experiments.
**Text S23**. Protocol for comparison of REE solubilization and plant elemental composition across studies.
**Text S24**. Descriptions of conditions used for each study in the REE solubilization and plant elemental composition comparison.
**Text S25**. Theoretical and experimental considerations for interpretation of cell-associated metal concentration data.
**Equations S1–S2**. Elemental concentrations of liquid and solid samples.
**Equations S3–S4**. Enrichment factors after thermal treatment of plant tissues.
**Equation S5**. XRD intensity normalization.
**Equations S6–S14**. FTIR baseline correction, normalization, and spectral-comparison calculations.
**Equations S15–S22**. Mass-balance, enrichment, and elemental-release calculations for water-washing experiments.
**Equations S23–S29**. Metal solubilization and reactant-to-metal-ratio calculations for solid–liquid extraction experiments.
**Equations S30–S31**. REE group amounts and feed-normalized HREE/LREE solubilization selectivity.
**Equations S32–S34**. Apparent solid–liquid distribution coefficients, surface-area normalization, and selectivity.
**Equations S35–S45**. REE reload-efficiency and corrected adsorption-capacity calculations.
**Equations S46–S57**. Bacterial REE association, dry-cell-weight normalization, and concentration-factor calculations.
**Equations S58–S63**. Geometric cell surface-area and dry-mass estimates.
**Equation S64**. Background correction for membrane-associated fluorescence assays.

**Equations S65–S76**. Buffering-capacity and integrated acid- and base-side contribution calculations.

**Equations S77–S84**. Mass, metal, and UV-metric recovery and partitioning calculations for centrifugal-ultrafiltration experiments.

**Equations S85–S86**. Logarithmic REE pool ratios and overall bioleaching mass-balance closure.

**Equations S87–S94**. Stage-specific and overall recovery calculations for plant residues, aqueous REEs, and *A. ferrooxidans*.

**Equations S95–S101**. Control correction and apparent REE adsorption calculations for (elemental) $S^0$.

**Equations S102–S104**. Ideal strong-monoprotic-acid and sulfuric-acid concentrations inferred from initial pH.

**Equations S105–S114**. Ionic activity corrections, EDTA protonation, conditional complexation, and free-metal concentrations.

**Equations S115–S117**. Illustrative first-order extraction kinetics and wash-duration sufficiency criteria.

**Equations S118–S122**. Tb depletion, recovery, mass-balance closure, and recovered-fraction distribution in phosphate-challenge experiments.

**Table S1**. Chemicals, materials, and equipment used in this study.

**Table S2**. Bacterial strains used in this study.

**Table S3**. Media formulations used in this study.

**Table S4**. Replicate-level elemental concentrations and molar ratios in lyophilized *Phytolacca* tissues.

**Table S5**. Two-way ANOVA and Holm–Bonferroni-adjusted contrasts of $\log_{10}$-transformed REE concentrations in *P. acinosa* roots and shoots after Ce/Dy/Nd/Tb supplementation.

**Table S6**. Two-way ANOVA and Šídák-adjusted comparisons of $\log_{10}$-transformed HREE concentrations in *P. acinosa* roots and shoots after Ce/Dy/Nd/Tb supplementation.

**Table S7**. Unpaired Welch's $t$ test of $\log_{10}$-transformed HREE/LREE molar ratios between Ce/Dy/Nd/Tb-supplemented *P. acinosa* shoots and roots.

**Table S8**. Exact one-sample Wilcoxon signed-rank tests of $\log_{10}$-transformed HREE/LREE molar ratios against zero in Ce/Dy/Nd/Tb-supplemented *P. acinosa* shoots and roots.

**Table S9**. Two-way ANOVA and Holm–Bonferroni-adjusted contrasts of $\log_{10}$-transformed Ni concentrations in *P. acinosa* roots and shoots after Ni supplementation.

**Table S10**. Brown–Forsythe and Welch ANOVAs with Dunnett's T3 comparisons of REE concentrations, and a Wilcoxon signed-rank test of HREE/LREE ratios, in *P. americana* shoots.

**Table S11**. Replicate-level BET surface areas and BJH cumulative pore volumes of control and Ce/Dy-enriched *P. americana* shoots and brown and yellow $S^0$.

**Table S12**. Thermal decomposition metrics extracted from TGA/DTG analysis of lyophilized *Phytolacca* tissues.

**Table S13**. Ordinary two-way ANOVA and Šídák-adjusted comparisons of DTG peak temperatures in Ce/Dy-supplemented and control *P. americana* shoots.

**Table S14**. Elemental concentrations and enrichment factors in post-TGA *Phytolacca* residues.

**Table S15**. Elemental concentrations and enrichment factors in *Phytolacca* residues after calcination at 600 °C.

**Table S16**. Two-way ANOVA and Tukey-adjusted comparisons of $\log_{10}$-transformed, baseline-corrected TREE/P molar ratios by thermochemical treatment and plant material.

**Table S17**. Paired $t$ test and repeated-measures one-way ANOVA with Šídák-adjusted comparisons of elemental release during washing of REE-enriched *P. americana* shoot ash.

**Table S18**. Log-linear regression and extra sum-of-squares $F$ test comparing REE solubilization from REE-enriched *P. americana* shoots to the nominal $H_2SO_4$-to-REE ratio obtained by varying $H_2SO_4$ dose or biomass loading.

**Table S19**. Log-linear regression and extra sum-of-squares *F* test comparing concentration-dependent REE solubilization from REE-enriched *P. americana* shoots using equimolar dosages of $H_2SO_4$, $K_2SO_4$, and $KNO_3$.

**Table S20**. Brown–Forsythe and Welch ANOVAs and extra sum-of-squares *F* tests of acid-dependent REE solubilization from REE-enriched *P. americana* shoots as a function of pH for some mineral and organic acids.

**Table S21**. Ordinary two-way ANOVA and Šídák-adjusted comparisons of acid and electrolyte effects on REE solubilization from REE-enriched *P. americana* shoots and its selectivity.

**Table S22**. Linear regression and extra sum-of-squares *F* test of the $Fe_2(SO_4)_3$ concentration effect on REE solubilization from REE-enriched *P. americana* shoots.

**Table S23**. Unpaired *t* test and one-way ANOVA with Dunnett comparisons of chelator-mediated REE solubilization from REE-enriched *P. americana* shoots against matched-pH $H_2SO_4$ controls.

**Table S24**. Two-way ANOVA and Šídák-adjusted comparisons of chelator identity and $In^{3+}$ pre-loading effects on REE solubilization from REE-enriched *P. americana* shoots.

**Table S25**. Multiple linear regression of $\log_2$-transformed HREE/LREE molar selectivity of REE solubilization from REE-enriched *P. americana* shoots by reagent, $KNO_3$ addition, and their interaction.

**Table S26**. Log-linear regression and extra sum-of-squares *F* test comparing Ni solubilization from Ni-enriched *P. acinosa* shoots versus reactant-to-Ni ratio obtained by varying the concentration of Ni supplementation during plant cultivation.

**Table S27**. Geisser–Greenhouse-corrected two-way repeated-measures ANOVA and Dunnett comparisons of electrolyte-dependent REE solubilization from REE-enriched *P. americana* shoots.

**Table S28**. One-sample *t* tests, Brown–Forsythe and Welch ANOVAs, and Dunnett's T3 comparisons of $\log_{10}$-transformed REE reload efficiency of (REE-enriched) *P. americana* shoots.

**Table S29**. Wilcoxon signed-rank tests of $\log_{10}$-transformed REE loading ratios relative to the non-leached REE-enriched *P. americana* reference loading after reloading leached REE-enriched *P. americana* residues.

**Table S30**. Brown–Forsythe and Welch ANOVAs with Dunnett's T3 comparisons of $\log_{10}$-transformed Tb retention and inferred binding capacity of *A. ferrooxidans* across incubation times.

**Table S31**. Brown–Forsythe and Welch ANOVAs with Dunnett's T3 comparisons of Tb retention by *A. ferrooxidans* across EDTA washing protocols after 24 h of binding.

**Table S32**. Ordinary two-way ANOVA of $\log_{10}$-transformed inferred Tb recovery by *A. ferrooxidans* across repeated adsorption–desorption cycles.

**Table S33**. Šídák- and Tukey-adjusted comparisons from two-way ANOVAs of $\log_{10}$-transformed Tb binding, citrate elution, and recovery across some *A. ferrooxidans* pretreatments and repeated adsorption–desorption cycles.

**Table S34**. Brown–Forsythe and Welch one-way ANOVAs with Dunnett's T3 comparisons of some membrane-associated fluorescence assay responses as a function of *A. ferrooxidans* pretreatment.

**Table S35**. One-way ANOVA and Šídák-adjusted comparisons of $\log_{10}$-transformed cell-associated Tb after *A. ferrooxidans* growth in Tb-supplemented medium and subsequent binding tests.

**Table S36**. Two-way ANOVA and Šídák-adjusted comparisons of $\log_{10}$-transformed REE-binding capacity in *A. ferrooxidans* after growth in F2S medium with or without Tb supplementation.

**Table S37**. Ordinary two-way ANOVA of lixiviant, tissue condition, and their interaction for $\Delta$pH and $\Delta$ORP during solid–liquid extraction of metals from (enriched) *P. acinosa* and *P. americana* shoots.

**Table S38**. Dunnett-adjusted comparisons of $\Delta$pH following two-way ANOVA between biomass treatments and no-shoot controls within each lixiviant after solid–liquid extraction of metals from (enriched) *P. acinosa* and *P. americana* shoots.

**Table S39**. Dunnett-adjusted comparisons of ΔORP following two-way ANOVA between biomass treatments and no-shoot controls within each lixiviant after solid–liquid extraction of metals from (enriched) *P. acinosa* and *P. americana* shoots.

**Table S40**. Two-tailed one-sample $t$ tests of mass retention against 50% after solid–liquid extraction of metals from (enriched) *P. acinosa* and *P. americana* shoots.

**Table S41**. Two-tailed one-sample $t$ tests of UV absorbance retention against 50% in ultrafiltration fractions after solid–liquid extraction of metals from (enriched) *P. acinosa* and *P. americana* shoots.

**Table S42**. Two-tailed one-sample $t$ tests of Ni or REE retention against 50% after solid–liquid extraction of metals from (enriched) *P. acinosa* and *P. americana* shoots.

**Table S43**. Ordinary least-squares regression models relating Ni retention during 3 kDa ultrafiltration of *P. acinosa* leachates to mass retention, UV absorbance retention, and pH.

**Table S44**. Ordinary least-squares regression models relating REE retention during 3 kDa ultrafiltration of *P. americana* leachates to mass retention, UV absorbance retention, and pH.

**Table S45**. Ordinary least-squares multiple regression of $\log_{10}$-transformed inferred REE-binding capacity by *A. ferrooxidans* across most binding tests.

**Table S46**. Ordinary least-squares regression models of endpoint REE solubilization in *P. americana* bioleaching experiments to some experimental factors and measurements.

**Table S47**. REML mixed-effects analysis of time, treatment, and their interaction for *P. americana* shoot bioleaching trajectories of REE solubilization, pH, ORP, and conductivity.

**Table S48**. REML mixed-effects analysis of time, inoculum group, and their interaction for REE solubilization, pH, ORP, and conductivity for the *P. americana* shoot bioleaching experiments.

**Table S49**. Ordinary least-squares regression of replicate-level bioleaching time-course summary metrics by inoculum group and pH metrics for the *P. americana* shoot bioleaching experiments.

**Figure S1**. Calibration of SYBR Green fluorescence and optical density for estimating planktonic *A. ferrooxidans* cell density.

**Figure S2**. Wet and dry cell mass of *A. ferrooxidans* and *E. coli* suspensions normalized to $OD_{600} = 1.0$.

**Figure S3**. SEM images of shoot tissue morphology in hydroponically grown, Ce/Dy-supplemented *P. americana* shoots.

**Figure S4**. SEM images of shoot tissue morphology in hydroponically grown control *P. americana* shoots.

**Figure S5**. SEM imaging and EDS mapping of a faceted structure associated with Ce/Dy-supplemented *P. americana* shoot tissue.

**Figure S6**. SEM imaging and EDS mapping of a shoot tissue fragment from hydroponically grown control *P. americana*.

**Figure S7**. TGA and DTG profiles of lyophilized *Phytolacca* tissues by tissue type, growth condition, and metal supplementation.

**Figure S8**. Raw X-ray diffraction patterns of dried and 600 °C-calcined *Phytolacca* shoots with a silicon-mount reference.

**Figure S9**. Comparison of FTIR spectra for dried *Phytolacca* and ash by treatment condition after regional baseline correction and vector normalization.

**Figure S10**. Deportment of Ca, K, Mg, and P during water washing of *Phytolacca* ash produced at 600 °C.

**Figure S11**. Time course of REE solubilization from *P. acinosa* tissues using 0.05 M $Na_2EDTA$ at pH 4.5.

**Figure S12**. Time course REE and Ni solubilization from metal-enriched *Phytolacca* shoots during acid and EDTA extraction.

**Figure S13**. Long-term REE and Ni solubilization from *Phytolacca* shoots following extended static storage, by pH and biomass loading.

**Figure S14**. Effects of mineral acid identity, pH, and electrolyte addition on REE solubilization and Dy/Ce selectivity from Ce/Dy-enriched *P. americana* shoots.

**Figure S15**. Apparent endpoint $Fe^{2+}$ speciation as a function of initial $Fe^{3+}$ concentrations incubated with and without Ce/Dy-enriched *P. americana* shoots.

**Figure S16**. Initial and final soluble In concentrations in citric acid and EDTA competition tests after incubation with and without *P. americana* shoots.

**Figure S17**. Dy and Ce solubilization and feed-normalized HREE/LREE selectivity from *P. americana* shoots after solid–liquid extraction with and without $KNO_3$ addition.

**Figure S18**. Ni and REE solubilization from enriched *P. acinosa* shoots as a function of reagent dosage during acid and chelator extraction.

**Figure S19**. Condition-specific apparent solid–liquid distribution coefficients and Dy/Ce selectivity during REE and Ni extraction from *Phytolacca* shoots as a function of reagent concentration.

**Figure S20**. Endpoint SYBR Green fluorescence in Ni- and REE-enriched *Phytolacca* shoot leachates and matched no-biomass controls across extraction conditions.

**Figure S21**. SYBR Green fluorescence of *P. americana* shoot lysates as a function of tissue mass after SDS and heat treatment.

**Figure S22**. Time course pH, ORP, dissolved sulfate, and SYBR Green fluorescence during water incubation of autoclaved and non-autoclaved *P. americana* shoots.

**Figure S23**. REE solubilization time courses, apparent solid–liquid distribution coefficients, and HREE/LREE selectivity during acid, chelator, and electrolyte extraction of *P. americana* shoots.

**Figure S24**. Ce and Dy uptake, release, and retention during sequential rebinding, unbinding, and rebinding of previously leached *P. americana* shoot residues.

**Figure S25**. Baseline-corrected and vector-normalized FTIR spectra of *P. americana* shoots before extraction, after leaching, and after Ce/Dy reloading.

**Figure S26**. Photographs of residues formed during attempted lyophilization of *P. americana* shoot leachates generated with 0.1 and 3 M $H_2SO_4$.

**Figure S27**. UV–visible absorbance spectra of diluted residues obtained by attempted lyophilization of $H_2SO_4$ leachates from *P. americana* shoots.

**Figure S28**. Time-dependent inferred REE loading, grade–recovery relationships, and apparent partitioning by *A. ferrooxidans* in mixed-REE simulants and *P. acinosa* shoot leachate at pH 1.8.

**Figure S29**. Inferred REE binding, grade–recovery relationships, and apparent HREE/LREE selectivity of *A. ferrooxidans* as a function of initial mixed-REE concentrations and cell densities.

**Figure S30**. Inferred Tb binding, grade–recovery relationships, and apparent solid–liquid distribution coefficients of *A. ferrooxidans* as a function of initial Tb concentrations and cell densities.

**Figure S31**. Effect of binding duration on inferred Tb association with *A. ferrooxidans* and Tb distribution across sequential EDTA washes, water washes, and final pellets.

**Figure S32**. Effect of EDTA washing duration and protocol on cell-associated Tb retention and recovery after 24 h binding by *A. ferrooxidans*.

**Figure S33**. Tb distribution and recovery after cell-free incubation with or without excess phosphate and subsequent sequential washing.

**Figure S34**. SEM images of *A. ferrooxidans* preparations after 24 h incubation with or without Tb and subsequent EDTA or water-only washing.

**Figure S35**. SEM imaging and EDS mapping of Tb-exposed, EDTA-washed *A. ferrooxidans* biomass mounted on carbon tape.

**Figure S36**. SEM imaging and EDS mapping of *A. ferrooxidans* biomass collected before Tb-binding experiments and mounted on carbon tape.

**Figure S37**. SEM imaging and EDS mapping of Tb-exposed, EDTA-washed *A. ferrooxidans* biomass mounted on a silicon wafer.

**Figure S38**. SEM imaging and EDS mapping of Tb-exposed, water-washed *A. ferrooxidans* biomass mounted on a silicon wafer.

**Figure S39**. Inferred Tb binding, citrate elution, and recovery across three adsorption–desorption cycles and different *A. ferrooxidans* pretreatments.

**Figure S40**. Membrane-associated fluorescence, (apparent) substrate oxidation, and growth of *A. ferrooxidans* following heat, UV-C, and Triton X-100 and sonication treatments.

**Figure S41**. Tb grade–recovery relationships, apparent partitioning, and lower-bound selectivity estimates for *A. ferrooxidans* and *E. coli* DH5$\alpha$ at pH 1.8.

**Figure S42**. Inferred Tb loading, recovery, and apparent partitioning by *E. coli* DH5$\alpha$ as a function of initial Tb concentration and cell density at pH 6.0.

**Figure S43**. $Fe^{2+}$ oxidation, growth, and subsequent REE association of *A. ferrooxidans* cultured with or without 30 $\mu$M $Tb^{3+}$ from $Tb_2(SO_4)_3$.

**Figure S44**. Biomass-associated shifts in pH and ORP relative to matched no-shoot controls during water, acid, and EDTA extraction of *Phytolacca* shoots.

**Figure S45**. Biomass-corrected acid- and base-side buffering profiles of *Phytolacca* shoot leachates as a function of initial pH and biomass loadings.

**Figure S46**. Integrated buffering contributions and their distribution across pH intervals in Ni- and Ce/Dy-enriched *Phytolacca* shoot leachates.

**Figure S47**. Mass, UV absorbance, and Ni or REE retention during 3 kDa centrifugal ultrafiltration of some *Phytolacca* shoot leachates.

**Figure S48**. Endpoint $Fe^{2+}$ oxidation and pH in *A. ferrooxidans* cultures grown with increasing *P. americana* shoot loading in $Fe^{2+}$- or $S^0$-containing media.

**Figure S49**. Time course REE solubilization, solution chemistry, and apparent cell densities during *P. americana* bioleaching at varied shoot loadings, $S^0$ dosages, and *A. ferrooxidans* inoculum densities.

**Figure S50**. Time course pH, ORP, conductivity, and apparent cell densities in bioleaching controls without *P. americana* shoots.

**Figure S51**. Inferred Ce and Dy loading, apparent partitioning, and Dy/Ce selectivity of brown and yellow $S^0$ at pH 2.0.

**Figure S52**. Ce and Dy mass balances and cell-associated recovery in EDTA-washed *A. ferrooxidans* pellets after bioleaching of *P. americana* shoots for some experiments.

**Figure S53**. Time course of planktonic *A. ferrooxidans* density and pH under 100 $\mu$M cation supplementation in the absence of plant biomass.

**Figure S54**. Time courses of pH, soluble Ce and Dy, and planktonic cell density across *P. americana* shoot-enrichment, dissolved-REE supplementation, and *A. ferrooxidans* inoculation conditions.

**Figure S55**. Partitioning of plant solids, REEs, and *A. ferrooxidans* during two-stage centrifugation of *P. acinosa* bioleaching cultures.

**Figure S56**. Photographic comparison of dried Ce/Dy-enriched *P. americana* shoots, ash produced at 600 °C, and monazite-bearing ore.

**Figure S57**. Literature comparison of REE solubilization, feed-normalized HREE/LREE selectivity, and REE and P composition in plant tissues and thermally treated residues.

### Supplementary text

#### Text S1. Protocol for routine cell culturing.

For routine culturing of *A. ferrooxidans* cells (ATCC 23270), frozen stocks (-80 °C, 6% (w/v) betaine) were revived in 10 mL of AFM1 prepared in 14 mL round bottom polystyrene test tubes (Falcon) incubated at 30 °C and 150 rpm until late-exponential phase was indicated by a reddish-orange appearance with rust-colored precipitation or a pH below 1.5 for iron- and sulfur-oxidizers, respectively, which typically occurred between 10 and 18 d in our hands. Viable "starter" cultures–defined as being stored at 4 °C for $<$ 2 weeks[1]–were used to inoculate fresh media in working cultures (1% (v/v), 100 mL, F2S) in 250 Delong shaker flasks (PYREEX), which were grown at 30 °C and 150 rpm for 120 $\pm$ 4 h to reach the stationary-phase (Figure S43).

For routine culturing of *E. coli DH*5$\alpha$ cells, frozen stocks (-80 °C, 40% glycerol) were revived in 5 mL starter cultures of Lysogeny Broth (LB) in 14 mL round bottom polystyrene test tubes incubated at 37 °C and 200 rpm for 8 h. The starter cultures were added to 150 mL Lysogeny Broth (LB cultures) in 250 mL flasks with 7.5 µg kanamycin and grown for 20 h at 37 °C.

Biological replicates indicate separate starter cultures revived from the same master cryovial of each strain on different days for different durations; each replicate was processes end-to-end under identical conditions. As such, these replicates assess cultivation variability and not lineage variability that may be captured by biological replicates started from separate cryovials or colonies.

#### Text S2. Protocol for cell harvesting and washing.

For the harvesting and washing of *A. ferrooxidans*, cultures were transferred to a sterile 50 mL centrifuge tube and centrifuged to sediment iron and sulfur particles (3,000 x g, 23 s, 4 °C). The supernatant was carefully transferred to a new sterile 50 mL centrifuge tube without disturbing the cell pellet. Cells were pelleted by centrifugation (4,695 x g, 7 min, 4 °C) and the supernatant was poured off as waste. Cell pellets were resuspended in 1 mL of AFM3 BS (1 mL), gently mixed via pippette (8x), and transferred to a 1.5 mL microcentrifuge tube. Residual iron and sulfur particles were sedimented by centrifugation (3,000 x g, 10 s) and a portion of the supernatant (750 µL) was transferred to a new 1.5 mL tube. Cells were washed five times by: centrifugation (17,000 x g, 1 min), aspiration, resuspension in AFM3 BS (1 mL) and gentle mixing (8x), and pelleting (17,000 x g, 1 min).

For the harvesting and washing of *E. coli DH*5$\alpha$ cells, cultures were transferred to a 500 mL bucket and centrifuged (4,000 rpm, 10 min, 20 °C). The pellet was resuspended in 10 mM MES (pH 6) and washed once prior to use.

#### Text S3. Protocol for cell density normalization and quantification.

After washing, cell suspensions were normalized by optical density at 600 nm ($OD_{600}$). Washed pellets were resuspended in the corresponding assay buffer and homogenized by gentle pipetting (8x) to remove visible clump; *A. ferrooxidans* were resuspended in AFM1 BS and *E. coli DH*5$\alpha$ in a 10 mM MES and 10 mM NaCl buffer (pH 6). A 1 mL buffer-only blank was used for background subtraction. A 10 µL portion of the cell resuspension was added to 990 µL of the assay buffer in a 1.5 mL PMMA cuvette (BrandTech), homogenized by gentle pipetting (8x), and measured immediately using a Genesys 10S UV-Vis spectrophotometer (ThermoFisher Scientific). Samples exceeding $OD_{600}$ of 1.0 were diluted further and remeasured. The nominal $OD_{600}$ used was taken as the average of at least five independently prepared and measured samples and these cell pellets were diluted to the target $OD_{600}$ using the relation $C_1 \cdot V_1 = C_2 \cdot V_2$, where $C_1$ and $V_1$ are the measured $OD_{600}$ and volume of the stock suspension and $C_2$ and $V_2$ are the desired $OD_{600}$ and final volume of the pellet not accounting for dilution, respectively. Normalized suspensions were stored at 4 °C for not more than 10 d before use and were mixed immediately before inoculation or use in adsorption assays to minimize settling.

The planktonic cell density of *A. ferrooxidans* was quantified using a SYBR Green I DNA fluorescence assay modified from the techniques the Banta lab has previously reported.[2] A 1 mL aliquot of sample containing cells was transferred to a 1.5 mL tube and centrifuged to remove iron, sulfur, and/or plant tissue solids (3,000 x g, 10 s). A 750 µL aliquot of the supernatant was transferred to a new 1.5 mL tube and centrifuged to pellet the cells (17,000 x g, 1 min). The supernatant was carefully aspirated without disturbing the cell pellet and either disposed of or used for other analyses, including $Fe^{2+}$ titrations and elemental analysis. The pelleted cells were resuspended in 750 mL of TE Buffer (10 mM Tris-HCl, 1 mM EDTA, pH 8.0), vortexed for 10 s, and lysed in a preheated hot plate set to 90 °C for 20 min, which appeared to be around 70 °C based on the reading from a thermometer placed within a hot plate well. After cooling for at least 30 min, the lysed cell suspension was centrifuged to pellet residual iron, sulfur, plant tissues, or cell debris (17,000 x g, 1 min). From the supernatant containing the soluble dsDNS, 200 µL aliquots were transferred to three wells in a black flat bottom polystyrene untreated 96-well plate (Corning). To each well containing samples, 50 µL of a 5X SYBR Green I nucleic acid stain, prepared by adding 20 µL of a 10,000X stock quickly thawed from -20 °C to 40 mL of TE Buffer which was covered in aluminum foil and stored at 4 °C for no more than 10 d, or when the orange hue began to fade, before use, was added, and the plate was covered with aluminum foil and incubated at room temperature for 20 min. The fluorescence intensity was measured with excitation at 497 nm and emission at 520 nm using a SpectraMax M2 plate reader (Molecular Devices) that had not been calibrated for some time.

Standard curves used to correlate the SYBR Green fluorescence intensity to planktonic cell density were obtained by measurements performed in our hands. For each concentration analyzed for each cell stock (n = 5), this process was repeated 3 times and the points shown indicate the mean of these technical replicates.

The fluorescence intensity and $OD_{600}$ were measured for a set of cell samples that were serially diluted from independently cultured and prepared stocks. Before each measurement, the cell suspensions were mixed gently via pipette (8x) before transferring a 10 µL portion into 990 µL of AFM1 BS in a 1.5 mL PMMA cuvette and measuring the absorbance at 600 nm immediately against an AFM1 BS blank. Additionally, a 10 µL portion of the cell suspension was similarly added to 990 µL of AFM1 BS in a 1.5 mL tube and subjected to the SYBR Green analysis procedure described earlier in this section. The data from matched pairs were fit to a power law model of the form fluorescence $= a \cdot OD_{600}^{b}$ (Figure S1a).[2]

In parallel, the $OD_{500}$ of these cell suspension were correlated to $OD_{600}$ by adding 10 µL portion into 990 µL of AFM1 BS in a 1.5 mL PMMA cuvette and measuring the absorbance at 500 nm immediately against an AFM1 BS blank. The data from matched pairs were fit to a straight line model of the form $OD_{600} = m \cdot \mathrm{OD}_{500}$ (Figure S1b).[2]

The relationship between $OD_{500}$ and total number of *A. ferrooxidans* per mL of culture was estimated from the slope of a straight line model of the form $\mathrm{OD}_{500} = m \cdot \rho_{cell}$ fitted to previously reported data[3] extracted using WebPlotDigitizer[4] (Figure S1c). However, this correlation is only used to describe the planktonic fraction of the *A. ferrooxidans* population as this is more representative of what the measurements derived from our specific sampling technique capture.

Together, these correlations were used to estimate the correlation between planktonic cell density and the SYBR Green fluorescence signal (Figure S1d). When needed, $OD_{600}$ was converted to estimate dry cell weight (DCW) using organism-specific conversion factors obtained from the measured mass of 10 mL of OD $OD_{600}$ = 1.0 solutions after lyophilization (Figure S2).

#### Text S4. Protocol for media preparation.

The media formulations used in this study were based on prior research performed in the Banta lab,[1,5,6] which are provided in Table S3. The non-iron or sulfur components were added to approximately 800 mL of ultrapure water in the order listed, before pH adjustment to the indicated setpoint. Upon the addition of iron, the pH was readjusted to the target pH using concentrated $H_2SO_4$ or KOH as needed. The volume of the solution was finalized to 1 L in a volumteric flask before final pH adjustments. The resulting media was then 0.22 µm filtered under flame into sterile containers and stored at 4 °C for no more than 2 weeks prior to use. For the SM4 media, the brown sulfur was added to the filtered media last.

#### Text S5. Protocol for *Phytolacca* cultivation.

*Phytolacca* seed germination, plant growth, and element treatment were carried out following method previously reported by the Doherty lab.[7] *Phytolacca* seeds were stratified by agitation in concentrated $H_2SO_4$ for 5 min and rinsed 5 times with deionized (DI) $H_2O$. The seeds were then surface-sterilized by agitation in 50% (v/v) bleach and 0.02% (v/v) Triton X for 7 min, and rinsed 3 times with DI water. After surface-sterilization, the seeds were kept in sterile DI water in the dark (25°C, 2 d) prior to exposure to 12/12 h light/dark cycles to germinate (7–10 d).

For plants grown in soil, germinating seeds were transferred to soil under 12/12 h light/dark cycles and plants were collected after 2 months of growth in soil.

For plants grown hydroponically in nutrient solution, germinating seeds were transferred to OASIS ®ROOTCUBES ®growing medium (Oasis grower solution, USA) containing 0.1x Hoagland solution (0.5 mM $KNO_3$, 0.5 mM $Ca(NO_3)_2 \cdot 4\,H_2O$, 0.6% Sprint®138 chelated iron, 0.2 mM $MgSO_4 \cdot 7\,H_2O$, 0.1 mM $NH_4NO_3$, 0.2 mM $KH_2PO_4$, 5 µM $H_3BO_3$, 1 µM $MnCl_2 \cdot 4\,H_2O$, 0.08 µM $ZnSO_4 \cdot 7\,H_2O$, 0.03 µM $CuSO_4 \cdot 5\,H_2O$, 0.01 µM $H_3MoO_4 \cdot H_2O$ under 12/12 h light/dark cycles. Once the plants had four fully expanded leaves, the plants were transferred to a 0.25x Hoagland solution. Plants were collected after 2 months of germination.

For hydroponic metal-enrichment, plants were transferred to a 0.25x Hoagland solution after developing 4 fully expanded leaves and supplemented with the selected element solution under 12/12 h light/dark cycles (5–7 d). The combination of elements included: (1) 10 mM of $CeCl_3$, $DyCl_3$, $NdCl_3$, and $TbCl_3$ each, (2) 25 mM of $NiCl_2$, (3) 50 mM of $NiCl_2$, and (4) 10 mM of $CeCl_3$ and $DyCl_3$ each. For the REE-supplemented conditions, $KH_2PO_4$ was excluded from the Hoagland solution to prevent REE precipitation with phosphate.

To generate the dried tissues, plants were harvested, rinsed with DI water, and dried overnight depending on the amount of tissue (70–110 °C). Representative subsamples were prepared by cone-and-quartering,[8] homogenized to approximately 0.5–2 mm passing size using a 4-piece cylindrical herb grinder, frozen overnight at −80 °C, and lyophilized (-80 °C, 0.5 mbar, 1-2 d) before storage or analysis.

#### Text S6. Protocol for digestion of solid samples.

To dissolve the plant tissues for bulk elemental analysis, 200-300 mg of representative subsamples were weighed into a 50 mL polyethylene digestion tube (Environmental Express). A 5 mL aliquot of freshly prepared aqua regia (1:3 v/v concentrated $HNO_3$:HCl) was added to each digestion tube within a fume hood, and the samples were allowed to pre-react at room temperature for 1-2 h. The tubes were loosely capped to vent to evolved gases and heated at 100 °C for 2 h. After cooling to room temperature, the digests were diluted to 50 mL with ultrapure water and filtered through 0.22 µm filters (CELLTREAT) to remove residual solids.

To dissolve bacteria for bulk elemental analysis, cells were resuspended in 500 µL of ultrapure water in 15 mL tubes, covered in perforated parafilm, frozen (-80 °C, >12 h), and lyophilized to constant mass, which typically took 1-2 d in our hands. The mass of the empty tube prior to cell addition was subtracted from that containing the dried cell pellets to determine the cell DCW. An aliquot of approximately 500 µL of trace-metal-grade $HNO_3$ was added to each sample using a polyethylene transfer pipette (Falcon). Loosely capped tubes were added to an aluminum hot block (USA Scientific) that was preheated to 90 °C to digest for 1 h.[9] To evaluate evaporative losses of $HNO_3$, we weighed samples before and after digestion and saw mass changes <1% (n = 40) so no volume corrections were applied. After cooling, ultrapure water was added to bring the total volume to 10 mL and the samples were 0.22 µm filtered prior to ICP-OES measurement, or further diluted to 1% (w/v) $HNO_3$ prior to ICP-MS measurements. Procedural blanks were processed in parallel for baseline subtraction. Cell-associated metal concentrations were determined per dry mass and converted to per cell using cell density and mass correlations described in Text S3 (Figure S2).

#### Text S7. Protocol for elemental analysis.

Elemental concentrations of most leachates and digested solids were measured using inductively coupled plasma optical emission spectroscopy (ICP-OES). Samples were diluted in 2% (w/v) nitric acid prior to analysis. Analyses were performed using a Horiba Ultima Expert (Piscataway, NJ) operating in radial mode. Calibration standards were prepared from a multi-element certified reference material solution (SM68 standard 1, VHG) at concentrations spanning the expected analyte range (0, 0.01, 0.05, 0.1, 0.5, 1, 5, 10, and 25 ppm). The following wavelength were selected to maximize sensitivity and minimize spectral interferences from these analyzed elements: Ca 393.366, K 766.49, Mg 280.271, P 177.433, Fe 259.94, Ni 230.3, Ce 416.561, Dy 338.502, Nd 386.333, and Tb 334.942. Quality controls—including blank subtraction, spike recovery tests (acceptable recovery: 100 ± 10%), analysis of independent check standards (acceptable accuracy: 100 ± 15%), and randomized replicate measurements (acceptable precision: 100 ± 10%)—were performed at least after every 20 experimental samples. Biomass-free leachates and blank acid samples were used as negative controls to account for matrix effects by subtraction under matched conditions, in triplicate, for every sample collected in this study.

For the analysis of ultracentrifugation fractions and the cell-associated metal concentrations, inductively coupled plasma-mass spectrometry (ICP-MS) was conducted using a Perkin Elmer Nexion 2000B ICP-MS with argon carrier gas and helium collision gas using methods developed and validated by the Austin lab (unpublished, Austin lab). This instrument was operated with a RF power of 1600 W and nebulizer, plasma, and collision gas flow rates of 0.98, 15, and 4.5 L·min$^{-1}$, respectively. For sample aquisition, the method used a 25 ms dwell time ·amu$^{-1}$ and a 500 ms integration time. Single element standard solutions were prepared in 1% (w/w) $HNO_3$ concentrations spanning the expected analyte range (0.1-10 ppb Ni, Ce, Dy, Tb; 1-200 ppb Nd). The calibration was confirmed using a continuing calibration verification standard. To check for runoff and contamination, 1% HNO3 rinses were analyzed between every third sample. Data was collected and processed using Syngistix version 3.4. Intensities were averaged from 3 instrumental replicates and corrected using an on-line internal standard containing Rh and In, with 1 reading per replicate and 20 sweeps per reading. Instrumental performance checks verified low oxide and doubly charged ion formation (CeO, $Ce^{++}$ <2.5%) prior to analysis. To confirm the removal of oxides via helium collision, standards containing Tb with Nd, Tb with Ce, and Tb with Dy were compared to the single element standards of the same concentrations.

##### Text S8. Protocol for thermogravimetric analysis.

Thermogravimetric analysis (TGA) was performed on dried tissue samples (ca. 30 mg, ground to ca. 100 µm via mortar and pestle, n = 1–3) under an oxygen atmosphere from 30 to 900 °C at 10 °C·min$^{-1}$. Residual mass fractions and derivative thermogravimetry metrics were extracted from normalized mass-loss curves. Statistical comparisons were performed on extracted thermal metrics rather than point-by-point TGA traces.

##### Text S9. Protocol for calcination at 600 °C.

For calcination experiments, ca. 1 g of mortar-ground dried tissue was heated to 600 °C at 10 °C·min$^{-1}$ in a tube furnace, held for 2 h, and cooled overnight (n = 1–3). Ash residues were analyzed by ICP-OES after aqua-regia digestion. These ash residues (n = 1–3) were also subjected to water-washing tests to assess whether the partitioning of REEs and major inorganic elements between soluble and residual ash fractions provided evidence of REE phosphate mineral decomposition (Text S13).[10] Equations used to calculate enrichment factors and elemental partitioning after thermal treatment are provided in Equations S3–S4 and Equations S15–S22, respectively.

##### Text S10. Protocol for X-ray diffraction on dried and calcined samples.

Dried tissues and calcined ash residues (n = 1) were mounted on the $\phi$10x0.2 mm cavity of a zero diffraction gradient holder and symmetric XRD scans were collected using Cu K$\alpha$ radiation (45 kV, 40 mA) in Bragg–Brentano geometry over 5–90° $2\theta$.[11] The equation used to normalize the XRD spectra of calcined ash residues is provided in Equation S5.

##### Text S11. Protocol for Fourier-transform infrared (FTIR) spectroscopy on dried and calcined samples.

For selected dried tissues, calcined ash residues, and post-leached tissues, samples (n = 1–3) were diluted in KBr (2 mg, 1:100 (w/w))[12] and flushed in $N_2$ (5 min) before Fourier-transform infrared (FTIR) spectra were collected in absorbance mode with a MCT detector from 400 to 4000 cm$^{-1}$ using 128 scans at a 4 cm$^{-1}$ resolution.

Since the full spectra contained broad overlapping matrix features, the raw FTIR spectra were processed within predefined diagnostic regions to support qualitative comparison of local spectral envelopes. For each diagnostic region, candidate low- and high-wavenumber baseline anchor windows were generated within region-specific search intervals. Candidate windows were scored from anchor noise, residual anchor absorbance after baseline correction, negative artifacts, and edge artifacts. In the workflow used for the exported spectra, anchor optimization was performed in global-anchor mode, in which the median spectrum across samples was used to select one low-anchor and one high-anchor window for each region. The selected anchor pair was then applied to all samples within that region to avoid sample-specific anchor choices that could change relative spectral intensities among samples. For each sample and region, the mean absorbance in the selected low- and high-wavenumber anchor windows was used to construct a linear baseline. This baseline was subtracted from the raw regional spectrum to obtain the baseline-corrected absorbance. The corrected regional spectrum was then vector-normalized by its Euclidean norm to compare spectral shape independent of total regional intensity. Baseline-corrected spectra were used for area and intensity metrics, whereas vector-normalized spectra were used only for shape comparison and figure display.

**Text S12. Protocol for scanning electron microscopy with energy dispersive X-ray spectroscopy on plant and bacteria samples.**

Control and Ce/Dy-enriched *P. americana* tissues and *A. ferrooxidans* samples with and without EDTA washing after 24 h incubation with and without soluble Tb were examined by scanning electron microscopy with energy dispersive X-ray spectroscopy (SEM–EDS).

Dried biomass was used for the SEM(-EDS) data collected on carbon tape. Dried plant samples were prepared as described in Text S5. After binding under indicated treatment conditions, bacterial samples were recovered and resuspended in $H_2O$ (about 100 µL; Text S16) before being frozen and lyophilized (Text S5). A small portion of the powder was added to carbon tape fixed on a 180° $\phi 12.7 \times 8$ mm standard SEM pin stub prior to blowing with air to remove loose material and sputter coating (20 s of 40% (w/w) Pd/Au; about 2 nm thickness). Images of the coated samples were collected under constant vacuum (about $9 \times 10^{-6}$ mbar) at varied beam energies (3–7 kV).

To remove the carbon mounting tape as a confounding factor in the EDS analyses, these samples were also mounted on silicon wafers for additional imaging using techniques that were modified from earlier studies on unrelated materials.[13,14] Silicon wafer segments (about $1 \times 1$ cm) were prepared from a 6" diameter, P-type, ⟨100⟩ silicon wafer by scoring with a straight-tip diamond scribe and cleaving with CleanBreak pliers. These wafer segments were placed within a polymer apparatus that held the wafers upright inside a beaker (250 mL). The wafer segments were completely submerged in methanol (100%) inside of a fume hood to prevent lung irritation. The methanol containing the wafer segments was sonicated continuously inside a fume hood using a probe sonicator equipped with a 1/16" diameter tip (10 min, 40% amplitude). The wafer segments were then removed using polymer-coated tweezers, dried using house $N_2$ gas, and attached to stubs using copper tape.

For all plant and bacteria samples, a portion of the dried biomass was added to the mounted wafer segments, $H_2O$ was added dropwise directly onto the solids (20 µL), and the solids were gently resuspended and spread across the wafer surface. These drop-cast samples were dried in an incubator overnight (40 °C) and blown with air prior to imaging. Because these samples had issues with excessive charging during our initial attempts at imaging (data not shown), we therefore sputter coated the samples as described above prior to imaging under constant vacuum (about $9 \times 10^{-6}$ mbar) at varied beam energies (5–10 kV).

For all images, the working distance was indicated on the reported images. EDS spectra and elemental maps were acquired for 180 s per analysis from point locations or defined areas. Data were not processed beyond what the instrument-native Quantax ESPIRIT software performed during acquisition. Analyses included one sample per condition and multiple fields or analysis regions per sample.

**Text S13. Protocol for inorganic residue washing experiments.**

A portion was transferred into a 50 mL centrifuge tubes and water was added with a ratio 10 mL per 10 mg residue to remove soluble salts. Tubes were vortexed for 10 s at the 10 setting and then mixed end-over-end for 40 min at room temperature using a mini tube rotator. Suspensions were centrifuged at 4,695 x g for 7 min at 4 °C to separate solid and liquid phases. A portion of the supernatant was diluted in 2% $HNO_3$ for ICP-OES analysis and the remainder was carefully aspirated and discarded. The residues were resuspended in a fresh aliquot of water and the wash procedure was repeated, generating two wash solution samples and the washed solids. Solids were dried at 30 °C to constant mass before aqua regia digestion (1–2 d in our hands).

#### Text S14. Protocol for BET surface area analysis for solid samples.

Specific surface areas were measured for dried *P. americana* shoots with and without Ce/Dy enrichment, dried *A. ferrooxidans*, and brown and yellow $S^0$.

Dried plant samples were prepared as described in Text S5. Dried bacterial samples were prepared as described in Text S12 $S^0$ samples were analyzed without additional preparation from the respective chemical storage containers.

This surface area analysis method was modified from an approach developed and validated by the Carneiro lab (unpublished, Carneiro lab). Representative subsamples (about 10–200 mg) were loaded into analysis tubes and degassed under vacuum (50 °C, 8 h, $5 \times 10^{-3}$ mmHg). Sample masses used for normalization were determined by subtracting the mass of empty physisorption tubes from the mass of the physisorption tubes after the sample was loading, degassed, and backfilled with $N_2$ gas.

$N_2$ adsorption–desorption isotherms were measured at −196 °C using a gas adsorption analyzer over a relative pressure range of 0–760 mmHg, where relative pressure is the equilibrium pressure divided by the saturation vapor pressure. Specific surface areas were calculated using the Brunauer–Emmett–Teller (BET) method[15] from relative pressure of 0.1 to 0.3, using 5–6 data points and a molecular cross-sectional area of 0.162 $nm^2$ for adsorbed $N_2$. Results were expressed as $m^2$ $g^{-1}$ of dry mass.

#### Text S15. Protocol for solid-liquid extraction experiments.

Unless otherwise specified, leaching tests were conducted in 50 mL centrifuge tubes containing 30 mL of solution and incubated under standard conditions (30 °C, 150 rpm, 7 d, n = 3–6). The initial and final pH and oxidation-reduction potential (ORP) of $H_2SO_4$ *Phytolacca* leachates and corresponding no-plant controls were measured using probes calibrated at least every 12 h, except under high-acid conditions that could damage probes. Initial and final $Fe^{2+}$ concentrations were measured using ferroin cerimetry,[16] where ferroin indicator (10 µL) was added to the diluted supernatant (250 µL into 750 µL $H_2O$) generated by using the two-stage centrifugation method (Text S3) and 0.1 N $Ce(SO_4)_2$ was incrementally added (5–10 µL) while vortexing until observing the color change from red to blue. Total dissolved Fe was measured by ICP-OES and $Fe^{3+}$ was calculated by difference between the total Fe and the $Fe^{2+}$ concentrations. Double-stranded DNA (dsDNA) was quantified using a SYBR Green spectrophotometric technique (Text S3). At each sampling point, 1 mL aliquots were centrifuged (17,000 x g, 1 min) and the supernatants were diluted in 2% $HNO_3$ for ICP-OES analysis (200 µL into 3 mL). Condition-matched no-plant controls were performed in parallel to distinguish plant-derived effects from reagent background or incubation effects. The remaining leachates were stored under benchtop conditions for some time after the initial 7 d experiments prior to post-leach characterization (1–300 d).

$H_2O$ was used to quantify the labile metal pool. Selected acids ($H_2SO_4$, $HNO_3$), chelators (citric acid, $Na_2EDTA$), and inorganic salts ($Fe_2(SO_4)_3$, $K_2SO_4$, $KNO_3$, $(NH_4)_2SO_4$) were used to test the effects of acidity, ligand chemistry, ferric iron, ionic strength, and background electrolyte composition on metal release. Tissue loadings ranged from 0.01–2% (w/v), and reagent concentrations ranged from 1 µM to 1 M depending on the objective of each experiment and reagent solubility. Reagent intensity was normalized to the nominal reactant-to-metal molar ratio calculated from the reagent concentration, biomass loading, and solution volume and the maximum amount of metal available for release determined from the metal concentrations measured for the bulk dried *Phytolacca* source material. This normalization was used as an operational feed-basis metric and does not imply the true reaction stoichiometry. The equation used to calculate nominal reactant-to-metal molar ratios is provided in Equations S23–S25. Nominal reagent-to-Ni molar ratios were calculated using the measured Ni concentration of the corresponding shoot material (Table S4). For some chelator-assisted extraction experiments, fixed-dose citric acid or $Na_2EDTA$ solutions were adjusted to defined pH values and supplemented with $KNO_3$ background electrolyte. Assuming that EDTA has a higher affinity for $In^{3+}$ than the supplemented REEs within the *Phytolacca* tissues[17,18] and that $In^{3+}$ binds strongly to citrate,[19] additional fixed-dose chelator solutions were pre-loaded with $In_2(SO_4)_3$ to test whether apparent ligand availability influenced REE release. Equations used to calculate the release, or solubilization, of target metal are provided in Equation S24 and Equations S26–S29.

The apparent solubilization values reported in this study were calculated using a cumulative mass-balance approach that accounted for the progressive decrease in liquid volume caused by serial sampling from the closed system (Equation S24). Since each leaching test began with 30 mL of solution and 1 mL aliquots were removed at each sampling point, the dissolved mass remaining in the reactor was evaluated using the residual liquid volume at that time point, and the mass of analyte removed in prior aliquots was included to estimate the cumulative amount released from the biomass. This approach avoids underestimating solubilization during time-course experiments and provides a consistent operational basis for comparing metal release across varied conditions. However, these values are reported as apparent solubilization because the calculation assumes that: (i) the elemental composition each sampling aliquot was representative of the bulk liquid phase, (ii) the nominal bulk tissue concentration measured from homogenized bulk *Phytolacca* tissues was consistently representative of all subsamples used in the experiments, (iii) and no *Phytolacca* tissues were removed from the system. The validity of the first two assumptions were not verified in this study, and we acknowledge the third was not true for all samples for all experiments due to unavoidable tissue removal in some cases. The uncertainty introduced from these assumptions become more impactful when cumulative aliquot removal become appreciable relative to the total test volume, changing the solution-to-solid ratio and reagent inventory. Additionally, these solubilization values are denoted as apparent since they were calculated by subtraction from the feed rather than from a full mass balance, which would have required elemental analysis of solid residues that was not routinely performed in this study. The variance observed across biological replicates in the leaching tests inherently includes this source of error alongside other experimental and biological variation. The calculated values are best interpreted as feed-basis, aliquot-corrected operational estimates of cumulative analyte release rather than exact fractions of the total tissue-associated analyte pool, which is reflected in the language used in claims herein.

##### Text S16. Protocol for REE binding and elution experiments.

Early-stationary phase cells were washed (Text S2) and normalized by $OD_{600}$ (Text S3) before a portion of this washed cell pellet was resuspended in acidic REE-sulfate simulant solutions or plant leachates to achieve the target $OD_{600}$ (1:100 (v/v)). Binding assays were performed in (micro)centrifuge tubes for $\leq$ 9 h under standard conditions unless otherwise noted (30 °C, 150 rpm, n = 3–9). After incubation, cells were pelleted by centrifugation (4,695 x g, 4 °C, 7 min), and the supernatant was diluted in 2% $HNO_3$ for ICP-OES analysis. Additional $Tb^{3+}$ binding tests were performed with *E. coli* DH5$\alpha$ under less acidic conditions (pH 6, 10 mM MES, 10 mM NaCl, $TbCl_3$)[20,21] to contextualize the pH and matrix dependence of REE binding. Similarly, the washed cell pellet were resuspended in the binding simulant (1 mL) within microcentrifuge tubes and mixed end-over-end via tube rotator (20, 30 min) prior to centrifugation (17,000 x g, 15 min) and metal analysis on the diluted supernatant.

For *A. ferrooxidans*, $Tb^{3+}$ binding and elution were also tested over repeated sorption–desorption cycles. After binding for up to 24 h, cells were pelleted by centrifugation and $Tb^{3+}$-depleted solutions were removed using a serological pipette (n = 6). The $Tb^{3+}$-enriched cells were re-suspended in citric acid (1:1 (v/v), 0.25 M, pH 3.5) and incubated under identical conditions. After 3 loading-desorption cycles, the *A. ferrooxidans* pellets from biological replicates were combined, resuspended in wash solution (1 mL, 0.1 M $Na_2$EDTA, 0.1 M NaCl), transferred into a microcentrifuge tube, and centrifuged (17,000 x g, 1 min). The wash solution was removed, and the cells were washed once again with wash solution before two rounds of $H_2O$ washing prior to lyophilization, digestion, and ICP-OES to quantify the concentration of cell-associated Tb.

The inferred binding capacity, referred to as binding capacity, was calculated as the difference between dissolved metal concentrations in no-cell controls and cell-containing treatments, and selected cell pellets were digested to compare inferred and measured cell-associated REE concentrations. Equations used to calculate the binding and elution parameters are provided in Equations S46–S51. The percentage of inferred prewash metal retained after washing and the wash-stage mass-balance closure were calculated using Equation S52 and Equation S53, respectively. For pellets washed after citrate elution, the inventory entering the washing sequence was determined using Equation S54. Readers who are interested in these measurements should consider the additional context for the EDTA washing methods provided in Text S25.

To estimate the specific surface area of *A. ferrooxidans*, the external surface area of *A. ferrooxidans* normalized to DCW was estimated by combining published cell-geometry measurements with the cell-number and dry-mass conversions used in this study. For *A. ferrooxidans* ATCC 23270 grown on $Fe^{2+}$, the average cell volume and spherical-equivalent surface area were measured as $0.31 \pm 0.01$ µm$^3$ and $2.1 \pm 0.05$ µm$^2$, respectively, in an earlier study.[22] The spherical-equivalent surface area was multiplied by 1.2 to account for the approximately 1:4 aspect ratio of the rod-shaped cells[22] to yield the rod-shaped surface area estimate of

$2.5 \times 10^{-12}$ $m^2 \cdot cell^{-1}$. The number of cells per unit suspension volume was estimated to be $4.3 \times 10^9$ cells $\cdot$ $mL^{-1}$ $\cdot$ unit $OD_{600}$ using the relationship between $OD_{600}$ and $OD_{500}$ measured in this study (Figure S1b) and the correlation between $OD_{500}$ and cell number obtained by fitting previously published data (Figure S1c).[3] The dry mass per *A. ferrooxidans* cell was estimated to be $1.1 \times 10^{-13}$ g $\cdot$ $cell^{-1}$ by dividing the DCW conversion measured in this study (Figure S2; Text S3) by the estimated number of cells per unit suspension volume. The specific surface area of the *A. ferrooxidans* was therefore estimated to be 22 $m^2 \cdot g^{-1}$ by dividing the rod-shaped surface area estimate by this dry mass per *A. ferrooxidans* cell estimate. This calculation assumes that the published cell geometry[22] and cell-number conversion[3] are applicable to the biomass examined herein, which was not verified.

For *E. coli*, a specific surface area of 19 $m^2 \cdot g^{-1}$ DCW was estimated by modeling cells as spherocylinders with an assumed total length of 2.0 µm and diameter of 0.8 µm and an intracellular dry-mass density of approximately 0.30 pg $\cdot$ $µm^{-3}$ reported for other *E. coli* strains.[23]

#### Text S17. Protocol for membrane fluorescence and growth assays performed on pretreated *A. ferrooxidans*.

Protocols to assess some membrane-associated responses of *A. ferrooxidans* to heat, UV-C, and coupled detergent and sonication pretreatment conditions prior to REE binding were adapted from a previous study from the Banta lab.[24]

Cells were grown to the stationary phase on F2S ($n = 3$; Text S1) and harvested and washed (Text S2). Washed cells were resuspended in AFM3 BS within microcentrifuge tubes (1 mL, $OD_{600}$ = 10.0). The suspensions were repeatedly (3×) pelleted by centrifugation (17,000 × g, 1 min), aspirated, and resuspended in 2:2 BS (1 mL).[24] The resulting resuspensions were $OD_{600}$-normalized (1.5 mL, $OD_{600}$ = 5.0),[24] and aliquoted into 4 separate microcentrifuge tubes prior to pretreatment (300 µL).

Pretreatment conditions were each incubated statically for the same duration (1 h). To establish a baseline for REE binding by *A. ferrooxidans*, untreated cells (AF) were incubated statically and capped room temperature. Heat treatment was used as a putative cell lysis[2] and metabolic inactivation[25] pretreatment to investigate the resulting effects on REE binding. Heat-treated cells (+ heat) were incubated in capped microcentrifuge tubes in a heating block (90 °C, 1 h). UV-C irradiation was selected as a putative metabolic inactivation pretreatment targeting nucleic acids[26] without cell lysis. UV-C-irradiated cells (+ UV-C) were exposed to short-wave ultraviolet light (254 nm) from a source placed directly above the uncapped microcentrifuge tubes (1 in). A UV-protective shield was placed around the irradiation apparatus as a safety measure to mitigate the risks of potential UV-C overexposure. A combined detergent and sonication treatment was included as a candidate membrane-disruption control[27,28] for the fluorescent assays, and was not used in subsequent REE binding experiments. Detergent- and sonication-treated cells (+ Triton X-100) were supplemented with about 1% (v/v) Triton X-100 (30 µL of about 10% in $H_2O$ stock), capped, and initially incubated on ice (30 min). After incubation on ice, the microcentrifuge tubes were subjected to water-bath sonication (5 min) and then incubated at room temperature for the remainder of the treatment period. For each of these four pretreatment conditions, medium-matched no-cell controls were subjected to the same conditions in parallel ($n = 3$). All sample aliquots were then capped and statically incubated at room temperature prior to measurements (about 2 h).

Stocks of propidium iodide (2.8 mM), NBD-PEN (5 mM), and $DiBAC_4(3)$ (30 mM) were prepared in DMSO, aliquoted, covered in foil for light protection, and frozen (-20 °C). Immediately before measurements, fresh working solutions (5× concentrations) of propidium iodide (150 µM), NBD-PEN (5 µM), and $DiBAC_4(3)$ (125 µM) were prepared in $H_2O$ from thawed aliquots.[24]

Cell suspensions were diluted in 2:2 BS to the cell density used in the corresponding REE binding experiments ($OD_{600} = 0.1$). These diluted suspensions were added to three wells within a black, flat-bottom untreated polystyrene 96-well plate (200 µL), and then the working solutions were added (50 µL) before immediately covering these plates for static incubation at room temperature.[24] For each condition, both the cell-containing and cell-free resuspensions were measured with both working solution and the respective DMSO-matched solution which did not contain the probe (50 µL). Technical triplicates from the three corresponding well measurements were averaged within each biological replicate. For each pretreatment, the mean fluorescence of the corresponding no-probe control was subtracted separately from the probe-containing cell and no-cell measurements to account for autofluorescence. The corrected no-cell signal was then subtracted from the corrected cell-containing signal to obtain the reported baseline-corrected fluorescence values.

For the propidium iodide cell permeability assay, the plates were incubated under foil (30 min) before measuring the fluorescence (excitation: 535 nm, emission: 617 nm).[24] For the NBD-PEN lipid-derived free radical assay,[29] the plates were incubated under foil (10 min) before measuring the fluorescence (excitation: 470 nm, emission: 530 nm).[24] For the $DiBAC_4(3)$ membrane-potential-sensitive fluorescence assay, the plates were incubated under foil (10 min) before measuring the fluorescence (excitation: 490 nm, emission: 516 nm).[24]

To evaluate the impact of these pretreatment conditions on the (apparent) rate of energy-substrate oxidation and planktonic cell growth, the treated cell suspensions were used in subsequent growth experiments ($n$ = 3 per condition). The $OD_{600}$-normalized cell suspensions which had not been subjected to the fluorescent staining procedures were inoculated into test tubes containing AFM1 or SM4 media ($OD_{600} = 0.02$, 10 mL) and incubated (30 °C, 150 rpm, 7–14 d), with no-cell abiotic controls performed in parallel. The initial pH of this SM4 media was about 1.8, and $H_2O$ was added to compensate for evaporation prior to each measurement. For the cultures in AFM1, an aliquot (250 µL) was removed at each sample for ferroin cerimetry (Text S15). For the cultures in SM4, the pH was measured. Endpoint planktonic cell density measurements were performed on all cultures (Text S3)

#### Text S18. Protocol for *A. ferrooxidans* time-course culture measurements.

Cultures were sampled at defined intervals, typically every 6-12 h or 12-36 h for iron or sulfur growth media, respectively. Evaporative looses were quantified gravimetrically and compensated with sterile $H_2O$ to maintain ionic strength and acidity. At each time point, the pH, redox potential (Eh), and conductivity measured and an aliquot of the culture (1 mL) was removed for further analysis of culture conditions. The aliquot was transferred into a microcentrifuge tube and centrifuged to sediment sulfur and plant solids (3,000 x g, 10 s). Some of the supernatant (750 µL) was transferred to another microcentrifuge tube and the cells were pelleted via centrifugation (17,000 x g, 1 min). The supernatant was aspirated and diluted in 2% $HNO_3$ for elemental analysis (Text S7) and in $H_2O$ for the $Fe^{2+}$ tiration (Text S15). The remaining cell sediment was re-suspended in TE buffer (750 µL; 1 mM EDTA, 10 mM Tris, pH 8.0) and processed with the same lysis and quantification workflow as for planktonic cells described in Text S3.

#### Text S19. Protocol for NaOH–HCl buffering capacity titrations.

After 0.22 µm filtration, a 2 mL portion of leachate was added to 6 mL water within a 100 mL beaker (PYREEX) equipped with a 0.5" hexagonal ribbed stir bar (Fisherbrand) set to stir atop a PC-420D stir plate (Corning) at 300 rpm. The diluted leachate was titrated by incremental addition of 0.001, 0.01, or 0.1 M NaOH until the solution reached a pH of approximately 9. The same nominal concentrations of HCl were then used to back-titrate the solution to a pH of approximately 3. Base- and acid-addition data were processed separately. NaOH and HCl additions were assigned signs of +1 and −1, respectively, so that the processed profiles retained the direction of acid or base addition. The cumulative equivalents were normalized to both the diluted mixture volume and the original leachate volume; the original-leachate-normalized value was used for interpolation and derivative calculations. For each replicate and branch, original-leachate-normalized cumulative equivalents were linearly interpolated onto a 0.1-pH-unit grid from pH 3.0 to 9.0. Only grid points within the measured pH range of that replicate and branch were retained; no extrapolation was performed. Apparent buffering profiles were calculated as the finite-difference derivative of original-leachate-normalized cumulative equivalents with respect to pH. Forward and backward one-sided differences were used at the first and last available pH-grid points, respectively, and central differences were used at internal pH-grid points. The derivative of the apparent buffering profile with respect to pH was calculated using the same one-sided and central finite-difference scheme.

#### Text S20. Protocol for centrifugal ultrafiltration experiments.

For the centrifugal ultrafiltration experiments, the empty filter device, filtrate collection tube, and concentrate collection tube were weighed before use. The loaded device was reweighed after sample addition, and the filtrate and recovered concentrate tubes were centrifuged (4,000 x g, 10 min, 23 °C) in a swinging bucket rotor until an intended retentate volume of 1000 µL was reached, which was selected to preserve sufficient volume for subsequent analyses in each fraction. Retained material was recovered immediately by inverting the device into the concentrate collection tube and centrifuging (1,000 x g, 2 min, 23 °C). For the subsequent analyses performed on each fraction, percent concentrate recovery, percent filtrate recovery, and total recovery were calculated by direct measurements of fraction weights and analyte concentrations. Filtrate, concentrate, and unfractionated leachate fractions were analyzed by ICP-MS (Text S7) to determine metal recovery and partitioning across the cutoff. For each experimental replicate (n = 3), UV absorbance spectra from 220 to 400 nm were measured in triplicate for 50 µL aliquots in UV-transparent 96-well plate as a semiquantitative proxy for chromophoric dissolved organic matter, with $A_{254}$ and the integrated 220–400 nm absorbance area ($AUC_{220-400}$) used as primary UV metrics.[30,31] Measured values were scatter-corrected by subtraction of the absorbance at 750 nm ($A_{750}$) and background corrected by subtracting the corresponding no-biomass matched control in parallel.

#### Text S21. Protocols for screening, bioleaching, cell recovery, and coproduct separation workflows.

For the screening tests, cells were inoculated into 10 mL of AFM1 or SM4 media containing 0—30% (w/v) shoot tissue at an initial $OD_{600}$ of 0.1 and incubated (30 °C, 150 rpm, 14 d), with no-cell abiotic controls performed in parallel. Tolerance was assessed from apparent substrate oxidation relative to no-*Phytolacca* controls, using $Fe^{2+}$ oxidation for iron-grown cultures and pH change for sulfur-grown cultures.

At the bioleaching endpoint, the remaining cell cultures were transferred into 50 mL centrifuge tubes and centrifuged to separate bulk sulfur and *P. americana* solids from the bulk solution (3,000 x g, 23 s, 4 °C). The supernatant was transferred into a new 50 mL tube and centrifuged (4,695 x g, 7 min, 4 °C) before the putative REE-enriched cells were prepared for elemental analysis using the same methods as the REE-binding tests using ICP-MS (Text S7).

For some *P. acinosa* shoots leachates, we tested the feasibility of co-product separation between the putative REE-enriched *A. ferrooxidans*, the REE-depleted plant tissues, and the aqueous REE leachate (Equations S87–S94). The same two-stage centrifugation approach used to recover *A. ferrooxidans* from bioleaching cultures for elemental analysis was used. The 50 mL tubes containing the wet REE-depleted plant tissues were lyophilized to constant mass and the overall recovery of plant tissues was calculated by dividing the lyophilized mass of the solid residues by the initial mass of the tissues added to the cultures. At each stage, the recovery of *A. ferrooxidans* was calculated by dividing the product of the SYBR Green fluorescence signal of the supernatant and the supernatant volume by product of the corresponding values prior to centrifugation. The overall recovery of the process was calculated as the product of the recovery at each stage. The recovery of REEs in the aqueous leachate was calculated analogously to the *A. ferrooxidans* recovery for each stage and overall.

#### Text S22. Protocol for $S^0$ adsorption experiments.

For $S^0$ adsorption experiments, brown and yellow $S^0$ suspensions (1 g/L) were prepared in bulk in SM4 BS without citric acid (2×, pH 2.2; Table S3). The suspensions were vortexed before an aliquot (0.5 mL) was added an aliquot of $Ce^{3+}$ or $Dy^{3+}$ stock solution (0.5 mL, 2×, pH 1.8) in a 1.5 mL microcentrifuge tube to acheive a final pH of 1.95–2.03. The two cations were tested separately using sulfate salts (Table S1) at nominally similar initial molar concentrations. Stock solutions and media were adjusted to the indicated pH using $H_2SO_4$.

In addition to the test conditions, controls containing medium and ions without $S^0$, medium and $S^0$ without added ions, and medium without $S^0$ or added ions were performed in parallel to account for non-$S^0$ effects on the concentration measurements ($n = 3$).

Tubes were mixed end-over-end using a mini tube rotator (setting 20, room temperature, up to 7 d). For time-course measurements at the lowest and highest concentrations tested for each cation and sulfur type, independent test and control experiments were performed and analyzed.

After incubation, tubes were centrifuged to separate the solid $S^0$ particles from the aqueous phase (17,000 x g, 15 min), and the supernatants were diluted in 2% (w/v) $HNO_3$ for ICP-OES analysis (Text S7). These centrifugation conditions were selected because the supernatant count rate of the baseline brown $S^0$ condition measured by dynamic light scattering after centrifugation was not distinguishable from the corresponding no-brown-$S^0$ medium control, whereas the suspension before centrifugation produced a notable signal (Banta lab, unpublished). Equivalent particle removal were assumed but not validated for yellow $S^0$ using the same centrifugation conditions.

Equations used for control correction and calculation of apparent adsorption, recovery, (surface-area-normalized) distribution coefficients, and Dy/Ce selectivity are provided in Equations S32–S34 and Equations S95–S101.

**Text S23. Protocol for comparison of REE solubilization and plant elemental composition across studies.**

Experimental records from this study were combined with published plant-composition and solubilization data from some other papers.[32–37] Numerical tables were transcribed, and values reported only graphically were extracted using digital tools.[4] Direct aqueous extraction, residual-solid depletion, resin-assisted mobilization, and extraction after thermal treatment were distinguished according to their analytical endpoints. For data from this study, only the 7 d standard solid–liquid extraction endpoint data were included. Reported preparation concentrations took precedence over pH-derived estimates. Where only initial or explicitly stated acid pH was available, ideal acid concentrations were estimated from pH (Equations S102–S104).

**Text S24. Descriptions of conditions used for each study in the REE solubilization and plant elemental composition comparison.**

**This study.** Details for the preparation of materials and solid–liquid extraction tests can be found in this article and its Supporting Information.

**Chour et al.: direct extraction from field-collected fern biomass.**[34] This study used *D. linearis* leaves and stems collected from a former REE mine in Jiangxi, China. Biomass was dried at (70 °C, 24 h) and blade-ground (median particle size of 369 $\mu$m). Extraction used dry biomass in solution using an unspecified extraction temperature and stirring speed (3 g, 100 mL, 2 h). Reactants included $H_2O$, $H_2SO_4$ (0.25 M), citric acid (1 M; pH 1.6, 4.0, 8.0), and $Na_2$EDTA (0.05 M, pH 4.5) Solids were separated using 0.45 $\mu$m cellulose filters. Microwave digestion (0.1 g in 5 mL concentrated $HNO_3$, 200 °C, 40 min) was used to generate liquid samples for elemental analysis.

**Chour et al.: nitric-acid extraction with and without resin.**[32] This study used *D. dichotoma* leaves and stems collected in Jiangxi during 2015 and 2016. Samples were dried at (70 °C, 24 h) and blade-ground. Solid–liquid extraction was performed under standard conditions (3 g biomass in 100 mL, 5 h, 200 rom, ambient temperature) with or without Amberlite IRN77 cation-exchange resin (25 mL). $HNO_3$ was used as the reactant (0.5 M). Solid assays used microwave digestion (about 0.3 g sample in 8.5 mL $HNO_3$ and 1.5 mL $H_2O_2$, 200 °C, 50 min), dilution (to 50 mL), and filtration before ICP-AES analysis.

**Li et al.: extraction from experimentally enriched grass leaves.**[36] This study used commercial grass seeds were grown (14 d) in fertilized potting mix under natural light (25 ± 3 °C, periodically watered (every 2 d) with 50 mg $L^{-1}$ each Y, La, Ce, and Dy, supplied as nitrate salts.[36] Leaves were harvested after 14 d, rinsed with deionized water, dried at 80 °C for 48 h, and ground predominantly to 53–297 $\mu$m. The material therefore represents experimentally enriched, substrate-grown leaves; it was not reported as a field-collected stem-and-leaf mixture. Solid–liquid extraction was performed under standard conditions using $HNO_3$, HCl, and $H_2SO_4$ as reactants (0.5 M acid, 5 wt% solids, 100 mL, 25 °C, 30 min, 450 rpm magnetic stirring, $n = 3$). $H_2SO_4$ was also tested under varied solids at fixed acid concentration (1, 2, 5, and 10 wt% solids; 0.5 M acid) and varied acid concentration at a fixed solids concentration (0.1, 0.5, 1, and 2 M acid; 5 wt% solids). After centrifugation (5000 rpm, 5 min), supernatants were analyzed, and residues were water-washed, dried at (80 °C, 48 h), and digested. Feed and residue digestion used microwave digestion before ICP-MS analysis (0.2 g solid in 12 mL $HNO_3$, 18.5 °C $min^{-1}$ ramp, 40 min at 185 °C). This study calculated each elemental extraction fraction as aqueous REE divided by aqueous plus residual-solid REE. For dose calculations, a solid mass fraction $w_s$ was converted to liquid-to-solid ratio as $(1 - w_s)/(\rho_l w_s)$, taking solution density $\rho_l = 1$ kg $L^{-1}$.

**Xu et al.: raw and thermally treated fern materials.**[37] This study used a *B. orientale* material collected in Guangdong, China, naturally air-dried, and ground to an unspecified particle-size distribution.[37] Raw biomass was compared with furnace-calcined material (550 °C, 3 h, air) and rapidly electrothermally calcined material (1000 °C, 20 s, ambient atmosphere; about 500 mg on carbon paper). Solid–liquid extraction was performed under standard conditions (1 g in 10 mL, 90 °C, 1 h). $H_2SO_4$ concentration was varied (0.001, 0.01, 0.1, 1 M). The Results and Supplementary information specified a 1 M standard condition, whereas the general Methods stated 0.1 M; condition-specific figure and table labels were retained. Total REE assays were obtained by microwave digestion (250 mg biomass in 10 mL 15 M $HNO_3$, 120 °C, 1 h) followed by filtration and ICP-OES analysis. This study calculates solubilization using the initial solids REE concentrations.

**Text S25. Theoretical and experimental considerations for interpretation of cell-associated metal concentration data.**

Sequential chelation and rinsing were designed to remove freely entrained Tb and challenge externally accessible, exchangeable Tb associated with *A. ferrooxidans.* The procedure uses two fresh 1 mL washes containing 0.1 M $Na_2EDTA$ and 0.1 M NaCl at nominal pH 4.5, followed by two isovolumetric $H_2O$ rinses.

Much lower concentrations of EDTA have been used to distinguish metal fractions which were putatively surface-sorbed (1× wash using $1-2\times10^{-4}$ M EDTA, $20-50$ mM Tris, 0.5 M sorbital, pH 7.5;[38,39] 3× wash using $1\times10^{-3}$ M EDTA in PBS[40]). Much lower concentrations of EDTA have also been validated as a method to remove EPS entirely rather than only metals bound therein (1× wash using $2.7\times10^{-2}$ M EDTA).[41] Additionally, much lower EDTA concentrations have been sufficient to redissolve acid-recalcitrant co-precipitated extracellular metals ($5\times10^{-2}$ M EDTA);[42] Importantly, the duration of this protocol (24 h) is much longer than what was used herein and the stability constant for the EDTA-$Fe^{3+}$ complex is much higher than those expected from any trivalent lanthanide. As such, it is plausible that the washing protocol used herein could redissolve acid-recalcitrant co-precipitated extracellular metals under some conditions. There is also precedent for inferring intracellular metal accumulation from EDTA-washed bacterial pellets for several metals,[43–45] including a REE,[46] under milder chemical intensities than what was used herein.

For each single incubation containing about 12.2 mM Tb in the wash-validation experiments (Figure S31; Figure S32), the total supplied Tb inventory in 5.02 mL was about 61 µmol. The EDTA supplied in each 1 mL wash aliquot was about 100 µmol. Assuming 1:1 Tb−EDTA complexation,[47,48] the first wash therefore supplied about 1.6 molar equivalents of EDTA relative to the entire Tb inventory of one incubation. In practice, aspiration of the supernatant prior to washing would be expected to reduce the Tb presented to the wash to the pellet-associated inventory plus entrained solution. Even the hypothetical transfer of the entire incubation Tb inventory into the first wash would leave about 38.8 µmol of EDTA before accounting for consumption by competing metals. The combined inventory of these competing multivalent cations in the AFM1 BS binding medium (Table S3) is about 41 µmol, which is predominantly is Mg. Thus, 100 µmol EDTA would be about 2.4 µmol below the combined Tb and competing-cation inventory if all these metals complexed quantitatively in a 1:1 ratio.

The concentration-based formation constant for complexation of free $Tb^{3+}$ with fully deprotonated EDTA is denoted by $\beta_{\mathrm{Tb-EDTA}}$. Ionic activity corrections, EDTA protonation, and conditional formation constants were calculated using Equations S105–S111. Here, we adjusted the $\log_{10}\beta_{\mathrm{Tb-EDTA}} = 17.92$ reference at 0.1 M ionic strength[48] using a Davies coefficient of 0.509 for an approximate 25°C activity correction[49] to calculate an estimated $\log_{10}\beta_{\mathrm{Tb-EDTA}} \approx 17.22$ at an ionic strength of 0.4 M, which is expected for the EDTA wash solution we used. Reference EDTA $pK_a$ values at an ionic strength of 0.1 M (2.00, 2.67, 6.16, and 10.26)[50,51] were treated as concentration-based constants and adjusted using the same Davies approximation (1.94, 2.55, 5.99, and 10.03, respectively). The resulting values were about 1.94, 2.55, 5.99, and 10.03. Using nominal pH 4.5 on a hydrogen-ion activity basis yields an estimated conditional formation constant of about $8\times10^{9}$ $M^{-1}$ for Tb−EDTA (Equations S109–S111). For a hypothetical 1 mL wash containing the entire 61.2 µmol Tb inventory and 100 µmol EDTA, with fixed pH 4.5 and ionic strength 0.4 M, no competing metals, and free $Tb^{3+}$ and Tb−EDTA as the only dissolved Tb species, the calculated free $Tb^{3+}$ concentration was low (about $2\times10^{-10}$ M; Equation S112; Equation S113).

The influence of the AFM1 BS medium metal inventory was examined with a simplified competition calculation. The same same activity and protonation corrections were applied to the reference Mg−EDTA formation constant of $\log_{10}\beta_{\mathrm{Mg-EDTA}} = 8.69$.[52] One EDTA equivalent was conservatively allocated to each of the other multivalent cations, together consuming about 0.13 µmol, without assuming that their actual speciation or complexation kinetics had been established. Solving the ligand balance with the complete

Tb and Mg inventories gives about 30.4 µmol uncomplexed EDTA, summed across its protonation states, and 8.32 µmol Mg−EDTA (Equation S112; Equation S114). The corresponding free $Tb^{3+}$ concentration was still low (about $2 \times 10^{-10}$ M; Equation S113). Thus, despite the small stoichiometric deficit obtained by assuming quantitative binding of every multivalent cation, this simplified model predicts substantial uncomplexed ligand because only a fraction of the Mg binds EDTA at the assumed pH.

To assess sensitivity to lanthanide identity, La and Lu were considered as lower- and upper-affinity cases for EDTA complexation using $\log_{10} \beta_{\mathrm{LaEDTA}} = 15.46$ and $\log_{10} \beta_{\mathrm{LuEDTA}} = 19.80$ at an ionic strength of 0.1 M.[48] The same calculations gives estimated conditional formation constants of $3 \times 10^7$ and $6 \times 10^{11}$ $M^{-1}$, respectively, at pH 4.5 and an ionic strength of 0.4 M. For separate hypothetical 61 µmol single-metal inventories supplemented with 100 µmol EDTA in 1 mL, the estimated free-ion concentrations are $6 \times 10^{-8}$ M for $La^{3+}$ and $2 \times 10^{-12}$ M for $Lu^{3+}$ without competing metals, or $7 \times 10^{-8}$ and $3 \times 10^{-12}$ M, respectively, when including the basal-medium competition model. These scenarios indicate favorable suppression of dissolved free-ion concentrations even for the lower-affinity lanthanide case under the stated assumptions. These calculations are illustrative fixed-pH, fixed-ionic-strength scenarios rather than complete speciation predictions for the contacted wash. In particular, transferring the entire incubation metal inventory and its accompanying ions into 1 mL would itself change the wash composition. The Davies approximation does not account for specific interactions in the concentrated wash matrix, and the actual post-contact pH, ionic strength, metal transfer from the incubation, and release of competing metals from biomass were not established. Complexation of protonated EDTA can release protons, so a decrease in wash pH would reduce the conditional affinities assumed here.

For reversibly bound, accessible Tb, suppression of the free-ion concentration by EDTA can favor release from bacterial binding sites and reduce rebinding.[53] However, the calculation does not represent Tb bound to bacterial surface sites, including carboxyl and phosphate groups,[54] or the time required to release Tb from those sites. Consequently, a low calculated free-ion concentration should not be equated with a measured percentage of extracellular Tb removal.

Physical replacement of the liquid provides a removal mechanism independent of the magnitude of the EDTA formation constant. Fresh EDTA renews the ligand supply, while removal of each wash fraction removes previously extracted Tb. Across two distinct sets of binding experiments, the summed Tb recovered in the two EDTA washes, two $H_2O$ rinses, and final pellet digest was about equal to the inferred prewash cell-associated Tb inventory (Figure S31b; Equation S53), and the cell-associated Tb signal in the absence of Tb exposure was low (Figure S43c). The first EDTA wash accounted for most of the extracted Tb, whereas the second EDTA wash and subsequent water rinses contained comparatively small fractions on the plotted scale.

The equilibrium calculation does not establish whether the washing periods were sufficient to remove accessible, exchangeable Tb. For two 15-min EDTA stages, negligible equilibrium retention, and the same first-order rate constant incorporating release and transport in both stages, 99% depletion of the modeled bound exchangeable pool, before accounting for dissolved carryover, would require an observed rate constant of at least 0.154 $\mathrm{min}^{-1}$ (Equations S115–S116). This hypothetical rate corresponds to a half-time no greater than 5 min (Equation S117). The comparatively small amounts of Tb recovered in the second EDTA wash and subsequent water rinses are consistent with diminishing extraction during successive stages, although they do not establish complete removal of extracellular Tb (Figure S31b). The wash-duration experiment directly tested whether extending EDTA exposure materially changed the retained fraction (Equation S52). After 24 h of Tb binding, the standard two-stage EDTA protocol decreased this fraction relative to water-only washing (Table S31). A single 1 min EDTA exposure retained a significantly larger fraction than the standard protocol, whereas no significant differences from the standard protocol were detected for single exposures of 5–60 min (Figure S32; Table S31). Together with the apparent flattening of the retention curve, these results support limited additional extraction during longer EDTA exposure under the tested conditions, but they do not establish the extent of removal of an independently quantified extracellular Tb pool.

Although the wash-duration experiments did not indicate that slow desorption, limited accessibility, or slowly dissolving extracellular (mineral) phases were causing a departure from the single-pool approximation, additional tests were performed to investigate the response of a putatively precipitated Tb pool to the washing protocol. After 24 h of incubation at a nominal 10:1 phosphate-to-Tb ratio, soluble Tb concentrations decreased by $68 \pm 10\%$ relative to the nominal initial concentration ( Figure S33a; Equations S118–S119).

Small white specks, putatively terbium phosphate, were visually observed at the conclusion of these experiments, although the identity of the solid was not determined because insufficient material was available for XRD. The EDTA washing protocol recovered $99.9 \pm 0.1\%$ of the putatively precipitated Tb across the two EDTA washes, two $H_2O$ rinses, and final residue (Figure S33b; Equation S122).

For an initial inventory of 61 µmol Tb and a 68% decrease in soluble Tb (Equation S122), the mean recovery estimate leaves about 0.41 µmol unaccounted for, equivalent to 0.7% of the initial soluble inventory. This unaccounted amount is not itself a measurement of residual precipitate. Using the measured biomass calibration of 0.49 mg DCW $mL^{-1}$ $OD_{600}^{-1}$ (Figure S2), a 5.02 mL incubation at $OD_{600} = 0.1$ corresponds to about 0.25 mg initial DCW. Assigning the entire unaccounted Tb amount to this biomass would therefore correspond to about $2.6 \times 10^5$ mg Tb $kg^{-1}$ initial DCW. This exceeds the measured washed-pellet Tb inventories by about 3–19-fold across the 2–24 h binding experiment (Figure S31). Consequently, this recovery, by itself, does not inherently exclude sufficient extracellular Tb to account for the pellet measurements.

The binding-duration experiment indicated that the proportion of cell-associated Tb resistant to EDTA washing depended on incubation time. The geometric-mean retained fraction significant increased between 2 and 24 h, while no significant difference in inferred binding capacity was detected for the same comparison (Figure S31; Table S30). In the phosphate-precipitated control tested in parallel, EDTA recovery of the putatively precipitated Tb showed no apparent dependence on incubation duration (Figure S33). This contrast is consistent with a time-dependent change in the accessibility or binding state of cell-associated Tb that was not reproduced by the tested phosphate preparation.

Pretreatment experiments provided additional context for interpreting the retained cell-associated Tb. Heat-treated cells exhibited higher inferred Tb binding capacities than untreated cells across three binding–elution cycles, despite little detectable substrate oxidation or growth in subsequent recovery assays (Figure S39; Figure S40; Table S33). Increased propidium iodide and $DiBAC_4(3)$ fluorescence relative to the untreated controls after heating supports altered cellular membrane integrity (Table S34). These heat-treated cells had minimal detectable Tb in either the EDTA wash eluate or the cell-associated solid fraction (data not shown). If the heat-treated cells retained substantially less Tb after washing despite their higher inferred binding, that contrast could be consistent with passive association that did not require subsequent growth or substrate oxidation and with greater accessibility of the associated Tb to elution or washing.

Apparent citrate elution recovery was higher for both heat- and UV-C-treated cells than for untreated cells in later sorption cycles, while recovery by untreated cells decreased between cycles 1 and 3 (Figure S39; Table S33). UV-C-treated cells exhibited delayed signs of metabolism in the recovery assays, no statistically detected increase in propidium iodide fluorescence, and increased $DiBAC_4(3)$ fluorescence relative to untreated cells (Figure S40; Table S34). The absence of a statistically detected propidium iodide difference does not establish unchanged membrane permeability. Together, these observations are compatible with passive Tb association and a possible physiology-dependent contribution to wash-resistant retention, but they do not independently establish the active intracellular lanthanide transport and intracellular storage that has been demonstrated in *Methylorubrum extorquens*.[55]

SEM-EDS provided additional observations for *A. ferrooxidans* prepared with and without Tb exposure, with each exposure condition subjected to either water-only or EDTA washing (Figure S34; Text S12). The apparent Tb response was higher in the Tb-exposed, water-washed preparation than in the corresponding EDTA-washed preparation, whose response was close to those of the unexposed controls in some of the images (Figures S35–S38). This qualitative difference was consistent with removal of material contributing to the signal during EDTA washing, but the spectral overlap prevented its attribution exclusively to Tb.

Many factors could have confounded these SEM–EDS results. An apparent Tb-assigned response was present in all four preparations, predominantly in the spectral region containing overlapping Tb $M_\alpha$ and Mg $K_\alpha$ emissions. Residual medium was one possible source of Mg, although the origin of the shared signal was not established, which could have resulted in false positives during Tb mapping. The samples were sputter-coated before EDS analysis, which can alter or complicate elemental analysis of biological specimens.[56] Complications from the coating could possibly explain the Sc-assigned responses observed in these data which were not expected. EDTA-washing could have released outer-membrane components and periplasmic proteins without cell lysis.[57] The membrane-associated fluorescence assays were not performed after the EDTA washing procedure (Text S17), and therefore do not establish membrane integrity during washing. Accordingly, preservation of cell-shaped objects in the SEM images does not establish an unchanged permeability barrier, and Tb in a wash fraction need not have been exclusively extracellular before washing.

Conversely, wash-resistant Tb could be intracellular, strongly bound externally, occluded in extracellular material, or present in a slowly dissolving phase. Loss of intracellular Tb during washing could cause underestimation of its prewash intracellular inventory, whereas retention of extracellular Tb could cause overestimation if the entire pellet measurement were interpreted as intracellular. The net direction and magnitude of these potential biases were not established. Dehydration during lyophilization could have also altered the distribution or retention of Tb or other elements.[58] Accordingly, these SEM–EDS results were interpreted as qualitative observations of Tb occurence and were not used to establish its intracellular localization.

Together, all of these results are compatible with an intracellular contribution to the retained Tb. If accessible extracellular Tb was largely removed and intracellular Tb was largely retained, the protocol could have plausibility provided a reasonable approximation of the apparent intracellular concentrations, which may have even been lower than the true intracellular metal concentrations. Critically, these measurements did not directly verify that the washed-pellet measurements represent intracellular concentrations,[53] so the EDTA-wash measurements are reported operationally as cell-associated metal concentrations.

### Supplementary equations

For all elemental analysis samples, the blank- and dilution-corrected sample concentration of analyte $i$ ($C_{i,\text{sample}}$; mg L$^{-1}$) was calculated as:

$$C_{i,\text{sample}} = (C'_i - C_{i,\text{matrix}} - C_{i,\text{HNO}_3}) \cdot DF \qquad \text{(Equation S1)}$$

where $C'_i$ is the measured analyte concentration in the prepared sample solution (mg L$^{-1}$), $C_{i,\text{matrix}}$ is the analyte concentration measured in the experimental matrix-matched blank control (mg L$^{-1}$), $C_{i,\text{HNO}_3}$ is the analyte concentration measured in the 1–2% $HNO_3$ elemental-analysis matrix (mg L$^{-1}$), and $DF$ is the dilution factor into the 1–2% $HNO_3$ matrix (unitless). For digested solid samples, the dry-mass-normalized bulk concentration of analyte $i$ in the solid material ($C_{i,\text{bulk}}$; mg kg$^{-1}$) was calculated as:

$$C_{i,\text{bulk}} = \frac{C_{i,\text{sample}} \cdot V_{\text{digest}}}{m_{\text{digest}}} \qquad \text{(Equation S2)}$$

where $C_{i,\text{sample}}$ is the blank- and dilution-corrected sample concentration of analyte $i$ (mg L$^{-1}$), $V_{\text{digest}}$ is the final digest volume (L), and $m_{\text{digest}}$ is the dry mass of the solid sample digested (kg).

The enrichment factor for analyte $i$ after thermochemical treatment ($EF_i$; unitless) was calculated as:

$$EF_i = \frac{C_{i,\text{residue}}}{C_{i,\text{dried}}} \qquad \text{(Equation S3)}$$

where $C_{i,\text{residue}}$ is the dry-mass-normalized bulk concentration of analyte $i$ in the thermally generated ash residue (mg kg$^{-1}$), and $C_{i,\text{dried}}$ is the dry-mass-normalized bulk concentration of analyte $i$ in the corresponding untreated tissue (mg kg$^{-1}$). When enrichment was calculated for an analyte ratio, such as REE/P, the same ratio-of-ratios structure was used:

$$EF_{\text{REE/P}} = \frac{(\text{REE/P})_{\text{residue}}}{(\text{REE/P})_{\text{dried}}} \qquad \text{(Equation S4)}$$

where $(\text{REE/P})_{\text{residue}}$ and $(\text{REE/P})_{\text{dried}}$ are the corresponding molar REE/P ratios in the thermally generated ash residue and dried tissue, respectively.

For qualitative comparison across XRD spectra of calcined ash residues, the normalized intensity at diffraction angle $2\theta$ ($I_{\text{norm}}(2\theta)$; unitless) was calculated as:

$$I_{\text{norm}}(2\theta) = \frac{I(2\theta)}{I_{\text{max}}} \qquad \text{(Equation S5)}$$

where $I(2\theta)$ is the raw intensity at diffraction angle $2\theta$, and $I_{\text{max}}$ is the maximum intensity observed within the selected $2\theta$ range for that spectrum. This normalization was used only for visual comparison of diffraction features and does not preserve absolute differences in diffraction intensity among samples or correct for non-peak baseline signals.

For FTIR analysis, each diagnostic region $r$ was selected using a binary window mask:

$$M_r(\nu_i) = \begin{cases} 1, & \nu_{r,\text{min}} \le \nu_i \le \nu_{r,\text{max}} \\ 0, & \nu_i < \nu_{r,\text{min}} \text{ or } \nu_i > \nu_{r,\text{max}} \end{cases} \qquad \text{(Equation S6)}$$

where $\nu_i$ is the wavenumber at point $i$ (cm$^{-1}$), $\nu_{r,\text{min}}$ and $\nu_{r,\text{max}}$ are the lower and upper bounds of diagnostic region $r$ (cm$^{-1}$), and $M_r(\nu_i)$ indicates whether a point was included in the regional analysis. For anchor selection, the median absorbance spectrum across all samples was calculated at each wavenumber:

$$A_{\text{med}}(\nu_i) = \text{median}_s\left[A_s(\nu_i)\right] \qquad \text{(Equation S7)}$$

where $A_s(\nu_i)$ is the raw absorbance of sample $s$ at wavenumber $\nu_i$, and $A_{\text{med}}(\nu_i)$ is the median spectrum used for region-level anchor selection. Low- and high-anchor windows were selected for each diagnostic region by minimizing anchor noise, residual anchor absorbance after baseline subtraction, negative corrected absorbance artifacts, and edge artifacts in the median spectrum. The selected anchor windows were then applied to each sample in the corresponding diagnostic region. For sample $s$ and region $r$, the mean raw absorbances in the selected low- and high-anchor windows were calculated as:

$$\bar{A}_{L,s,r} = \frac{1}{n_L} \sum_{\nu_i \in L_r} A_s(\nu_i) \qquad \text{(Equation S8)}$$

$$\bar{A}_{H,s,r} = \frac{1}{n_H} \sum_{\nu_i \in H_r} A_s(\nu_i) \qquad \text{(Equation S9)}$$

where $\bar{A}_{L,s,r}$ and $\bar{A}_{H,s,r}$ are the sample-specific mean absorbances in the selected low- and high-anchor windows for sample $s$ and region $r$, and $n_L$ and $n_H$ are the number of wavenumber points in the selected low- and high-anchor windows, respectively. The sample- and region-specific linear-baseline slope was calculated as:

$$m_{s,r} = \frac{\bar{A}_{H,s,r} - \bar{A}_{L,s,r}}{\nu_{H,r} - \nu_{L,r}} \qquad \text{(Equation S10)}$$

where $\nu_{L,r}$ and $\nu_{H,r}$ are the midpoint wavenumbers of the selected low- and high-anchor windows, respectively. The local baseline for sample $s$ in region $r$ was calculated as:

$$B_{s,r}(\nu_i) = \bar{A}_{L,s,r} + m_{s,r}\,(\nu_i - \nu_{L,r}) \qquad \text{(Equation S11)}$$

where $B_{s,r}(\nu_i)$ is the local linear baseline at wavenumber $\nu_i$. The baseline-corrected absorbance was calculated as:

$$C_{s,r}(\nu_i) = A_s(\nu_i) - B_{s,r}(\nu_i) \qquad \text{(Equation S12)}$$

where $C_{s,r}(\nu_i)$ is the baseline-corrected absorbance for sample $s$ in region $r$. The Euclidean norm of the baseline-corrected regional spectrum was calculated as:

$$\|C_{s,r}\|_2 = \sqrt{\sum_{\nu_i \in r} C_{s,r}(\nu_i)^2} \qquad \text{(Equation S13)}$$

where $\|C_{s,r}\|_2$ is the vector norm of the baseline-corrected spectrum in region $r$. The vector-normalized absorbance was calculated as:

$$V_{s,r}(\nu_i) = \frac{C_{s,r}(\nu_i)}{\|C_{s,r}\|_2} \qquad \text{(Equation S14)}$$

where $V_{s,r}(\nu_i)$ is the vector-normalized absorbance. Vector-normalized spectra were used only to compare spectral shape and were not used for quantitative integration.

After water washing of calcined residues, the amount of analyte $i$ recovered in wash step $\ell$ ($n_{i,W_\ell}$; mmol) was calculated as:

$$n_{i,W_\ell} = \frac{C_{i,W_\ell} \cdot V_{W_\ell}}{M_i} \qquad \text{(Equation S15)}$$

where $C_{i,W_\ell}$ is the blank- and dilution-corrected concentration of analyte $i$ in wash step $\ell$ (mg L$^{-1}$), $V_{W_\ell}$ is the recovered wash volume (L), and $M_i$ is the atomic or formula mass used for analyte $i$ (mg mmol$^{-1}$). Standard atomic weights or the appropriate formula masses were used consistently for these conversions, typically obtained using Google AI. The total amount of analyte $i$ removed by water washing ($n_{i,\mathrm{water}}$; mmol) was calculated as:

$$n_{i,\mathrm{water}} = \sum_{\ell=1}^{N_W} n_{i,W_\ell} \qquad \text{(Equation S16)}$$

where $N_W$ is the number of sequential wash steps. The initial amount of analyte $i$ in the pre-washed calcined ash residue ($n_{i,\text{initial}}$; mmol) was calculated as:

$$n_{i,\text{initial}} = \frac{C_{i,\text{calcined}} \cdot m_{\text{wash}}}{M_i} \qquad \text{(Equation S17)}$$

where $C_{i,\text{calcined}}$ is the dry-mass-normalized bulk concentration of analyte $i$ in the pre-washed calcined residue (mg kg$^{-1}$), $m_{\text{wash}}$ is the dry mass of calcined residue used in the wash test (kg), and $M_i$ is the atomic or formula mass used for analyte $i$ (mg mmol$^{-1}$). The percentage of analyte $i$ removed by water washing ($R_{i,\text{water}}$; %) was calculated as:

$$R_{i,\text{water}} = 100 \cdot \frac{n_{i,\text{water}}}{n_{i,\text{initial}}} \qquad \text{(Equation S18)}$$

where $n_{i,\text{water}}$ is the total amount of analyte $i$ recovered across all wash steps (mmol), and $n_{i,\text{initial}}$ is the initial amount of analyte $i$ in the washed calcined residue (mmol). The light REE, heavy REE, and total REE amounts were calculated from the measured REE amounts as:

$$n_{\text{LREE}} = n_{\text{Ce}} + n_{\text{Nd}} \qquad \text{(Equation S19)}$$

$$n_{\text{HREE}} = n_{\text{Dy}} + n_{\text{Tb}} \qquad \text{(Equation S20)}$$

$$n_{\text{REE}} = n_{\text{LREE}} + n_{\text{HREE}} \qquad \text{(Equation S21)}$$

respectively. The molar ratio of REE to P (REE/P; unitless) was calculated as:

$$\frac{\text{REE}}{\text{P}} = \frac{n_{\text{REE}}}{n_{\text{P}}} \qquad \text{(Equation S22)}$$

where $n_{\text{REE}}$ is the total measured REE amount (mmol), and $n_{\text{P}}$ is the measured P amount (mmol). A REE/P ratio near unity was used only as an idealized stoichiometric reference for a monazite-like orthophosphate composition and was not interpreted as direct evidence of mineral identity.

For leaching tests, the initial amount of leaching reactant $j$ ($n_{j,0}$; mmol) was calculated as:

$$n_{j,0} = C_j \cdot V_{\text{leach}} \qquad \text{(Equation S23)}$$

where $C_j$ is the leaching reactant concentration (mmol L$^{-1}$), and $V_{\text{leach}}$ is the leaching volume (L). The initial amount of target metal $i$ in the plant tissue ($n_{i,\text{tissue}}$; mmol) was calculated as:

$$n_{i,\text{tissue}} = \frac{C_{i,\text{tissue}} \cdot m_{\text{tissue}}}{M_i} \qquad \text{(Equation S24)}$$

where $C_{i,\text{tissue}}$ is the dry-mass-normalized bulk concentration of target metal $i$ in the tissue (mg kg$^{-1}$), $m_{\text{tissue}}$ is the dry mass of tissue used in the leaching test (kg), and $M_i$ is the atomic or formula mass used for analyte $i$ (mg mmol$^{-1}$). The nominal molar ratio of reactant $j$ to target metal $i$ ($x_{j,i}$; unitless) was then calculated as:

$$x_{j,i} = \frac{n_{j,0}}{n_{i,\text{tissue}}} \qquad \text{(Equation S25)}$$

where $n_{j,0}$ is the initial amount of leaching reactant $j$ (mmol), and $n_{i,\text{tissue}}$ is the initial amount of target metal $i$ in the plant tissue (mmol). The dissolved amount of analyte $i$ in a leachate assumed to be homogeneous ($n_{i,\text{diss}}$; mmol) was calculated as:

$$n_{i,\text{diss}} = \frac{C_{i,\text{leach}} \cdot V_{\text{leach}}}{M_i} \qquad \text{(Equation S26)}$$

where $C_{i,\text{leach}}$ is the blank- and dilution-corrected analyte concentration in the leachate (mg L$^{-1}$), $V_{\text{leach}}$ is the leachate volume (L), and $M_i$ is the atomic or formula mass used for analyte $i$ (mg mmol$^{-1}$). The apparent solubilization of analyte $i$ from the plant tissue ($S_i$; %) was calculated as:

$$S_i = 100 \cdot \frac{n_{i,\text{diss}}}{n_{i,\text{tissue}}} \qquad \text{(Equation S27)}$$

where $n_{i,\text{diss}}$ is the dissolved amount of analyte $i$ measured in the leachate (mmol), and $n_{i,\text{tissue}}$ is the initial amount of analyte $i$ in the plant tissue (mmol). For derived metrics such as LREE, HREE, and REE solubilization, the numerator and denominator were each calculated from the summed amounts of the constituent analytes. For time-course tests in which multiple aliquots were removed, the cumulative dissolved amount of analyte $i$ at time point $t_n$ ($n_{i,\text{diss}}(t_n)$; mmol) was calculated as:

$$n_{i,\text{diss}}(t_n) = \frac{C_{i,n} \cdot [V_{\text{leach}} - (n-1) \cdot V_{\text{sample}}] + V_{\text{sample}} \cdot \sum_{k=1}^{n-1} C_{i,k}}{M_i} \qquad \text{(Equation S28)}$$

where $C_{i,n}$ is the blank- and dilution-corrected concentration of analyte $i$ in the leachate at sampling point $n$ (mg L$^{-1}$), $C_{i,k}$ is the corresponding concentration at prior sampling point $k$ (mg L$^{-1}$), $V_{\text{leach}}$ is the initial leaching volume (L), $V_{\text{sample}}$ is the aliquot volume removed at each prior sampling point (L), and $M_i$ is the atomic or formula mass used for analyte $i$ (mg mmol$^{-1}$). The cumulative apparent solubilization at time point $t_n$ ($S_i(t_n)$; %) was then calculated as:

$$S_i(t_n) = 100 \cdot \frac{n_{i,\text{diss}}(t_n)}{n_{i,\text{tissue}}} \qquad \text{(Equation S29)}$$

where $n_{i,\text{diss}}(t_n)$ is the cumulative dissolved amount of analyte $i$ at time point $t_n$ (mmol), and $n_{i,\text{tissue}}$ is the initial amount of analyte $i$ in the plant tissue (mmol).

For REE comparisons involving different sets of quantified elements, the molar amount of group $G$ in phase or material $p$ ($n_{G,p}$; mmol) was calculated as:

$$n_{G,p} = \sum_{i \in \mathcal{I}_G} n_{i,p} \qquad \text{(Equation S30)}$$

where $\mathcal{I}_G$ is the stated set of quantified LREEs, HREEs, or total REEs for that dataset, and $p$ identifies the feed, aqueous phase, or solid-associated pool. For this study, the group assignments in Equations S19–S21 were retained. For literature datasets, the quantified element sets and any inclusion of Y were retained for each source. The same element sets were used for the feed and aqueous amounts in each within-dataset comparison.

Feed-normalized HREE/LREE solubilization selectivity ($\alpha_{\text{HREE/LREE}}$; unitless), distinct from the distribution-coefficient ratio, was calculated as:

$$\alpha_{\text{HREE/LREE}} = \frac{n_{\text{HREE,aq}}/n_{\text{LREE,aq}}}{n_{\text{HREE,feed}}/n_{\text{LREE,feed}}} \qquad \text{(Equation S31)}$$

The group amounts were calculated using Equation S30. Literature total REE mass concentration is the sum of quantified elemental mass concentrations, whereas molar ratios use the corresponding sums after mass-to-mole conversion. P/REE is the reciprocal of Equation S22, evaluated for the relevant REE set.

The condition-specific apparent solid–liquid distribution coefficient ($K_{\text{d,app},i}$; L kg$^{-1}$ dry reference solid) was calculated as:

$$K_{\text{d,app},i} = \frac{q_{\text{app},i}}{c_{\text{e,app},i}} \qquad \text{(Equation S32)}$$

where $q_{\text{app},i}$ is the apparent solid-associated molar loading (mmol kg$^{-1}$) and $c_{\text{e,app},i}$ is the corresponding corrected aqueous concentration (mmol L$^{-1}$), converted from mass concentration using Equation S101. For plant-leaching endpoints, the loading is the inferred retained loading from Equation S37; for rebinding endpoints, it is the total corrected retained loading from Equation S39, applied separately to each stage for sequential tests. For bacterial binding, the inferred loading from Equation S48 is divided by the elemental molar mass to convert mg kg$^{-1}$ DCW to mmol kg$^{-1}$ DCW; for sulfur, the loading is calculated using Equation S100. For a combined REE group, constituent molar loadings and aqueous molar concentrations are summed separately before taking the ratio.

The surface-area-normalized apparent distribution coefficient ($K^A_{\mathrm{d,app},i}$; L m$^{-2}$) was calculated as:

$$K^A_{\mathrm{d,app},i} = \frac{K_{\mathrm{d,app},i}}{a_{\mathrm{S}}} \qquad \text{(Equation S33)}$$

In this common definition, $a_{\mathrm{S}}$ denotes the specific surface area of the reference solid on the same dry-mass basis as the loading (m$^2$ kg$^{-1}$); values in m$^2$ g$^{-1}$ are multiplied by 1000. For sulfur and dried plant samples, this is the mean BET specific surface area of the corresponding sample preparation, whereas for bacterial comparisons, the reference area is the estimated exterior cell area.

The apparent selectivity for Dy relative to Ce ($\alpha_{\mathrm{app,Dy/Ce}}$; unitless) was calculated as:

$$\alpha_{\mathrm{app,Dy/Ce}} = \frac{K_{\mathrm{d,app,Dy}}}{K_{\mathrm{d,app,Ce}}} \qquad \text{(Equation S34)}$$

The same ratio defines HREE/LREE distribution selectivity when the group coefficients replace the Dy and Ce coefficients, respectivelym according to Equations S19–S21.

For the REE repartitioning experiments, the initial loading of analyte $i$ in the *Phytolacca* tissue before leaching and/or rebinding ($q_{i,0}$; $\mu$mol g$^{-1}$) was calculated from the corresponding dry-mass-normalized bulk tissue concentration as:

$$q_{i,0} = \frac{C_{i,\mathrm{bulk}}}{M_i} \qquad \text{(Equation S35)}$$

where $C_{i,\mathrm{bulk}}$ is the dry-mass-normalized bulk concentration of analyte $i$ in the starting tissue or post-leach residue basis material (mg kg$^{-1}$), and $M_i$ is the atomic or formula mass used for analyte $i$ (mg mmol$^{-1}$). The resulting units are mmol kg$^{-1}$, which are equivalent to $\mu$mol g$^{-1}$. For samples that were leached before rebinding, the apparent amount of analyte $i$ released during the preceding leaching step ($q_{i,\mathrm{release}}$; $\mu$mol g$^{-1}$) was calculated as:

$$q_{i,\mathrm{release}} = q_{i,0} \cdot \frac{S_i}{100} \qquad \text{(Equation S36)}$$

where $q_{i,0}$ is the initial tissue loading of analyte $i$ before leaching and/or rebinding ($\mu$mol g$^{-1}$), and $S_i$ is the apparent solubilization of analyte $i$ during the preceding leaching experiment (%). For direct rebinding tests performed without a preceding leaching step, $q_{i,\mathrm{release}}$ was set to 0 $\mu$mol g$^{-1}$. The apparent amount of analyte $i$ remaining on the plant tissue immediately before the rebinding step ($q_{i,\mathrm{pre,ads}}$; $\mu$mol g$^{-1}$) was calculated as:

$$q_{i,\mathrm{pre,ads}} = q_{i,0} - q_{i,\mathrm{release}} \qquad \text{(Equation S37)}$$

where $q_{i,0}$ is the initial tissue loading of analyte $i$ before leaching and/or rebinding ($\mu$mol g$^{-1}$), and $q_{i,\mathrm{release}}$ is the apparent amount of analyte $i$ released during the preceding leaching step ($\mu$mol g$^{-1}$). To account for solution-chemistry effects and experimental artifacts, condition-matched no-plant controls were performed in parallel. The corrected adsorption of analyte $i$ from the reload solution onto the plant tissue during rebinding ($q_{i,\mathrm{ads,corr}}$; $\mu$mol g$^{-1}$) was calculated as:

$$q_{i,\mathrm{ads,corr}} = \frac{(C_{i,\mathrm{reload},f,\mathrm{no\ plant}} - C_{i,\mathrm{reload},f,\mathrm{plant}}) \cdot V_{\mathrm{reload}}}{m_{\mathrm{tissue,reload}}} \qquad \text{(Equation S38)}$$

where $C_{i,\mathrm{reload},f,\mathrm{no\ plant}}$ is the final molar concentration of analyte $i$ in the condition-matched no-plant control ($\mu$mol L$^{-1}$), $C_{i,\mathrm{reload},f,\mathrm{plant}}$ is the final molar concentration of analyte $i$ after incubation with plant tissue ($\mu$mol L$^{-1}$), $V_{\mathrm{reload}}$ is the reload solution volume (L), and $m_{\mathrm{tissue,reload}}$ is the dry mass of plant tissue used in the rebinding experiment (g). This corrected value was calculated only for tests containing plant tissue. The final corrected loading of analyte $i$ on the plant tissue after rebinding ($q_{i,\mathrm{after}}$; $\mu$mol g$^{-1}$) was calculated as:

$$q_{i,\mathrm{after}} = q_{i,\mathrm{pre,ads}} + q_{i,\mathrm{ads,corr}} \qquad \text{(Equation S39)}$$

where $q_{i,\mathrm{pre,ads}}$ is the apparent loading of analyte $i$ immediately before the rebinding step ($\mu$mol g$^{-1}$), and $q_{i,\mathrm{ads,corr}}$ is the corrected adsorption of analyte $i$ during the rebinding step ($\mu$mol g$^{-1}$). The notation $q_{\mathrm{after,corr}}$ used in the main text denotes the final corrected total REE loading obtained by summing the element-specific final loadings on a molar basis. When the final corrected loading was reported on a dry-mass concentration basis ($C_{i,\text{after}}$; mg kg$^{-1}$), it was calculated as:

$$C_{i,\text{after}} = q_{i,\text{after}} \cdot M_i \qquad \text{(Equation S40)}$$

where $q_{i,\text{after}}$ is the final corrected loading of analyte $i$ after rebinding ($\mu$mol g$^{-1}$, equivalent to mmol kg$^{-1}$), and $M_i$ is the atomic or formula mass used for analyte $i$ (mg mmol$^{-1}$). The operational reload efficiency for analyte $i$ ($R_{i,\text{reload}}$; unitless) was calculated as:

$$R_{i,\text{reload}} = \frac{q_{i,\text{ads,corr}}}{q_{i,\text{release}}} \qquad \text{(Equation S41)}$$

where $q_{i,\text{ads,corr}}$ is the corrected adsorption of analyte $i$ during the rebinding step ($\mu$mol g$^{-1}$), and $q_{i,\text{release}}$ is the apparent amount of analyte $i$ released during the preceding leaching step ($\mu$mol g$^{-1}$). $R_{i,\text{reload}}$ was calculated only for leached samples where $q_{i,\text{release}} > 0$. For sequential first-rebinding, unbinding, and second-rebinding tests, stages were indexed by $s = 1$, 2, and 3, respectively. The aqueous amount accounted for by time $t_n$ within each stage, $n_{i,s,r,\text{aq,accounted}}(t_n)$ (mmol), was calculated using Equation S28, substituting that stage's initial volume, aqueous concentrations, liquid-sample volume, and sampling count. The subscript $r$ identifies the plant-containing experiment or its condition-matched no-plant control; each control uses its own sampling history. For a stage with only an endpoint measurement, the prior-sample sum is zero. The initial aqueous amount, $n_{i,s,r,\text{aq},0}$ (mmol), was calculated using Equation S26 with the stage-specific initial concentration and volume. The stage input includes all liquid present after transfer, including any carried-over liquid. The control-corrected apparent net transfer from the aqueous phase to the plant-associated pool ($\Delta n_{i,s,\text{uptake}}(t_n)$; mmol) is:

$$\Delta n_{i,s,\text{uptake}}(t_n) = [n_{i,s,\text{plant,aq},0} - n_{i,s,\text{plant,aq,accounted}}(t_n)] - \overline{[n_{i,s,\text{control,aq},0} - n_{i,s,\text{control,aq,accounted}}(t_n)]} \qquad \text{(Equation S42)}$$

where the overbar denotes the mean of the matched no-plant controls. Positive values represent apparent net uptake and negative values represent apparent net release. For unsampled stages with identical initial solution inventories and final volumes, the expression reduces to the concentration-difference numerator in Equation S38 after some unit conversion. The signed loading change ($\Delta q_{i,s}(t_n)$; mmol kg$^{-1}$) is:

$$\Delta q_{i,s}(t_n) = \frac{\Delta n_{i,s,\text{uptake}}(t_n)}{m_\text{ref}} \qquad \text{(Equation S43)}$$

where $m_\text{ref}$ is the reference initial dried tissue mass (kg). The within-stage retained loading is then calculated using Equation S39, substituting the stage-start loading and this signed loading change for its pre-adsorption loading and corrected adsorption terms. The first stage starts from Equation S37. Conservation of the inferred plant-associated inventory between consecutive stages was maintained using:

$$q_{i,s+1,\text{start}} = q_{i,s,\text{end}} + \frac{\Delta n_{i,s,\text{transfer}}}{m_\text{ref}} \qquad \text{(Equation S44)}$$

where $q_{i,s,\text{end}}$ is the preceding stage's endpoint retained loading, and $\Delta n_{i,s,\text{transfer}}$ is the signed net change in plant-associated amount during interstage handling (mmol). The retained loading normalized to the reference surface area ($q^A_{i,s,\text{retained}}$; mmol m$^{-2}$) was calculated as:

$$q^A_{i,s,\text{retained}} = \frac{q_{i,s,\text{retained}}}{a_\text{S}} \qquad \text{(Equation S45)}$$

where $a_\text{S}$ has the same reference-area meaning and units as in Equation S33. Group loadings are summed on a molar basis before normalization; endpoint distribution coefficients use the total retained loading and endpoint aqueous concentration in Equation S32.

For bacterial REE binding experiments, the inferred concentration of metal $i$ removed from solution during the binding step ($\Delta C_{i,\text{bound}}$; mg L$^{-1}$) was calculated as:

$$\Delta C_{i,\text{bound}} = C_{i,\text{control}} - C_{i,\text{sup}} \qquad \text{(Equation S46)}$$

where $C_{i,\text{control}}$ is the metal concentration measured in the matched no-bacteria control (mg $L^{-1}$), and $C_{i,\text{sup}}$ is the metal concentration remaining in the supernatant after incubation with bacteria (mg $L^{-1}$). The matched no-bacteria controls were prepared using the same dilution ratio of metal solution and cell-suspension buffer used in the binding tests. The inferred mass of metal $i$ associated with the bacterial pellet ($m_{i,\text{bound}}$; mg) was calculated as:

$$m_{i,\text{bound}} = \Delta C_{i,\text{bound}} \cdot V_{\text{binding}} \qquad \text{(Equation S47)}$$

where $V_{\text{binding}}$ is the total liquid volume used during the binding step (L). The inferred binding capacity normalized to dry cell weight ($q_{i,\text{bound}}$; mg $kg^{-1}$ DCW) was calculated as:

$$q_{i,\text{bound}} = \frac{m_{i,\text{bound}}}{m_{\text{DCW}}} \qquad \text{(Equation S48)}$$

where $m_{\text{DCW}}$ is the dry cell mass present in the binding test based on the $OD_{600}$-to-DCW conversion (kg DCW; Figure S2). For elution experiments, the mass of metal $i$ recovered in elution fraction $k$ ($m_{i,\text{eluted},k}$; mg) was calculated as:

$$m_{i,\text{eluted},k} = C_{i,\text{eluent},k} \cdot V_{\text{eluent},k} \qquad \text{(Equation S49)}$$

where $C_{i,\text{eluent},k}$ is the blank- and dilution-corrected concentration of metal $i$ in elution fraction $k$ (mg $L^{-1}$), and $V_{\text{eluent},k}$ is the recovered eluent volume for that fraction (L). The total mass of metal $i$ recovered across elution fractions ($m_{i,\text{eluted}}$; mg) was calculated as:

$$m_{i,\text{eluted}} = \sum_{k=1}^{N_E} m_{i,\text{eluted},k} \qquad \text{(Equation S50)}$$

where $N_E$ is the number of elution fractions. The elution capacity normalized to dry cell weight ($q_{i,\text{eluted}}$; mg $kg^{-1}$ DCW) was calculated as:

$$q_{i,\text{eluted}} = \frac{m_{i,\text{eluted}}}{m_{\text{DCW}}} \qquad \text{(Equation S51)}$$

where $m_{\text{DCW}}$ is the dry cell mass present in the elution test (kg DCW).

For evaluation of the washing procedures described in Text S25, the mass of metal $i$ retained in the final washed pellet ($m_{i,\text{pellet}}$; mg) was calculated using Equation S78, with the blank- and dilution-corrected digest concentration ($C_{i,\text{digest}}$; mg $L^{-1}$) and final digest volume ($V_{\text{digest}}$; L) as the concentration and volume terms, respectively. Digest concentrations were corrected using Equation S1, and the digest represented the entire recovered pellet. The measured pellet capacity ($q_{i,\text{measured}}$; mg $kg^{-1}$ initial DCW) was calculated using Equation S48, substituting the measured pellet inventory for the inferred bound inventory and retaining the initial dry cell mass ($m_{\text{DCW}}$; kg DCW) estimated from the $OD_{600}$-to-DCW calibration (Figure S2) as the denominator. For pellets washed directly after a single binding incubation, the retained percentage ($R_{i,\text{retained}}$; %) and corresponding unitless retained fraction ($f_{i,\text{retained}}$) were defined as:

$$R_{i,\text{retained}} = 100 \cdot f_{i,\text{retained}} = 100 \cdot \frac{m_{i,\text{pellet}}}{m_{i,\text{bound}}} = 100 \cdot \frac{q_{i,\text{measured}}}{q_{i,\text{inferred}}} \qquad \text{(Equation S52)}$$

where $m_{i,\text{bound}}$ is the inferred prewash inventory calculated using Equation S47, and $q_{i,\text{inferred}}$ denotes the inferred binding capacity defined as $q_{i,\text{bound}}$ in Equation S48. Both capacities assume the same initial dry cell mass, $m_{\text{DCW}}$, as their denominator. The mass of metal $i$ in each recovered wash fraction ($m_{i,\text{wash},k}$; mg) was calculated using Equation S49, with the corresponding wash concentration and recovered wash volume replacing the eluent terms. Wash-stage mass-balance closure ($B_{i,\text{wash}}$; %) was calculated as:

$$B_{i,\text{wash}} = 100 \cdot \frac{m_{i,\text{pellet}} + \sum_{k=1}^{N_W} m_{i,\text{wash},k}}{m_{i,\text{prewash}}} \qquad \text{(Equation S53)}$$

where $N_W$ is the number of collected wash fractions, and $m_{i,\text{prewash}} > 0$ is the estimated metal inventory entering the washing sequence (mg). For the standard protocol, the four fractions were the first and second EDTA washes and the first and second water rinses. For pellets washed directly after binding, the prewash inventory was the inferred bound inventory from Equation S47. Fraction-resolved recoveries were calculated using Equation S55, with the inventory in the corresponding wash or final pellet as the recovered inventory and the prewash inventory as the input inventory. For pellets washed after citrate elution in sorption cycle $c$, the wash-entry inventory accounted for the preceding removals using:

$$m_{i,\text{prewash},c} = m_{i,\text{carry},c} + m_{i,\text{feed},c} - m_{i,\text{sup},c} - m_{i,\text{citrate},c} \quad \text{(Equation S54)}$$

where $m_{i,\text{carry},c}$ is the metal inventory carried into cycle $c$ in the retained solids and liquid (mg), $m_{i,\text{feed},c}$ is the inventory introduced in fresh binding medium (mg), $m_{i,\text{sup},c}$ is the inventory removed in the binding supernatant (mg), and $m_{i,\text{citrate},c}$ is the inventory removed by citrate elution (mg).

Fractional recovery of analyte $i$ in the concentrate or filtrate relative to the starting leachate inventory ($R_{i,c/f}$; %) was calculated as:

$$R_{i,c/f} = 100 \cdot \frac{m_{i,c/f}}{m_{i,0}} \quad \text{(Equation S55)}$$

where $m_{i,0}$, $m_{i,c}$, and $m_{i,f}$ are the masses of analyte $i$ in the starting leachate, concentrate, and filtrate, respectively (mg). For comparisons on a total-REE molar basis, the concentration factor relative to the reference plant material ($CF$; unitless) is defined as:

$$CF = \frac{\sum_{i \in \mathcal{I}_{\text{cell}}} q_{i,\text{cell}}/M_i}{\sum_{j \in \mathcal{I}_{\text{plant}}} q_{j,0}} \quad \text{(Equation S56)}$$

where $q_{i,\text{cell}}$ is the reported bacterial loading (mg kg$^{-1}$ DCW), $M_i$ is the elemental molar mass (mg mmol$^{-1}$), and $q_{j,0}$ is the reference plant molar loading calculated using Equation S35 ($\mu$mol g$^{-1}$, numerically equivalent to mmol kg$^{-1}$). The sets $\mathcal{I}_{\text{cell}}$ and $\mathcal{I}_{\text{plant}}$ identify the REEs included in the bacterial and reference plant totals, respectively. The bacterial loading retains the dry-cell-mass basis used for its measurement or inference. For the Ce/Dy-enriched plant reference, its Tb-equivalent loading ($q_{\text{Tb,eq,plant}}$; mg kg$^{-1}$) is:

$$q_{\text{Tb,eq,plant}} = M_{\text{Tb}} \left(q_{\text{Ce},0} + q_{\text{Dy},0}\right) \quad \text{(Equation S57)}$$

For a Tb-only bacterial loading, the same concentration factor is the ratio of the reported Tb loading to this Tb-equivalent plant reference.

For the geometric surface-area estimates described in Text S16, the exterior area per *A. ferrooxidans* cell ($A_{\text{cell,AF}}$; m$^2$ cell$^{-1}$) was estimated from the published spherical-equivalent area as:

$$A_{\text{cell,AF}} = 1.2 \cdot A_{\text{spherical,AF}} \quad \text{(Equation S58)}$$

where $A_{\text{spherical,AF}}$ is the spherical-equivalent area per cell (m$^2$ cell$^{-1}$), and the factor of 1.2 is the correction used for the approximately 1:4 cell aspect ratio.[22] The estimated dry mass per *A. ferrooxidans* cell ($m_{\text{cell,AF}}$; g cell$^{-1}$) was calculated as:

$$m_{\text{cell,AF}} = \frac{\kappa_{\text{DCW,AF}}}{\kappa_{N,\text{AF}}} \quad \text{(Equation S59)}$$

where $\kappa_{\text{DCW,AF}}$ is the measured dry-mass conversion (g mL$^{-1}$ unit OD$_{600}^{-1}$), and $\kappa_{N,\text{AF}}$ is the cell-number conversion (cells mL$^{-1}$ unit OD$_{600}^{-1}$) obtained using the measured OD relationship and the literature cell-number relationship described in Text S3.[3]

For the *E. coli* estimate, the exterior area per cell ($A_{\text{cell,EC}}$; m$^2$ cell$^{-1}$) and volume per cell ($V_{\text{cell,EC}}$; m$^3$ cell$^{-1}$) were calculated using the assumed spherocylinder geometry as:

$$A_{\text{cell,EC}} = \pi d \ell_{\text{total}} \quad \text{(Equation S60)}$$

$$V_{\text{cell,EC}} = \frac{\pi d^2}{4}\left(\ell_{\text{total}} - d\right) + \frac{\pi d^3}{6} \quad \text{(Equation S61)}$$

where $d$ is the cell diameter and $\ell_{\text{total}}$ is the total end-to-end cell length including the two hemispherical ends, both expressed in m, with $\ell_{\text{total}} \geq d$. The corresponding dry mass per cell ($m_{\text{cell,EC}}$; g cell$^{-1}$) was estimated as:

$$m_{\text{cell,EC}} = \rho_{\text{DCW,EC}} \cdot V_{\text{cell,EC}} \qquad \text{(Equation S62)}$$

where $\rho_{\text{DCW,EC}}$ is the assumed intracellular dry-mass density described in Text S16, expressed in g $m^{-3}$ for use with the volume in $m^3$ $cell^{-1}$.[23]

For either organism, the estimated geometric exterior cell area normalized to dry cell mass ($a_{\text{geometric}}$; $m^2$ $g^{-1}$ DCW) was calculated as:

$$a_{\text{geometric}} = \frac{A_{\text{cell}}}{m_{\text{cell}}} \qquad \text{(Equation S63)}$$

where $A_{\text{cell}}$ and $m_{\text{cell}}$ are the corresponding organism-specific area and dry mass per cell, respectively.

For the membrane-associated fluorescence assays described in Text S17, the baseline-corrected fluorescence ($F_{\text{corr}}$; RFU) was calculated as:

$$F_{\text{corr}} = \left(F_{\text{cell},+} - \overline{F}_{\text{cell},-}\right) - \left(F_{\text{no cell},+} - \overline{F}_{\text{no cell},-}\right) \qquad \text{(Equation S64)}$$

where $+$ denotes the probe-containing measurement and $-$ denotes the corresponding DMSO-matched no-probe control. The cell-containing and no-cell probe signals are the technical-triplicate means, and each overbar denotes the mean of the corresponding no-probe control for the same pretreatment and cell-presence condition. The separate no-probe corrections account for autofluorescence before subtraction of the corrected no-cell signal.

For acid-base titrations, the signed titrant factor ($s_j$; unitless) for measurement step $j$ was defined as:

$$s_j = \begin{cases} +1, & \text{for NaOH} \\ -1, & \text{for HCl} \end{cases} \qquad \text{(Equation S65)}$$

where $s_j$ defines whether the titrant addition corresponds to base addition ($+1$) or acid addition ($-1$). The signed amount of titrant added at step $j$ ($\Delta n_j$; mmol) was calculated as:

$$\Delta n_j = C_{\text{tit},j} \cdot V_j \cdot s_j \qquad \text{(Equation S66)}$$

where $C_{\text{tit},j}$ is the titrant concentration at step $j$ (mmol $L^{-1}$), $V_j$ is the incremental titrant volume (L), and $s_j$ is the signed titrant factor. Cumulative signed titrant equivalents were calculated independently within each titration branch as:

$$n_{\text{cum},j} = \sum_{k=1}^{j} \Delta n_k \qquad \text{(Equation S67)}$$

where $n_{\text{cum},j}$ is the cumulative signed amount of titrant added through step $j$ within a given titration branch (mmol), and $\Delta n_k$ is the signed amount of titrant added at step $k$ (mmol). The cumulative signed titrant equivalents were normalized to the original leachate volume as:

$$E_{\text{sample},j} = \frac{n_{\text{cum},j}}{V_{\text{sample}}} \qquad \text{(Equation S68)}$$

where $E_{\text{sample},j}$ is the cumulative titrant demand normalized to the original leachate volume (mmol $L^{-1}$, equivalent to mM), and $V_{\text{sample},L}$ is the original leachate volume used in the diluted titration mixture (L). The common interpolation grid was defined as:

$$\text{pH}^*_g = \text{pH}_{\text{min}} + g \cdot \Delta\text{pH} \qquad \text{(Equation S69)}$$

where $\text{pH}^*_g$ is the pH value at grid index $g$, $\text{pH}_{\text{min}} = 3.0$, $g$ is a unitless integer grid index, and $\Delta\text{pH} = 0.1$ pH units. Grid values were retained only when $\text{pH}^*_g \leq 9.0$ and when $\text{pH}^*_g$ fell within the measured pH range of the corresponding replicate and titration branch. For grid point $\text{pH}^*_g$ bracketed by measured points $(\text{pH}_a, E_a)$ and $(\text{pH}_b, E_b)$, the interpolated cumulative titrant demand was calculated by linear interpolation as:

$$E^*_g = E_a + \frac{E_b - E_a}{\text{pH}_b - \text{pH}_a} \cdot \left(\text{pH}^*_g - \text{pH}_a\right) \qquad \text{(Equation S70)}$$

where $E_g^*$ is the interpolated original-leachate-normalized cumulative titrant demand at $\mathrm{pH}_g^*$ (mM), $\mathrm{pH}_a$ and $\mathrm{pH}_b$ are the measured pH values bracketing $\mathrm{pH}_g^*$, and $E_a$ and $E_b$ are the corresponding original-leachate-normalized cumulative titrant demands (mM). No extrapolation was performed outside the measured pH range. The apparent buffering derivative at pH-grid point $g$ ($\beta_g$; mM $\mathrm{pH}^{-1}$) was calculated by finite differences as:

$$\beta_g = \begin{cases} \dfrac{E_1^* - E_0^*}{\mathrm{pH}_1^* - \mathrm{pH}_0^*}, & g = 0 \\ \dfrac{E_{g+1}^* - E_{g-1}^*}{\mathrm{pH}_{g+1}^* - \mathrm{pH}_{g-1}^*}, & 0 < g < n \\ \dfrac{E_n^* - E_{n-1}^*}{\mathrm{pH}_n^* - \mathrm{pH}_{n-1}^*}, & g = n \end{cases} \qquad \text{(Equation S71)}$$

where $n$ is the final pH-grid index for the corresponding replicate and titration branch. The derivative of the apparent buffering profile was calculated using the same finite-difference structure applied to $\beta$:

$$\left(\frac{d\beta}{d\mathrm{pH}}\right)_g = \begin{cases} \dfrac{\beta_1 - \beta_0}{\mathrm{pH}_1^* - \mathrm{pH}_0^*}, & g = 0 \\ \dfrac{\beta_{g+1} - \beta_{g-1}}{\mathrm{pH}_{g+1}^* - \mathrm{pH}_{g-1}^*}, & 0 < g < n \\ \dfrac{\beta_n - \beta_{n-1}}{\mathrm{pH}_n^* - \mathrm{pH}_{n-1}^*}, & g = n \end{cases} \qquad \text{(Equation S72)}$$

where $(d\beta/d\mathrm{pH})_g$ is the derivative of the apparent buffering profile at pH-grid point $g$ (mM $\mathrm{pH}^{-2}$). For plant-containing leachates, the biomass-associated buffering contribution at pH-grid point $g$ ($\Delta\beta_g$; mM $\mathrm{pH}^{-1}$) was calculated relative to the matched no-plant control as:

$$\Delta\beta_g = \beta_{g,\mathrm{plant}} - \beta_{g,\mathrm{no\ plant}} \qquad \text{(Equation S73)}$$

where $\beta_{g,\mathrm{plant}}$ and $\beta_{g,\mathrm{no\ plant}}$ are the apparent buffering derivatives of the plant-containing leachate and matched no-plant control, respectively.

For each replicate $r$, the integrated acid-side buffering contribution over the pH interval $[a, b]$ was defined from the corresponding matched-control difference in Equation S73 as:

$$B_{r,\mathrm{acid}}[a, b] = -\int_a^b \Delta\beta_{r,\mathrm{acid}}(p)\, dp \qquad \text{(Equation S74)}$$

The integrated base-side contribution was defined analogously as:

$$B_{r,\mathrm{base}}[a, b] = \int_a^b \Delta\beta_{r,\mathrm{base}}(p)\, dp \qquad \text{(Equation S75)}$$

Here, $p$ denotes pH, and $\Delta\beta_{r,\mathrm{acid}}$ and $\Delta\beta_{r,\mathrm{base}}$ denote the plant-minus-control buffering profiles for the acid- and base-addition branches, respectively (mM $\mathrm{pH}^{-1}$ on the original-leachate-volume basis). The integrated contributions $B_{r,\mathrm{acid}}$ and $B_{r,\mathrm{base}}$ therefore have units of mM acid or base equivalents. The total interval in Figure S46 is pH 3.0–9.0; the subintervals are pH 3.0–4.0, 4.0–5.5, 5.5–8.0, and 8.0–9.0. The contribution of subinterval $j$, with bounds $[a_j, b_j]$, to the total for branch $s$ was expressed as:

$$f_{r,j,s}(\%) = 100 \cdot \frac{B_{r,s}[a_j, b_j]}{B_{r,s}[3.0, 9.0]} \qquad \text{(Equation S76)}$$

where $s$ denotes the acid or base branch, and the denominator must be nonzero. These signed contributions follow the plotted definitions $-\Delta\beta$ and $\Delta\beta$; they do not replace a profile by its absolute value. Comparable totals require the same integration bounds and a consistent treatment of available pH coverage.

Centrifugal ultrafiltration recoveries were calculated by direct weighing of the starting leachate, recovered concentrate, and filtrate fractions as specified by the manufacturer. For each fraction $j$, the recovered liquid volume ($V_j$; L) was calculated from the measured liquid mass as:

$$V_j = \frac{W_j}{\rho_j} \qquad \text{(Equation S77)}$$

where $W_j$ is the measured mass of liquid in fraction $j$ (g), and $\rho_j$ is the solution density (g $L^{-1}$). Because the separated fractions were dilute aqueous leachates, $\rho_j$ was approximated as 1000 g $L^{-1}$ unless otherwise indicated. The mass of elemental analyte $i$ in each fraction $j$ ($m_{i,j}$; mg) was calculated as:

$$m_{i,j} = C_{i,j} \cdot V_j \qquad \text{(Equation S78)}$$

where $C_{i,j}$ is the blank- and dilution-corrected concentration of analyte $i$ in fraction $j$ measured by elemental analysis (mg $L^{-1}$), and $V_j$ is the recovered liquid volume of fraction $j$ (L). The total analytical recovery of analyte $i$ ($R_{i,\text{total}}$; %) was calculated as:

$$R_{i,\text{total}} = R_{i,c} + R_{i,f} \qquad \text{(Equation S79)}$$

Elemental partitioning of analyte $i$ between the recovered concentrate and filtrate ($P_{i,c/f}$; %) was calculated from the recovered mass in each separated fraction as:

$$P_{i,c/f} = 100 \cdot \frac{m_{i,c/f}}{m_{i,c} + m_{i,f}} \qquad \text{(Equation S80)}$$

where $P_{i,c}$ and $P_{i,f}$ represent the percentages of recovered analyte $i$ present in the concentrate and filtrate, respectively. Total recovery was used to evaluate mass-balance closure relative to the starting leachate, whereas concentrate-filtrate partitioning was used to evaluate whether recovered analyte was preferentially associated with the retained or permeate fraction. For UV absorbance measurements, the scatter- and background-corrected absorbance at wavelength $\lambda$ in fraction $j$ ($A_{\text{corr},j}(\lambda)$; absorbance units) was calculated as:

$$A_{\text{corr},j}(\lambda) = [A_{\text{meas},j}(\lambda) - A_{\text{meas},j}(750)] - [A_{\text{blank},j}(\lambda) - A_{\text{blank},j}(750)] \qquad \text{(Equation S81)}$$

where $A_{\text{meas},j}(\lambda)$ is the measured absorbance of fraction $j$ at wavelength $\lambda$, $A_{\text{meas},j}(750)$ is the corresponding absorbance at 750 nm used for scatter correction, $A_{\text{blank},j}(\lambda)$ is the absorbance of the matched no-biomass or matrix blank at wavelength $\lambda$, and $A_{\text{blank},j}(750)$ is the corresponding blank absorbance at 750 nm. The integrated UV absorbance from 220 to 400 nm for fraction $j$ ($U_j$; absorbance·nm) was calculated by trapezoidal integration as:

$$U_j = \sum_{\ell=1}^{N_\lambda - 1} \frac{A_{\text{corr},j}(\lambda_\ell) + A_{\text{corr},j}(\lambda_{\ell+1})}{2} \cdot (\lambda_{\ell+1} - \lambda_\ell) \qquad \text{(Equation S82)}$$

where $\lambda_\ell$ are the measured wavelengths between 220 and 400 nm sorted in increasing order, and $N_\lambda$ is the number of wavelengths in the integration range. UV-metric recovery in the concentrate or filtrate relative to the starting leachate ($R_{U,c/f}$; %) was calculated as:

$$R_{U,c/f} = 100 \cdot \frac{U_{c/f} \cdot V_{c/f}}{U_0 \cdot V_0} \qquad \text{(Equation S83)}$$

where $U_0$, $U_c$, and $U_f$ are the integrated UV absorbance metrics for the starting leachate, concentrate, and filtrate, respectively, and $V_0$, $V_c$, and $V_f$ are the corresponding volumes. UV-metric partitioning between the recovered concentrate and filtrate ($P_{U,c/f}$; %) was calculated as:

$$P_{U,c/f} = 100 \cdot \frac{U_{c/f} \cdot V_{c/f}}{U_c \cdot V_c + U_f \cdot V_f} \qquad \text{(Equation S84)}$$

where $P_{U,c}$ and $P_{U,f}$ represent the percentages of recovered UV-active material present in the concentrate and filtrate, respectively.

For time-dependent comparisons, $D(t)_{\mathrm{REE}}$ denotes the corresponding operational distribution ratio using the calculation in Equation S32, with the inferred nonsolubilized REE loading as the solid-associated numerator and the corresponding dissolved REE concentration as the aqueous denominator. Surface-area normalization follows Equation S33, using the reference plant specific surface area instead of the sulfur specific surface area. For a cumulative feed-basis calculation, the nonsolubilized inventory is the initial inventory less the cumulative dissolved inventory from Equation S28; the corresponding cumulative dissolved inventory is divided by the chosen reference volume to obtain the aqueous concentration used in the operational ratio. The logarithmic ratio of the nonsolubilized to solubilized REE inventory ($Z$; unitless)–which removes the explicit dry-mass, volume, and surface-area normalization factors used to express the distribution ratio–is defined as:

$$Z(t) = \log_{10}\left[\frac{n_{\mathrm{REE},0} - n_{\mathrm{REE,diss}}(t)}{n_{\mathrm{REE,diss}}(t)}\right] = \log_{10}\left[\frac{100 - S_{\mathrm{REE}}(t)}{S_{\mathrm{REE}}(t)}\right] \quad \text{(Equation S85)}$$

where $n_{\mathrm{REE},0}$ is the initial summed molar REE inventory, $n_{\mathrm{REE,diss}}(t)$ is its corresponding cumulative dissolved inventory, and $S_{\mathrm{REE}}(t)$ is the cumulative solubilization in percent calculated using Equation S29 from summed REE inventories, which is constrained by $0 < S_{\mathrm{REE}}(t) < 100$.

For each analyte $i$, overall bioleaching mass-balance closure ($B_{i,\mathrm{bio}}$; %) is defined relative to the input inventory as:

$$B_{i,\mathrm{bio}} = 100 \cdot \frac{m_{i,\mathrm{aq}} + m_{i,\mathrm{coarse\ residue}} + m_{i,\mathrm{cells}} + m_{i,\mathrm{other\ recovered}}}{m_{i,\mathrm{in}}}. \quad \text{(Equation S86)}$$

All inventory terms are in mg and represent nonoverlapping fractions. The input inventory includes the initial plant-associated analyte, obtained from Equation S24 by conversion from mmol to mg, and all separately quantified analyte introduced in medium, inoculum, or subsequent additions. The aqueous inventory, $m_{i,\mathrm{aq}}$, includes final recovered solution and previously withdrawn aqueous samples using the inventory-summation approach in Equation S28. The coarse-residue inventory represents recovered plant/sulfur-containing solids, and the cell inventory represents the separately recovered bacterial fraction. Solid inventories are calculated from their complete representative digest concentrations and the corresponding recovered dry mass using Equation S2. The other-recovered term includes separately collected washes or other measured fractions not already counted in the preceding terms. For comparison with a balance excluding the cell-associated inventory, the same expression is evaluated without $m_{i,\mathrm{cells}}$ in the numerator. This whole-incubation calculation is not the same as the stage-specific separation recovery calculations described in Equations S87–S94.

For the coproduct-separation tests, operational recovery of the REE-depleted plant-solid residue ($R_{\mathrm{plant,res}}$; %) was calculated from the lyophilized mass of the recovered stage-1 residue as:

$$R_{\mathrm{plant,res}} = 100 \cdot \frac{m_{\mathrm{res},1}}{m_{\mathrm{plant},0}} \quad \text{(Equation S87)}$$

where $m_{\mathrm{res},1}$ is the lyophilized mass of the recovered stage-1 residue, and $m_{\mathrm{plant},0}$ is the initial dry mass of plant tissue added to the culture. This value was treated as an operational plant-solid residue recovery because the recovered residue could also include insoluble sulfur or cell-associated solids depending on the test condition. For dissolved REEs, the mass of analyte $i$ in aqueous fraction $x$ during centrifugation stage $s$ ($m_{i,s,x}$; mg) was calculated as:

$$m_{i,s,x} = C_{i,s,x} \cdot V_{s,x} \quad \text{(Equation S88)}$$

where $C_{i,s,x}$ is the blank- and dilution-corrected concentration of analyte $i$ in fraction $x$ at stage $s$ (mg $\mathrm{L}^{-1}$), and $V_{s,x}$ is the corresponding recovered volume (L). When total REE recovery was reported, the total recovered REE mass was calculated as:

$$m_{\mathrm{REE},s,x} = \sum_i C_{i,s,x} \cdot V_{s,x} \quad \text{(Equation S89)}$$

where $i$ represents the individual REEs quantified in that experiment. For *A. ferrooxidans*, the SYBR Green signal inventory in fraction $x$ during centrifugation stage $s$ ($S_{\mathrm{AF},s,x}$; RFU·L) was calculated as:

$$S_{\mathrm{AF},s,x} = F_{s,x} \cdot V_{s,x} \quad \text{(Equation S90)}$$

where $F_{s,x}$ is the background-corrected SYBR Green fluorescence signal of fraction $x$, and $V_{s,x}$ is the recovered fraction volume (L). For either an aqueous REE inventory or a SYBR Green bacterial signal inventory, the stage-specific supernatant recovery was calculated as:

$$R_{X,s,\text{sup}} = 100 \cdot \frac{I_{X,s,\text{sup}}}{I_{X,s,\text{in}}} \qquad \text{(Equation S91)}$$

where $X$ represents either dissolved REE mass or *A. ferrooxidans* SYBR Green signal, $I_{X,s,\text{in}}$ is the corresponding inventory entering centrifugation stage $s$, and $I_{X,s,\text{sup}}$ is the inventory recovered in the supernatant after that stage. The corresponding stage-specific residue retention was calculated as:

$$R_{X,s,\text{res}} = 100 - R_{X,s,\text{sup}} \qquad \text{(Equation S92)}$$

The overall recovery of dissolved REE analyte $i$ in the final aqueous leachate after the two-stage centrifugation workflow was calculated as:

$$R_{i,\text{aq,overall}} = \frac{R_{i,1,\text{sup}} \cdot R_{i,2,\text{sup}}}{100} \qquad \text{(Equation S93)}$$

where $R_{i,1,\text{sup}}$ is the stage-1 supernatant recovery of dissolved analyte $i$ and $R_{i,2,\text{sup}}$ is the stage-2 supernatant recovery of dissolved analyte $i$. The overall recovery of *A. ferrooxidans* in the final microbial pellet was calculated as:

$$R_{\text{AF,pellet,overall}} = \frac{R_{\text{AF},1,\text{sup}} \cdot R_{\text{AF},2,\text{res}}}{100} \qquad \text{(Equation S94)}$$

where $R_{\text{AF},1,\text{sup}}$ describes bacterial carryover through the plant-solid removal step and $R_{\text{AF},2,\text{res}}$ describes bacterial retention in the cell-pellet fraction during the second centrifugation step.

For the $S^0$ adsorption experiments, the concentration of aqueous analyte $i$ in the no-$S^0$ control ($C_{\text{ref},i}$; mg $L^{-1}$) was calculated as:

$$C_{\text{ref},i} = C_{\text{ref},i}^{\text{meas}} - \overline{C}_{\text{medium},i} \qquad \text{(Equation S95)}$$

where $C_{\text{ref},i}^{\text{meas}}$ is the analyte concentration in the control containing medium and added ions without $S^0$ (mg $L^{-1}$), and $\overline{C}_{\text{medium},i}$ is the mean concentration in the corresponding medium-only controls (mg $L^{-1}$). The concentration of aqueous analyte $i$ after incubation with $S^0$ ($C_{\text{aq},i}$; mg $L^{-1}$) was calculated as:

$$C_{\text{aq},i} = C_{\text{aq},i}^{\text{meas}} - \overline{C}_{\text{S},i} \qquad \text{(Equation S96)}$$

where $C_{\text{aq},i}^{\text{meas}}$ is the analyte concentration in the supernatant of the $S^0$-containing sample, and $\overline{C}_{\text{S},i}$ is the mean concentration in the corresponding controls containing medium and $S^0$ without added ions (mg $L^{-1}$). The $S^0$-attributed change in the soluble concentration of analyte $i$ ($\Delta C_{\text{app},i}$; mg $L^{-1}$) was calculated as:

$$\Delta C_{\text{app},i} = \overline{C}_{\text{ref},i} - C_{\text{aq},i} \qquad \text{(Equation S97)}$$

where $\overline{C}_{\text{ref},i}$ is the mean corrected concentration in the matched no-$S^0$ controls. The apparent recovery, or extent of apparent removal, of analyte $i$ by $S^0$ ($R_{\text{app},i}$; %) was calculated as:

$$R_{\text{app},i} = 100 \cdot \frac{\Delta C_i}{\overline{C}_{\text{ref},i}} \qquad \text{(Equation S98)}$$

The inferred mass loading of analyte $i$ on $S^0$ ($q_{\text{mass,app},i}$; mg $kg^{-1}$ $S^0$) was calculated as:

$$q_{\text{mass,app},i} = \frac{\Delta C_i}{m_{\text{S,dry}}} \qquad \text{(Equation S99)}$$

where $m_{\text{S,dry}}$ is the dry $S^0$ mass per unit final solution volume (kg $L^{-1}$). The apparent molar adsorption capacity of analyte $i$ on $S^0$ ($q_{\text{app},i}$; mmol $kg^{-1}$ $S^0$) was calculated as:

$$q_{\text{app},i} = \frac{q_{\text{mass},i}}{M_i} \qquad \text{(Equation S100)}$$

where $M_i$ is the elemental molar mass of analyte $i$ (mg mmol$^{-1}$). The apparent equilibrium solution concentration of analyte $i$ after incubation ($c_{\text{e,app},i}$; mmol L$^{-1}$) was calculated as:

$$c_{\text{e,app},i} = \frac{C_{\text{aq},i}}{M_i} \qquad \text{(Equation S101)}$$

For determination of nominal reactant to REE molar ratios for the literature comparisons, reported concentrations took precedence over pH-derived estimates. Where only initial or explicitly stated acid pH was available, the ideal concentration (mol L$^{-1}$) was estimated using:

$$h = 10^{-\text{pH}_0} \qquad \text{(Equation S102)}$$

For strong monoprotic acids, the nominal reactant concentration was assumed to be:

$$C^{\text{ideal}}_{\text{acid},0} = h. \qquad \text{(Equation S103)}$$

For sulfuric acid, complete first dissociation and a second-dissociation constant of $K_{a,2} = 0.0102$ mol L$^{-1}$[159] were assumed:

$$C^{\text{ideal}}_{\text{H}_2\text{SO}_4,0} = \frac{h(h + K_{a,2})}{h + 2K_{a,2}}. \qquad \text{(Equation S104)}$$

For the illustrative EDTA complexation calculations in Text S25, ionic activity coefficients were approximated using the Davies expression:[49]

$$\log_{10} \gamma_j(I) = -Az_j^2 \left( \frac{\sqrt{I/c^\circ}}{1 + \sqrt{I/c^\circ}} - 0.3\frac{I}{c^\circ} \right) \qquad \text{(Equation S105)}$$

where $\gamma_j$ is the activity coefficient of species $j$ (unitless), $z_j$ is its charge number, $I$ is the ionic strength (mol L$^{-1}$), $c^\circ = 1$ mol L$^{-1}$ is the standard concentration, and $A = 0.509$ was used for the approximate 25°C correction.[60] The reference ionic strength was $I_{\text{ref}} = 0.1$ mol L$^{-1}$, and the illustrative wash calculation used $I = 0.4$ mol L$^{-1}$. The formation constants describe the 1:1 complex MY formed from free metal M and fully deprotonated EDTA, denoted by $Y^{4-}$.[47,48] The concentration-based formation constant was adjusted between ionic strengths as:[49]

$$\log_{10} \beta_{\text{MY}}(I) = \log_{10} \beta_{\text{MY}}(I_{\text{ref}}) + \log_{10} \left[ \frac{\gamma_{\text{M}}(I)\gamma_{\text{Y}}(I)\gamma_{\text{MY}}(I_{\text{ref}})}{\gamma_{\text{MY}}(I)\gamma_{\text{M}}(I_{\text{ref}})\gamma_{\text{Y}}(I_{\text{ref}})} \right] \qquad \text{(Equation S106)}$$

where $\beta_{\text{MY}}$ is the concentration-based formation constant (M$^{-1}$), and the activity coefficients correspond to the charges of the free metal, $Y^{4-}$, and the complex. The complex charge is −1 for a trivalent lanthanide and −2 for $Mg^{2+}$. The four reference EDTA $pK_a$ values were the selected literature values for 0.1 M KCl.[50,51] For each acid-dissociation step of the $H_4Y$ approximation, the reference concentration-based $pK_a$ was adjusted as:[49]

$$pK_{a,r}(I) = pK_{a,r}(I_{\text{ref}}) + \log_{10} \left[ \frac{\gamma_{\text{H}^+}(I)\gamma_{\text{H}_{4-r}\text{Y}}(I)\gamma_{\text{H}_{5-r}\text{Y}}(I_{\text{ref}})}{\gamma_{\text{H}_{5-r}\text{Y}}(I)\gamma_{\text{H}^+}(I_{\text{ref}})\gamma_{\text{H}_{4-r}\text{Y}}(I_{\text{ref}})} \right], \qquad r = 1, \ldots, 4 \qquad \text{(Equation S107)}$$

where the protonated reactant and deprotonated product in step $r$ have charges $-(r-1)$ and $-r$, respectively. The corresponding concentration-based dissociation constants ($K_{a,r}$; mol L$^{-1}$) were calculated as:

$$K_{a,r} = c^\circ 10^{-pK_{a,r}} \qquad \text{(Equation S108)}$$

With pH defined by hydrogen-ion activity, the free hydrogen-ion concentration ($h$; mol L$^{-1}$) and the fraction of uncomplexed EDTA present as $Y^{4-}$ ($\alpha_{\text{Y}}$; unitless) were calculated as:

$$h = \frac{c^\circ 10^{-\text{pH}}}{\gamma_{\text{H}^+}}, \qquad \text{(Equation S109)}$$

$$\alpha_{\mathrm{Y}} = \left(1 + \frac{h}{K_{a,4}} + \frac{h^2}{K_{a,3}K_{a,4}} + \frac{h^3}{K_{a,2}K_{a,3}K_{a,4}} + \frac{h^4}{K_{a,1}K_{a,2}K_{a,3}K_{a,4}}\right)^{-1} \quad \text{(Equation S110)}$$

where all $K_{a,r}$ values are concentration-based constants at the assumed wash ionic strength. The conditional formation constant for metal M ($K'_{\mathrm{MY}}$; $\mathrm{M}^{-1}$) was calculated as:

$$K'_{\mathrm{MY}} = \beta_{\mathrm{MY}} \cdot \alpha_{\mathrm{Y}} \quad \text{(Equation S111)}$$

which assumes no additional dissolved metal complexes, surface-binding equilibria, or precipitated phases. For the simplified competition calculation, the concentration of uncomplexed EDTA summed across its protonation states ($L_f$; mol $\mathrm{L}^{-1}$) was obtained by solving the ligand balance:

$$L_T - L_{\mathrm{reserve}} = L_f + \sum_{\mathrm{M}} C_{\mathrm{M},T} \frac{K'_{\mathrm{MY}} L_f}{1 + K'_{\mathrm{MY}} L_f} \quad \text{(Equation S112)}$$

where $L_T$ is the total supplied EDTA concentration, $L_{\mathrm{reserve}}$ is the concentration of ligand equivalents conservatively allocated to other multivalent cations, and $C_{\mathrm{M},T}$ is the total concentration of each explicitly modeled metal (all in mol $\mathrm{L}^{-1}$ of wash solution). The reference lanthanide and Mg formation constants were those tabulated by Hu et al.[48] and Smith et al.,[52] respectively. For the basal-medium competition scenario, the sum included Tb and Mg, whereas the no-competition scenario included only Tb and used $L_{\mathrm{reserve}} = 0$. The La and Lu sensitivity calculations substituted the selected lanthanide for Tb. The metal and ligand concentrations were obtained by rearranging Equation S23 for concentration, using the corresponding amount and assumed wash volume and expressing the result in mol $\mathrm{L}^{-1}$. The free-metal and metal–EDTA concentrations were then calculated as:

$$[\mathrm{M}] = \frac{C_{\mathrm{M},T}}{1 + K'_{\mathrm{MY}} L_f}, \quad \text{(Equation S113)}$$

$$[\mathrm{MY}] = \frac{C_{\mathrm{M},T} K'_{\mathrm{MY}} L_f}{1 + K'_{\mathrm{MY}} L_f} \quad \text{(Equation S114)}$$

where [M] denotes the free metal-ion concentration (M) rather than total dissolved metal. The calculations assumed fixed pH and ionic strength, equilibrium 1:1 complexation, and only free metal and its EDTA complex as dissolved forms of each modeled metal. For an illustrative first-order model of release from an accessible, exchangeable Tb pool, the fraction remaining after two wash stages was expressed as:

$$\frac{m_{\mathrm{exch,after}}}{m_{\mathrm{exch,before}}} = \exp\left[-k_{\mathrm{observed}}(t_1 + t_2)\right] \quad \text{(Equation S115)}$$

where $m_{\mathrm{exch,before}}$ and $m_{\mathrm{exch,after}}$ are the inventories in the modeled exchangeable bound pool before and after washing (mg), respectively, $k_{\mathrm{observed}}$ is the first-order release rate constant ($\mathrm{min}^{-1}$), and $t_1$ and $t_2$ are the contact durations of the two stages (min). The expression assumes negligible equilibrium retention, the same rate constant in both stages, and no contribution from dissolved carryover or other retained pools. For a target remaining fraction $r_{\mathrm{target}}$ (unitless), the minimum required rate constant ($k_{\mathrm{min}}$; $\mathrm{min}^{-1}$) was obtained by rearranging Equation S115:

$$k_{\mathrm{min}} = \frac{-\ln(r_{\mathrm{target}})}{t_1 + t_2} \quad \text{(Equation S116)}$$

where $0 < r_{\mathrm{target}} < 1$. The corresponding maximum half-time ($t_{1/2,\mathrm{max}}$; min) was calculated as:

$$t_{1/2,\mathrm{max}} = \frac{\ln 2}{k_{\mathrm{min}}} \quad \text{(Equation S117)}$$

For abiotic incubations of Tb with phosphate, the inferred decrease in soluble Tb inventory ($m_{\mathrm{Tb,depleted}}$;

mg) was calculated as:

$$m_{\mathrm{Tb,depleted}} = m_{\mathrm{Tb},0} - C_{\mathrm{Tb,sup}} \cdot V_{\mathrm{liquid}} \quad \text{(Equation S118)}$$

where $m_{\mathrm{Tb},0}$ is the initial Tb inventory (mg), $C_{\mathrm{Tb,sup}}$ is the post-incubation soluble Tb concentration (mg $\mathrm{L}^{-1}$), and $V_{\mathrm{liquid}}$ is the total liquid volume present at the end of incubation before supernatant removal (L). The corresponding percentage of the initial inventory ($D_{\mathrm{Tb}}$; %) was calculated as:

$$D_{\mathrm{Tb}} = 100 \cdot \frac{m_{\mathrm{Tb,depleted}}}{m_{\mathrm{Tb},0}}. \quad \text{(Equation S119)}$$

The apparent recovery in the complete washing sequence relative to that depleted inventory ($R_{\mathrm{Tb,P,app}}$; %) was calculated by:

$$R_{\mathrm{Tb,P,app}} = 100 \cdot \frac{\sum_{k=1}^{N_W} m_{\mathrm{Tb,wash},k}}{m_{\mathrm{Tb,depleted}}} \quad \text{(Equation S120)}$$

The calculation of $R_{\mathrm{Tb,P,app}}$ is constrained by $m_{\mathrm{Tb,depleted}} > 0$. The corresponding overall incubation mass-balance closure ($B_{\mathrm{Tb,P}}$; %) was defined as:

$$B_{\mathrm{Tb,P}} = 100 \cdot \frac{m_{\mathrm{Tb,sup,removed}} + \sum_{k=1}^{N_W} m_{\mathrm{Tb,wash},k} + m_{\mathrm{Tb,residue}}}{m_{\mathrm{Tb},0}} \quad \text{(Equation S121)}$$

where $m_{\mathrm{Tb,sup,removed}}$ is the Tb inventory in the supernatant actually removed (mg) and $m_{\mathrm{Tb,residue}}$ is the inventory measured by digestion of the final washed residue (mg).

The percentage of recovered wash-plus-residue Tb present in the two EDTA washes ($P_{\mathrm{Tb,EDTA}}$; %) was calculated using a separate denominator:

$$P_{\mathrm{Tb,EDTA}} = 100 \cdot \frac{m_{\mathrm{Tb,EDTA},1} + m_{\mathrm{Tb,EDTA},2}}{\sum_{k=1}^{N_W} m_{\mathrm{Tb,wash},k} + m_{\mathrm{Tb,residue}}} \quad \text{(Equation S122)}$$

where $m_{\mathrm{Tb,EDTA},1}$ and $m_{\mathrm{Tb,EDTA},2}$ are the inventories in the first and second EDTA washes (mg), respectively, and the denominator includes both EDTA washes, both water rinses, and the final residue.

### Supplementary tables

**Table S1. Chemicals, materials, and equipment used in this study.**

| Item | Manufacturer | Purity/model | Lot no. (Source no.) | Use |
|---|---|---|---|---|
| **Chemicals and reagents** | | | | |
| Acidophile cultivation and media preparation | | | | |
| Ammonium sulfate ($(NH_4)_2SO_4$) | Sigma Aldrich | $\geq$ 99.0% | 0000414475 (0000399014) | Media, solid–liquid extraction |
| Potassium phosphate monobasic ($KH_2PO_4$) | Sigma Aldrich | $\geq$ 99.0% | 1002190558 (SLBP479V) | Media |
| Magnesium sulfate heptahydrate ($MgSO_4 \cdot 7\,H_2O$) | Sigma Aldrich | $\geq$ 99.0% | 102475240 (BCCG2453) | Media |
| Trace Mineral Supplement (MD-TMS) | ATCC | | 70071354 | Media |
| Citric acid ($C_6H_8O_7$) | Sigma Aldrich | $\geq$ 99.5% | 1003178896 (MKCL2222) | Media, solid–liquid extraction |
| Iron(II) sulfate heptahydrate ($FeSO_4 \cdot 7\,H_2O$) | Sigma Aldrich | $\geq$ 99% | 0000417390 (0000398320) | Media |
| Iron(III) sulfate pentahydrate ($Fe_2(SO_4)_3 \cdot 5\,H_2O$) | Honeywell Fluka | 21–23% Fe basis | K314A (10094524) | Media |
| Sulfur, brown ($S^0$) | Toronto Research Chemicals Inc. | $\geq$ 99% | 1-LWF-28-1 | Media, $S^0$ binding |
| Sulfur, yellow ($S^0$) | Sigma Aldrich | $\geq$ 99.0% | 1003576369 (MKCT0576) | $S^0$ binding |
| Sulfuric acid ($H_2SO_4$) | Sigma Aldrich | 95.0–98.0% | 4104104872 (MKCX5032) | Media, solid–liquid extraction |
| Plant elemental treatment | | | | |
| Nickel(II) chloride hexahydrate ($NiCl_2 \cdot 6\,H_2O$) | Ward's Science | 100% | (7791-20-0) | Plant elemental treatment |
| Cerium (III) chloride heptahydrate ($CeCl_3 \cdot 7\,H_2O$) | Oakwood Chemical | 99% | (18618-55-8) | Plant elemental treatment |
| Neodymium chloride ($NdCl_3$) | Oakwood Chemical | 99% | (10024-93-8) | Plant elemental treatment |
| Terbium(III) chloride hexahydrate ($TbCl_3 \cdot 6\,H_2O$) | Sigma Aldrich | 99.9% | (13798-24-8) | Plant elemental treatment |
| Dysprosium (III) chloride hexahydrate ($DyCl_3 \cdot 6\,H_2O$) | Oakwood Chemical | 99% | (15059-52-6) | Plant elemental treatment |
| Solid–liquid extraction and titration | | | | |
| Nitric acid ($HNO_3$) | Sigma Aldrich | 70% | 4104110432 (MKCX6119) | Solid–liquid extraction |
| Potassium sulfate ($K_2SO_4$) | Sigma Aldrich | $\geq$ 99.0% | 0000461956 (0000382469) | Solid–liquid extraction |
| Potassium nitrate ($KNO_3$) | Sigma Aldrich | $\geq$ 99.0% | MKCR3008 (1003458844) | Solid–liquid extraction |
| Disodium ethylenediaminetetraacetate ($Na_2EDTA$) | Sigma | 99.0–101.0% | SLBM2279V (1001987424) | Solid–liquid extraction, washing cells |
| Indium sulfate ($In_2(SO_4)_3$) | aablocks | 99% | 1554335 (AA003QX6) | Solid–liquid extraction |
| Sodium hydroxide (NaOH) | Sigma Aldrich | $\geq$ 98% | SLBT8498 (1002488407) | Titrations, media |
| Hydrochloric acid (HCl) | Fisher Scientific | 37.5% | 133511 | Titrations |
| Methanesulfonic acid (MSA) | Sigma Aldrich | $\geq$ 99.0% | 102948407 (BCCP1470) | Solid–liquid extraction |
| Cell binding, REE repartitioning, and molecular analysis | | | | |
| 4-Morpholineethanesulfonic acid (MES) | Bio Basic | "high purity" | SB626010 (MB0341) | Cell binding |
| Sodium chloride (NaCl) | Sigma Aldrich | $\geq$ 99% | SLCJ2677 (1003281350) | Cell binding |
| Terbium chloride ($TbCl_3$) | Sigma Aldrich | 99.9% | 1003410557 (MKCP4743) | Cell binding |
| Terbium sulfate octahydrate ($Tb_2(SO_4)_3 \cdot 8\,H_2O$) | Sigma Aldrich | 99.9% | 1003446134 (19416KI) | Media, cell binding |
| Cerium(III) sulfate ($Ce_2(SO_4)_3$) | Thermo Scientific | 99.9% | A0434313 (378660500) | Cell binding, plant binding, $S^0$ binding |
| Neodymium sulfate octahydrate ($Nd_2(SO_4)_3 \cdot 8\,H_2O$) | Thermo Scientific | 99.9% | T24J010 (040253.14) | Cell binding |
| Dysprosium sulfate octahydrate ($Dy_2(SO_4)_3 \cdot 8\,H_2O$) | ChemCruz | | SC-223955 | Cell binding, plant binding, $S^0$ binding |

*Continued on next page*

*Table S1 continued*

| Item | Manufacturer | Purity/model | Lot no. (Source no.) | Use |
|---|---|---|---|---|
| Tris(hydroxymethyl)aminomethane hydrochloride (Tris−HCl) | Sigma | ≥99.0% | SLBR1378V (1002286292) | dsDNA measurements |
| SYBR™Green I Nucleic Acid Gel Stain | Sigma Aldrich | 10,000X concentrate in DMSO | 105377 | dsDNA measurements |
| Propidium iodide | Sigma-Aldrich | ≥94.0% | 102970296 (WXBF7582V) | Bacteria pretreatment assay |
| Triton X-100 | Sigma-Aldrich | ∼10% in $H_2O$ | 102583054 (BCCJ9119) | Bacteria pretreatment assay |
| NBD-PEN | TargetMol | 98.27% | 235740 (T33602) | Bacteria pretreatment assay |
| $DiBAC_4(3)$, sodium salt | Dojindo Laboratories | – | HV675 (D545) | Bacteria pretreatment assay |
| **Materials and consumables** | | | | |
| 1.5 mL microcentrifuge tubes | Eppendorf | – | – | Water-washing experiments |
| 15 mL centrifuge tubes | CELLTREAT | – | – | Lyophilizing plant samples |
| 50 mL sterile centrifuge tubes | CELLTREAT | – | – | Water-washing experiments |
| 150 mL porcelain crucibles | Fisherbrand | – | – | Calcination experiments |
| Silicon wafer, 6-inch diameter | Ted Pella Inc. | P-type, ⟨100⟩ | – (16015; Fisher Scientific NC0815962) | SEM–EDS sample substrate |
| Au/Pd, 60/40 wt% sputtering target | Kurt J. Lesker Company | 99.95% purity | KJL81003 | Conductive coating for SEM–EDS |
| **Equipment and instruments** | | | | |
| I26 shaking incubator | New Brunswick | – | – | Chemical leaching experiments |
| Vortex-Genie | Scientific Industries | – | – | Water-washing experiments |
| Mini tube rotator | Fisherbrand | – | – | Water-washing experiments |
| Sorvall ST Plus centrifuge | Thermo Scientific | – | – | Centrifuging samples |
| pH 700 benchtop meter | Oakton | – | – | pH and Eh measurements |
| FreeZone 6 Plus freeze dryer | Labconco | – | – | Lyophilizer |
| Discovery Hi-Res TGA | TA Instruments | – | – | Thermogravimetric analysis |
| OTF-1200X tube furnace | MTI Corporation | – | – | Calcination experiments |
| X'Pert3 Powder diffractometer | Malvern Panalytical | – | – | XRD spectra collection |
| $\phi$32 × 2.0 mm sample holder | MTI Corporation | – | – | XRD spectra collection |
| iS50 spectrometer | Nicolet | – | – | FTIR spectra collection |
| Diamond scribe | Ted Pella Inc. | Straight tip | NC9983656 | SEM–EDS imaging |
| Wafer-cleaving pliers | Ted Pella Inc. | 6" | 7295 | SEM–EDS imaging |
| Probe sonicator | | 1/16-inch-diameter probe tip | | SEM–EDS imaging |
| Sputter coater | Cressington | 108 Manual | – | SEM–EDS imaging |
| Scanning electron microscope | ZEISS | SIGMA VP | – | SEM–EDS imaging |
| Energy-dispersive X-ray detector | Bruker | XFlash 6\|30 | – | SEM–EDS imaging |
| Gas adsorption analyzer | Micromeritics | 3Flex | – | BET analysis |
| Sample degassing system | Micromeritics | VacPrep 061 | – | BET analysis |

*Continued on next page*

*Table S1 continued*

| **Item** | **Manufacturer** | **Purity/model** | **Lot no. (Source no.)** | **Use** |
|---|---|---|---|---|
| Ultrasonic water bath | Branson Ultrasonics (Emerson) | 3800 | – | Sonication during combined detergent and sonication pretreatment |
| Inductively coupled plasma mass spectrometer | PerkinElmer | NexION 2000B | – | Elemental analysis of separated leachate fractions and cell-associated metals |
| Inductively coupled plasma optical emission spectrometer | HORIBA | ULTIMA Expert | – | Elemental analysis of leachates and digested solids |
| UV–Vis spectrophotometer | Thermo Fisher Scientific | GENESYS 10S | – | Optical-density measurements and cell-density normalization |
| Fluorescence plate reader | Molecular Devices | SpectraMax M2 | – | SYBR Green fluorescence measurements |
| Aluminum digestion hot block | USA Scientific | 2512-0000 | – | Digestion |
| Stir plate | Corning | PC-420D | – | Titrations |
| Short-wave ultraviolet source | | 254 nm | – | UV-C pretreatment of bacteria |
| Plant grinder | | Four-piece cylindrical herb grinder | – | Plant tissue homogenization |

**Table S2. Bacterial strains used in this study.**

| **Strain** | **Description** | **Source** |
|---|---|---|
| *A. ferrooxidans* (ATCC 23270) | Type strain | ATCC |
| *E. coli* DH5$\alpha$ | pET-24a(+) | NEB |

**Table S3. Media formulations used in this study.** AFM1[2] is modified from ATCC Medium 2039 (Acidithiobacillus ferrooxidans Medium) (American Type Culture Collection [ATCC], Manassas, VA, USA). The composition of F2S used in this study[61] was not the same as that used in earlier studies.[1] Modifications made from the reported media compositions are indicated by *.

| **Component** | **AFM1**[2] (pH 1.8) | **AFM1 BS**[2] (pH 1.8) | **AFM3 BS**[5] (pH 1.8) | **F2S**[61] (pH 1.8) | modified **SM4**[1] (pH 2.0* or 5.0) |
|---|---|---|---|---|---|
| $(NH_4)_2SO_4$ | 0.8 g/L | 0.8 g/L | 0.8 g/L | 0.8 g/L | 0.8 g/L |
| $KH_2PO_4$ | 0.1 g/L | 0.1 g/L | 0.1 g/L | 0.1 g/L | 0.1 g/L |
| $MgSO_4 \cdot 7\,H_2O$ | 2.0 g/L | 2.0 g/L | 2.0 g/L | 2.0 g/L | 2.0 g/L |
| MD-TMS | 5 mL/L | 5 mL/L | 5 mL/L | 5 mL/L | 5 mL/L |
| Citric acid | – | – | 1.92 g/L | 1.92 g/L | 0 g/L |
| $FeSO_4 \cdot 7\,H_2O$ | 20 g/L | – | – | 27.8 g/L | – |
| Sulfur, brown | – | – | – | 1 g/L | 1, 2, or 5 g/L* |

**Table S4. Replicate-level elemental concentrations and molar ratios in lyophilized *Phytolacca* tissues.** Elemental concentrations are reported in mg kg$^{-1}$ dry biomass. LREE, HREE, and REE were calculated from measured elemental concentrations. REE/P and HREE/LREE were calculated from the corresponding molar ratios. Unmeasured values are indicated by –.

| Species | Tissue | Growth treatment | *n* | Ca | K | Mg | P | Fe | Ni | Ce | Dy | Nd | Tb | LREE | HREE | REE | REE/ P | HREE/ LREE |
|---|---|---|---|---|---|---|---|---|---|---|---|---|---|---|---|---|---|---|
| *P. acinosa* | root | hydroponic baseline (control) | 1 | 21,000 | 44,400 | 2,480 | 7,410 | 629 | 16 | 115 | 78.5 | 0 | 0 | 115 | 79 | 194 | 0.01 | 0.59 |
| | | | 2 | 27,000 | 36,300 | 3,390 | 6,550 | 102 | 31.1 | 0 | 4.80 | 0 | 0 | – | 5 | 5 | 0.00 | – |
| | | | 3 | 32,300 | 37,600 | 3,540 | 7,020 | 1,150 | 35 | 509 | 136 | 0 | 0 | 509 | 136 | 645 | 0.02 | 0.23 |
| *P. acinosa* | root | +10 mM each Ce/Dy/Nd/Tb | 1 | 32,400 | 35,000 | 2,730 | 6,040 | 72 | 18 | 5,850 | 5,860 | 4,930 | 5,280 | 10,780 | 11,140 | 21,920 | 0.74 | 0.91 |
| | | | 2 | 31,100 | 36,600 | 2,960 | 6,870 | 81 | 24 | 6,280 | 6,340 | 5,250 | 5,410 | 11,530 | 11,750 | 23,280 | 0.70 | 0.90 |
| | | | 3 | 30,500 | 38,400 | 3,970 | 6,330 | 122 | 29 | 6,780 | 6,780 | 5,610 | 5,940 | 12,390 | 12,720 | 25,110 | 0.81 | 0.91 |
| | | | 4 | – | – | – | – | – | – | 3,360 | 3,380 | 3,460 | 3,700 | 6,820 | 7,080 | 13,900 | – | 0.92 |
| | | | 5 | – | – | – | – | – | – | 4,850 | 4,820 | 4,520 | 5,090 | 9,370 | 9,910 | 19,280 | – | 0.94 |
| | | | 6 | – | – | – | – | – | – | 2,460 | 2,270 | 2,330 | 2,460 | 4,790 | 4,730 | 9,520 | – | 0.87 |
| | | | 7 | – | – | – | – | – | – | 3,290 | 3,220 | 3,090 | 3,450 | 6,380 | 6,670 | 13,050 | – | 0.92 |
| | | | 8 | – | – | – | – | – | – | 3,540 | 3,700 | 4,730 | 3,970 | 8,270 | 7,670 | 15,940 | – | 0.82 |
| *P. acinosa* | root | +25 mM Ni | 1 | 38,900 | 5,440 | 2,460 | 614 | 174 | 12,000 | 26 | 4 | 1 | 0 | 27 | 4 | 31 | 0.01 | 0.12 |
| | | | 2 | 34,000 | 5,200 | 1,550 | 649 | 162 | 12,000 | 15 | 6 | 1 | 0 | 16 | 6 | 21 | 0.01 | 0.32 |
| | | | 3 | 28,500 | 6,080 | 2,390 | 628 | 151 | 9,860 | 15 | 6 | 1 | 0 | 16 | 6 | 22 | 0.01 | 0.30 |
| *P. acinosa* | root | +50 mM Ni | 1 | 33,800 | 16,100 | 1,700 | 491 | 157 | 9,870 | 5 | 3 | 1 | 0 | 6 | 3 | 8 | 0.00 | 0.40 |
| | | | 2 | 28,200 | 19,600 | 2,520 | 544 | 206 | 11,900 | 15 | 8 | 1 | 0 | 15 | 8 | 23 | 0.01 | 0.43 |
| | | | 3 | 33,700 | 18,000 | 2,390 | 483 | 192 | 12,200 | 24 | 2 | 1 | 0 | 25 | 2 | 27 | 0.01 | 0.06 |
| *P. acinosa* | shoot | hydroponic baseline (control) | 1 | 11,100 | 25,800 | 9,950 | 4,120 | 75 | 13 | 39 | 51 | 0 | 0 | 39 | 51 | 90 | 0.00 | 1.13 |
| | | | 2 | 12,200 | 28,000 | 9,830 | 4,880 | 89 | 12 | 56 | 51 | 0 | 0 | 56 | 51 | 107 | 0.00 | 0.80 |
| | | | 3 | 11,500 | 28,000 | 10,500 | 4,230 | 47 | 57 | 83 | 74 | 0 | 0 | 83 | 74 | 157 | 0.01 | 0.77 |
| *P. acinosa* | shoot | +10 mM each Ce/Dy/Nd/Tb | 1 | 11,000 | 44,400 | 12,200 | 4,840 | 0 | 119 | 8,840 | 9,800 | 5,570 | 7,380 | 14,410 | 17,180 | 31,590 | 1.33 | 1.05 |
| | | | 2 | – | – | – | – | – | – | 4,190 | 5,140 | 4,260 | 4,980 | 8,450 | 10,120 | 18,570 | – | 1.06 |
| | | | 3 | – | – | – | – | – | – | 6,190 | 7,540 | 4,810 | 6,280 | 11,000 | 13,820 | 24,820 | – | 1.11 |
| | | | 4 | – | – | – | – | – | – | 8,050 | 9,430 | 6,640 | 8,080 | 14,690 | 17,510 | 32,200 | – | 1.05 |
| | | | 5 | – | – | – | – | – | – | 6,150 | 6,890 | 5,100 | 6,670 | 11,250 | 13,560 | 24,810 | – | 1.06 |
| | | | 6 | – | – | – | – | – | – | 3,070 | 3,780 | 3,010 | 2,960 | 6,080 | 6,740 | 12,820 | – | 0.98 |
| | | | 7 | – | – | – | – | – | – | 7,560 | 8,860 | 8,250 | 8,440 | 15,810 | 17,300 | 33,110 | – | 0.97 |
| *P. acinosa* | shoot | +25 mM Ni | 1 | 10,500 | 33,700 | 9,830 | 6,980 | 44 | 31,200 | 11 | 2 | 1 | 0 | 12 | 2 | 14 | 0.00 | 0.18 |
| | | | 2 | 10,100 | 32,000 | 10,300 | 7,210 | 47 | 25,400 | 0 | 3 | 1 | 0 | 1 | 3 | 3 | 0.00 | 4.46 |
| | | | 3 | 10,100 | 32,100 | 9,980 | 7,240 | 56 | 27,000 | 6 | 5 | 0 | 0 | 6 | 5 | 11 | 0.00 | 0.66 |
| | | | 4 | – | – | – | – | – | 16,800 | – | – | – | – | – | – | – | – | – |
| | | | 5 | – | – | – | – | – | 19,300 | – | – | – | – | – | – | – | – | – |
| | | | 6 | – | – | – | – | – | 23,800 | – | – | – | – | – | – | – | – | – |

*Continued on next page*

*Table S4 continued*

| Species | Tissue | Growth treatment | *n* | Ca | K | Mg | P | Fe | Ni | Ce | Dy | Nd | Tb | LREE | HREE | REE | REE/ P | HREE/ LREE |
|---|---|---|---|---|---|---|---|---|---|---|---|---|---|---|---|---|---|---|
| *P. acinosa* | shoot | +50 mM Ni | 1 | 10,100 | 29,400 | 9,750 | 6,770 | 61 | 31,300 | 2 | 2 | 0 | 0 | 2 | 2 | 4 | 0.00 | 0.71 |
| | | | 2 | 9,200 | 30,400 | 9,530 | 6,110 | 54 | 30,300 | 0 | 3 | 1 | 0 | 1 | 3 | 3 | 0.00 | 4.24 |
| | | | 3 | 10,900 | 32,700 | 10,700 | 6,700 | 51 | 45,600 | 0 | 3 | 1 | 0 | 1 | 3 | 4 | 0.00 | 3.46 |
| | | | 4 | – | – | – | – | – | 42,500 | – | – | – | – | – | – | – | – | – |
| | | | 5 | – | – | – | – | – | 32,400 | – | – | – | – | – | – | – | – | – |
| | | | 6 | – | – | – | – | – | 42,600 | – | – | – | – | – | – | – | – | – |
| *P. americana* | shoot | hydroponic baseline (control) | 1 | 8,980 | 54,000 | 8,160 | 6,180 | 26 | 4 | 0 | 3 | 0 | 0 | 0 | 3 | 3 | 0.00 | 15.20 |
| | | | 2 | 8,780 | 65,800 | 12,000 | 6,350 | 41 | 4 | 0 | 2 | 0 | 0 | 1 | 2 | 3 | 0.00 | 2.43 |
| | | | 3 | 9,660 | 53,000 | 8,480 | 5,700 | 96 | 4 | 0 | 2 | 0 | 0 | 0 | 2 | 2 | 0.00 | 8.09 |
| | | | 4 | 7,530 | 38,800 | 11,600 | 6,100 | 0 | 2 | 5 | 0 | 0 | 0 | 5 | – | 5 | 0.00 | – |
| *P. americana* | shoot | soil-grown baseline (control) | 1 | 11,900 | 49,500 | 14,900 | 2,620 | 73 | 3 | 3 | 2 | 0 | 0 | 3 | 2 | 5 | 0.00 | 0.63 |
| | | | 2 | 9,580 | 45,900 | 10,400 | 2,560 | 93 | 4 | 0 | 2 | 0 | 0 | 1 | 2 | 3 | 0.00 | 2.91 |
| | | | 3 | 12,900 | 49,800 | 16,500 | 2,520 | 91 | 3 | 0 | 0 | 0 | 0 | 0 | – | 0 | 0.00 | – |
| *P. americana* | shoot | +10 mM each Ce/Dy | 1 | 9,820 | 38,100 | 9,710 | 4,200 | 59 | 22 | 7,330 | 7,970 | 8 | 48 | 7,338 | 8,018 | 15,356 | 0.75 | 0.94 |
| | | | 2 | 12,700 | 32,800 | 12,700 | 2,700 | 46 | 22 | 9,200 | 9,870 | 8 | 46 | 9,208 | 9,916 | 19,124 | 1.45 | 0.93 |
| | | | 3 | 14,200 | 33,200 | 13,900 | 2,950 | 53 | 25 | 10,900 | 12,000 | 7 | 52 | 10,907 | 12,052 | 22,959 | 1.60 | 0.95 |
| | | | 4 | – | – | – | – | – | – | 4,600 | 5,420 | – | – | 4,600 | 5,420 | 10,020 | – | 1.02 |
| | | | 5 | – | – | – | – | – | – | 4,340 | 5,560 | – | – | 4,340 | 5,560 | 9,900 | – | 1.10 |
| | | | 6 | – | – | – | – | – | – | 4,330 | 5,380 | – | – | 4,330 | 5,380 | 9,710 | – | 1.07 |
| | | | 7 | – | – | – | – | – | – | 5,400 | 6,520 | – | – | 5,400 | 6,520 | 11,920 | – | 1.04 |
| | | | 8 | – | – | – | – | – | – | 5,410 | 6,240 | – | – | 5,410 | 6,240 | 11,650 | – | 0.99 |

**Table S5. Two-way ANOVA and Holm–Bonferroni-adjusted contrasts of $\log_{10}$-transformed REE concentrations in *P. acinosa* roots and shoots after Ce/Dy/Nd/Tb supplementation.** REE concentrations were $\log_{10}$ transformed prior to analysis. Omnibus effects were tested using two-way ANOVA with REE supplementation and tissue type as factors. Planned comparisons tested REE-supplemented versus non-supplemented tissues within each tissue type and root versus shoot concentrations within each supplementation condition. Estimates and confidence intervals are reported on the $\log_{10}$ scale. Fold changes were calculated as $10^{\Delta \log_{10}}$. Pairwise comparisons used contrasts from the two-way ANOVA, with Holm–Bonferroni adjustment across the four planned comparisons in this table. Pairwise $P$ values are multiplicity adjusted; the reported 95% confidence intervals are pointwise and unadjusted.

| Test/model | Result type | Effect or comparison | Estimate [95% CI] | Fold change | Test statistic | *P* value | *n* |
|---|---|---|---|---|---|---|---|
| Two-way ANOVA | Omnibus term | REE supplementation | – | – | $F_{1,17} = 142$ | $< 0.0001$ | 21 |
| Two-way ANOVA | Omnibus term | Tissue type | – | – | $F_{1,17} = 0.535$ | 0.4745 | 21 |
| Two-way ANOVA | Omnibus term | Supplementation × tissue | – | – | $F_{1,17} = 0.00522$ | 0.9433 | 21 |
| Holm-adjusted contrast | Planned comparison | +REE vs. −REE, root | 2.30 [1.72, 2.87] | 200× | $t_{17} = 8.46$ | $< 0.0004$ | 8 vs. 3 |
| Holm-adjusted contrast | Planned comparison | +REE vs. −REE, shoot | 2.32 [1.74, 2.91] | 210× | $t_{17} = 8.40$ | $< 0.0004$ | 7 vs. 3 |
| Holm-adjusted contrast | Planned comparison | root vs. shoot, +REE | −0.156 [−0.593, 0.282] | 0.70× | $t_{17} = 0.751$ | 0.9264 | 8 vs. 7 |
| Holm-adjusted contrast | Planned comparison | root vs. shoot, −REE | −0.128 [−0.818, 0.563] | 0.75× | $t_{17} = 0.390$ | 0.9264 | 3 vs. 3 |

**Table S6. Two-way ANOVA and Šídák-adjusted comparisons of $\log_{10}$-transformed HREE concentrations in *P. acinosa* roots and shoots after Ce/Dy/Nd/Tb supplementation.** HREE concentrations were $\log_{10}$ transformed prior to analysis. Omnibus effects were tested using two-way ANOVA with REE supplementation and tissue type as factors. Planned comparisons tested shoot versus root HREE concentrations within each supplementation condition. Estimates and confidence intervals are reported on the $\log_{10}$ scale. Fold changes were calculated as $10^{\Delta \log_{10}}$. Pairwise comparisons were performed using Šídák's multiple comparisons test.

| **Test/model** | **Result type** | **Effect or comparison** | **Estimate** [95% CI] | **Fold change** | **Test statistic** | ***P* value** | ***n*** |
|---|---|---|---|---|---|---|---|
| Two-way ANOVA | Omnibus term | REE supplementation | – | – | $F_{1,17} = 270$ | $< 0.0001$ | 21 |
| Two-way ANOVA | Omnibus term | Tissue type | – | – | $F_{1,17} = 1.67$ | 0.2130 | 21 |
| Two-way ANOVA | Omnibus term | Supplementation × tissue | – | – | $F_{1,17} = 2.41 \times 10^{-5}$ | 0.9961 | 21 |
| Šídák | Planned comparison | shoot vs. root, +REE | 0.186 [−0.190, 0.562] | 1.53× | $t_{17} = 1.21$ | 0.4250 | 7 vs. 8 |
| Šídák | Planned comparison | shoot vs. root, −REE | 0.185 [−0.409, 0.778] | 1.53× | $t_{17} = 0.763$ | 0.7039 | 3 vs. 3 |

**Table S7. Unpaired Welch's $t$ test of $\log_{10}$-transformed HREE/LREE molar ratios between Ce/Dy/Nd/Tb-supplemented *P. acinosa* shoots and roots.** HREE/LREE molar ratios were $\log_{10}$ transformed prior to analysis. A two-tailed unpaired Welch's $t$ test was used to compare REE-supplemented shoot and root tissues. Estimates and confidence intervals are reported on the $\log_{10}$ scale as shoot minus root. Fold change was calculated as $10^{\Delta \log_{10}}$.

| **Comparison** | **Estimate** [95% CI] | **Fold change** | **Test statistic** | ***P* value** | ***n*** |
|---|---|---|---|---|---|
| shoot vs. root, +REE | 0.0631 [0.0402, 0.0861] | 1.16× | $t_{11.7} = 6.01$ | $< 0.0001$ | 7 vs. 8 |

**Table S8. Exact one-sample Wilcoxon signed-rank tests of $\log_{10}$-transformed HREE/LREE molar ratios against zero in Ce/Dy/Nd/Tb-supplemented *P. acinosa* shoots and roots.** HREE/LREE molar ratios were $\log_{10}$ transformed prior to analysis, so a value of 0 corresponds to HREE/LREE = 1. Two-tailed exact Wilcoxon signed-rank tests were used to test whether each tissue type differed from the theoretical median of 0. Median HREE/LREE values were calculated as $10^{\text{median } \log_{10}(\text{HREE/LREE})}$.

| **Tissue** | **Theoretical median** | **Observed median** | **Median HREE/LREE** | **W** | ***P* value** | ***n*** |
|---|---|---|---|---|---|---|
| *P. acinosa* shoot | 0 | 0.0755 | 1.19 | 28.0 | 0.0156 | 7 |
| *P. acinosa* root | 0 | 0.0128 | 1.03 | 18.0 | 0.2344 | 8 |

**Table S9. Two-way ANOVA and Holm–Bonferroni-adjusted contrasts of $\log_{10}$-transformed Ni concentrations in *P. acinosa* roots and shoots after Ni supplementation.** Ni concentrations were $\log_{10}$ transformed prior to analysis. Omnibus effects were tested using two-way ANOVA with Ni supplementation and tissue type as factors. Planned comparisons tested shoot versus root Ni concentrations within each supplementation condition and Ni supplementation effects within each tissue type. Estimates and confidence intervals are reported on the $\log_{10}$ scale in the direction stated in the comparison column. Fold changes were calculated as $10^{\Delta \log_{10}}$. Pairwise comparisons used contrasts from the two-way ANOVA, with Holm–Bonferroni adjustment across the nine planned comparisons in this table. Pairwise $P$ values are multiplicity adjusted; the reported 95% confidence intervals are pointwise and unadjusted.

| **Test/model** | **Result type** | **Effect or comparison** | **Estimate** [95% CI] | **Fold change** | **Test statistic** | ***P* value** | ***n*** |
|---|---|---|---|---|---|---|---|
| Two-way ANOVA | Omnibus term | Ni supplementation | – | – | $F_{2,18} = 736$ | $< 0.0001$ | 24 |
| Two-way ANOVA | Omnibus term | Tissue type | – | – | $F_{1,18} = 13.1$ | 0.0020 | 24 |
| Two-way ANOVA | Omnibus term | Supplementation × tissue | – | – | $F_{2,18} = 6.58$ | 0.0072 | 24 |
| Holm-adjusted contrast | Planned comparison | shoot vs. root, −Ni | −0.0973 [−0.368, 0.173] | 0.80× | $t_{18} = 0.756$ | 0.9192 | 3 vs. 3 |
| Holm-adjusted contrast | Planned comparison | shoot vs. root, +25 mM Ni | 0.319 [0.0846, 0.553] | 2.08× | $t_{18} = 2.86$ | 0.0416 | 6 vs. 3 |
| Holm-adjusted contrast | Planned comparison | shoot vs. root, +50 mM Ni | 0.515 [0.281, 0.750] | 3.27× | $t_{18} = 4.62$ | 0.0010 | 6 vs. 3 |
| Holm-adjusted contrast | Planned comparison | +25 mM Ni vs. −Ni, shoot | 3.05 [2.82, 3.29] | 1120× | $t_{18} = 27.4$ | $< 0.0009$ | 6 vs. 3 |
| Holm-adjusted contrast | Planned comparison | +50 mM Ni vs. −Ni, shoot | 3.25 [3.02, 3.49] | 1780× | $t_{18} = 29.2$ | $< 0.0009$ | 6 vs. 3 |
| Holm-adjusted contrast | Planned comparison | +50 mM Ni vs. +25 mM Ni, shoot | 0.198 [0.00645, 0.389] | 1.58× | $t_{18} = 2.17$ | 0.1305 | 6 vs. 6 |
| Holm-adjusted contrast | Planned comparison | +25 mM Ni vs. −Ni, root | 2.64 [2.37, 2.91] | 437× | $t_{18} = 20.5$ | $< 0.0009$ | 3 vs. 3 |
| Holm-adjusted contrast | Planned comparison | +50 mM Ni vs. −Ni, root | 2.64 [2.37, 2.91] | 437× | $t_{18} = 20.5$ | $< 0.0009$ | 3 vs. 3 |
| Holm-adjusted contrast | Planned comparison | +50 mM Ni vs. +25 mM Ni, root | 0.00133 [−0.269, 0.272] | 1.00× | $t_{18} = 0.0103$ | 0.9919 | 3 vs. 3 |

**Table S10. Brown–Forsythe and Welch ANOVAs with Dunnett's T3 comparisons of REE concentrations, and a Wilcoxon signed-rank test of HREE/LREE ratios, in *P. americana* shoots.** REE concentrations and HREE/LREE molar ratios were $\log_{10}$ transformed prior to analysis. REE concentrations were analyzed using Brown–Forsythe and Welch ANOVA tests, with Dunnett's T3 multiple comparisons used to compare REE-supplemented shoots against hydroponic and soil-grown non-supplemented controls. The HREE/LREE ratio in REE-supplemented shoots was tested against $\log_{10}(\text{HREE/LREE}) = 0$, corresponding to HREE/LREE $= 1$, using a two-tailed exact Wilcoxon signed-rank test. Estimates and confidence intervals are reported on the $\log_{10}$ scale in the direction stated in the comparison column. Fold changes were calculated as $10^{\Delta \log_{10}}$ or $10^{\text{median}}$ for the one-sample HREE/LREE test.

| **Response** | **Test/model** | **Result type** | **Effect or comparison** | **Estimate** [95% CI] | **Fold change** | **Test statistic** | ***P* value** | ***n*** |
|---|---|---|---|---|---|---|---|---|
| REE | Brown–Forsythe ANOVA | Omnibus term | Treatment | – | – | $F^{*}_{2.00,4.12} = 912$ | $< 0.0001$ | 14 |
| REE | Welch's ANOVA | Omnibus term | Treatment | – | – | $W_{2.00,2.75} = 763$ | 0.0002 | 14 |
| REE | Dunnett's T3 | Planned comparison | +REE vs. −REE | 3.63 [3.34, 3.93] | 4270× | $t_{5.37} = 38.0$ | $< 0.0001$ | 8 vs. 4 |
| REE | Dunnett's T3 | Planned comparison | −REE vs. −REE soil | −0.0995 [−0.866, 0.667] | 0.80× | $t_{2.16} = 0.723$ | 0.7525 | 4 vs. 2 |
| HREE/LREE | Wilcoxon signed-rank test | One-sample comparison | +REE vs. unity | 0.0663 [0.0334, 0.107] | 1.16× | $W = 36.0$ | 0.0078 | 8 |

**Table S11. Replicate-level BET surface areas and BJH cumulative pore volumes of control and Ce/Dy-enriched *P. americana* shoots and brown and yellow $S^0$.** All measurements were performed on dry (bio)mass. Independent replicate-level measurements are tabulated ($n = 3 - 9$). BJH adsorption cumulative pore volumes cover pore widths of 1.7–300 nm.

| **Sample** | **n** | **BET SA** ($m^2/g$) | **BJH adsorption cumulative pore volume** ($cm^3/g$) |
|---|---|---|---|
| *P. americana* shoot hydroponic baseline (control) | 1 | 14.9482 | 0.012221 |
| | 2 | 15.3200 | 0.012575 |
| | 3 | 14.1388 | 0.011810 |
| *P. americana* shoot hydrponic + 10 mM each Ce/Dy | 1 | 24.2641 | 0.021764 |
| | 2 | 25.4545 | 0.022758 |
| | 3 | 23.9238 | 0.021300 |
| Brown $S^0$ | 1 | 0.9179 | 0.002345 |
| | 2 | 1.2108 | 0.002408 |
| | 3 | 1.6017 | 0.003539 |
| | 4 | 1.4666 | 0.002882 |
| | 5 | 1.1491 | 0.002356 |
| Yellow $S^0$ | 1 | 4.0033 | 0.003422 |
| | 2 | 1.5029 | 0.002698 |
| | 3 | 2.1598 | 0.007707 |
| | 4 | 2.4560 | 0.002986 |
| | 5 | 4.7140 | 0.003874 |
| | 6 | 4.1524 | 0.004858 |
| | 7 | 3.1507 | 0.003658 |
| | 8 | 5.6932 | 0.005146 |
| | 9 | 4.3530 | 0.005793 |

**Table S12. Thermal decomposition metrics extracted from TGA/DTG analysis of lyophilized *Phytolacca* tissues.** Metrics are reported for the indicated growth and metal-supplementation conditions. $T_{\mathrm{max},i}$ denotes the temperature of local DTG minima. $\Delta m$ values denote interval-specific mass losses calculated from normalized mass-remaining curves. Residual mass was taken as the mass remaining at 1000 °C. Values are reported as mean ± standard deviation of biological replicates.

| Species | Tissue | Growth condition | *n* | $T_{\mathrm{max},1}$ (°C) | $T_{\mathrm{max},2}$ (°C) | $\Delta m_{20-200}$ (%) | $\Delta m_{200-400}$ (%) | $\Delta m_{400-1000}$ (%) | $m_{\mathrm{residual},1000}$ (%) |
|---|---|---|---|---|---|---|---|---|---|
| *P. acinosa* | root | hydroponic baseline (control) | 1 | 261 | 409 | 7.7 | 59.5 | 24.3 | 6.4 |
| | | +10 mM each Ce/Dy/Nd/Tb | 1 | 266 | 446 | 7.4 | 59.0 | 25.0 | 6.4 |
| *P. acinosa* | shoot | hydroponic baseline (control) | 1 | 273 | 430 | 7.6 | 67.3 | 19.0 | 3.5 |
| | | +25 mM Ni | 1 | 256 | 443 | 9.7 | 59.4 | 22.6 | 6.6 |
| | | +50 mM Ni | 1 | 279 | 445 | 10.1 | 56.2 | 24.8 | 7.0 |
| | | +10 mM each Ce/Dy/Nd/Tb | 1 | 279 | 463 | 10.2 | 59.1 | 22.8 | 5.8 |
| *P. americana* | shoot | hydroponic baseline (control) | 3 | 267 ± 5 | 399 ± 4 | 7.8 ± 0.7 | 58.1 ± 2.7 | 26.2 ± 3.1 | 7.1 ± 0.2 |
| | | soil-grown baseline (control) | 1 | 269 | 399 | 6.2 | 57.2 | 27.2 | 8.4 |
| | | +10 mM each Ce/Dy | 3 | 264 ± 15 | 452 ± 4 | 8.7 ± 0.5 | 46.2 ± 1.4 | 38.6 ± 1.7 | 5.2 ± 0.6 |

**Table S13. Ordinary two-way ANOVA and Šídák-adjusted comparisons of DTG peak temperatures in Ce/Dy-supplemented and control *P. americana* shoots.** The lower- and higher-temperature DTG peak temperatures (°C), $T_{\mathrm{max},1}$ and $T_{\mathrm{max},2}$, were analyzed using ordinary two-way ANOVA with peak identity, Ce/Dy supplementation, and their interaction as fixed effects. Planned comparisons compared Ce/Dy-supplemented shoots against the corresponding hydroponic *P. americana* controls separately for each peak. Estimates and confidence intervals were calculated as supplemented minus control. Pairwise 95% confidence intervals and $P$ values were Šídák adjusted across one family of two comparisons; omnibus $P$ values are unadjusted. Subscripts on test statistics indicate degrees of freedom, and pairwise $t$ statistics are reported as absolute values. For omnibus rows, $n$ reports the total number of analyzed peak-temperature values; for pairwise rows, $n$ reports the number of values in each group. Residual passed residual tests, and Spearman's test indicated heteroscedasticity ($P = 0.0226$).

| Test/model | Result type | Effect or comparison | Estimate [95% CI] | Test statistic | *P* value | *n* |
|---|---|---|---|---|---|---|
| Two-way ANOVA | Omnibus term | Peak identity | – | $F_{1,8} = 1031$ | < 0.0001 | 12 |
| Two-way ANOVA | Omnibus term | Ce/Dy supplementation | – | $F_{1,8} = 25.4$ | 0.0010 | 12 |
| Two-way ANOVA | Omnibus term | Peak identity × supplementation | – | $F_{1,8} = 31.8$ | 0.0005 | 12 |
| Šídák comparison | Planned comparison | +Ce/Dy vs. control, $T_{\mathrm{max},1}$ | −2.97 [−22.3, 16.3] | $t_8 = 0.422$ | 0.9003 | 3 vs. 3 |
| Šídák comparison | Planned comparison | +Ce/Dy vs. control, $T_{\mathrm{max},2}$ | 53.1 [33.8, 72.4] | $t_8 = 7.55$ | 0.0001 | 3 vs. 3 |

**Table S14. Elemental concentrations and enrichment factors in post-TGA *Phytolacca* residues.** Enrichment was calculated as the post-TGA concentration divided by the average corresponding dried tissue concentration, with errors determined by error propagation. Elemental concentrations are reported in mg $kg^{-1}$. Unmeasured values are indicated by –.

| Species | Tissue | Growth treatment | *n* | Ca | K | Mg | P | Fe | Ni | Ce | Dy | Nd | Tb | LREE | HREE | REE | REE/ P | HREE/ LREE |
|---|---|---|---|---|---|---|---|---|---|---|---|---|---|---|---|---|---|---|
| *P. acinosa* | root | hydroponic baseline (control) | 1 | 357,000 | 333,400 | 42,600 | 89,100 | 10,600 | 326 | 5 | 16 | 0 | 0 | – | – | – | 0.00 | 2.66 |
| | | | enrichment | 13.3 | 8.5 | 13.6 | 12.7 | 16.9 | – | – | – | – | – | – | – | – | 0.0 | – |
| *P. acinosa* | root | +10 mM each Ce/Dy/Nd/Tb | 1 | 222,000 | 245,000 | 32,800 | 61,800 | 1,270 | 622 | 54,800 | 55,300 | 46,300 | 46,300 | 101,100 | 101,600 | 202,700 | 0.67 | 0.89 |
| | | | enrichment | 7.1 | 6.7 | 10.2 | 9.6 | 13.8 | – | 12.0 | 12.2 | 10.9 | 10.5 | – | – | 11.4 | 0.9 | 0.87 |
| *P. acinosa* | shoot | hydroponic baseline (control) | 1 | 18,400 | 36,000 | 18,500 | 7,420 | 112 | 72 | 34 | 14 | 0 | 0 | 34 | 14 | 47 | 0.00 | 0.35 |
| | | | enrichment | 1.6 | 1.3 | 1.8 | 1.7 | 1.6 | – | – | – | – | – | – | – | – | 0.2 | – |
| *P. acinosa* | shoot | +10 mM each Ce/Dy/Nd/Tb | 1 | 136,000 | 157,000 | 131,000 | 59,400 | 595 | 405 | 128,000 | 148,000 | 95,000 | 142,000 | 223,000 | 290,000 | 513,000 | 1.76 | 1.15 |
| | | | enrichment | 12.4 | 3.5 | 10.7 | 12.3 | – | – | 20.3 | 20.1 | 17.7 | 22.2 | – | – | 25.9 | 1.3 | 0.98 |
| *P. acinosa* | shoot | +25 mM Ni | 1 | 128,000 | 167,000 | 123,000 | 86,800 | 445 | 331,000 | 0 | 13 | 4 | 0 | – | – | – | 0.00 | 2.63 |
| | | | enrichment | 12.5 | 5.3 | 12.3 | 12.3 | 8.6 | 13.3 | – | – | – | – | – | – | – | 0.2 | – |
| *P. acinosa* | shoot | +50 mM Ni | 1 | 95,000 | 84,700 | 97,600 | 72,400 | 1,180 | 487,000 | 0 | 7 | 23 | 0 | – | – | – | 0.00 | 0.26 |
| | | | enrichment | 9.4 | 2.7 | 9.8 | 11.1 | 21.4 | 13.0 | – | – | – | – | – | – | – | 0.8 | – |
| *P. americana* | shoot | hydroponic baseline (control) | 1 | 200,000 | 529,000 | 191,000 | 96,300 | 508 | 464 | 241 | 101 | 4 | 0 | 245 | 101 | 346 | 0.00 | 0.36 |
| | | | 2 | 136,000 | 387,000 | 120,000 | 73,800 | 378 | 207 | 95 | 34 | 24 | 0 | 119 | 34 | 153 | 0.00 | 0.25 |
| | | | 3 | 198,000 | 447,060 | 194,604 | 96,700 | 497 | 237 | 33 | 16 | 3 | 0 | 35 | 16 | 51 | 0.00 | 0.39 |
| | | | enrichment | 20.4 ± 3.8 | 8.6 ± 1.9 | 16.8 ± 4.5 | 14.6 ± 1.9 | 11.3 ± 9.8 | – | – | – | – | – | – | – | – | 4.1 ± 3 | – |
| *P. americana* | shoot | soil-grown baseline (control) | 1 | 181,000 | 344,000 | 183,000 | 31,800 | 806 | 158 | 52 | 17 | 2 | 0 | 53 | 17 | 70 | 0.00 | 0.27 |
| | | | enrichment | 15.8 | 7.1 | 13.1 | 12.4 | 9.4 | – | – | – | – | – | – | – | – | 2.2 | – |
| *P. americana* | shoot | +10 mM each Ce/Dy | 1 | 191,000 | 150,000 | 170,000 | 39,100 | 720 | 364 | 21,700 | 48,000 | 16 | 0 | 21,716 | 48,000 | 69,716 | 0.36 | 1.91 |
| | | | 2 | 206,000 | 134,000 | 203,000 | 42,100 | 775 | 875 | 33,800 | 66,000 | 5 | 0 | 33,805 | 66,000 | 99,805 | 0.48 | 1.68 |
| | | | 3 | 182,000 | 110,000 | 211,000 | 38,500 | 1,490 | 346 | 28,800 | 58,300 | 22 | 0 | 28,822 | 58,300 | 87,122 | 0.45 | 1.74 |
| | | | enrichment | 15.8 ± 2.5 | 3.8 ± 0.5 | 16.1 ± 2.8 | 12.2 ± 2.5 | 18.9 ± 6.9 | – | 4.4 ± 1.7 | 7.8 ± 2.6 | – | – | – | – | 6.2 ± 2.2 | 0.3 ± 0.1 | – |

**Table S15. Elemental concentrations and enrichment factors in *Phytolacca* residues after calcination at 600 °C.** Enrichment was calculated as the calcined-residue concentration divided by the average corresponding lyophilized tissue concentration, with errors determined by error propagation. Elemental concentrations are reported in mg $kg^{-1}$. Unmeasured values are indicated by –.

| **Species** | **Tissue** | **Growth treatment** | ***n*** | **Ca** | **K** | **Mg** | **P** | **Fe** | **Ni** | **Ce** | **Dy** | **Nd** | **Tb** | **LREE** | **HREE** | **REE** | **REE/ P** | **HREE/ LREE** |
|---|---|---|---|---|---|---|---|---|---|---|---|---|---|---|---|---|---|---|
| *P. acinosa* | shoot | hydroponic baseline (control) | 1 | 134,000 | 504,000 | 123,000 | 50,100 | 435 | 229 | 0 | 0 | 16 | 0 | 16 | – | 16 | $6.86 \times 10^{-5}$ | – |
| | | | enrichment | 11.6 | 18.5 | 12.2 | 11.4 | 6.2 | – | – | – | – | – | – | – | – | 0.012 | – |
| *P. acinosa* | shoot | +10 mM each Ce/Dy/Nd/Tb | 1 | 89,100 | 292,000 | 104,000 | 37,600 | 886 | 617 | 86,900 | 101,000 | 58,500 | 81,500 | 145,400 | 182,500 | 327,900 | 1.78 | 1.26 |
| | | | enrichment | 8.1 | 6.6 | 8.5 | 7.8 | – | 5.2 | 13.8 | 13.7 | 10.9 | 12.7 | – | – | – | 1.33 | 1.2 |
| *P. americana* | shoot | hydroponic baseline (control) | 1 | 162,000 | 504,000 | 123,000 | 50,100 | 435 | 229 | 0 | 0 | 2 | 0 | – | – | – | – | – |
| | | | 2 | 157,000 | 422,000 | 106,000 | 40,600 | 437 | 38 | 4 | 27 | 2 | 0 | – | – | – | – | – |
| | | | 3 | 118,000 | 430,000 | 108,000 | 41,700 | 429 | 63 | 6 | 3 | 4 | 0 | – | – | – | – | – |
| | | | enrichment | 16.7 ± 2.7 | 8.5 ± 1.7 | 11.2 ± 2.1 | 7.3 ± 0.8 | 10.6 ± 9.1 | – | – | – | – | – | – | – | – | – | – |
| *P. americana* | shoot | soil-grown baseline (control) | 1 | 145,000 | 368,000 | 123,000 | 20,500 | 694 | 17 | 23 | 34 | 4 | 0 | – | – | – | – | – |
| | | | 2 | 112,000 | 282,000 | 95,700 | 16,100 | 519 | 36 | 127 | 27 | 3 | 0 | – | – | – | – | – |
| | | | 3 | 149,000 | 373,000 | 122,000 | 21,400 | 722 | 61 | 3 | 36 | 2 | 0 | – | – | – | – | – |
| | | | enrichment | 11.8 ± 2.0 | 7.0 ± 0.9 | 8.2 ± 1.8 | 7.5 ± 0.9 | 7.5 ± 1.3 | – | – | – | – | – | – | – | – | – | – |
| *P. americana* | shoot | +10 mM each Ce/Dy | 1 | 144,000 | 317,000 | 146,000 | 25,000 | 613 | 172 | 57,900 | 89,800 | 1 | 0 | 57,901 | 89,800 | 147,701 | 1.20 | 1.55 |
| | | | 2 | 132,000 | 293,000 | 128,000 | 24,500 | 536 | 172 | 45,400 | 75,900 | 3 | 1 | 45,403 | 75,901 | 121,303 | 1.00 | 1.67 |
| | | | 3 | 142,000 | 295,000 | 133,000 | 24,400 | 569 | 158 | 53,800 | 86,000 | 4 | 0 | 53,804 | 86,000 | 139,804 | 1.16 | 1.60 |
| | | | enrichment | 11.4 ± 1.7 | 8.7 ± 0.7 | 11.2 ± 1.7 | 7.5 ± 1.5 | 10.9 ± 1.2 | 7.3 ± 0.6 | 8.1 ± 3.0 | 11.4 ± 3.6 | – | 0 ± 0 | – | – | – | 0.88 ± 0.33 | 1.6 ± 0.1 |

**Table S16. Two-way ANOVA and Tukey-adjusted comparisons of $\log_{10}$-transformed, baseline-corrected TREE/P molar ratios by thermochemical treatment and plant material.** Baseline-corrected TREE/P molar ratios were $\log_{10}$ transformed prior to analysis. Omnibus effects were tested using two-way ANOVA with thermochemical treatment and supplemented plant material as factors. Planned comparisons tested differences between plant materials within each treatment condition and differences between thermochemical treatments within each plant material. Estimates and confidence intervals are reported on the $\log_{10}$ scale in the direction stated in the comparison column. Pairwise comparisons were performed using Tukey's multiple comparisons test.

| Test/model | Result type | Effect or comparison | Estimate [95% CI] | Test statistic | *P* value | *n* |
|---|---|---|---|---|---|---|
| Two-way ANOVA | Omnibus term | Thermochemical treatment | – | $F_{2,6} = 2.84$ | 0.1355 | 12 |
| Two-way ANOVA | Omnibus term | Plant material | – | $F_{1,6} = 12.1$ | 0.0131 | 12 |
| Two-way ANOVA | Omnibus term | Treatment × plant material | – | $F_{2,6} = 5.04$ | 0.0519 | 12 |
| Tukey | Planned comparison | +Ce/Dy *P. americana* vs. +Ce/Dy/Nd/Tb *P. acinosa*, dried | −0.0224 [−0.339, 0.295] | $q_6 = 0.244$ | 0.8687 | 3 vs. 1 |
| Tukey | Planned comparison | +Ce/Dy *P. americana* vs. +Ce/Dy/Nd/Tb *P. acinosa*, post-TGA | −0.585 [−0.902, −0.268] | $q_6 = 6.38$ | 0.0040 | 3 vs. 1 |
| Tukey | Planned comparison | +Ce/Dy *P. americana* vs. +Ce/Dy/Nd/Tb *P. acinosa*, calcined | −0.174 [−0.491, 0.143] | $q_6 = 1.90$ | 0.2275 | 3 vs. 1 |
| Tukey | Planned comparison | dried vs. post-TGA, +Ce/Dy *P. americana* | 0.441 [0.160, 0.722] | $q_6 = 6.81$ | 0.0071 | 3 vs. 3 |
| Tukey | Planned comparison | dried vs. calcined, +Ce/Dy *P. americana* | 0.0267 [−0.254, 0.308] | $q_6 = 0.412$ | 0.9546 | 3 vs. 3 |
| Tukey | Planned comparison | post-TGA vs. calcined, +Ce/Dy *P. americana* | −0.414 [−0.695, −0.133] | $q_6 = 6.39$ | 0.0095 | 3 vs. 3 |
| Tukey | Planned comparison | dried vs. post-TGA, +Ce/Dy/Nd/Tb *P. acinosa* | −0.122 [−0.609, 0.365] | $q_6 = 1.08$ | 0.7357 | 1 vs. 1 |
| Tukey | Planned comparison | dried vs. calcined, +Ce/Dy/Nd/Tb *P. acinosa* | −0.125 [−0.612, 0.362] | $q_6 = 1.11$ | 0.7233 | 1 vs. 1 |
| Tukey | Planned comparison | post-TGA vs. calcined, +Ce/Dy/Nd/Tb *P. acinosa* | −0.00350 [−0.490, 0.483] | $q_6 = 0.0312$ | 0.9997 | 1 vs. 1 |



**Table S17. Paired *t* test and repeated-measures one-way ANOVA with Šídák-adjusted comparisons of elemental release during washing of REE-enriched *P. americana* shoot ash.** Wash-release percentages were analyzed for ash generated from REE-enriched *P. americana* shoots. REE and P release were first compared directly using a paired two-tailed *t* test on untransformed release percentages. Differences between REE or P release and major inorganic element release were then evaluated using repeated-measures one-way ANOVA on $\log_{10}$-transformed wash-release percentages, with Geisser–Greenhouse correction for nonsphericity and Šídák-corrected multiple comparisons. Estimates for the repeated-measures ANOVA comparisons are reported on the $\log_{10}$ scale in the direction shown.

| Test/model | Result type | Effect or comparison | Estimate [95% CI] | Test statistic | *P* value | *n* |
|---|---|---|---|---|---|---|
| Paired t-test | Planned comparison | REE release vs. P release | −2.73 [−7.40, 1.97] | – | 0.1301 | 3 |
| Repeated-measures ANOVA | Omnibus term | Element released | – | $F_{1.12,2.23} = 51.7$ | 0.0138 | 3 |
| Repeated-measures ANOVA | Matching term | Individual ash replicate | – | $F_{2,8} = 1.25$ | 0.3360 | 3 |
| Šídák | Planned comparison | REE vs. Ca | −0.959 [−2.61, 0.693] | $t_2 = 6.25$ | 0.1390 | 3 vs. 3 |
| Šídák | Planned comparison | REE vs. K | −2.09 [−3.90, −0.267] | $t_2 = 12.4$ | 0.0383 | 3 vs. 3 |
| Šídák | Planned comparison | REE vs. Mg | −1.52 [−2.90, −0.134] | $t_2 = 11.8$ | 0.0418 | 3 vs. 3 |
| Šídák | Planned comparison | P vs. Ca | −0.136 [−1.70, 1.43] | $t_2 = 0.930$ | 0.9724 | 3 vs. 3 |
| Šídák | Planned comparison | P vs. K | −1.26 [−2.90, 0.376] | $t_2 = 8.30$ | 0.0824 | 3 vs. 3 |
| Šídák | Planned comparison | P vs. Mg | −0.694 [−2.59, 1.21] | $t_2 = 3.93$ | 0.3054 | 3 vs. 3 |

**Table S18. Log-linear regression and extra sum-of-squares $F$ test comparing REE solubilization from REE-enriched *P. americana* shoots to the nominal $H_2SO_4$-to-REE ratio obtained by varying $H_2SO_4$ dose or biomass loading.** REE solubilization was fit as a function of the nominal $H_2SO_4$-to-REE ratio using the log-linear model $y = a\log_{10}(H_2SO_4 : REE + 1) + b$, where $y$ is REE solubilization (%). Fits were compared between datasets in which the nominal $H_2SO_4$-to-REE ratio was varied by changing $H_2SO_4$ concentration at fixed biomass loading or by changing biomass loading at fixed $H_2SO_4$ concentration. An extra sum-of-squares $F$ test compared the null model with one shared curve for both datasets against the alternative model with separate curves for each dataset. Parameter confidence intervals are profile-likelihood 95% confidence intervals.

| **Analysis** | **Model or dataset** | **Parameter or comparison** | **Estimate** [95% CI] | **Fit statistic** | ***P* value** | ***n*** |
|---|---|---|---|---|---|---|
| Extra sum-of-squares F test | Shared vs. separate curves | One shared curve for both datasets | Preferred model | $F_{2,68} = 0.789$ | 0.4584 | 72 |
| Global shared fit | Shared curve | Slope, $a$ | 18.1 [15.7, 20.6] | $R^2 = 0.760$ | – | 72 |
| Global shared fit | Shared curve | Intercept, $b$ | −0.736 [−5.90, 4.43] | $S_{y.x} = 15.3$ | – | 72 |
| Separate fit | $H_2SO_4$ varied at fixed biomass | Slope, $a$ | 17.6 [14.7, 20.5] | $R^2 = 0.741$ | – | 54 |
| Separate fit | $H_2SO_4$ varied at fixed biomass | Intercept, $b$ | 1.19 [−5.68, 8.07] | $S_{y.x} = 16.9$ | – | 54 |
| Separate fit | Biomass varied at fixed $H_2SO_4$ | Slope, $a$ | 22.6 [16.0, 29.2] | $R^2 = 0.768$ | – | 18 |
| Separate fit | Biomass varied at fixed $H_2SO_4$ | Intercept, $b$ | −7.21 [−14.0, −0.414] | $S_{y.x} = 8.20$ | – | 18 |

**Table S19. Log-linear regression and extra sum-of-squares $F$ test comparing concentration-dependent REE solubilization from REE-enriched *P. americana* shoots using equimolar dosages of $H_2SO_4$, $K_2SO_4$, and $KNO_3$.** REE solubilization was fit at the same nominal reagent concentrations shown in the highlighted band of Figure S14a using [REE solubilization $= a \cdot \log([\text{reactant}] + 1) + b$], where REE solubilization is expressed in % and [reactant] in M. An extra sum-of-squares $F$ test compared one curve shared by all three datasets with separate curves for each dataset. Parameter intervals are profile-likelihood 95% confidence intervals.

| **Analysis** | **Model or dataset** | **Parameter or comparison** | **Estimate** [95% CI] | **Fit statistic** | ***P* value** | ***n*** |
|---|---|---|---|---|---|---|
| Extra SS $F$ test | Shared vs. separate curves | At least one curve differs | — | $F_{4,30} = 43.1$ | $< 0.0001$ | 36 |
| Shared fit | Global shared curve | Slope, $a$ | 868 [30.8, 1710] | $R^2 = 0.115$ | — | 36 |
| Shared fit | Global shared curve | Intercept, $b$ | 15.4 [−7.13, 38.0] | $S_{y.x} = 37.5$ | — | 36 |
| Separate fit | $H_2SO_4$ | Slope, $a$ | 1620 [929, 2310] | $R^2 = 0.607$ | — | 18 |
| Separate fit | $H_2SO_4$ | Intercept, $b$ | 30.2 [11.6, 48.9] | $S_{y.x} = 21.0$ | — | 18 |
| Separate fit | $K_2SO_4$ | Slope, $a$ | 195 [115, 276] | $R^2 = 0.826$ | — | 9 |
| Separate fit | $K_2SO_4$ | Intercept, $b$ | −0.00110 [−2.16, 2.16] | $S_{y.x} = 1.54$ | — | 9 |
| Separate fit | $KNO_3$ | Slope, $a$ | 33.4 [2.00, 64.7] | $R^2 = 0.475$ | — | 9 |
| Separate fit | $KNO_3$ | Intercept, $b$ | 1.36 [0.510, 2.20] | $S_{y.x} = 0.603$ | — | 9 |

**Table S20. Brown–Forsythe and Welch ANOVAs and extra sum-of-squares $F$ tests of acid-dependent REE solubilization from REE-enriched *P. americana* shoots as a function of pH for some mineral and organic acids.** Brown–Forsythe and Welch tests compared $H_2SO_4$, $HNO_3$, HCl, and $CH_3SO_3H$ separately at each measured pH; their $P$ values are not adjusted across the four pH values. Extra sum-of-squares $F$ tests compared one shared pH-dependent curve with separate curves for the four acids. The matched-pH fit analyzed 54 observations; the expanded fit analyzed 78 observations, including additional $H_2SO_4$ observations. In both fits, $Y_0$ was fixed at 9.54 based on water-labile extraction (Figure S12). Parameter intervals are profile-likelihood 95% confidence intervals.

| **Dataset** | **Test/model** | **Effect or comparison** | **Estimate** [95% CI] | **Statistic or $R^2$** | ***P* value** | ***n*** |
|---|---|---|---|---|---|---|
| pH 1.19 | Brown–Forsythe | Four acids | — | $F^*_{3,5.65} = 2.42$ | 0.1702 | 12 |
| pH 1.19 | Welch | Four acids | — | $W_{3,4.13} = 3.94$ | 0.1057 | 12 |
| pH 1.29 | Brown–Forsythe | Four acids | — | $F^*_{3,9.68} = 0.792$ | 0.5266 | 15 |
| pH 1.29 | Welch | Four acids | — | $W_{3,5.24} = 1.11$ | 0.4230 | 15 |
| pH 1.90 | Brown–Forsythe | Four acids | — | $F^*_{3,3.07} = 2.07$ | 0.2791 | 15 |
| pH 1.90 | Welch | Four acids | — | $W_{3,4.85} = 0.660$ | 0.6121 | 15 |
| pH 2.99 | Brown–Forsythe | Four acids | — | $F^*_{3,4.26} = 2.67$ | 0.1762 | 12 |
| pH 2.99 | Welch | Four acids | — | $W_{3,3.44} = 7.85$ | 0.0493 | 12 |
| Matched-pH set | Extra SS $F$ test | Separate vs. shared curves | Separate curves favored | $F_{6,46} = 2.52$ | 0.0340 | 54 |
| Matched-pH set | Separate fit | $H_2SO_4$ | $A = 1080$ [354, 6305]; $k = 0.913$ [0.553, 1.52] | $R^2 = 0.800$ | — | 18 |
| Matched-pH set | Separate fit | $HNO_3$ | $A = 174$ [77.1, 581]; $k = 0.421$ [0.193, 0.826] | $R^2 = 0.700$ | — | 12 |
| Matched-pH set | Separate fit | HCl | $A = 706$ [279, 3406]; $k = 0.849$ [0.539, 1.41] | $R^2 = 0.917$ | — | 12 |
| Matched-pH set | Separate fit | $CH_3SO_3H$ | $A = 612$ [215, 4933]; $k = 0.768$ [0.430, 1.50] | $R^2 = 0.854$ | — | 12 |
| Matched-pH set | Shared fit | Four acids | $A = 497$ [292, 982]; $k = 0.705$ [0.532, 0.937] | $R^2 = 0.768$ | — | 54 |
| Expanded pool | Extra SS $F$ test | Separate vs. shared curves | Shared curve not rejected | $F_{6,64} = 0.881$ | 0.5141 | 78 |
| Expanded pool | Separate fit | $H_2SO_4$ | $A = 323$ [202, 540]; $k = 0.575$ [0.422, 0.762] | $R^2 = 0.785$ | — | 36 |
| Expanded pool | Separate fit | $HNO_3$ | $A = 175$ [77.1, 585]; $k = 0.422$ [0.194, 0.829] | $R^2 = 0.699$ | — | 12 |
| Expanded pool | Separate fit | HCl | $A = 544$ [203, 2862]; $k = 0.754$ [0.438, 1.33] | $R^2 = 0.866$ | — | 12 |
| Expanded pool | Separate fit | $CH_3SO_3H$ | $A = 617$ [220, 3515]; $k = 0.765$ [0.433, 1.37] | $R^2 = 0.855$ | — | 12 |
| Expanded pool | Shared fit | Four acids | $A = 336$ [238, 490]; $k = 0.593$ [0.480, 0.724] | $R^2 = 0.788$ | — | 78 |

**Table S21. Ordinary two-way ANOVA and Šídák-adjusted comparisons of acid and electrolyte effects on REE solubilization from REE-enriched *P. americana* shoots and its selectivity.** Ordinary two-way ANOVAs included the $H_2SO_4$ and $HNO_3$ formulations at pH 1.9, with and without 0.1 M $K_2SO_4$ or 0.2 M $KNO_3$. Estimates are the first condition minus the second. REE solubilization differences are in percentage points, and selectivity differences are in index units. Confidence intervals and $P$ values for cell comparisons are Šidák adjusted. The heteroscedasticity diagnostic was positive for REE solubilization ($P = 0.0023$).

| Outcome | Test/model | Result type | Effect or comparison | Estimate [95% CI] | Test statistic | *P* value | *n* |
|---|---|---|---|---|---|---|---|
| REE solubilization | Two-way ANOVA | Omnibus term | Acid identity | – | $F_{1,15} = 0.000371$ | 0.9849 | 21 |
| REE solubilization | Two-way ANOVA | Omnibus term | Electrolyte condition | – | $F_{2,15} = 8.31$ | 0.0037 | 21 |
| REE solubilization | Two-way ANOVA | Omnibus term | Acid × electrolyte | – | $F_{2,15} = 4.21$ | 0.0353 | 21 |
| REE solubilization | Šidák | Cell comparison | $H_2SO_4$: $+K_2SO_4$ vs. baseline | +26.7 [0.00444, 53.3] | – | 0.0499 | 3 vs. 6 |
| REE solubilization | Šidák | Cell comparison | $H_2SO_4$: $+KNO_3$ vs. baseline | −2.36 [−29.0, 24.3] | – | > 0.9999 | 3 vs. 6 |
| REE solubilization | Šidák | Cell comparison | $H_2SO_4$: $+K_2SO_4$ vs. $+KNO_3$ | +29.0 [−1.76, 59.8] | – | 0.0742 | 3 vs. 3 |
| REE solubilization | Šidák | Cell comparison | $HNO_3$: $+K_2SO_4$ vs. baseline | −5.14 [−35.9, 25.7] | – | > 0.9999 | 3 vs. 3 |
| REE solubilization | Šidák | Cell comparison | $HNO_3$: $+KNO_3$ vs. baseline | −27.0 [−57.8, 3.81] | – | 0.1166 | 3 vs. 3 |
| REE solubilization | Šidák | Cell comparison | $HNO_3$: $+K_2SO_4$ vs. $+KNO_3$ | +21.8 [−8.95, 52.6] | – | 0.3301 | 3 vs. 3 |
| Selectivity index | Two-way ANOVA | Omnibus term | Acid identity | – | $F_{1,15} = 3.17$ | 0.0955 | 21 |
| Selectivity index | Two-way ANOVA | Omnibus term | Electrolyte condition | – | $F_{2,15} = 2.22$ | 0.1432 | 21 |
| Selectivity index | Two-way ANOVA | Omnibus term | Acid × electrolyte | – | $F_{2,15} = 4.57$ | 0.0281 | 21 |
| Selectivity index | Šidák | Cell comparison | $H_2SO_4$: $+K_2SO_4$ vs. baseline | −0.0642 [−0.461, 0.332] | – | > 0.9999 | 3 vs. 6 |
| Selectivity index | Šidák | Cell comparison | $H_2SO_4$: $+KNO_3$ vs. baseline | +0.0278 [−0.369, 0.424] | – | > 0.9999 | 3 vs. 6 |
| Selectivity index | Šidák | Cell comparison | $H_2SO_4$: $+K_2SO_4$ vs. $+KNO_3$ | −0.0920 [−0.550, 0.366] | – | > 0.9999 | 3 vs. 3 |
| Selectivity index | Šidák | Cell comparison | $HNO_3$: $+K_2SO_4$ vs. baseline | +0.162 [−0.296, 0.620] | – | 0.9831 | 3 vs. 3 |
| Selectivity index | Šidák | Cell comparison | $HNO_3$: $+KNO_3$ vs. baseline | −0.306 [−0.764, 0.152] | – | 0.4125 | 3 vs. 3 |
| Selectivity index | Šidák | Cell comparison | $HNO_3$: $+K_2SO_4$ vs. $+KNO_3$ | +0.468 [0.0101, 0.926] | – | 0.0429 | 3 vs. 3 |

**Table S22. Linear regression and extra sum-of-squares $F$ test of the $Fe_2(SO_4)_3$ concentration effect on REE solubilization from REE-enriched *P. americana* shoots.** REE solubilization from Ce/Dy-enriched *P. americana* shoots was analyzed as a function of $Fe_2(SO_4)_3$ concentration at pH 1.8. A linear model with an unconstrained slope was compared against a constrained model with slope fixed to zero. Parameter confidence intervals are profile-likelihood 95% confidence intervals.

| Test/model | Result type | Effect or parameter | Estimate [95% CI] | Fit statistic | $P$ value | $n$ |
|---|---|---|---|---|---|---|
| Extra sum-of-squares F test | Model comparison | Unconstrained slope vs. slope fixed to zero | Slope = 0 preferred | $F_{1,19} = 0.122$ | 0.7309 | 21 |
| Unconstrained linear fit | Fit parameter | Slope | −0.00436 [−0.0305, 0.0218] | $R^2 = 0.00637$ | – | 21 |
| Unconstrained linear fit | Fit parameter | Intercept | 79.0 [69.8, 88.2] | $S_{y.x} = 15.2$ | – | 21 |
| Slope-constrained linear fit | Fit parameter | Intercept, slope fixed to zero | 78.0 [71.2, 84.8] | $S_{y.x} = 14.9$ | – | 21 |

**Table S23. Unpaired $t$ test and one-way ANOVA with Dunnett comparisons of chelator-mediated REE solubilization from REE-enriched *P. americana* shoots against matched-pH $H_2SO_4$ controls.** REE solubilization was compared between each chelator and the matched-pH $H_2SO_4$ condition. At pH 1.8, only $H_2SO_4$ and citric acid were tested, so an unpaired two-tailed $t$ test was used. At pH 3.0 and 4.5, one-way ANOVA followed by Dunnett's multiple comparisons test was used with $H_2SO_4$ as the control. $\overline{H_2SO_4}$ denotes the mean of the corresponding $H_2SO_4$ baseline control, and $\overline{\text{Chelator}}$ denotes the mean response using the corresponding chelator. Estimates and confidence intervals are reported as chelator minus $H_2SO_4$.

| pH | Test/model | Comparison | $\overline{H_2SO_4}$ (%) | $\overline{\text{Chelator}}$ (%) | Estimate [95% CI] | Test statistic | $P$ value | $n$ |
|---|---|---|---|---|---|---|---|---|
| 1.8 | Unpaired t test | citric acid vs. $H_2SO_4$ | 58.4 | 47.5 | −10.9 [−27.3, 5.56] | $t_4 = 1.84$ | 0.1402 | 3 vs. 3 |
| 3.0 | Dunnett | citric acid vs. $H_2SO_4$ | 12.5 | 6.90 | −5.63 [−29.6, 18.4] | $q_6 = 0.672$ | 0.7381 | 3 vs. 3 |
| 3.0 | Dunnett | $Na_2EDTA$ vs. $H_2SO_4$ | 12.5 | 81.7 | 69.2 [45.2, 93.2] | $q_6 = 8.25$ | 0.0003 | 3 vs. 3 |
| 4.5 | Dunnett | citric acid vs. $H_2SO_4$ | 0.00 | 50.9 | 50.9 [23.8, 77.9] | $q_6 = 5.38$ | 0.0031 | 3 vs. 3 |
| 4.5 | Dunnett | $Na_2EDTA$ vs. $H_2SO_4$ | 0.00 | 80.5 | 80.5 [53.4, 108] | $q_6 = 8.51$ | 0.0003 | 3 vs. 3 |

**Table S24. Two-way ANOVA and Šídák-adjusted comparisons of chelator identity and $In^{3+}$ pre-loading effects on REE solubilization from REE-enriched *P. americana* shoots.** REE solubilization from Ce/Dy-enriched *P. americana* shoots was measured after extraction with fixed-dose citric acid or $Na_2EDTA$ solutions pre-loaded with the indicated $In^{3+}$ concentrations. Omnibus effects were tested using two-way ANOVA with chelator identity and $In^{3+}$ dose as factors. Planned simple-effects comparisons tested each $In^{3+}$-preloaded condition against the corresponding no-$In^{3+}$ condition within each chelator. Estimates and confidence intervals are reported as no $In^{3+}$ minus the $In^{3+}$-preloaded condition, so positive values indicate lower REE solubilization after $In^{3+}$ pre-loading. Pairwise comparisons were performed using Šídák's multiple comparisons test.

| **Test/model** | **Result type** | **Effect or comparison** | **Estimate** [95% CI] | **Test statistic** | ***P* value** | ***n*** |
|---|---|---|---|---|---|---|
| Two-way ANOVA | Omnibus term | Chelator identity | – | $F_{1,12} = 84.5$ | $< 0.0001$ | 18 |
| Two-way ANOVA | Omnibus term | $In^{3+}$ dose | – | $F_{2,12} = 1.76$ | 0.2129 | 18 |
| Two-way ANOVA | Omnibus term | Chelator identity × $In^{3+}$ dose | – | $F_{2,12} = 5.50$ | 0.0201 | 18 |
| Šídák | Planned comparison | citric acid, no $In^{3+}$ vs. 1.25 mM $In^{3+}$ | 4.30 [−21.8, 30.4] | $t_{12} = 0.421$ | 0.8982 | 3 vs. 3 |
| Šídák | Planned comparison | citric acid, no $In^{3+}$ vs. 7.5 mM $In^{3+}$ | 28.6 [2.57, 54.7] | $t_{12} = 2.81$ | 0.0315 | 3 vs. 3 |
| Šídák | Planned comparison | $Na_2EDTA$, no $In^{3+}$ vs. 1.25 mM $In^{3+}$ | −21.6 [−47.7, 4.46] | $t_{12} = 2.12$ | 0.1087 | 3 vs. 3 |
| Šídák | Planned comparison | $Na_2EDTA$, no $In^{3+}$ vs. 7.5 mM $In^{3+}$ | −19.2 [−45.3, 6.86] | $t_{12} = 1.88$ | 0.1617 | 3 vs. 3 |

**Table S25. Multiple linear regression of $\log_2$-transformed HREE/LREE molar selectivity of REE solubilization from REE-enriched *P. americana* shoots by reagent, $KNO_3$ addition, and their interaction.** The response was $z = \log_2(\alpha_{\mathrm{HREE/LREE}})$, where $\alpha_{\mathrm{HREE/LREE}} = (n_{\mathrm{HREE,aq}}/n_{\mathrm{LREE,aq}})/(n_{\mathrm{HREE,feed}}/n_{\mathrm{LREE,feed}})$ is the feed-normalized molar selectivity. Coefficients are differences in $z$, with $H_2SO_4$ without added $KNO_3$ as the reference. Reagent coefficients compare lixiviants in the absence of $KNO_3$; the $KNO_3$ coefficient is its effect within $H_2SO_4$; interactions describe differences in the $KNO_3$ effect relative to $H_2SO_4$. The ordinary least-squares fit used 228 observations. Intervals are the 95% confidence intervals reported by Prism, and $P$ values are unadjusted. Residual normality was rejected (Shapiro–Wilk $W = 0.920$, $P < 0.0001$), so model-based intervals and $P$ values should be interpreted cautiously.

| **Model term** | **Interpretation** | $\beta$ [95% CI] | **Test statistic** | $P$ **value** |
|---|---|---|---|---|
| Intercept | $H_2SO_4$, without $KNO_3$ | 0.800 [0.618, 0.982] | $\lvert t\rvert = 8.67$ | $< 0.0001$ |
| Citric acid | Citric acid vs. $H_2SO_4$, without $KNO_3$ | 0.0756 [−0.292, 0.443] | $\lvert t\rvert = 0.405$ | 0.6856 |
| $Na_2EDTA$ | $Na_2EDTA$ vs. $H_2SO_4$, without $KNO_3$ | 0.438 [0.0704, 0.805] | $\lvert t\rvert = 2.35$ | 0.0197 |
| $HNO_3$ | $HNO_3$ vs. $H_2SO_4$, without $KNO_3$ | −0.936 [−1.42, −0.449] | $\lvert t\rvert = 3.79$ | 0.0002 |
| $KNO_3$ | With vs. without $KNO_3$, within $H_2SO_4$ | −0.950 [−1.25, −0.646] | $\lvert t\rvert = 6.15$ | $< 0.0001$ |
| Citric acid × $KNO_3$ | Difference in $KNO_3$ effect: citric acid vs. $H_2SO_4$ | 0.400 [−0.134, 0.934] | $\lvert t\rvert = 1.48$ | 0.1410 |
| $Na_2EDTA$ × $KNO_3$ | Difference in $KNO_3$ effect: $Na_2EDTA$ vs. $H_2SO_4$ | −0.243 [−0.791, 0.306] | $\lvert t\rvert = 0.871$ | 0.3844 |
| $HNO_3$ × $KNO_3$ | Difference in $KNO_3$ effect: $HNO_3$ vs. $H_2SO_4$ | 0.319 [−0.735, 1.37] | $\lvert t\rvert = 0.596$ | 0.5517 |
| Overall model | Eight-parameter regression, $n = 228$ | – | $F_{7,220} = 12.6$ | $< 0.0001$ |
| Goodness of fit | Model $R^2$ | 0.287 | – | – |

**Table S26. Log-linear regression and extra sum-of-squares $F$ test comparing Ni solubilization from Ni-enriched *P. acinosa* shoots versus reactant-to-Ni ratio obtained by varying the concentration of Ni supplementation during plant cultivation.** Ni solubilization from *P. acinosa* shoots supplemented with 25 or 50 mM $NiCl_2$ was fit as a function of the nominal reactant-to-Ni ratio using the log-linear model $\text{Ni solubilization} = a\log_{10}([\text{reactant}] : \text{Ni} + 1) + b$, where Ni solubilization is expressed in % and [reactant]:Ni as a molar ratio. An extra sum-of-squares $F$ test compared the null model with one shared curve across datasets against the alternative model with separate curves for each dataset. Parameter confidence intervals are profile-likelihood 95% confidence intervals.

| **Analysis** | **Model or dataset** | **Parameter or comparison** | **Estimate** [95% CI] | **Fit statistic** | ***P* value** | ***n*** |
|---|---|---|---|---|---|---|
| Extra sum-of-squares F test | Shared vs. separate curves | One shared curve across datasets | Preferred model | $F_{2,32} = 0.532$ | 0.5924 | 36 |
| Global shared fit | Shared curve | Slope, $a$ | 60.9 [43.3, 78.5] | $R^2 = 0.5928$ | – | 36 |
| Global shared fit | Shared curve | Intercept, $b$ | 47.3 [42.4, 52.2] | $S_{y.x} = 12.7$ | – | 36 |
| Separate fit | +25 mM $NiCl_2$ shoots | Slope, $a$ | 63.7 [43.1, 84.3] | $R^2 = 0.728$ | – | 18 |
| Separate fit | +25 mM $NiCl_2$ shoots | Intercept, $b$ | 44.7 [38.1, 51.4] | $S_{y.x} = 11.6$ | – | 18 |
| Separate fit | +50 mM $NiCl_2$ shoots | Slope, $a$ | 58.0 [22.5, 93.5] | $R^2 = 0.429$ | – | 18 |
| Separate fit | +50 mM $NiCl_2$ shoots | Intercept, $b$ | 49.7 [41.7, 57.7] | $S_{y.x} = 14.0$ | – | 18 |

**Table S27. Geisser–Greenhouse-corrected two-way repeated-measures ANOVA and Dunnett comparisons of electrolyte-dependent REE solubilization from REE-enriched *P. americana* shoots.** REE solubilization in the one-day pH 4.5 electrolyte-leaching experiment was analyzed by two-way repeated-measures ANOVA with time and electrolyte condition as factors. Sphericity was not assumed, and Geisser–Greenhouse correction was applied to time-dependent terms. Simple-effects comparisons were performed within each time point using Dunnett's multiple comparisons test against the $H_2SO_4$ baseline. Estimates are reported as $H_2SO_4$ minus the indicated electrolyte condition, so negative values indicate greater REE release than the $H_2SO_4$ baseline.

| **Test/model** | **Result type** | **Time (h)** | **Effect or comparison** | **Estimate [95% CI]** | **Test statistic** | ***P* value** | ***n*** |
|---|---|---|---|---|---|---|---|
| Two-way RM ANOVA | Omnibus term | – | Time × electrolyte | – | $F_{7.46,19.9} = 9.21$ | < 0.0001 | 12 |
| Two-way RM ANOVA | Omnibus term | – | Time | – | $F_{2.49,19.9} = 28.7$ | < 0.0001 | 12 |
| Two-way RM ANOVA | Omnibus term | – | Electrolyte | – | $F_{3,8} = 118$ | < 0.0001 | 12 |
| Two-way RM ANOVA | Omnibus term | – | Subject | – | $F_{8,56} = 2.00$ | 0.0629 | 12 |
| Dunnett | Simple effect | 0.25 | $H_2SO_4$ vs. +200 mM $KNO_3$ | −0.133 [−0.697, 0.430] | $q_{3.67} = 0.894$ | 0.7216 | 3 vs. 3 |
| Dunnett | Simple effect | 0.25 | $H_2SO_4$ vs. +100 mM $K_2SO_4$ | −2.10 [−2.75, −1.45] | $q_{3.96} = 11.7$ | 0.0007 | 3 vs. 3 |
| Dunnett | Simple effect | 0.25 | $H_2SO_4$ vs. +100 mM $(NH_4)_2SO_4$ | −0.167 [−0.752, 0.418] | $q_{2.88} = 1.25$ | 0.5448 | 3 vs. 3 |
| Dunnett | Simple effect | 0.5 | $H_2SO_4$ vs. +200 mM $KNO_3$ | −1.03 [−3.77, 1.71] | $q_{2.08} = 2.19$ | 0.2853 | 3 vs. 3 |
| Dunnett | Simple effect | 0.5 | $H_2SO_4$ vs. +100 mM $K_2SO_4$ | −2.33 [−2.90, −1.76] | $q_{3.12} = 17.0$ | 0.0008 | 3 vs. 3 |
| Dunnett | Simple effect | 0.5 | $H_2SO_4$ vs. +100 mM $(NH_4)_2SO_4$ | −0.467 [−0.871, −0.0623] | $q_{2.00} = 7.00$ | 0.0380 | 3 vs. 3 |
| Dunnett | Simple effect | 1 | $H_2SO_4$ vs. +200 mM $KNO_3$ | −0.467 [−1.54, 0.604] | $q_{2.43} = 2.19$ | 0.2627 | 3 vs. 3 |
| Dunnett | Simple effect | 1 | $H_2SO_4$ vs. +100 mM $K_2SO_4$ | −2.27 [−2.84, −1.70] | $q_{3.12} = 16.5$ | 0.0008 | 3 vs. 3 |
| Dunnett | Simple effect | 1 | $H_2SO_4$ vs. +100 mM $(NH_4)_2SO_4$ | −0.667 [−0.989, −0.345] | $q_{2.94} = 8.94$ | 0.0068 | 3 vs. 3 |
| Dunnett | Simple effect | 2 | $H_2SO_4$ vs. +200 mM $KNO_3$ | −0.433 [−0.960, 0.0932] | $q_{2.44} = 4.11$ | 0.0768 | 3 vs. 3 |
| Dunnett | Simple effect | 2 | $H_2SO_4$ vs. +100 mM $K_2SO_4$ | −1.90 [−2.41, −1.39] | $q_{4.00} = 13.4$ | 0.0004 | 3 vs. 3 |
| Dunnett | Simple effect | 2 | $H_2SO_4$ vs. +100 mM $(NH_4)_2SO_4$ | −0.567 [−1.09, −0.0402] | $q_{2.44} = 5.38$ | 0.0424 | 3 vs. 3 |
| Dunnett | Simple effect | 4 | $H_2SO_4$ vs. +200 mM $KNO_3$ | −0.467 [−0.919, −0.0145] | $q_{2.56} = 4.95$ | 0.0464 | 3 vs. 3 |
| Dunnett | Simple effect | 4 | $H_2SO_4$ vs. +100 mM $K_2SO_4$ | −1.53 [−2.02, −1.05] | $q_{3.94} = 11.5$ | 0.0008 | 3 vs. 3 |
| Dunnett | Simple effect | 4 | $H_2SO_4$ vs. +100 mM $(NH_4)_2SO_4$ | −0.467 [−0.881, −0.0521] | $q_{3.72} = 4.22$ | 0.0348 | 3 vs. 3 |
| Dunnett | Simple effect | 8 | $H_2SO_4$ vs. +200 mM $KNO_3$ | −0.167 [−1.44, 1.11] | $q_{2.09} = 0.754$ | 0.8055 | 3 vs. 3 |
| Dunnett | Simple effect | 8 | $H_2SO_4$ vs. +100 mM $K_2SO_4$ | −1.07 [−2.09, −0.0456] | $q_{3.48} = 4.06$ | 0.0440 | 3 vs. 3 |
| Dunnett | Simple effect | 8 | $H_2SO_4$ vs. +100 mM $(NH_4)_2SO_4$ | −0.333 [−1.53, 0.868] | $q_{2.28} = 1.47$ | 0.4704 | 3 vs. 3 |
| Dunnett | Simple effect | 24 | $H_2SO_4$ vs. +200 mM $KNO_3$ | −0.267 [−1.31, 0.779] | $q_{3.04} = 1.08$ | 0.6269 | 3 vs. 3 |
| Dunnett | Simple effect | 24 | $H_2SO_4$ vs. +100 mM $K_2SO_4$ | −1.03 [−1.72, −0.345] | $q_{3.81} = 5.57$ | 0.0133 | 3 vs. 3 |
| Dunnett | Simple effect | 24 | $H_2SO_4$ vs. +100 mM $(NH_4)_2SO_4$ | −0.0333 [−0.658, 0.592] | $q_{2.33} = 0.277$ | 0.9826 | 3 vs. 3 |

**Table S28. One-sample $t$ tests, Brown–Forsythe and Welch ANOVAs, and Dunnett's T3 comparisons of $\log_{10}$-transformed REE reload efficiency of (REE-enriched) *P. americana* shoots.** Operational reload efficiency, $R_{\mathrm{REE,reload}}$, was calculated as the amount of REE taken up during the reload step divided by the amount released during the preceding leaching step. Values were analyzed after $\log_{10}$ transformation, so a value of 0 corresponds to $R_{\mathrm{REE,reload}} = 1$. Two-tailed one-sample $t$ tests assessed whether mean $\log_{10}(R_{\mathrm{REE,reload}})$ differed from 0. For individual conditions, estimates are mean $\log_{10}(R_{\mathrm{REE,reload}})$ values with 95% confidence intervals; geometric means were obtained by back-transformation. One-sample $t$ statistics are reported as absolute values, and their $P$ values are unadjusted. Conditions marked with # were not statistically analyzed because corrected reload values were non-positive or insufficiently replicated. The four matched 1 d, pH 4.5 leaching conditions were additionally compared using Brown–Forsythe and Welch one-way ANOVAs. Dunnett's T3 comparisons with the matched $H_2SO_4$ baseline are reported as differences in mean $\log_{10}(R_{\mathrm{REE,reload}})$, calculated as the baseline minus the supplemented condition. Dunnett's T3 $P$ values are multiplicity adjusted. Confidence intervals are reported for the one-sample analyses only. For omnibus analyses, $n$ is the total number of observations; for pairwise comparisons, sample sizes are shown for both groups.

| Test/model | Condition or comparison | Estimate [95% CI] | Geometric mean $R_{\mathrm{REE,reload}}$ | Test statistic | $P$ value | $n$ | Interpretation |
|---|---|---|---|---|---|---|---|
| One-sample $t$ test | $H_2SO_4$, pH 4.5 | 1.81 [0.745, 2.87] | 64.6 | $t_2 = 7.31$ | 0.0182 | 3 | Reload efficiency above unity |
| One-sample $t$ test | +200 mM $KNO_3$ | 1.71 [0.903, 2.51] | 51.3 | $t_2 = 9.13$ | 0.0118 | 3 | Reload efficiency above unity |
| One-sample $t$ test | +100 mM $K_2SO_4$ | −1.80 [−2.23, −1.37] | 0.0158 | $t_2 = 17.9$ | 0.0031 | 3 | Reload efficiency below unity |
| One-sample $t$ test | +100 mM $(NH_4)_2SO_4$ | 1.73 [1.13, 2.33] | 53.7 | $t_2 = 12.4$ | 0.0065 | 3 | Reload efficiency above unity |
| Not analyzed | $H_2SO_4$, pH 1.8# | – | – | – | – | – | Not log-plotted; corrected reload $\leq 0$ |
| One-sample $t$ test | $H_2SO_4$, pH 3.0 | 0.164 [−0.258, 0.587] | 1.46 | $t_2 = 1.67$ | 0.2364 | 3 | Not resolved as different from unity |
| One-sample $t$ test | Citric acid | −1.48 [−2.08, −0.882] | 0.0331 | $t_2 = 10.6$ | 0.0087 | 3 | Reload efficiency below unity |
| Descriptive | $Na_2EDTA$# | −0.387 | 0.410 | – | – | 1 | Reported descriptively; insufficient replication for one-sample test |
| Brown–Forsythe ANOVA | Four matched 1 d, pH 4.5 conditions | – | – | $F^*_{3,5.80} = 100$ | < 0.0001 | 12 | Difference among condition means |
| Welch's ANOVA | Four matched 1 d, pH 4.5 conditions | – | – | $W_{3,4.25} = 155$ | < 0.0001 | 12 | Difference among condition means |
| Dunnett's T3 | $H_2SO_4$ vs. +200 mM $KNO_3$ | 0.102 | – | – | 0.9808 | 3 vs. 3 | Difference not statistically resolved |
| Dunnett's T3 | $H_2SO_4$ vs. +100 mM $K_2SO_4$ | 3.61 | – | – | 0.0021 | 3 vs. 3 | Reload efficiency lower after $K_2SO_4$ leaching than after the matched $H_2SO_4$ baseline |
| Dunnett's T3 | $H_2SO_4$ vs. +100 mM $(NH_4)_2SO_4$ | 0.0798 | – | – | 0.9872 | 3 vs. 3 | Difference not statistically resolved |

**Table S29. Wilcoxon signed-rank tests of $\log_{10}$-transformed REE loading ratios relative to the non-leached REE-enriched *P. americana* reference loading after reloading leached REE-enriched *P. americana* residues.** Final corrected REE loadings after the standardized reload step were normalized to the non-leached Ce/Dy-enriched *P. americana* reference loading and $\log_{10}$ transformed. For each prior-leaching condition, a two-tailed exact one-sample Wilcoxon signed-rank test evaluated departure from zero, corresponding to a loading ratio of unity. Median $\log_{10}$ ratios are reported with 75% confidence intervals, and median ratios were obtained by base-10 back-transformation. Each condition included three replicate values. With $n = 3$, the smallest attainable two-tailed exact $P$ value is 0.25; therefore, the absence of a statistically detected difference does not establish preservation or equivalence of endpoint REE loading. Rows are ordered to match Figure 3c.

| **Prior treatment** | **Median $\log_{10}$ ratio** [75% CI] | **Median ratio** | **Signed-rank statistic** | ***P* value** | ***n*** | **Interpretation** |
|---|---|---|---|---|---|---|
| **1 d leach, 0 d storage** | | | | | | |
| $H_2SO_4$, pH 4.5 | −0.224 [−0.549, 0.101] | 0.60 | $W = -4.00$ | 0.5000 | 3 | Not resolved as different from reference loading |
| +200 mM $KNO_3$ | −0.380 [−0.419, 0.249] | 0.42 | $W = -4.00$ | 0.5000 | 3 | Not resolved as different from reference loading |
| +100 mM $K_2SO_4$ | −0.554 [−0.663, −0.0358] | 0.28 | $W = -6.00$ | 0.2500 | 3 | Not resolved as different from reference loading |
| +100 mM $(NH_4)_2SO_4$ | −0.233 [−0.468, 0.276] | 0.58 | $W = -2.00$ | 0.7500 | 3 | Not resolved as different from reference loading |
| **7 d leach, >100 d storage** | | | | | | |
| $H_2SO_4$, pH 1.8 | −0.841 [−1.02, −0.389] | 0.14 | $W = -6.00$ | 0.2500 | 3 | Not resolved as different from reference loading |
| $H_2SO_4$, pH 3.0 | −0.398 [−0.475, 0.258] | 0.40 | $W = -4.00$ | 0.5000 | 3 | Not resolved as different from reference loading |
| Citric acid | −0.397 [−1.01, −0.140] | 0.40 | $W = -6.00$ | 0.2500 | 3 | Not resolved as different from reference loading |
| $Na_2EDTA$ | −0.884 [−0.889, −0.605] | 0.13 | $W = -6.00$ | 0.2500 | 3 | Not resolved as different from reference loading |

**Table S30. Brown–Forsythe and Welch ANOVAs with Dunnett's T3 comparisons of $\log_{10}$-transformed Tb retention and inferred binding capacity of *A. ferrooxidans* across incubation times.** Statistical analysis for time-course measurements are shown in Figure S31. Retention, expressed on a 0–1 scale for these statistical analyses, and inferred binding capacity, $q_{\text{inferred}}$, were analyzed separately after $\log_{10}$ transformation. The corresponding retained percentage is defined in Equation S52. Here, $q_{\text{measured}}$ is the final pellet capacity after two EDTA and two water washes. Both capacities are expressed in mg Tb $\text{kg}^{-1}$ initial DCW. Unpaired Brown–Forsythe and Welch ANOVAs compared four incubation times (2, 4, 8, and 24 h; $n = 3$ per time). Dunnett's T3 comparisons used one family of three comparisons against 2 h for each response. Estimates are mean differences on the $\log_{10}$ scale, expressed as the later time minus 2 h; pairwise confidence intervals and $P$ values are multiplicity-adjusted within each response, but not across responses. Fold changes are ratios of geometric means, calculated by base-10 back-transformation of the reported differences. The $t$ statistics are absolute values. For omnibus rows, $n$ is the total number of observations; for pairwise rows, sample sizes are shown for the two groups.

| Response | Test/model | Effect or comparison | Estimate [95% CI] | Fold change | Test statistic | $P$ value | $n$ |
|---|---|---|---|---|---|---|---|
| $f$ | Brown–Forsythe ANOVA | Incubation time | – | – | $F^*_{3.00,5.01} = 12.9$ | 0.0085 | 12 |
| $f$ | Welch's ANOVA | Incubation time | – | – | $W_{3.00,3.92} = 23.1$ | 0.0059 | 12 |
| $f$ | Dunnett's T3 | 4 vs. 2 h | 0.178 [−0.438, 0.795] | 1.51× | $t_{3.92} = 1.08$ | 0.6582 | 3 vs. 3 |
| $f$ | Dunnett's T3 | 8 vs. 2 h | 0.314 [−0.224, 0.852] | 2.06× | $t_{2.99} = 2.59$ | 0.1822 | 3 vs. 3 |
| $f$ | Dunnett's T3 | 24 vs. 2 h | 0.743 [0.0341, 1.45] | 5.53× | $t_{2.29} = 6.65$ | 0.0457 | 3 vs. 3 |
| $q_{\text{inferred}}$ | Brown–Forsythe ANOVA | Incubation time | – | – | $F^*_{3.00,5.96} = 0.661$ | 0.6055 | 12 |
| $q_{\text{inferred}}$ | Welch's ANOVA | Incubation time | – | – | $W_{3.00,4.34} = 0.384$ | 0.7705 | 12 |
| $q_{\text{inferred}}$ | Dunnett's T3 | 4 vs. 2 h | 0.0749 [−0.306, 0.456] | 1.19× | $t_{3.55} = 0.737$ | 0.8427 | 3 vs. 3 |
| $q_{\text{inferred}}$ | Dunnett's T3 | 8 vs. 2 h | 0.0769 [−0.335, 0.489] | 1.19× | $t_{2.89} = 0.827$ | 0.7989 | 3 vs. 3 |
| $q_{\text{inferred}}$ | Dunnett's T3 | 24 vs. 2 h | 0.116 [−0.310, 0.542] | 1.31× | $t_{3.16} = 1.21$ | 0.6039 | 3 vs. 3 |

**Table S31. Brown–Forsythe and Welch ANOVAs with Dunnett's T3 comparisons of Tb retention by *A. ferrooxidans* across EDTA washing protocols after 24 h of binding.** Following 24 h of binding at the highest $Tb_2(SO_4)_3$ concentration tested in Figure 4, retention was compared among seven washing conditions ($n = 3$ per condition). Retention was expressed on a 0–1 scale for these statistical analyses; the corresponding retained percentage is defined in Equation S52. Here, $q_{\text{measured}}$ is the final pellet binding capacity determined by digestion, and $q_{\text{inferred}}$ is the inferred prewash binding capacity; both use the same initial dry cell weight basis. Single EDTA exposures of 1, 5, 15, 30, or 60 min and the standard two-stage EDTA protocol were each followed by two water washes. Water-only controls received two water washes without EDTA. Unpaired Brown–Forsythe and Welch ANOVAs analyzed the untransformed retained fraction. Dunnett's T3 comparisons comprised one family of 11 comparisons against the water-only and standard two-stage protocols. Estimates are differences in mean retained fraction (first listed condition minus second); pairwise confidence intervals and $P$ values are multiplicity-adjusted across this family. The $t$ statistics are absolute values. For omnibus rows, $n$ is the total number of observations; for pairwise rows, sample sizes are shown for the two groups.

| Response | Test/model | Effect or comparison | Estimate [95% CI] | Test statistic | $P$ value | $n$ |
|---|---|---|---|---|---|---|
| $f$ | Brown–Forsythe ANOVA | Washing condition | – | $F^*_{6.00,7.21} = 30.0$ | $< 0.0001$ | 21 |
| $f$ | Welch's ANOVA | Washing condition | – | $W_{6.00,6.00} = 13.8$ | 0.0028 | 21 |
| $f$ | Dunnett's T3 | 1 min vs. water-only | −0.165 [−0.391, 0.0600] | $t_{3.41} = 4.34$ | 0.1128 | 3 vs. 3 |
| $f$ | Dunnett's T3 | 5 min vs. water-only | −0.270 [−0.472, −0.0669] | $t_{2.56} = 7.87$ | 0.0226 | 3 vs. 3 |
| $f$ | Dunnett's T3 | 15 min vs. water-only | −0.309 [−0.513, −0.106] | $t_{2.59} = 9.00$ | 0.0154 | 3 vs. 3 |
| $f$ | Dunnett's T3 | 30 min vs. water-only | −0.299 [−0.584, −0.0131] | $t_{2.15} = 9.14$ | 0.0458 | 3 vs. 3 |
| $f$ | Dunnett's T3 | 60 min vs. water-only | −0.316 [−0.552, −0.0809] | $t_{3.95} = 6.58$ | 0.0177 | 3 vs. 3 |
| $f$ | Dunnett's T3 | Two-stage vs. water-only | −0.317 [−0.524, −0.110] | $t_{2.74} = 9.07$ | 0.0150 | 3 vs. 3 |
| $f$ | Dunnett's T3 | 1 min vs. two-stage | 0.152 [0.0300, 0.274] | $t_{3.52} = 6.10$ | 0.0233 | 3 vs. 3 |
| $f$ | Dunnett's T3 | 5 min vs. two-stage | 0.0477 [−0.0428, 0.138] | $t_{3.92} = 2.58$ | 0.3189 | 3 vs. 3 |
| $f$ | Dunnett's T3 | 15 min vs. two-stage | 0.00783 [−0.0840, 0.0997] | $t_{3.95} = 0.418$ | $> 0.9999$ | 3 vs. 3 |
| $f$ | Dunnett's T3 | 30 min vs. two-stage | 0.0188 [−0.0720, 0.110] | $t_{2.77} = 1.22$ | 0.8759 | 3 vs. 3 |
| $f$ | Dunnett's T3 | 60 min vs. two-stage | 0.000847 [NR] | $t_{2.60} = 0.0220$ | $> 0.9999$ | 3 vs. 3 |

**Table S32. Ordinary two-way ANOVA of $\log_{10}$-transformed inferred Tb recovery by *A. ferrooxidans* across repeated adsorption–desorption cycles.** Inferred Tb values were $\log_{10}$ transformed prior to analysis. Omnibus effects were tested by ordinary two-way ANOVA with sorption–elution cycle and recovery step as factors. The adsorption–desorption contrast summarizes the marginal effect of recovery step across all cycles. Estimates and confidence intervals are reported on the $\log_{10}$ scale.

| Effect or comparison | Result type | Estimate [95% CI] | Test statistic | $P$ value | n |
|---|---|---|---|---|---|
| Cycle | Omnibus term | – | $F_{2,30} = 0.764$ | 0.4746 | 36 |
| Recovery step | Omnibus term | – | $F_{1,30} = 1.40$ | 0.2459 | 36 |
| Cycle × recovery step | Omnibus term | – | $F_{2,30} = 0.00257$ | 0.9974 | 36 |
| Adsorption vs. desorption | Marginal contrast | 0.0791 [−0.0574, 0.216] | – | – | 36 |

**Table S33. Šídák- and Tukey-adjusted comparisons from two-way ANOVAs of $\log_{10}$-transformed Tb binding, citrate elution, and recovery across some *A. ferrooxidans* pretreatments and repeated adsorption–desorption cycles.** Inferred binding capacity ($q_B$), citrate-eluted capacity ($q_E$), and apparent citrate recovery ($r_{E/B} = q_E/q_B$) were analyzed after $\log_{10}$ transformation using the supplied ordinary two-way ANOVAs (treatment × cycle; $n = 3$ per treatment–cycle combination). Ratios compare geometric means of the first listed group with the second. The reported multiplicity-adjusted 95% confidence intervals and $P$ values are retained from the original analyses: Šídák comparisons for capacities and Tukey comparisons for $r_{E/B}$. Adjustments covered three comparisons per family, with three cycle-specific families per capacity analysis and six row/column families for recovery; they were not recalculated for this selected subset and do not extend across families or responses. Samples sizes $n$ are shown for the two groups for pairwise rows.

| **Response** | **Comparison** | **Geometric mean ratio** [95% CI] | **Adjusted *P*** | **n** |
|---|---|---|---|---|
| Inferred binding, $q_B$ | Heat vs. untreated, cycle 1 | 1.58 [1.03, 2.43] | 0.0354 | 3 vs. 3 |
| Inferred binding, $q_B$ | Heat vs. untreated, cycle 2 | 1.71 [1.11, 2.62] | 0.0128 | 3 vs. 3 |
| Inferred binding, $q_B$ | Heat vs. untreated, cycle 3 | 1.85 [1.2, 2.84] | 0.0042 | 3 vs. 3 |
| Citrate elution, $q_E$ | Heat vs. untreated, cycle 1 | 1.82 [1.28, 2.6] | 0.0009 | 3 vs. 3 |
| Citrate elution, $q_E$ | Heat vs. untreated, cycle 2 | 2.23 [1.56, 3.18] | $< 0.0001$ | 3 vs. 3 |
| Citrate elution, $q_E$ | Heat vs. untreated, cycle 3 | 2.59 [1.82, 3.7] | $< 0.0001$ | 3 vs. 3 |
| Apparent recovery, $r_{E/B}$ | Heat vs. untreated, cycle 2 | 1.31 [1.11, 1.54] | 0.0015 | 3 vs. 3 |
| Apparent recovery, $r_{E/B}$ | UV-C vs. untreated, cycle 2 | 1.22 [1.03, 1.43] | 0.0175 | 3 vs. 3 |
| Apparent recovery, $r_{E/B}$ | Heat vs. untreated, cycle 3 | 1.4 [1.19, 1.65] | 0.0001 | 3 vs. 3 |
| Apparent recovery, $r_{E/B}$ | UV-C vs. untreated, cycle 3 | 1.33 [1.13, 1.57] | 0.0007 | 3 vs. 3 |
| Apparent recovery, $r_{E/B}$ | Untreated: cycle 3 vs. cycle 1 | 0.832 [0.706, 0.979] | 0.0251 | 3 vs. 3 |

**Table S34. Brown–Forsythe and Welch one-way ANOVAs with Dunnett's T3 comparisons of some membrane-associated fluorescence assay responses as a function of *A. ferrooxidans* pretreatment.** Baseline-corrected fluorescence from propidium iodide (PI), NBD-PEN, and $DiBAC_4(3)$ assays was analyzed using Brown–Forsythe and Welch one-way ANOVA tests without assuming equal variances. Dunnett's T3 multiple comparisons tests compared each treatment against untreated cells (AF). Estimates are mean differences in baseline-corrected relative fluorescence units (RFU), calculated as untreated cells (AF) minus the indicated pretreated cells. Negative estimates therefore indicate greater fluorescence after treatment. Pairwise confidence intervals and $P$ values are adjusted across the three comparisons within each assay, with familywise $\alpha = 0.05$; no adjustment across assays was applied. Omnibus $P$ values are unadjusted. Each assay included three analyzed replicate values per condition. The $n$ column reports the total number of values for omnibus tests and the numbers in the two groups for pairwise comparisons. Triton X-100 denotes detergent treatment followed by sonication.

| **Response** | **Test/model** | **Result type** | **Effect or comparison** | **Estimate** [95% CI] | **Test statistic** | ***P* value** | ***n*** |
|---|---|---|---|---|---|---|---|
| PI | Brown–Forsythe ANOVA | Omnibus term | Treatment | – | $F^*_{3.00,5.50} = 23.0$ | 0.0016 | 12 |
| PI | Welch's ANOVA | Omnibus term | Treatment | – | $W_{3.00,4.07} = 11.5$ | 0.0188 | 12 |
| PI | Dunnett's T3 | Planned comparison | AF vs. heat | −8.63 [−16.7, −0.533] | $t_{2.39} = 6.76$ | 0.0443 | 3 vs. 3 |
| PI | Dunnett's T3 | Planned comparison | AF vs. UV-C | −0.633 [−4.15, 2.88] | $t_{3.10} = 0.798$ | 0.8134 | 3 vs. 3 |
| PI | Dunnett's T3 | Planned comparison | AF vs. Triton X-100 | −0.637 [−5.00, 3.73] | $t_{2.68} = 0.646$ | 0.8834 | 3 vs. 3 |
| NBD-PEN | Brown–Forsythe ANOVA | Omnibus term | Treatment | – | $F^*_{3.00,5.47} = 0.0908$ | 0.9622 | 12 |
| NBD-PEN | Welch's ANOVA | Omnibus term | Treatment | – | $W_{3.00,4.13} = 0.0688$ | 0.9737 | 12 |
| NBD-PEN | Dunnett's T3 | Planned comparison | AF vs. heat | −0.147 [−3.03, 2.74] | $t_{2.76} = 0.225$ | 0.9932 | 3 vs. 3 |
| NBD-PEN | Dunnett's T3 | Planned comparison | AF vs. UV-C | −0.220 [−5.54, 5.10] | $t_{2.44} = 0.262$ | 0.9888 | 3 vs. 3 |
| NBD-PEN | Dunnett's T3 | Planned comparison | AF vs. Triton X-100 | 0.160 [−2.09, 2.41] | $t_{3.31} = 0.315$ | 0.9823 | 3 vs. 3 |
| $DiBAC_4(3)$ | Brown–Forsythe ANOVA | Omnibus term | Treatment | – | $F^*_{3.00,7.37} = 2764$ | < 0.0001 | 12 |
| $DiBAC_4(3)$ | Welch's ANOVA | Omnibus term | Treatment | – | $W_{3.00,4.42} = 1627$ | < 0.0001 | 12 |
| $DiBAC_4(3)$ | Dunnett's T3 | Planned comparison | AF vs. heat | −1985 [−2089, −1881] | $t_{3.65} = 71.8$ | < 0.0001 | 3 vs. 3 |
| $DiBAC_4(3)$ | Dunnett's T3 | Planned comparison | AF vs. UV-C | −2038 [−2143, −1933] | $t_{3.71} = 72.8$ | < 0.0001 | 3 vs. 3 |
| $DiBAC_4(3)$ | Dunnett's T3 | Planned comparison | AF vs. Triton X-100 | −1634 [−1738, −1529] | $t_{3.72} = 58.3$ | < 0.0001 | 3 vs. 3 |

**Table S35. One-way ANOVA and Šídák-adjusted comparisons of $\log_{10}$-transformed cell-associated Tb after *A. ferrooxidans* growth in Tb-supplemented medium and subsequent binding tests.** Cell-associated Tb values were $\log_{10}$ transformed prior to analysis. Omnibus effects were tested using ordinary one-way ANOVA across the five indicated conditions. Planned pairwise comparisons were performed using Šídák's multiple comparisons test. Estimates are reported on the $\log_{10}$ scale in the direction listed for each comparison. Fold changes were calculated as $10^{\Delta \log_{10}}$. The presence and absence of 30 µM $Tb^{3+}$ in the F2S medium during prior growth is indicated by + $Tb^{3+}$ and – $Tb^{3+}$, respectively. All binding tests were performed using 12 mM $Tb^{3+}$.

| **Test/model** | **Result type** | **Effect or comparison** | **Estimate** [95% CI] | **Fold change** | **Test statistic** | ***P* value** | ***n*** |
|---|---|---|---|---|---|---|---|
| One-way ANOVA | Omnibus term | Treatment | – | – | $F_{4,17} = 318$ | < 0.0001 | 22 |
| Šídák | Planned comparison | Growth endpoint + $Tb^{3+}$ vs. growth endpoint – $Tb^{3+}$ | 1.02 [0.793, 1.26] | 10.5× | $t_{17} = 12.8$ | < 0.0001 | 6 vs. 6 |
| Šídák | Planned comparison | $Tb^{3+}$ binding – $Tb^{3+}$ vs. $Tb^{3+}$ binding + $Tb^{3+}$ | 0.0453 [−0.282, 0.373] | 1.11× | $t_{17} = 0.400$ | 0.9973 | 3 vs. 3 |
| Šídák | Planned comparison | $Tb^{3+}$ binding + $Tb^{3+}$ vs. mixed-REE binding + $Tb^{3+}$ | 0.674 [0.367, 0.980] | 4.72× | $t_{17} = 6.35$ | < 0.0001 | 3 vs. 4 |
| Šídák | Planned comparison | $Tb^{3+}$ binding – $Tb^{3+}$ vs. growth endpoint + $Tb^{3+}$ | 1.71 [1.43, 2.00] | 51.3× | $t_{17} = 17.5$ | < 0.0001 | 3 vs. 6 |
| Šídák | Planned comparison | $Tb^{3+}$ binding + $Tb^{3+}$ vs. growth endpoint + $Tb^{3+}$ | 1.67 [1.39, 1.95] | 46.8× | $t_{17} = 17.0$ | < 0.0001 | 3 vs. 6 |

**Table S36. Two-way ANOVA and Šídák-adjusted comparisons of $\log_{10}$-transformed REE-binding capacity in *A. ferrooxidans* after growth in F2S medium with or without Tb supplementation.** *A. ferrooxidans* were grown in F2S medium with or without 30 $\mu$M $Tb^{3+}$, then exposed to 0.95 mM $Tb^{3+}$, 9.5 mM $Tb^{3+}$, or 5.9 mM equimolar Ce/Dy/Nd/Tb mixed-REE simulant solutions at pH 1.8. Inferred cell-associated REE values were $\log_{10}$ transformed prior to analysis. Omnibus effects were tested using ordinary two-way ANOVA with binding challenge and growth condition as factors. Pairwise comparisons were performed using Šídák's multiple comparisons test. Estimates and confidence intervals are reported on the $\log_{10}$ scale. Fold changes were calculated as $10^{\Delta \log_{10}}$.

| **Test/model** | **Result type** | **Effect or comparison** | **Estimate** [95% CI] | **Fold change** | **Test statistic** | ***P* value** | ***n*** |
|---|---|---|---|---|---|---|---|
| Two-way ANOVA | Omnibus term | Binding challenge | – | – | $F_{2,48} = 61.4$ | < 0.0001 | 54 |
| Two-way ANOVA | Omnibus term | Growth condition | – | – | $F_{1,48} = 1.01$ | 0.3207 | 54 |
| Two-way ANOVA | Omnibus term | Challenge × growth condition | – | – | $F_{2,48} = 0.195$ | 0.8233 | 54 |
| Šídák | Planned comparison | F2S − Tb vs. F2S + Tb, 0.95 mM $Tb^{3+}$ | 0.162 [−0.178, 0.501] | 1.45× | $t_{48} = 0.959$ | 0.3425 | 9 vs. 9 |
| Šídák | Planned comparison | F2S − Tb vs. F2S + Tb, 9.5 mM $Tb^{3+}$ | 0.116 [−0.224, 0.455] | 1.31× | $t_{48} = 0.685$ | 0.4967 | 9 vs. 9 |
| Šídák | Planned comparison | F2S − Tb vs. F2S + Tb, 5.9 mM Ce/Dy/Nd/Tb | 0.0159 [−0.324, 0.355] | 1.04× | $t_{48} = 0.0942$ | 0.9253 | 9 vs. 9 |
| Šídák | Planned comparison | 0.95 mM vs. 9.5 mM $Tb^{3+}$, F2S − Tb | −1.14 [−1.56, −0.720] | 0.072× | $t_{48} = 6.74$ | < 0.0001 | 9 vs. 9 |
| Šídák | Planned comparison | 0.95 mM $Tb^{3+}$ vs. 5.9 mM Ce/Dy/Nd/Tb, F2S − Tb | −1.06 [−1.48, −0.640] | 0.087× | $t_{48} = 6.26$ | < 0.0001 | 9 vs. 9 |
| Šídák | Planned comparison | 9.5 mM $Tb^{3+}$ vs. 5.9 mM Ce/Dy/Nd/Tb, F2S − Tb | 0.0801 [−0.338, 0.498] | 1.20× | $t_{48} = 0.474$ | 0.9524 | 9 vs. 9 |
| Šídák | Planned comparison | 0.95 mM vs. 9.5 mM $Tb^{3+}$, F2S + Tb | −1.18 [−1.60, −0.766] | 0.066× | $t_{48} = 7.01$ | < 0.0001 | 9 vs. 9 |
| Šídák | Planned comparison | 0.95 mM $Tb^{3+}$ vs. 5.9 mM Ce/Dy/Nd/Tb, F2S + Tb | −1.20 [−1.62, −0.786] | 0.063× | $t_{48} = 7.13$ | < 0.0001 | 9 vs. 9 |
| Šídák | Planned comparison | 9.5 mM $Tb^{3+}$ vs. 5.9 mM Ce/Dy/Nd/Tb, F2S + Tb | −0.0197 [−0.437, 0.398] | 0.96× | $t_{48} = 0.117$ | 0.9992 | 9 vs. 9 |

**Table S37. Ordinary two-way ANOVA of lixiviant, tissue condition, and their interaction for ΔpH and ΔORP during solid–liquid extraction of metals from (enriched) *P. acinosa* and *P. americana* shoots.** Biomass effects were calculated from 7 d leaching controls containing $H_2O$, $H_2SO_4$, citric acid, or $Na_2EDTA$. $\Delta pH$ and $\Delta ORP$ were analyzed separately by ordinary two-way ANOVA with lixiviant and tissue condition as fixed effects. The $\Delta pH$ and $\Delta ORP$ analyses included 306 observations across 16 lixiviant–tissue combinations, with 3–48 observations per combination.

| Response | Effect | SS | df | MS | Test statistic | *P* value |
|---|---|---|---|---|---|---|
| $\Delta pH$ | Lixiviant × tissue | 3.08 | 9 | 0.343 | $F_{9,290} = 2.87$ | 0.0030 |
| $\Delta pH$ | Lixiviant | 7.10 | 3 | 2.37 | $F_{3,290} = 19.8$ | $< 0.0001$ |
| $\Delta pH$ | Tissue | 10.4 | 3 | 3.46 | $F_{3,290} = 28.9$ | $< 0.0001$ |
| $\Delta pH$ | Residual | 34.7 | 290 | 0.120 | – | – |
| $\Delta ORP$ | Lixiviant × tissue | 67169 | 9 | 7463 | $F_{9,290} = 9.39$ | $< 0.0001$ |
| $\Delta ORP$ | Lixiviant | 60285 | 3 | 20095 | $F_{3,290} = 25.3$ | $< 0.0001$ |
| $\Delta ORP$ | Tissue | 49231 | 3 | 16410 | $F_{3,290} = 20.6$ | $< 0.0001$ |
| $\Delta ORP$ | Residual | 230552 | 290 | 795 | – | – |

**Table S38. Dunnett-adjusted comparisons of ΔpH following two-way ANOVA between biomass treatments and no-shoot controls within each lixiviant after solid–liquid extraction of metals from (enriched) *P. acinosa* and *P. americana* shoots.** Values are reported as the biomass-containing treatment minus the corresponding no-shoot control within each lixiviant, so positive values indicate that biomass increased the final pH change relative to the matched no-shoot control. Comparisons were performed within each lixiviant using Dunnett's multiple comparisons test against the matched no-shoot control.

| Lixiviant | Tissue condition | Biomass effect on $\Delta pH$ [95% CI] | Test statistic | Adjusted *P* value | n |
|---|---|---|---|---|---|
| $H_2O$ | +25 mM Ni *P. acinosa* | 0.670 [0.0869, 1.25] | $q_{290} = 2.74$ | 0.0188 | 6 vs. 3 |
| $H_2O$ | +50 mM Ni *P. acinosa* | 0.637 [0.0536, 1.22] | $q_{290} = 2.60$ | 0.0278 | 6 vs. 3 |
| $H_2O$ | +Ce/Dy *P. americana* | -0.380 [-0.963, 0.203] | $q_{290} = 1.55$ | 0.3021 | 6 vs. 3 |
| $H_2SO_4$ | +25 mM Ni *P. acinosa* | 0.486 [0.241, 0.730] | $q_{290} = 4.75$ | $< 0.0001$ | 48 vs. 15 |
| $H_2SO_4$ | +50 mM Ni *P. acinosa* | 0.481 [0.237, 0.725] | $q_{290} = 4.70$ | $< 0.0001$ | 48 vs. 15 |
| $H_2SO_4$ | +Ce/Dy *P. americana* | 0.133 [-0.0379, 0.305] | $q_{290} = 1.86$ | 0.1709 | 48 vs. 45 |
| Citric acid | +25 mM Ni *P. acinosa* | 0.540 [0.289, 0.791] | $q_{290} = 5.14$ | $< 0.0001$ | 39 vs. 15 |
| Citric acid | +50 mM Ni *P. acinosa* | 0.509 [0.258, 0.760] | $q_{290} = 4.85$ | $< 0.0001$ | 39 vs. 15 |
| Citric acid | +Ce/Dy *P. americana* | -0.185 [-0.399, 0.0288] | $q_{290} = 2.07$ | 0.1091 | 39 vs. 24 |
| $Na_2EDTA$ | +25 mM Ni *P. acinosa* | 0.219 [-0.0913, 0.530] | $q_{290} = 1.69$ | 0.2418 | 33 vs. 9 |
| $Na_2EDTA$ | +50 mM Ni *P. acinosa* | 0.126 [-0.185, 0.437] | $q_{290} = 0.970$ | 0.6854 | 33 vs. 9 |
| $Na_2EDTA$ | +Ce/Dy *P. americana* | -0.249 [-0.470, -0.0268] | $q_{290} = 2.68$ | 0.0227 | 33 vs. 24 |

**Table S39. Dunnett-adjusted comparisons of ΔORP following two-way ANOVA between biomass treatments and no-shoot controls within each lixiviant after solid–liquid extraction of metals from (enriched) *P. acinosa* and *P. americana* shoots.** Values are reported as the biomass-containing treatment minus the corresponding no-shoot control within each lixiviant, so negative values indicate that biomass lowered ORP relative to the matched no-shoot control. ORP was measured versus an Ag/AgCl reference electrode with 3 M KCl. Comparisons were performed within each lixiviant using Dunnett's multiple comparisons test against the matched no-shoot control.

| Lixiviant | Tissue condition | Biomass effect on $\Delta ORP$ (mV) [95% CI] | Test statistic | Adjusted $P$ value | n |
|---|---|---|---|---|---|
| $H_2O$ | +25 mM Ni *P. acinosa* | -17.0 [-64.6, 30.6] | $q_{290} = 0.853$ | 0.7561 | 6 vs. 3 |
| $H_2O$ | +50 mM Ni *P. acinosa* | 106 [58.4, 154] | $q_{290} = 5.32$ | $< 0.0001$ | 6 vs. 3 |
| $H_2O$ | +Ce/Dy *P. americana* | -3.67 [-51.2, 43.9] | $q_{290} = 0.184$ | 0.9964 | 6 vs. 3 |
| $H_2SO_4$ | +25 mM Ni *P. acinosa* | -76.0 [-95.9, -56.1] | $q_{290} = 9.11$ | $< 0.0001$ | 48 vs. 15 |
| $H_2SO_4$ | +50 mM Ni *P. acinosa* | -77.4 [-97.3, -57.5] | $q_{290} = 9.28$ | $< 0.0001$ | 48 vs. 15 |
| $H_2SO_4$ | +Ce/Dy *P. americana* | -40.6 [-54.6, -26.7] | $q_{290} = 6.94$ | $< 0.0001$ | 48 vs. 45 |
| Citric acid | +25 mM Ni *P. acinosa* | -39.8 [-60.3, -19.3] | $q_{290} = 4.65$ | $< 0.0001$ | 39 vs. 15 |
| Citric acid | +50 mM Ni *P. acinosa* | -52.1 [-72.5, -31.6] | $q_{290} = 6.08$ | $< 0.0001$ | 39 vs. 15 |
| Citric acid | +Ce/Dy *P. americana* | -37.9 [-55.3, -20.4] | $q_{290} = 5.18$ | $< 0.0001$ | 39 vs. 24 |
| $Na_2EDTA$ | +25 mM Ni *P. acinosa* | -59.3 [-84.7, -34.0] | $q_{290} = 5.60$ | $< 0.0001$ | 33 vs. 9 |
| $Na_2EDTA$ | +50 mM Ni *P. acinosa* | -57.8 [-83.1, -32.4] | $q_{290} = 5.45$ | $< 0.0001$ | 33 vs. 9 |
| $Na_2EDTA$ | +Ce/Dy *P. americana* | -25.0 [-43.1, -6.92] | $q_{290} = 3.31$ | 0.0032 | 33 vs. 24 |

**Table S40. Two-tailed one-sample $t$ tests of mass retention against 50% after solid–liquid extraction of metals from (enriched) *P. acinosa* and *P. americana* shoots.** All entries correspond to shoot tissues. Mass retained was compared against the theoretical value of 50% using two-tailed one-sample $t$ tests. Differences are reported as observed mean minus 50%.

| Species | Growth treatment | Leaching condition | $m_{\mathbf{retained}}$ (%) | Difference (%) [95% CI] | Test statistic | $P$ value | n |
|---|---|---|---|---|---|---|---|
| *P. americana* | soil-grown baseline | – | 43.7 | −6.26 [−8.77, −3.75] | $t_2 = 10.7$ | 0.0086 | 3 |
| *P. americana* | hydroponic baseline | – | 50.9 | 0.874 [−4.82, 6.56] | $t_2 = 0.661$ | 0.5767 | 3 |
| *P. americana* | + Ce/Dy | pH 1 | 46.1 | −3.92 [−5.09, −2.76] | $t_2 = 14.5$ | 0.0047 | 3 |
| *P. americana* | + Ce/Dy | +20× | 48.1 | −1.95 [−13.1, 9.21] | $t_2 = 0.751$ | 0.5309 | 3 |
| *P. americana* | + Ce/Dy | +0.1× | 45.7 | −4.34 [−6.68, −2.00] | $t_2 = 7.97$ | 0.0154 | 3 |
| *P. americana* | + Ce/Dy | pH 3 | 50.6 | 0.614 [−1.82, 3.04] | $t_2 = 1.09$ | 0.3907 | 3 |
| *P. americana* | + Ce/Dy | pH 5 | 51.3 | 1.31 [−0.461, 3.08] | $t_2 = 3.18$ | 0.0861 | 3 |
| *P. americana* | + Ce/Dy | $H_2O$ | 50.2 | 0.190 [−0.491, 0.871] | $t_2 = 1.20$ | 0.3536 | 3 |
| *P. acinosa* | +25 mM Ni | pH 1 | 51.6 | 1.61 [−0.468, 3.68] | $t_2 = 3.33$ | 0.0795 | 3 |
| *P. acinosa* | +25 mM Ni | pH 3 | 47.4 | −2.58 [−5.76, 0.599] | $t_2 = 3.49$ | 0.0731 | 3 |
| *P. acinosa* | +25 mM Ni | pH 5 | 48.8 | −1.22 [−3.75, 1.30] | $t_2 = 2.09$ | 0.1722 | 3 |
| *P. acinosa* | +25 mM Ni | $H_2O$ | 47.5 | −2.52 [−2.94, −2.10] | $t_2 = 25.8$ | 0.0015 | 3 |
| *P. acinosa* | +50 mM Ni | pH 1 | 53.9 | 3.87 [2.22, 5.51] | $t_2 = 10.1$ | 0.0097 | 3 |
| *P. acinosa* | +50 mM Ni | pH 3 | 41.0 | −9.00 [−10.5, −7.49] | $t_2 = 25.7$ | 0.0015 | 3 |
| *P. acinosa* | +50 mM Ni | pH 5 | 49.2 | −0.780 [−3.48, 1.92] | $t_2 = 1.25$ | 0.3391 | 3 |
| *P. acinosa* | +50 mM Ni | $H_2O$ | 49.2 | −0.810 [−1.64, 0.0192] | $t_2 = 4.20$ | 0.0522 | 3 |

**Table S41. Two-tailed one-sample $t$ tests of UV absorbance retention against 50% in ultrafiltration fractions after solid–liquid extraction of metals from (enriched) *P. acinosa* and *P. americana* shoots.** All entries correspond to shoot tissues. $AUC_{220-400}$ retained was compared against the theoretical value of 50% using two-tailed one-sample $t$ tests. Differences are reported as observed mean minus 50%.

| Species | Growth treatment | Leaching condition | $AUC_{220\text{-}400,retained}$ (%) | Difference (%) [95% CI] | Test statistic | $P$ value | n |
|---|---|---|---|---|---|---|---|
| *P. americana* | soil-grown baseline | – | 79.1 | 29.1 [22.1, 36.0] | $t_2 = 17.9$ | 0.0031 | 3 |
| *P. americana* | hydroponic baseline | – | 82.4 | 32.4 [11.1, 53.8] | $t_2 = 6.53$ | 0.0227 | 3 |
| *P. americana* | + Ce/Dy | pH 1 | 60.6 | 10.6 [−3.58, 24.8] | $t_2 = 3.22$ | 0.0846 | 3 |
| *P. americana* | + Ce/Dy | +20× | 91.0 | 41.0 [27.5, 54.5] | $t_2 = 13.1$ | 0.0058 | 3 |
| *P. americana* | + Ce/Dy | +0.1× | 111 | 61.3 [−2.81, 125] | $t_2 = 4.11$ | 0.0543 | 3 |
| *P. americana* | + Ce/Dy | pH 3 | 73.4 | 23.4 [2.97, 43.8] | $t_2 = 4.93$ | 0.0388 | 3 |
| *P. americana* | + Ce/Dy | pH 5 | 91.4 | 41.4 [31.4, 51.5] | $t_2 = 17.7$ | 0.0032 | 3 |
| *P. americana* | + Ce/Dy | $H_2O$ | 92.4 | 42.4 [38.9, 46.0] | $t_2 = 51.4$ | 0.0004 | 3 |
| *P. acinosa* | +25 mM Ni | pH 1 | 62.0 | 12.0 [10.0, 14.1] | $t_2 = 25.6$ | 0.0015 | 3 |
| *P. acinosa* | +25 mM Ni | pH 3 | 64.5 | 14.5 [5.18, 23.9] | $t_2 = 6.69$ | 0.0216 | 3 |
| *P. acinosa* | +25 mM Ni | pH 5 | 82.2 | 32.2 [18.8, 45.6] | $t_2 = 10.3$ | 0.0093 | 3 |
| *P. acinosa* | +25 mM Ni | $H_2O$ | 86.3 | 36.3 [30.8, 41.8] | $t_2 = 28.2$ | 0.0013 | 3 |
| *P. acinosa* | +50 mM Ni | pH 1 | 63.2 | 13.2 [7.06, 19.3] | $t_2 = 9.28$ | 0.0114 | 3 |
| *P. acinosa* | +50 mM Ni | pH 3 | 51.9 | 1.90 [1.00, 2.80] | $t_2 = 9.13$ | 0.0118 | 3 |
| *P. acinosa* | +50 mM Ni | pH 5 | 79.3 | 29.3 [22.4, 36.1] | $t_2 = 18.4$ | 0.0029 | 3 |
| *P. acinosa* | +50 mM Ni | $H_2O$ | 84.2 | 34.2 [33.1, 35.3] | $t_2 = 129$ | $< 0.0001$ | 3 |

**Table S42. Two-tailed one-sample $t$ tests of Ni or REE retention against 50% after solid–liquid extraction of metals from (enriched) *P. acinosa* and *P. americana* shoots.** All entries correspond to shoot tissues. Target-metal retention of analyte(s) $i$ was compared against the theoretical value of 50% using two-tailed one-sample $t$ tests. Differences are reported as observed mean minus 50%.

| Species | Growth treatment | Leaching condition | $i_{retention}$ (%) | Difference (%)[95% CI] | Test statistic | $P$ value | n |
|---|---|---|---|---|---|---|---|
| *P. americana* | + Ce/Dy | pH 1 | 51.0 | 0.997 [–2.56, 4.55] | $t_2 = 1.21$ | 0.3512 | 3 |
| *P. americana* | + Ce/Dy | +20× | 82.9 | 32.9 [–0.287, 66.1] | $t_2 = 4.27$ | 0.0508 | 3 |
| *P. americana* | + Ce/Dy | +0.1× | 53.8 | 3.85 [–25.2, 32.9] | $t_2 = 0.571$ | 0.6259 | 3 |
| *P. americana* | + Ce/Dy | pH 3 | 57.8 | 7.76 [5.75, 9.76] | $t_2 = 16.7$ | 0.0036 | 3 |
| *P. americana* | + Ce/Dy | pH 5 | 95.9 | 45.9 [41.3, 50.5] | $t_2 = 43.1$ | 0.0005 | 3 |
| *P. americana* | + Ce/Dy | $H_2O$ | 97.9 | 47.9 [46.2, 49.5] | $t_2 = 122$ | $< 0.0001$ | 3 |
| *P. acinosa* | +25 mM Ni | pH 1 | 56.8 | 6.80 [3.39, 10.2] | $t_2 = 8.56$ | 0.0134 | 3 |
| *P. acinosa* | +25 mM Ni | pH 3 | 50.7 | 0.697 [–3.87, 5.26] | $t_2 = 0.657$ | 0.5788 | 3 |
| *P. acinosa* | +25 mM Ni | pH 5 | 55.8 | 5.80 [–1.65, 13.2] | $t_2 = 3.35$ | 0.0787 | 3 |
| *P. acinosa* | +25 mM Ni | $H_2O$ | 55.2 | 5.23 [2.39, 8.07] | $t_2 = 7.94$ | 0.0155 | 3 |
| *P. acinosa* | +50 mM Ni | pH 1 | 57.5 | 7.52 [4.05, 11.0] | $t_2 = 9.32$ | 0.0113 | 3 |
| *P. acinosa* | +50 mM Ni | pH 3 | 43.0 | –6.95 [–7.90, –6.01] | $t_2 = 31.7$ | 0.0010 | 3 |
| *P. acinosa* | +50 mM Ni | pH 5 | 52.5 | 2.50 [–3.36, 8.35] | $t_2 = 1.83$ | 0.2080 | 3 |
| *P. acinosa* | +50 mM Ni | $H_2O$ | 53.8 | 3.84 [1.45, 6.23] | $t_2 = 6.92$ | 0.0203 | 3 |

**Table S43. Ordinary least-squares regression models relating Ni retention during 3 kDa ultrafiltration of *P. acinosa* leachates to mass retention, UV absorbance retention, and pH.** Ordinary least-squares models were fit using Ni retention as the dependent variable. Estimates are reported with 95% confidence intervals. AUC denotes integrated UV absorbance used as a proxy for plant-derived solution components. VIF values are reported for predictor terms only. Residual normality was marked as passed only when all reported residual normality tests passed at $\alpha = 0.05$.

| **Model** | **Term** | **Estimate** [95% CI] | **Term** $P$ | **VIF** | **Model statistic** | **Model** $P$ | $R^2$ | **Residual normality** | $n$ |
|---|---|---|---|---|---|---|---|---|---|
| pH only | Intercept | 53.2 [48.5, 57.9] | $< 0.0001$ | – | $F_{1,22} = 4.49 \times 10^{-6}$ | 0.9983 | $2.04 \times 10^{-7}$ | No | 24 |
| | pH | -0.00113 [-1.10, 1.10] | 0.9983 | 1.00 | | | | | |
| AUC retention only | Intercept | 40.0 [29.4, 50.5] | $< 0.0001$ | – | $F_{1,22} = 6.91$ | 0.0153 | 0.239 | Yes | 24 |
| | AUC retained (%) | 0.184 [0.0389, 0.330] | 0.0153 | 1.00 | | | | | |
| Mass retention only | Intercept | -2.76 [-15.3, 9.75] | 0.6514 | – | $F_{1,22} = 86.4$ | $< 0.0001$ | 0.797 | Yes | 24 |
| | Mass retained (%) | 1.15 [0.895, 1.41] | $< 0.0001$ | 1.00 | | | | | |
| Mass + AUC retention | Intercept | -6.08 [-16.4, 4.22] | 0.2329 | – | $F_{2,21} = 72.9$ | $< 0.0001$ | 0.874 | Yes | 24 |
| | Mass retained (%) | 1.06 [0.846, 1.28] | $< 0.0001$ | 1.06 | | | | | |
| | AUC retained (%) | 0.108 [0.0453, 0.171] | 0.0017 | 1.06 | | | | | |
| Mass retention + pH | Intercept | -8.76 [-20.7, 3.20] | 0.1426 | – | $F_{2,21} = 59.5$ | $< 0.0001$ | 0.850 | Yes | 24 |
| | Mass retained (%) | 1.23 [0.994, 1.46] | $< 0.0001$ | 1.07 | | | | | |
| | pH | 0.593 [0.140, 1.05] | 0.0128 | 1.07 | | | | | |
| AUC retention + pH | Intercept | 24.6 [16.3, 32.9] | $< 0.0001$ | – | $F_{2,21} = 28.2$ | $< 0.0001$ | 0.729 | No | 24 |
| | AUC retained (%) | 0.561 [0.406, 0.717] | $< 0.0001$ | 3.04 | | | | | |
| | pH | -3.05 [-4.08, -2.02] | $< 0.0001$ | 3.04 | | | | | |
| Mass + AUC retention + pH | Intercept | -2.26 [-14.6, 10.1] | 0.7062 | – | $F_{3,20} = 49.8$ | $< 0.0001$ | 0.882 | Yes | 24 |
| | Mass retained (%) | 0.896 [0.529, 1.26] | $< 0.0001$ | 3.15 | | | | | |
| | AUC retained (%) | 0.202 [0.0210, 0.383] | 0.0305 | 8.98 | | | | | |
| | pH | -0.664 [-1.86, 0.536] | 0.2619 | 8.99 | | | | | |

**Table S44. Ordinary least-squares regression models relating REE retention during 3 kDa ultrafiltration of *P. americana* leachates to mass retention, UV absorbance retention, and pH.** Ordinary least-squares models were fit using REE retention as the dependent variable. Estimates are reported with 95% confidence intervals. AUC denotes integrated UV absorbance used as a proxy for plant-derived solution components. VIF values are reported for predictor terms only. Residual normality was marked as passed only when all reported residual normality tests passed at $\alpha = 0.05$.

| Model | Term | Estimate [95% CI] | Term $P$ value | VIF | Model statistic | Model $P$ value | $R^2$ | Residual normality | $n$ |
|---|---|---|---|---|---|---|---|---|---|
| pH only | Intercept | 36.2 [21.5, 51.0] | $< 0.0001$ | – | $F_{1,16} = 34.2$ | $< 0.0001$ | 0.681 | Yes | 18 |
| | pH | 10.6 [6.74, 14.4] | $< 0.0001$ | 1.00 | | | | | |
| AUC retention only | Intercept | 52.6 [1.85, 103] | 0.0431 | – | $F_{1,16} = 0.780$ | 0.3903 | 0.0465 | No | 18 |
| | AUC retention (%) | 0.238 [−0.334, 0.810] | 0.3903 | 1.00 | | | | | |
| Mass retention only | Intercept | −83.3 [−261, 94.4] | 0.3352 | – | $F_{1,16} = 3.50$ | 0.0798 | 0.179 | No | 18 |
| | Mass retention (%) | 3.22 [−0.429, 6.86] | 0.0798 | 1.00 | | | | | |
| Mass + AUC retention | Intercept | −107 [−293, 78.4] | 0.2369 | – | $F_{2,15} = 2.25$ | 0.1394 | 0.231 | Yes | 18 |
| | Mass retention (%) | 3.26 [−0.402, 6.93] | 0.0772 | 1.00 | | | | | |
| | AUC retention (%) | 0.251 [−0.282, 0.785] | 0.3314 | 1.00 | | | | | |
| Mass retention + pH | Intercept | 62.7 [−67.9, 193] | 0.3226 | – | $F_{2,15} = 16.3$ | 0.0002 | 0.685 | Yes | 18 |
| | Mass retention (%) | −0.584 [−3.45, 2.28] | 0.6703 | 1.49 | | | | | |
| | pH | 11.1 [6.30, 16.0] | 0.0002 | 1.49 | | | | | |
| AUC retention + pH | Intercept | 50.1 [21.0, 79.3] | 0.0023 | – | $F_{2,15} = 18.2$ | $< 0.0001$ | 0.708 | Yes | 18 |
| | AUC retention (%) | −0.202 [−0.567, 0.164] | 0.2580 | 1.24 | | | | | |
| | pH | 11.6 [7.36, 15.8] | $< 0.0001$ | 1.24 | | | | | |
| Mass + AUC retention + pH | Intercept | 117 [−33.0, 266] | 0.1169 | – | $F_{3,14} = 12.4$ | 0.0003 | 0.727 | Yes | 18 |
| | Mass retention (%) | −1.36 [−4.37, 1.65] | 0.3478 | 1.75 | | | | | |
| | AUC retention (%) | −0.271 [−0.669, 0.128] | 0.1676 | 1.45 | | | | | |
| | pH | 13.3 [7.62, 18.9] | 0.0002 | 2.17 | | | | | |

**Table S45. Ordinary least-squares multiple regression of $\log_{10}$-transformed inferred REE-binding capacity by *A. ferrooxidans* across most binding tests.** Endpoint binding data were analyzed by ordinary least-squares regression using $\log_{10}$-transformed DCW-normalized REE binding capacity as the dependent variable. Solution class, $\log_{10}$-transformed initial REE concentration, $\log_{10}$-transformed OD600, and the $\log 10(\text{TREE}) \times \log_{10}(\text{OD}_{600})$ interaction were included as predictors. $Tb^{3+}$-only binding tests were used as the reference class. Parameter estimates are reported with 95% confidence intervals. Residual normality was marked as passed only when all reported residual normality tests passed at $\alpha = 0.05$. The data from pretreated *A. ferrooxidans* were excluded from this analysis because these tests were designed to alter the REE-binding capacity.

| Model term | Estimate [95% CI] | Term SS | DF | Test statistic | $P$ value | VIF | Notes |
|---|---|---|---|---|---|---|---|
| Model | – | 341 | 5 | $F_{5,203} = 1.92 \times 10^6$ | $< 0.0001$ | – | $R^2 = 1.000$; $n = 209$ |
| Class | – | 0.025 | 2 | $F_{2,203} = 358$ | $< 0.0001$ | – | Reference: $Tb^{3+}$ only |
| mixed REE simulant | 0.029 [0.027, 0.031] | – | – | – | $< 0.0001$ | 1.7 | Relative to $Tb^{3+}$ only |
| plant leachate | 0.032 [0.028, 0.036] | – | – | – | $< 0.0001$ | 2.8 | Relative to $Tb^{3+}$ only |
| $\log_{10}(\text{REE}_0)$ | 1.00 [1.00, 1.00] | 82.0 | 1 | $F_{1,203} = 2.31 \times 10^6$ | $< 0.0001$ | 4.2 | Initial REE concentration |
| $\log_{10}(\text{OD}_{600})$ | −0.0080 [−0.015, −0.00048] | 0.00016 | 1 | $F_{1,203} = 4.4$ | 0.0371 | 17 | Small negative association |
| $\log_{10}(\text{REE}_0) \times \log_{10}(\text{OD}_{600})$ | 0.0022 [0.00020, 0.0041] | 0.00017 | 1 | $F_{1,203} = 4.7$ | 0.0307 | 21 | Small interaction term |
| Residual | – | 0.0072 | 203 | – | – | – | Residual normality: No |

**Table S46. Ordinary least-squares regression models of endpoint REE solubilization in *P. americana* bioleaching experiments to some experimental factors and measurements.** Ordinary least-squares models were fit using endpoint REE solubilization as the dependent variable. Endpoint pH, ORP, and conductivity were centered prior to regression and are denoted with subscript $c$. For treatment-identity models, treatment identity was included as a categorical predictor and the abiotic treatment using an initial pH of 5.0, 1 g/L brown $S^0$, and 1% (w/v) *P. americana* was used as the reference class. Estimates are reported with 95% confidence intervals. VIF values are reported for predictor terms only; for treatment identity, the maximum VIF among the treatment-indicator variables is shown. Residual normality was marked as passed only when all reported residual normality tests passed at $\alpha = 0.05$.

| Model | Term | Estimate [95% CI] | Term $P$ value | VIF | Model statistic | Model $P$ value | $R^2$ | Residual normality | $n$ |
|---|---|---|---|---|---|---|---|---|---|
| pH only | Intercept | 35.7 [30.6, 40.7] | < 0.0001 | – | $F_{1,40} = 41.5$ | < 0.0001 | 0.509 | Yes | 42 |
| | $pH_c$ | −7.93 [−10.4, −5.44] | < 0.0001 | 1.00 | | | | | |
| Treatment + pH | Intercept | 30.5 [2.88, 58.1] | 0.0317 | – | $F_{14,27} = 105$ | < 0.0001 | 0.982 | Yes | 42 |
| | Treatment identity | – | < 0.0001 | 82.5 | | | | | |
| | $pH_c$ | −5.89 [−13.5, 1.74] | 0.1248 | 168 | | | | | |
| Inoculum density + pH | Intercept | 39.3 [33.6, 45.1] | < 0.0001 | – | $F_{2,39} = 26.0$ | < 0.0001 | 0.571 | Yes | 42 |
| | Inoculum density | −151 [−280, −22.1] | 0.0228 | 1.11 | | | | | |
| | $pH_c$ | −8.82 [−11.3, −6.35] | < 0.0001 | 1.11 | | | | | |
| Sulfur dosage + pH | Intercept | 34.7 [27.4, 41.9] | < 0.0001 | – | $F_{2,39} = 20.4$ | < 0.0001 | 0.511 | Yes | 42 |
| | Sulfur dosage | 0.891 [−3.82, 5.60] | 0.7039 | 1.20 | | | | | |
| | $pH_c$ | −7.72 [−10.5, −4.97] | < 0.0001 | 1.20 | | | | | |
| Plant dosage + pH | Intercept | 34.6 [28.0, 41.2] | < 0.0001 | – | $F_{2,39} = 20.5$ | < 0.0001 | 0.513 | Yes | 42 |
| | log(plant dosage) | −2.08 [−9.86, 5.70] | 0.5921 | 1.23 | | | | | |
| | $pH_c$ | −7.60 [−10.4, −4.81] | < 0.0001 | 1.23 | | | | | |
| Treatment + pH + ORP | Intercept | 30.7 [2.35, 59.1] | 0.0349 | – | $F_{15,26} = 94.7$ | < 0.0001 | 0.982 | Yes | 42 |
| | Treatment identity | – | < 0.0001 | 83.6 | | | | | |
| | $pH_c$ | −5.88 [−13.7, 1.91] | 0.1327 | 168 | | | | | |
| | $ORP_c$ | 0.00313 [−0.0429, 0.0491] | 0.8897 | 10.8 | | | | | |
| Treatment + pH + conductivity | Intercept | 37.5 [−0.470, 75.4] | 0.0527 | – | $F_{15,26} = 95.8$ | < 0.0001 | 0.982 | Yes | 42 |
| | Treatment identity | – | < 0.0001 | 130 | | | | | |
| | $pH_c$ | −7.92 [−18.7, 2.83] | 0.1418 | 324 | | | | | |
| | $Conductivity_c$ | −0.803 [−3.75, 2.14] | 0.5799 | 21.1 | | | | | |

**Table S47. REML mixed-effects analysis of time, treatment, and their interaction for *P. americana* shoot bioleaching trajectories of REE solubilization, pH, ORP, and conductivity.** Fourteen treatments, including four abiotic control conditions, were analyzed with three independent replicate flasks per treatment ($n = 42$). Models included categorical time, treatment identity, and their interaction as fixed effects, with a random intercept for replicate flask, and were fit by restricted maximum likelihood. The Geisser–Greenhouse correction was applied to time and time × treatment tests. Common sampling times were used across treatments. Matching $P$ values were calculated by comparing models with and without the flask random intercept using a restricted likelihood-ratio test referenced to a $\chi^2$ distribution with one degree of freedom. SDs are followed by variances in brackets.

| Response | Fixed effect | Test statistic | $P$ value | G–G $\epsilon$ | Replicate SD [variance] | Residual SD [variance] | Matching $P$ | Notes |
|---|---|---|---|---|---|---|---|---|
| REE solubilization | Time | $F_{3.64,102} = 433$ | $< 0.0001$ | 0.455 | 0.993 [0.986] | 3.64 [13.3] | 0.0534 | 9 time points; $n = 42$ |
| | Treatment identity | $F_{13,28} = 372$ | $< 0.0001$ | – | | | | |
| | Time × treatment identity | $F_{47.3,102} = 21.2$ | $< 0.0001$ | 0.455 | | | | |
| pH | Time | $F_{1.62,45.3} = 132$ | $< 0.0001$ | 0.202 | 0.181 [0.0326] | 0.231 [0.0532] | $< 0.0001$ | 9 time points; $n = 42$ |
| | Treatment identity | $F_{13,28} = 173$ | $< 0.0001$ | – | | | | |
| | Time × treatment identity | $F_{21.0,45.3} = 39.3$ | $< 0.0001$ | 0.202 | | | | |
| ORP | Time | $F_{4.26,119} = 69.7$ | $< 0.0001$ | 0.533 | 20.3 [412] | 20.5 [420] | $< 0.0001$ | 9 time points; $n = 42$ |
| | Treatment identity | $F_{13,28} = 29.5$ | $< 0.0001$ | – | | | | |
| | Time × treatment identity | $F_{55.4,119} = 15.8$ | $< 0.0001$ | 0.533 | | | | |
| Conductivity | Time | $F_{3.63,102} = 125$ | $< 0.0001$ | 0.518 | 0.275 [0.0755] | 0.461 [0.212] | $< 0.0001$ | 8 time points; $n = 42$ |
| | Treatment identity | $F_{13,28} = 35.0$ | $< 0.0001$ | – | | | | |
| | Time × treatment identity | $F_{47.2,102} = 13.5$ | $< 0.0001$ | 0.518 | | | | |

**Table S48. REML mixed-effects analysis of time, inoculum group, and their interaction for REE solubilization, pH, ORP, and conductivity for the *P. americana* shoot bioleaching experiments.** Time-course data from the matched inoculum-density series were analyzed using mixed-effects models fit by restricted maximum likelihood with matched values stacked into replicate-flask subcolumns. The subset included four inoculum groups measured at fixed sulfur dosage and plant dosage. Time, inoculum group, and the time × inoculum group interaction were included as fixed effects. The Geisser–Greenhouse correction was used for time and time × inoculum group effects. Replicate flask was modeled as the matched subject. Matching was marked as effective when the corresponding Prism matching test was significant at $\alpha = 0.05$.

| Response | Fixed effect | Test statistic | $P$ value | G–G $\epsilon$ | Replicate SD [variance] | Residual SD [variance] | Matching $P$ | Notes |
|---|---|---|---|---|---|---|---|---|
| REE solubilization | Time | $F_{1.33,10.6} = 74.2$ | $< 0.0001$ | 0.166 | 2.26 [5.11] | 2.96 [8.74] | $< 0.0001$ | 9 time points; $n = 12$ |
| | Inoculum group | $F_{3,8} = 7.10$ | 0.0121 | – | | | | |
| | Time × inoculum group | $F_{3.99,10.6} = 7.77$ | 0.0035 | 0.166 | | | | |
| pH | Time | $F_{1.69,13.5} = 48.6$ | $< 0.0001$ | 0.211 | 0.185 [0.0342] | 0.360 [0.130] | 0.0076 | 9 time points; $n = 12$ |
| | Inoculum group | $F_{3,8} = 62.3$ | $< 0.0001$ | – | | | | |
| | Time × inoculum group | $F_{5.07,13.5} = 8.09$ | 0.0010 | 0.211 | | | | |
| ORP | Time | $F_{2.59,20.7} = 159$ | $< 0.0001$ | 0.324 | 12.8 [163] | 11.8 [140] | $< 0.0001$ | 9 time points; $n = 12$ |
| | Inoculum group | $F_{3,8} = 25.6$ | 0.0002 | – | | | | |
| | Time × inoculum group | $F_{7.77,20.7} = 12.6$ | $< 0.0001$ | 0.324 | | | | |
| Conductivity | Time | $F_{3.40,27.2} = 98.9$ | $< 0.0001$ | 0.486 | 0.166 [0.0276] | 0.257 [0.0660] | 0.0011 | 8 time points; $n = 12$ |
| | Inoculum group | $F_{3,8} = 137$ | $< 0.0001$ | – | | | | |
| | Time × inoculum group | $F_{10.2,27.2} = 18.6$ | $< 0.0001$ | 0.486 | | | | |

**Table S49. Ordinary least-squares regression of replicate-level bioleaching time-course summary metrics by inoculum group and pH metrics for the *P. americana* shoot bioleaching experiments.** Each biological replicate flask was summarized using time-integrated REE release, time-integrated pH decrease, REE-versus-pH slope, maximum REE solubilization, and minimum pH. Ordinary least-squares regression was performed on the matched S1/P1 inoculum-density series using abiotic controls as the reference inoculum group. Inoculum group was modeled as a categorical predictor. Parameter estimates are reported with 95% confidence intervals. VIF values are reported for predictor terms only. The model $R^2$ is shown as reported by Prism; residual normality was marked as passed only when all reported residual normality tests passed at $\alpha = 0.05$.

| Model term | Estimate [95% CI] | Term SS | DF | Test statistic | $P$ value | VIF | Notes |
|---|---|---|---|---|---|---|---|
| REE AUC vs pH-decrease AUC | – | 2989 | 1 | $F_{1,10} = 2.13$ | 0.1753 | – | $R^2 = 0.175$; $n = 12$ |
| Intercept | 55.0 [−52.3, 162] | – | – | – | 0.2799 | – | – |
| pH-decrease AUC | 1.92 [−1.01, 4.86] | 2989 | 1 | $F_{1,10} = 2.13$ | 0.1753 | 1.00 | Not significant |
| Residual | – | 14046 | 10 | – | – | – | Residual normality: Yes |
| REE AUC vs inoculum group + pH-decrease AUC | – | 16126 | 4 | $F_{4,7} = 31.0$ | 0.0002 | – | $R^2 = 0.947$; $n = 12$ |
| Intercept | 557 [374, 740] | – | – | – | 0.0002 | – | Reference: abiotic inoculum group |
| Inoculum group | – | 13137 | 3 | $F_{3,7} = 33.7$ | 0.0002 | – | Categorical predictor |
| 0.001 | −43.0 [−66.8, −19.2] | – | – | – | 0.0037 | 1.76 | Relative to abiotic reference |
| 0.01 | −4.97 [−27.1, 17.1] | – | – | – | 0.6108 | 1.51 | Relative to abiotic reference |
| 0.1 | −253 [−338, −168] | – | – | – | 0.0002 | 22.4 | Relative to abiotic reference |
| pH-decrease AUC | −10.1 [−14.7, −5.43] | 3426 | 1 | $F_{1,7} = 26.4$ | 0.0013 | 23.9 | Significant; high collinearity |
| Residual | – | 909 | 7 | – | – | – | Residual normality: Yes |
| REE-versus-pH slope vs inoculum group | – | 93.0 | 3 | $F_{3,8} = 4.96$ | 0.0312 | – | $R^2 = 0.650$; $n = 12$ |
| Intercept | −9.37 [−12.7, −6.04] | – | – | – | 0.0002 | – | Reference: abiotic inoculum group |
| Inoculum group | – | 93.0 | 3 | $F_{3,8} = 4.96$ | 0.0312 | – | Categorical predictor |
| 0.001 | 4.47 [−0.233, 9.18] | – | – | – | 0.0598 | 1.50 | Relative to abiotic reference |
| 0.01 | −0.947 [−5.65, 3.76] | – | – | – | 0.6551 | 1.50 | Relative to abiotic reference |
| 0.1 | 5.53 [0.827, 10.2] | – | – | – | 0.0266 | 1.50 | Relative to abiotic reference |
| Residual | – | 50.0 | 8 | – | – | – | Residual normality: No |
| Maximum REE solubilization vs minimum pH | – | 212 | 1 | $F_{1,10} = 3.46$ | 0.0925 | – | $R^2 = 0.257$; $n = 12$ |
| Intercept | −0.125 [−30.9, 30.6] | – | – | – | 0.9929 | – | – |
| Minimum pH | 8.35 [−1.65, 18.4] | 212 | 1 | $F_{1,10} = 3.46$ | 0.0925 | 1.00 | Not significant |
| Residual | – | 614 | 10 | – | – | – | Residual normality: Yes |
| Maximum REE solubilization vs inoculum group + minimum pH | – | 754 | 4 | $F_{4,7} = 18.2$ | 0.0008 | – | $R^2 = 0.912$; $n = 12$ |
| Intercept | 64.0 [7.69, 120] | – | – | – | 0.0312 | – | Reference: abiotic inoculum group |
| Inoculum group | – | 541 | 3 | $F_{3,7} = 17.4$ | 0.0013 | – | Categorical predictor |
| 0.001 | −13.0 [−19.7, −6.35] | – | – | – | 0.0024 | 1.73 | Relative to abiotic reference |
| 0.01 | 0.603 [−5.73, 6.94] | – | – | – | 0.8282 | 1.56 | Relative to abiotic reference |
| 0.1 | −26.9 [−46.1, −7.66] | – | – | – | 0.0130 | 14.4 | Relative to abiotic reference |
| Minimum pH | −9.57 [−26.9, 7.76] | 17.7 | 1 | $F_{1,7} = 1.71$ | 0.2328 | 15.8 | Not significant; high collinearity |
| Residual | – | 72.5 | 7 | – | – | – | Residual normality: Yes |

### Supplementary figures

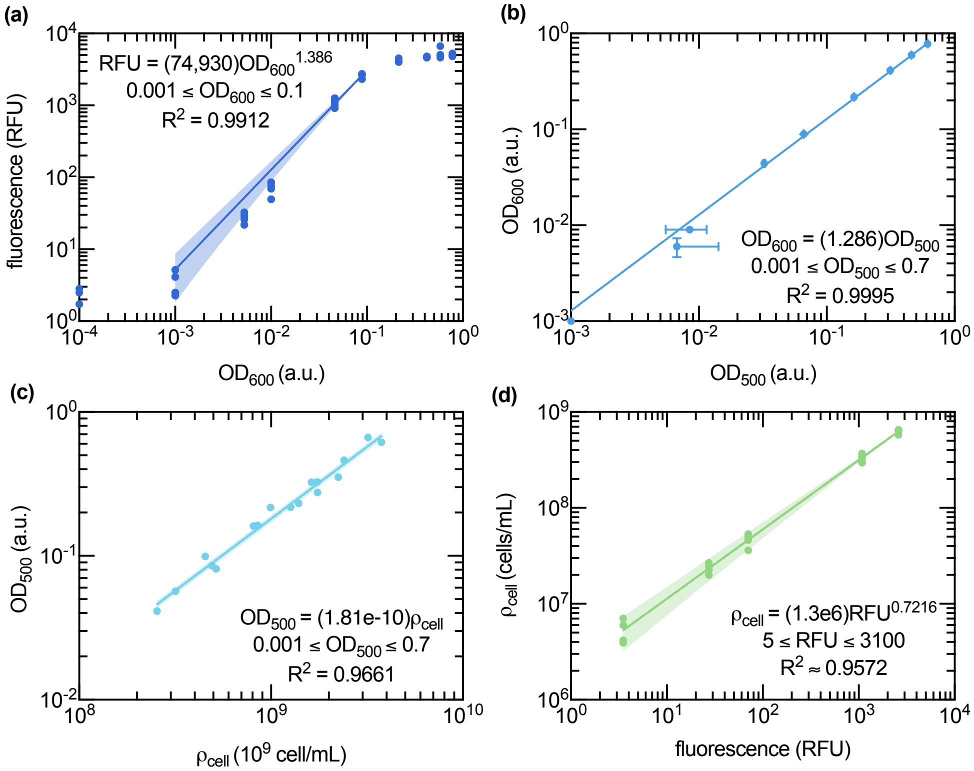


**Figure S1. Calibration of SYBR Green fluorescence and optical density for estimating planktonic *A. ferrooxidans* cell density.** (a) Background-corrected SYBR Green fluorescence increased with $OD_{600}$ according to $RFU = 74{,}930 \cdot OD_{600}^{1.386}$ over the non-saturating calibration range of $0.001 \leq OD_{600} \leq 0.1$. (b) $OD_{600}$ and $OD_{500}$ were linearly related over $0.001 \leq OD_{500} \leq 0.7$, allowing $OD_{500}$-based measurements to be converted to $OD_{600}$ using $OD_{600} = 1.286 \cdot OD_{500}$. (c) Planktonic cell density ($\rho_{cell}$) was linearly related to $OD_{500}$ by $OD_{500} = 1.81 \cdot 10^{-10} \cdot \rho_{cell}$ based on data reported in another study.[3] (d) Combining these calibrations yielded the working conversion from background-corrected fluorescence to planktonic cell density, $\rho_{cell} = 1.3 \cdot 10^6 \cdot RFU^{0.7216}$, over $5 \leq RFU \leq 3100$. Points show the mean of technical triplicates for independent calibration measurements (n = 5), solid lines show fitted relationships, and shaded regions show fitted asymmetric 95% confidence intervals.

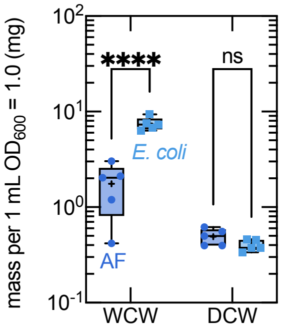


**Figure S2. Wet and dry cell mass of *A. ferrooxidans* and *E. coli* suspensions normalized to $OD_{600} = 1.0$.** Washed cell suspensions were normalized to 1 mL at $OD_{600} = 1.0$ before wet cell weight (WCW) and dry cell weight (DCW) were measured. The measured value of 0.49 mg $mL^{-1}$ $OD_{600}^{-1}$ for *A. ferrooxidans* was close to a previous estimate of 0.53 mg DCW for 1 mL of cell sample at $OD_{600} = 1.0$ from the Banta lab,[2] and the measured value of 0.40 mg $mL^{-1}$ $OD_{600}^{-1}$ for *E. coli* is consistent with reports from other studies.[62] *A. ferrooxidans* had significantly lower WCW than *E. coli* (mean difference = -5.71 mg, 95% CI [-6.75, -4.68], $P_{adj} < 0.0002$), whereas DCW was not significantly different between organisms (mean difference = 0.0934 mg, 95% CI [-0.941, 1.13], $P_{adj} = 0.8505$). Points show independent measurements (n = 5), and boxes show replicate distributions. Organism differences within WCW and DCW were tested using contrasts from the two-way ANOVA, with Holm–Bonferroni correction across the two comparisons. Reported $P$ values are multiplicity adjusted; the reported 95% confidence intervals are pointwise and unadjusted.

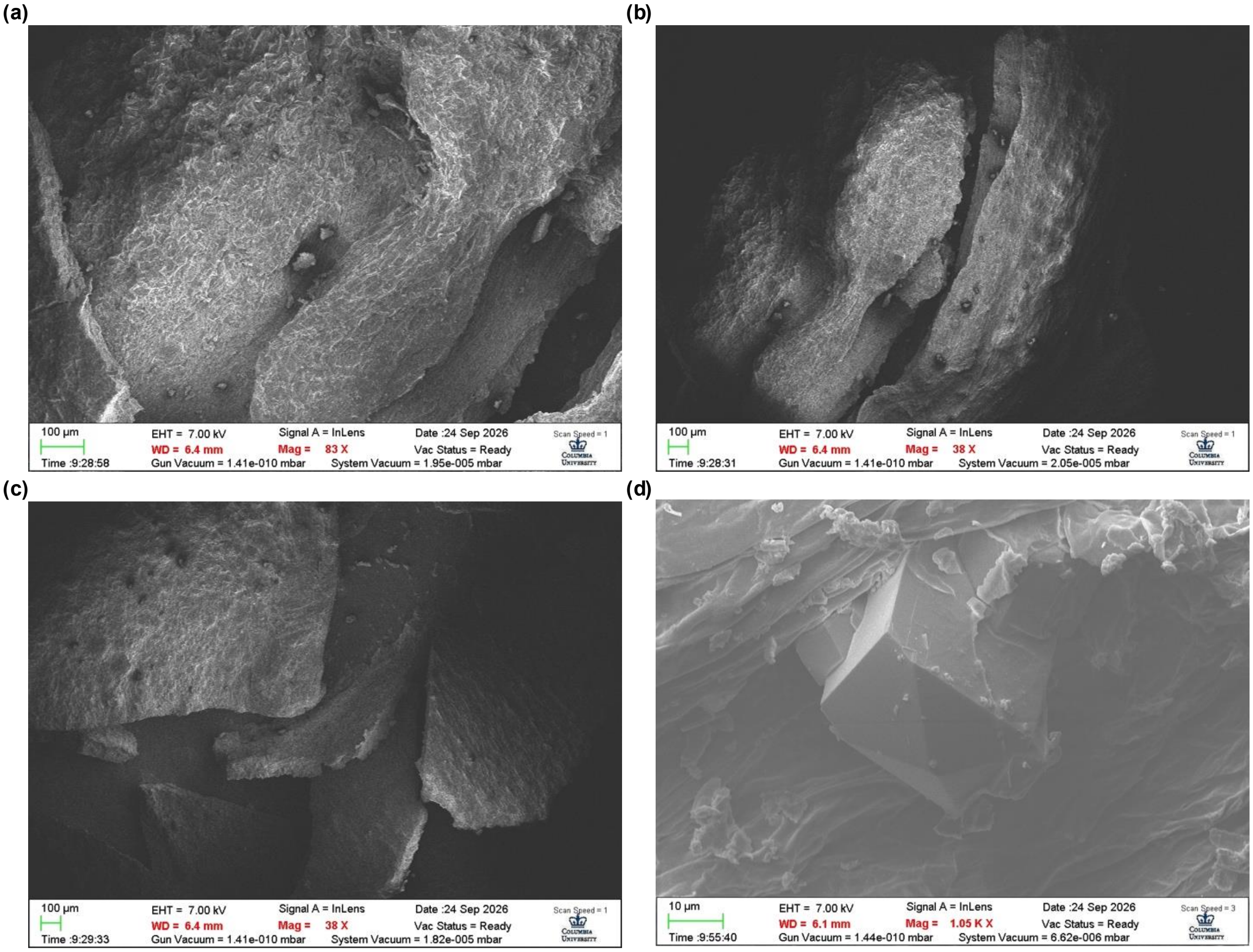


**Figure S3. SEM images of shoot tissue morphology in hydroponically grown, Ce/Dy-supplemented *P. americana* shoots.** (a–c) Low- and (d) higher-magnification views of shoot tissue fragments mounted on carbon tape. Selected images from one representative sample are shown here.

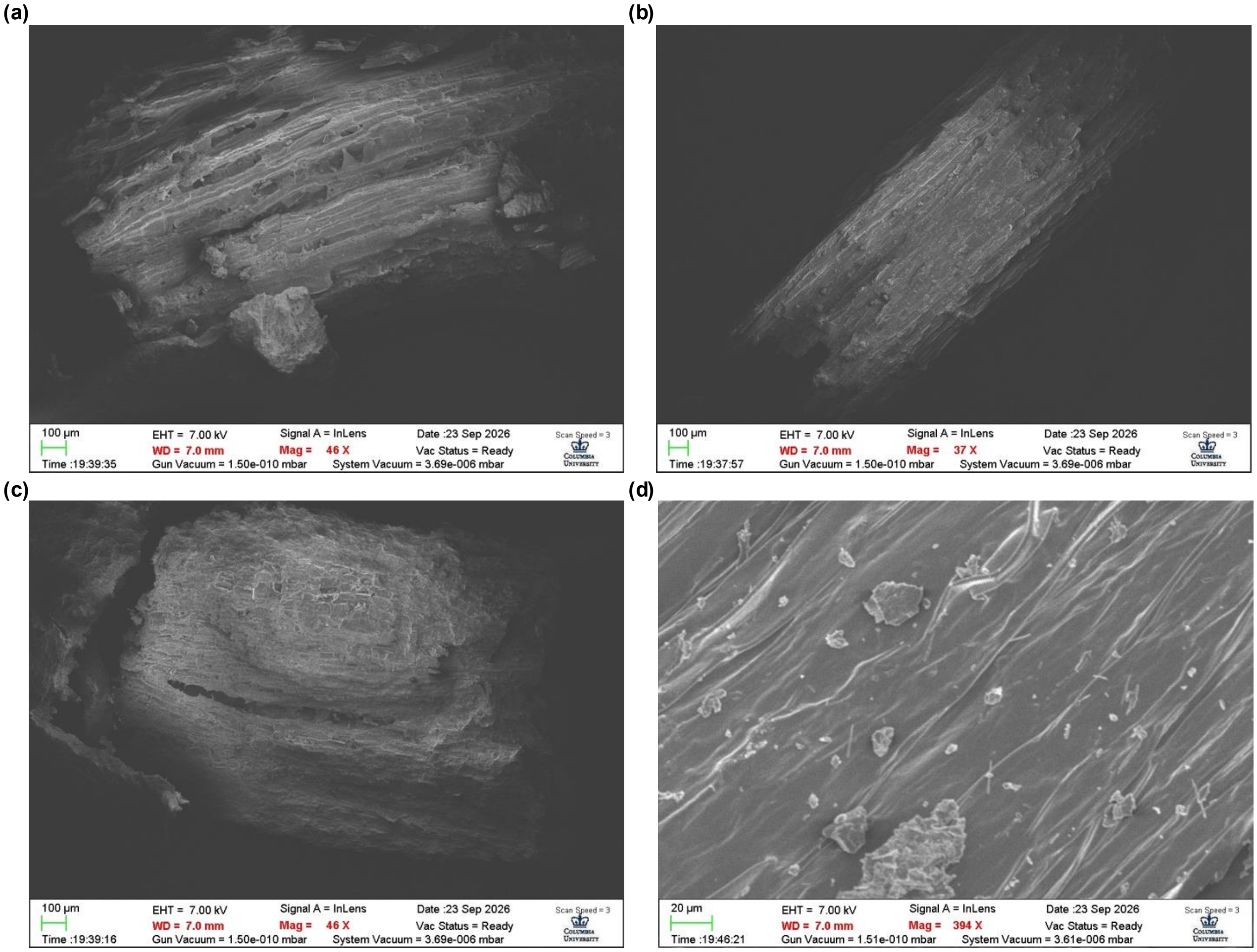


**Figure S4. SEM images of shoot tissue morphology in hydroponically grown control *P. americana* shoots.** (a–c) Low- and (d) higher-magnification views of shoot tissue fragments mounted on carbon tape. Selected images from one representative sample are shown here.

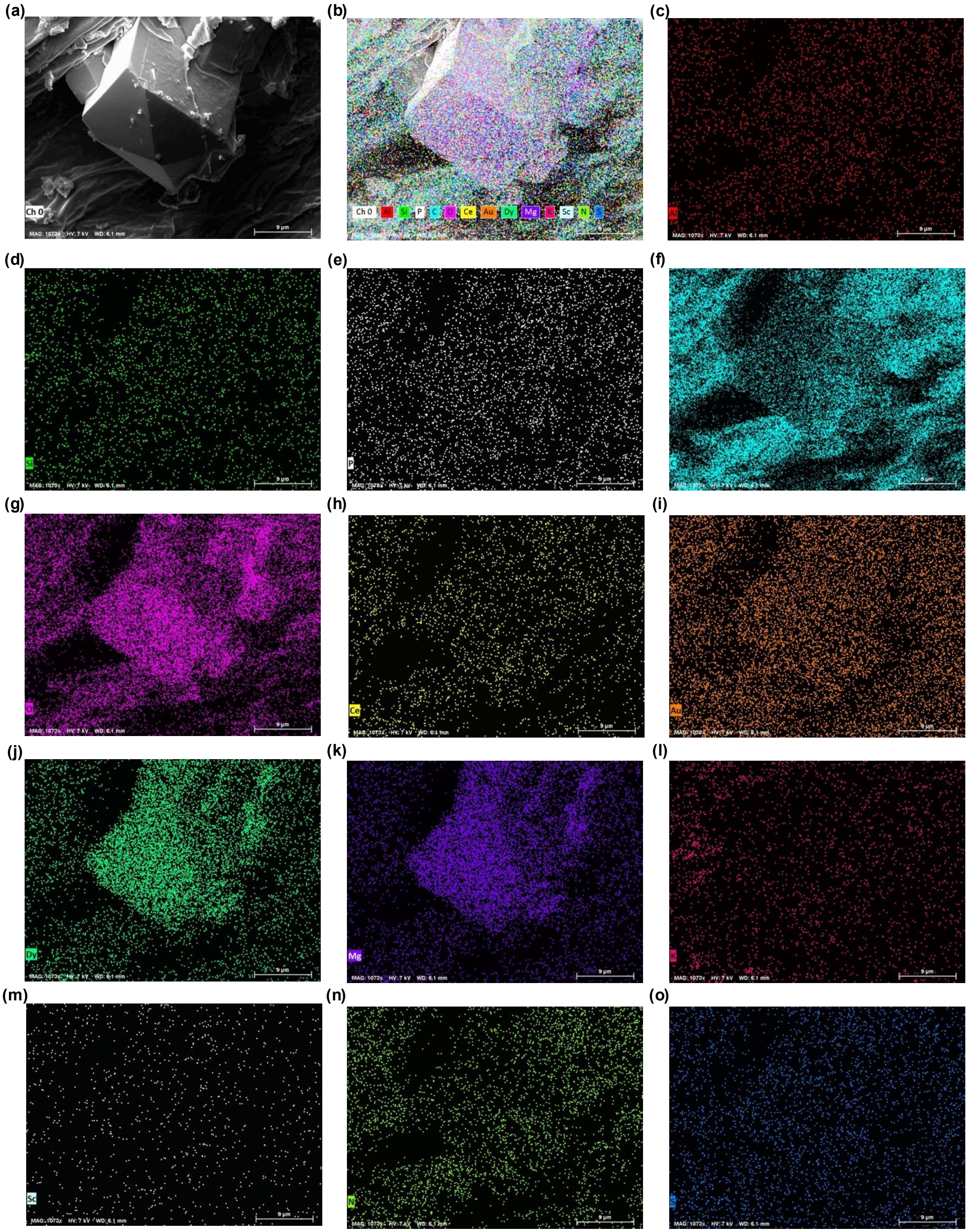


**Figure S5. SEM imaging and EDS mapping of a faceted structure associated with Ce/Dy-supplemented *P. americana* shoot tissue.** (a) Electron image of a faceted structure associated with a shoot tissue fragment and (b) composite overlay of the corresponding EDS maps for dried biomass mounted on carbon tape and sputter-coated (Text S12). Individual maps show software-assigned X-ray channels assigned to (c) Al, (d) silicon, (e) P, (f) C, (g) O, (h) Ce, (i) Au, (j) Dy, (k) Mg, (l) K, (m) Sc, (n) N, and (o) S. All panels show the same field of view from one representative sample.

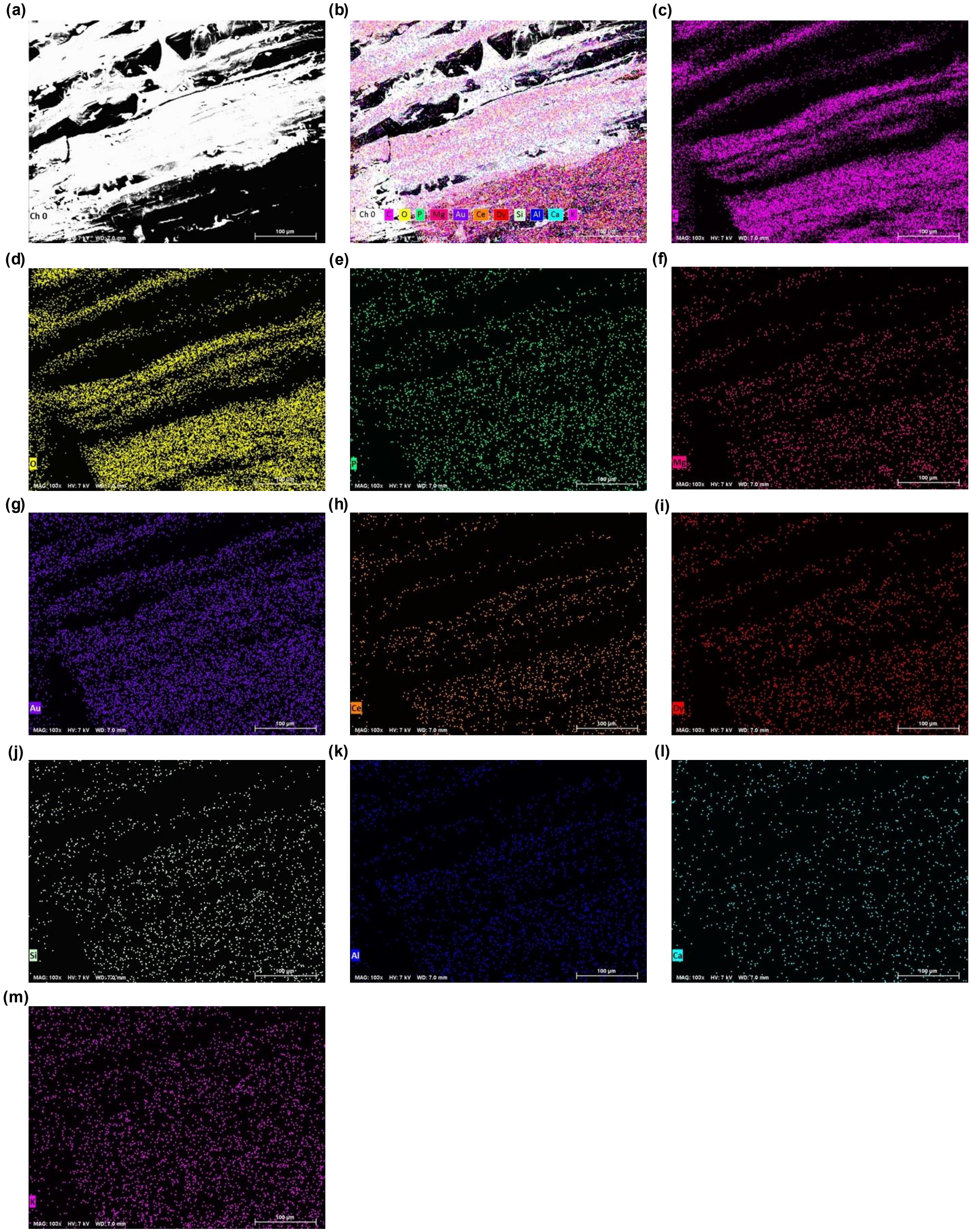


**Figure S6. SEM imaging and EDS mapping of a shoot tissue fragment from hydroponically grown control *P. americana*.** (a) Electron image of a shoot tissue fragment and (b) composite overlay of the corresponding EDS maps for dried biomass mounted on carbon tape and sputter-coated (Text S12). Individual maps show software-assigned X-ray channels assigned to (c) C, (d) O, (e) P, (f) Mg, (g) Au, (h) Ce, (i) Dy, (j) silicon, (k) Al, (l) Ca, and (m) K. All panels show the same field of view from one representative sample.

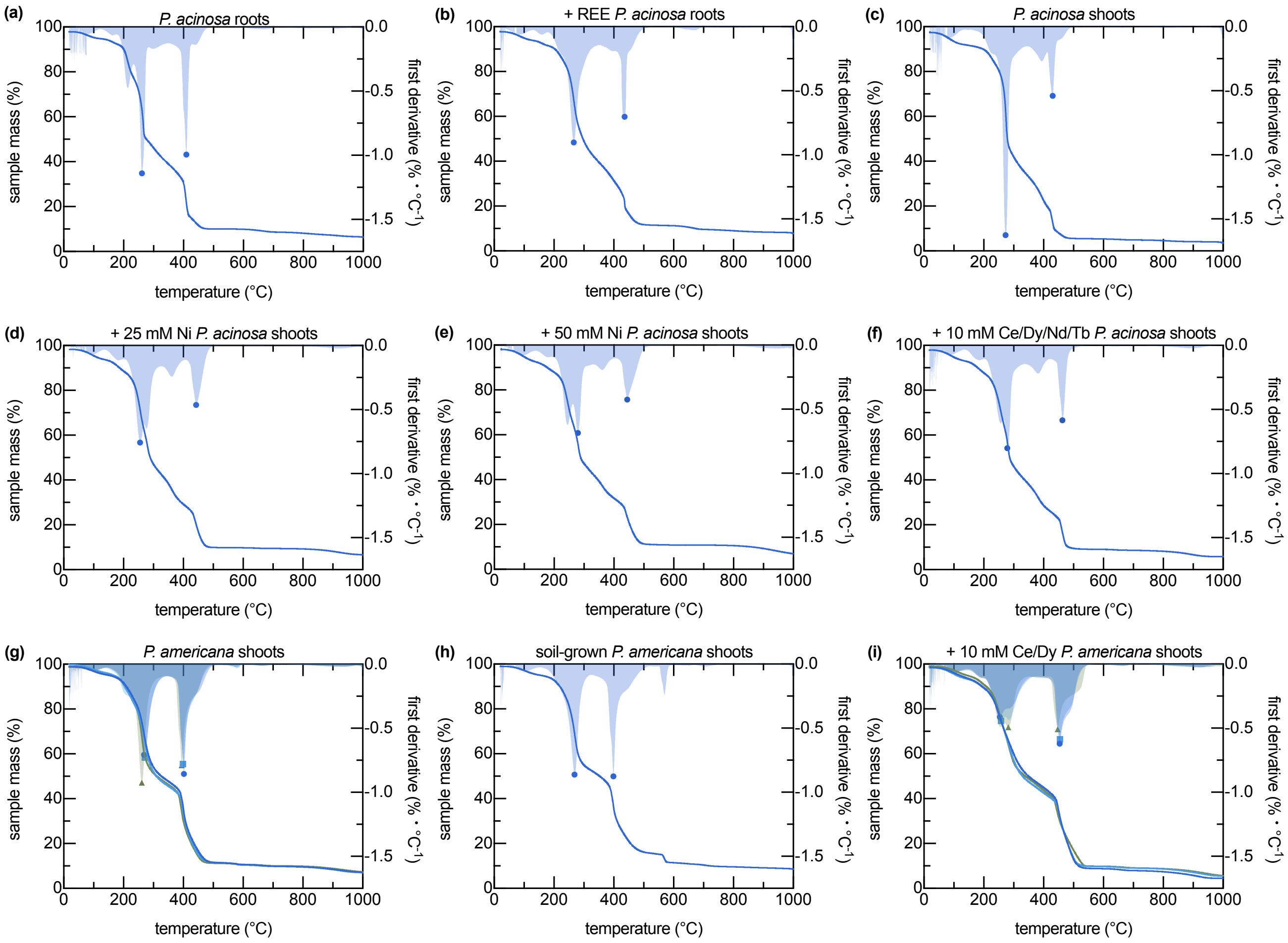


**Figure S7. TGA and DTG profiles of lyophilized *Phytolacca* tissues by tissue type, growth condition, and metal supplementation.** Normalized mass-remaining profiles (solid lines, left y-axis) and first-derivative thermogravimetric profiles (DTG; shaded traces, right y-axis) are shown for (a) *P. acinosa* roots, (b) REE-supplemented *P. acinosa* roots, (c) *P. acinosa* shoots, (d) 25 mM Ni-supplemented *P. acinosa* shoots, (e) 50 mM Ni-supplemented *P. acinosa* shoots, (f) REE-supplemented *P. acinosa* shoots, (g) hydroponic *P. americana* shoots, (h) soil-grown *P. americana* shoots, and (i) REE-supplemented *P. americana* shoots. Markers indicate local DTG minima used to extract characteristic decomposition temperatures.

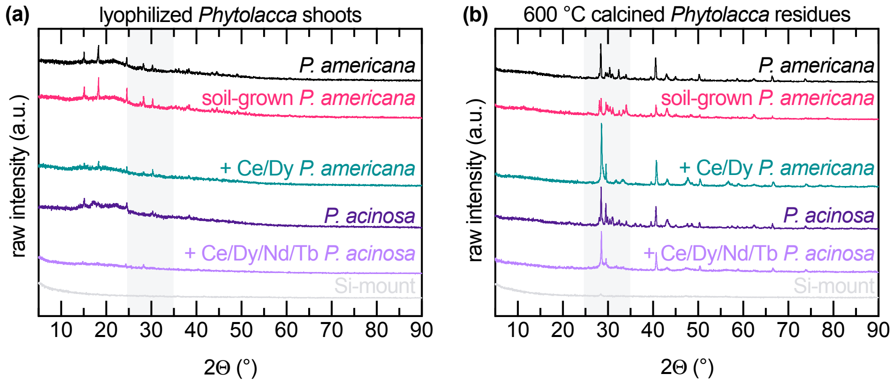


**Figure S8. Raw X-ray diffraction patterns of dried and 600 °C-calcined *Phytolacca* shoots with a silicon-mount reference.** XRD spectra are shown for (a) dried and (b) 600 °C calcined *Phytolacca* shoots (n = 1). Spectra are plotted with the same sample color order and vertical offsets for clarity; raw intensity differences should not be interpreted quantitatively as phase abundance. The Si-mount spectrum is included as a baseline to identify substrate-associated features.

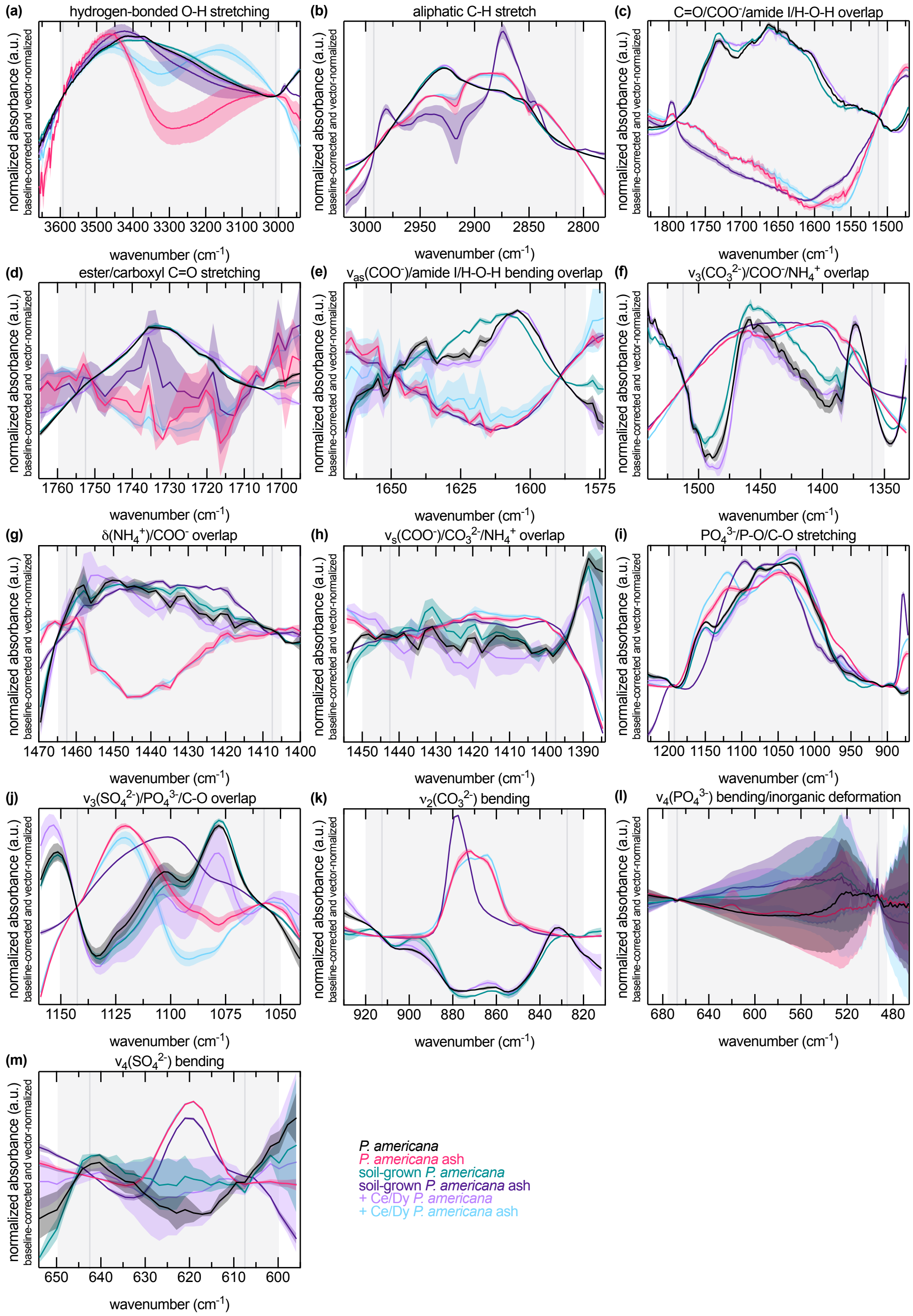


**Figure S9. Comparison of FTIR spectra for dried *Phytolacca* and ash by treatment condition after regional baseline correction and vector normalization.** Local baseline correction between the values indicated by dark gray vertical lines improved comparison of regional band shapes and intensities across samples, but the expanded comparison showed that spectral variation was not specific to REE supplementation. Lines and shaded regions show mean ± standard deviation from independent samples ($n = 3$).

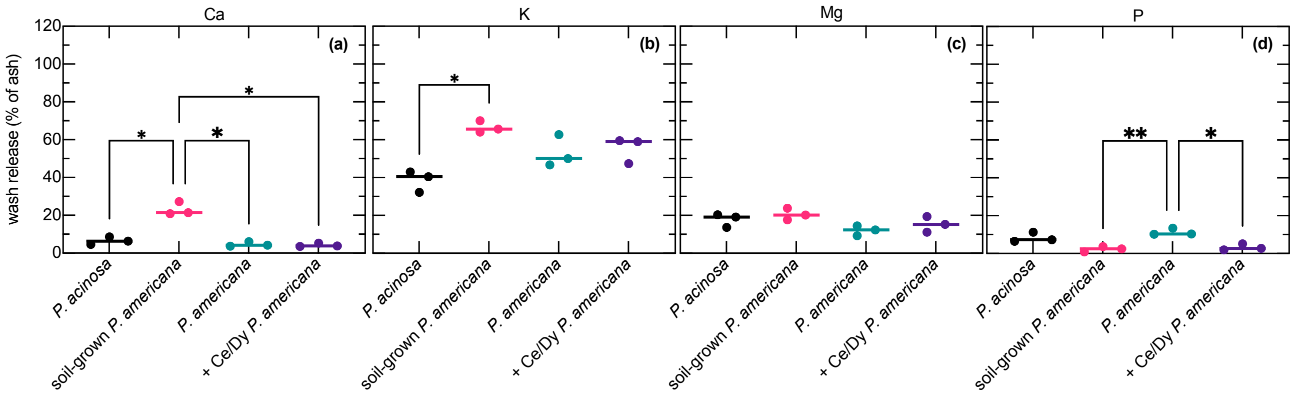


**Figure S10. Deportment of Ca, K, Mg, and P during water washing of *Phytolacca* ash produced at 600 °C.** Calcined ash generated at 600 °C from *P. acinosa*, soil- and hydroponically grown *P. americana*, and REE-enriched *P. americana* was washed with water. Elemental release was calculated as a percentage of the corresponding calcined-residue inventory for (a) Ca, (b) K, (c) Mg, and (d) P. Points show independent calcined-residue replicates, and horizontal bars show means (n = 3). Statistical comparisons were performed independently for each element using Brown–Forsythe/Welch ANOVA with Dunnett's T3 multiple comparisons. Asterisks indicate Dunnett's T3-adjusted pairwise differences ($^*P < 0.05$; $^{**}P < 0.01$).

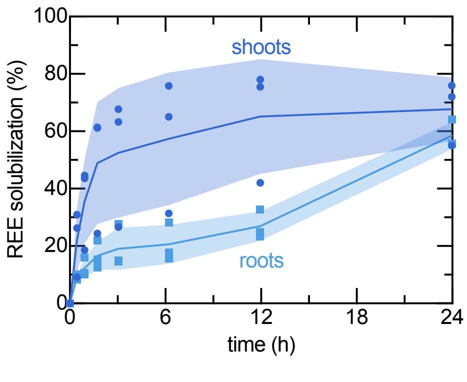


**Figure S11. Time course of REE solubilization from *P. acinosa* tissues using 0.05 M $Na_2$EDTA at pH 4.5.** Time-course measurements of REE solubilization from *P. acinosa* tissues were collected during extraction with $Na_2$EDTA (pH 4.5, 0.05 M) under shaker-flask conditions (30 °C, 150 rpm).[34] Lines and shaded regions show mean ± standard deviation, with biological replicates overlaid (n = 3).

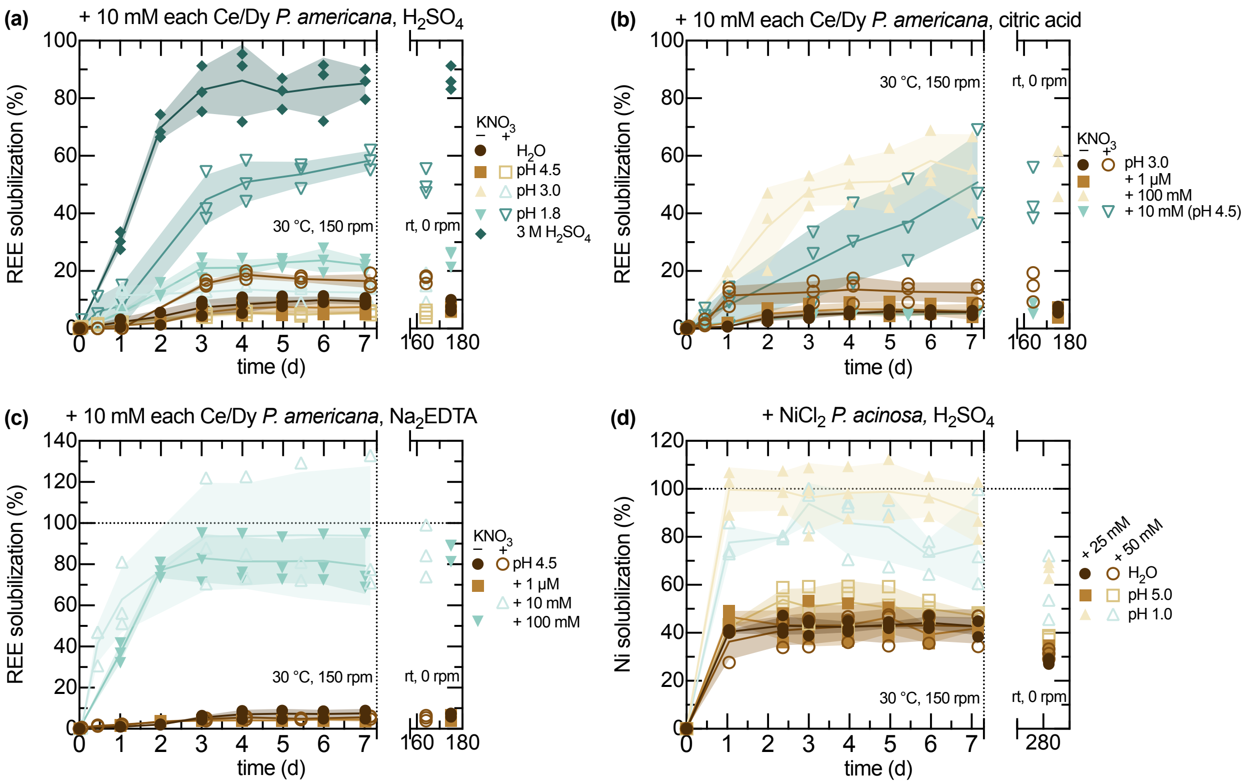


**Figure S12. Time course REE and Ni solubilization from metal-enriched *Phytolacca* shoots during acid and EDTA extraction.** Time-course REE solubilization from REE-enriched *P. americana* shoots bracketed for some low and high concentrations of (a) $H_2SO_4$, (b) citric acid (pH 3.0), and (c) $Na_2EDTA$ (pH 4.5). (d) Ni solubilization from Ni-enriched *P. acinosa* under some $H_2SO_4$ conditions. Solid–liquid extraction tests were all performed at 0.1% (w/v) solids loadings. Lines and shaded regions show mean ± standard deviation, with biological replicates overlaid (n = 3). Long-term target metal solubilization were measured after extended static storage at room temperature, which are shown as symbols not connected by the lines.

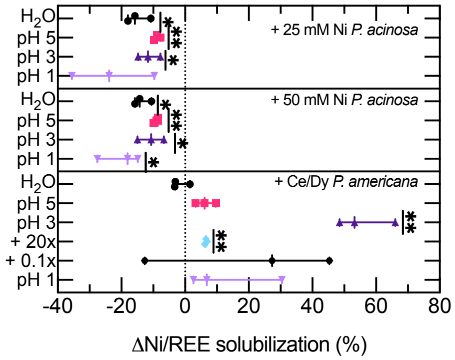


**Figure S13. Long-term REE and Ni solubilization from *Phytolacca* shoots following extended static storage, by pH and biomass loading.** After the 7 d leaching experiments at 30 °C and 150 rpm, tubes were stored under static benchtop conditions for an additional 168 d for the REE leaching tests or 274 d for the Ni leaching tests. Points show individual biological replicates (n = 3), and boxes show min-to-max distributions. $\Delta E$ values were compared against zero using two-sided one-sample $t$ tests. Asterisks indicate nominal, unadjusted significance relative to zero ($*P < 0.05$; $**P < 0.01$).

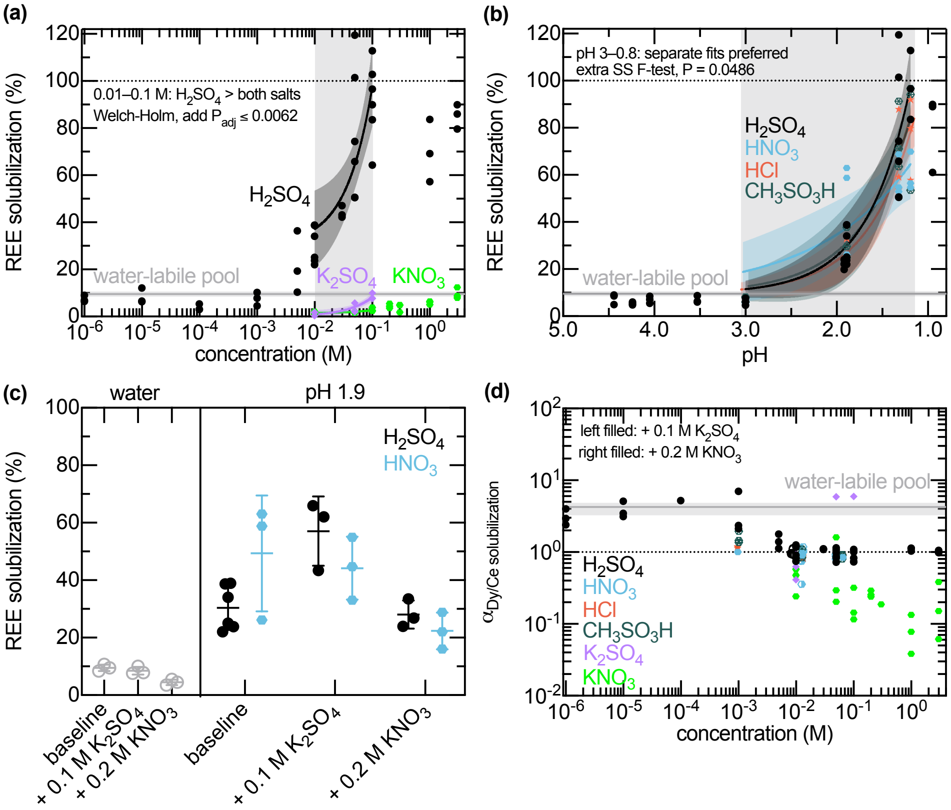


**Figure S14. Effects of mineral acid identity, pH, and electrolyte addition on REE solubilization and Dy/Ce selectivity from Ce/Dy-enriched *P. americana* shoots.** All tests used 0.1% (w/v) *P. americana* shoots. (a) REE solubilization as a function of $H_2SO_4$, $K_2SO_4$, and $KNO_3$ concentration. Solid lines show fitted logarithmic models [REE solubilization $= a \cdot \log([\text{reactant}] + 1) + b$], and shaded regions show 95% confidence intervals. (b) REE solubilization as a function of pH for $H_2SO_4$, $HNO_3$, HCl, and $CH_3SO_3H$. Solid lines show fitted exponential models [REE solubilization $=$ REE solubilization$_{H_2O} + A \cdot 10^{-k \cdot \text{pH}}$], and shaded regions show 95% confidence intervals. (c) REE solubilization in $H_2O$ and some mineral acids at pH 1.9 under varied electrolyte supplementation conditions. (d) Apparent selectivity of Dy to Ce solubilization as a function of reactant concentrations for the data underlying (a-c). Points show individual biological replicates (n = 3–6), and the horizontal band indicates the $H_2O$ baseline condition.

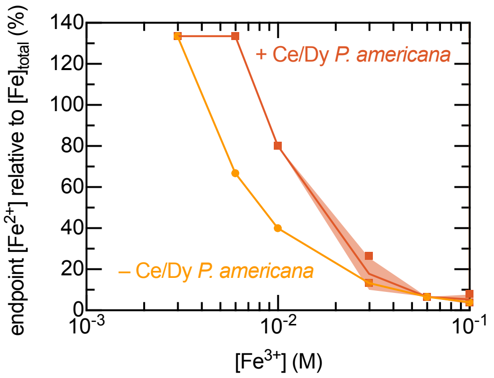


**Figure S15. Apparent endpoint $Fe^{2+}$ speciation as a function of initial $Fe^{3+}$ concentrations incubated with and without Ce/Dy-enriched *P. americana* shoots.** Endpoint $Fe^{2+}$ was measured by cerium sulfate titration and normalized to total Fe measured by ICP-OES as $[Fe^{2+}]$final/[Fe]final $\cdot$ 100%. oints show apparent endpoint $Fe^{2+}$ recovery for samples incubated with or without Ce/Dy-enriched *P. americana* shoots across the indicated initial $Fe^{3+}$ range relevant to conventional *A. ferrooxidans* bioleaching experiments. Lines and shaded regions show mean and standard deviation, respectively, with biological replicates overlaid as points (n = 3). Apparent recoveries exceeding 100% at the lowest $Fe^{3+}$ concentrations reflect limited sensitivity of the titration near the lower end of the tested $Fe^{2+}$ range and should not be interpreted as $Fe^{2+}$ production exceeding the initial iron pool.

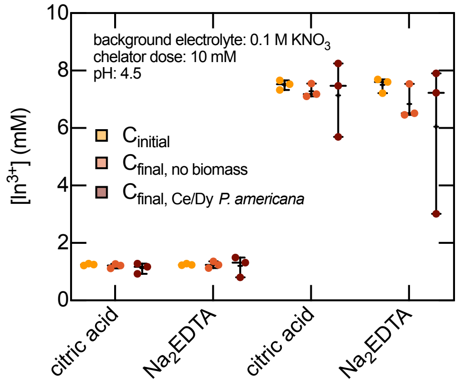


**Figure S16. Initial and final soluble In concentrations in citric acid and EDTA competition tests after incubation with and without *P. americana* shoots.** Bulk In concentrations measured initially did not significantly differ from concentrations measured after 7 d incubation with or without *P. americana* shoots for either citric acid or $Na_2$EDTA at either tested initial In loading (two-way ANOVA with Tukey's multiple-comparisons test, all adjusted $P > 0.13$). Points show individual replicates (n = 3), black bars show min–median–max distributions, and plus signs indicate means.

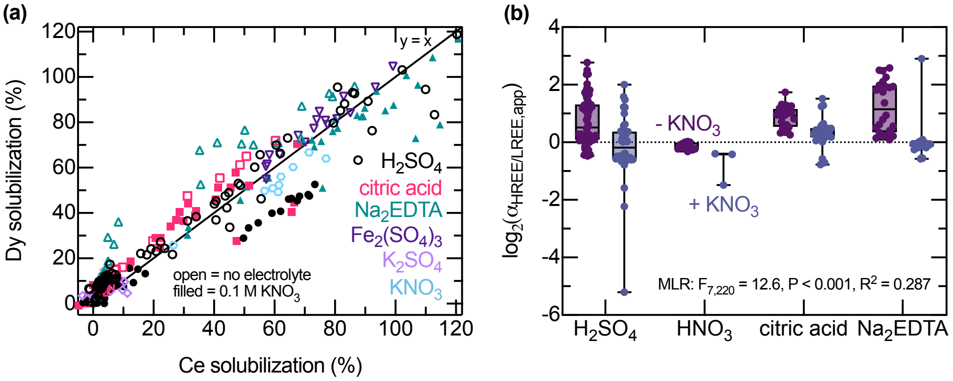


**Figure S17. Dy and Ce solubilization and feed-normalized HREE/LREE selectivity from *P. americana* shoots after solid–liquid extraction with and without $KNO_3$ addition.** (a) Paired Dy and Ce solubilization across the indicated leaching conditions; the solid $y = x$ line denotes equal fractional Dy and Ce release. Open symbols indicate no added background electrolyte, whereas filled symbols indicate 0.1 M $KNO_3$. (b) Feed-normalized molar selectivity, $\alpha_{HREE/LREE} = (n_{HREE,aq}/n_{LREE,aq})/(n_{HREE,feed}/n_{LREE,feed})$, is plotted as $\log_2(\alpha_{HREE/LREE})$ for the displayed conditions. Purple and blue symbols indicate conditions without and with 0.1 M $KNO_3$, respectively. The dotted line at zero indicates equal fractional HREE and LREE release relative to the feed; positive values indicate preferential HREE solubilization. Individual observations are overlaid on box plots. A regression of HREE/LREE selectivity that also includes $HNO_3$ conditions is reported in Table S25.

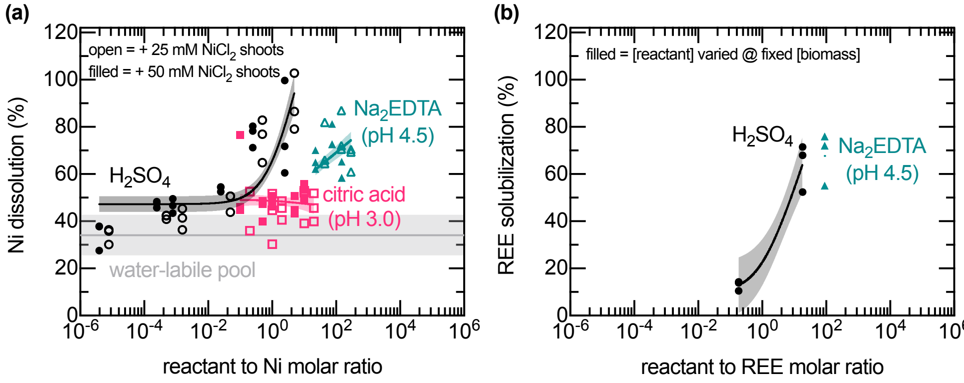


**Figure S18. Ni and REE solubilization from enriched *P. acinosa* shoots as a function of reagent dosage during acid and chelator extraction.** (a) Ni solubilization was plotted as a function of reactant:Ni molar ratio for varied doses of $H_2SO_4$, citric acid (pH 3.0), and $Na_2EDTA$ (pH 4.5), compared with the water-extractable pool at a fixed 0.1% (w/v) biomass loading. Open symbols denote +25 mM $NiCl_2$-supplemented tissues, and filled symbols denote +50 mM $NiCl_2$-supplemented shoots. (b) REE solubilization from REE-supplemented shoots increased with $H_2SO_4$ dose and was also observed under the single tested $Na_2EDTA$ condition (pH 4.5). Points show replicate-level measurements (n = 3), solid lines show fitted logarithmic models [$Y = a \cdot \log(X + 1) + b$], and shaded regions show 95% confidence intervals.

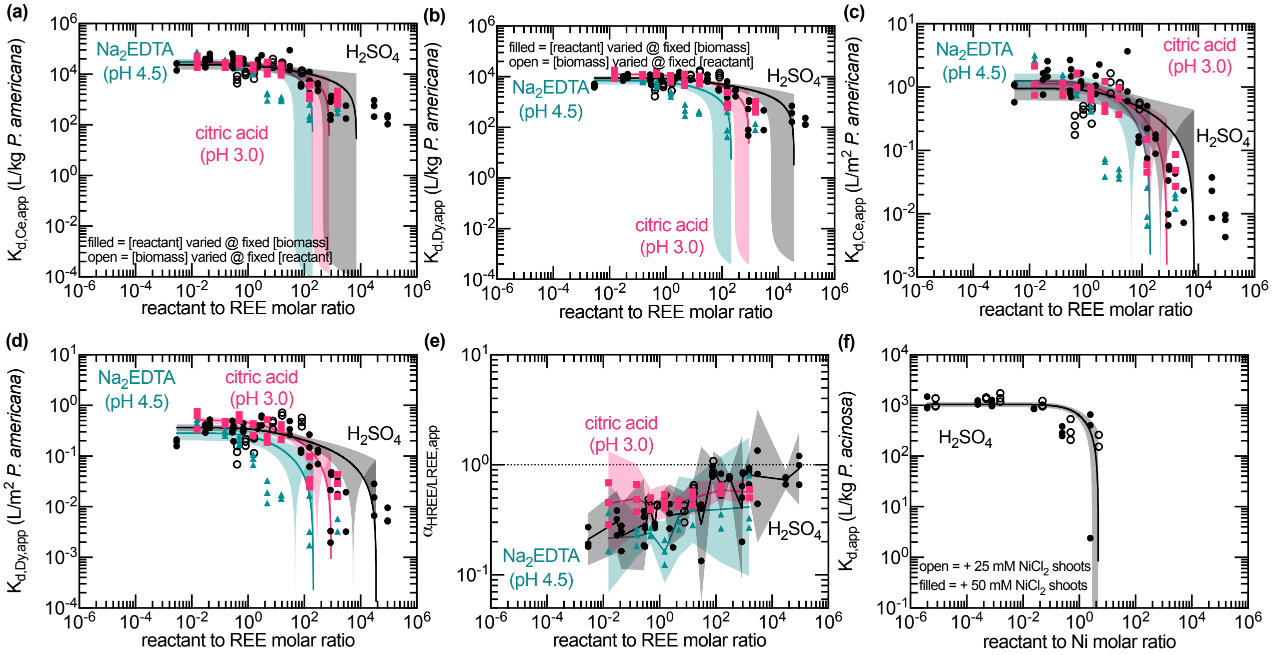


**Figure S19. Condition-specific apparent solid–liquid distribution coefficients and Dy/Ce selectivity during REE and Ni extraction from *Phytolacca* shoots as a function of reagent concentration.** Endpoint measurements of target metal solubilization for solid–liquid extraction reactant with bounds of time-course data shown in Figure S12. Condition-specific apparent solid–liquid distribution coefficients ($K_{\mathrm{d,i,app}}$) as a function of initial reactant concentration for Ce and Dy solubilized from *P. americana* (a-b) before and (c-d) after surface-area normalization, respectively. Solid lines show fitted logarithmic models [$K_{\mathrm{d,i,app}} = a \cdot \log(\mathrm{reactant{:}i} + 1) + b$]. (e) Condition-specific apparent selectivity of HREE over LREE, with $Dy^{3+}$ and $Ce^{3+}$ representing the HREE and LREE, respectively, as a function of initial reactant concentration for the data plotted in (a-d). Solid lines show geometric means. (f) Condition-specific apparent $K_{\mathrm{d,Ni,app}}$ for Ni solubilized from *P. acinosa*. Solid lines show fitted logarithmic models [$K_{\mathrm{d,Ni,app}} = a \cdot \log(\mathrm{reactant{:}Ni} + 1) + b$]. Shaded regions show 95% confidence intervals in each panel, with independent replicate values overlaid as points ($n = 3$ per condition).

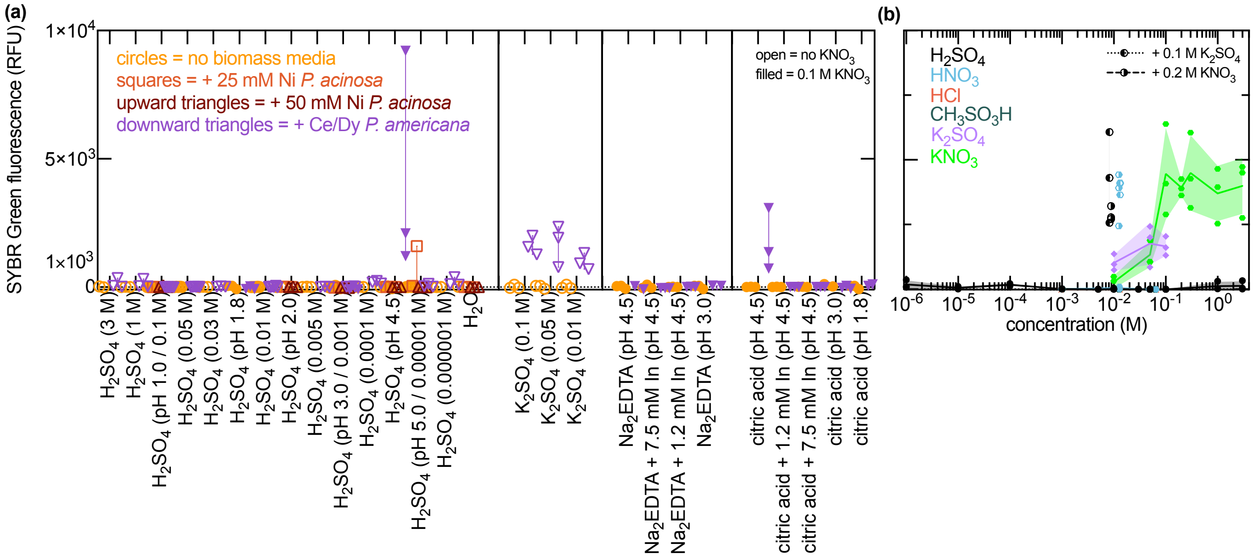


**Figure S20. Endpoint SYBR Green fluorescence in Ni- and REE-enriched *Phytolacca* shoot leachates and matched no-biomass controls across extraction conditions.** Endpoint leachates and matched no-biomass media controls were assayed with SYBR Green to evaluate fluorescence signals associated with plant-derived nucleic-acid or matrix components released during extraction. (a) Circles indicate no-biomass media controls, squares indicate leachates from *P. acinosa* shoots supplemented with 25 mM $NiCl_2$, upward triangles indicate leachates from *P. acinosa* shoots supplemented with 50 mM $NiCl_2$, and downward triangles indicate leachates from Ce/Dy-supplemented *P. americana* shoots. Open symbols indicate conditions without added $KNO_3$; filled symbols indicate conditions containing 0.1 M $KNO_3$. Extraction conditions are grouped by reagent identity and separated by vertical lines. Points show replicate measurements reported as relative fluorescence units (RFU; n = 3–6). (b) Symbols correspond to the scheme from Figure S14d.

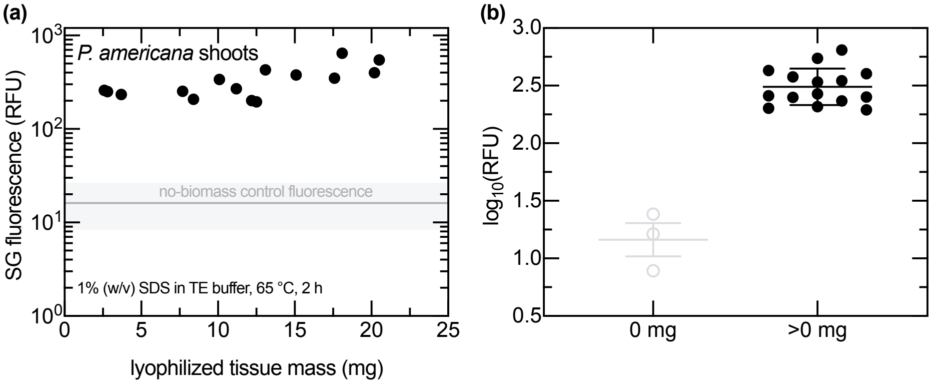


**Figure S21. SYBR Green fluorescence of *P. americana* shoot lysates as a function of tissue mass after SDS and heat treatment.** Lyophilized *P. americana* shoot tissues were lysed in 1% (w/v) SDS in TE buffer at 65 °C for 2 h, and lysates were analyzed by SYBR Green fluorescence. (a) Fluorescence of biomass-containing lysates plotted as a function of lyophilized tissue mass; the horizontal reference band indicates the fluorescence range of no-biomass controls processed under the same lysis conditions. (b) $\log_{10}$-transformed fluorescence values for no-biomass controls (0 mg) and biomass-containing lysates (>0 mg). Biomass-containing lysates showed significantly higher fluorescence than no-biomass controls by two-tailed unpaired Welch's $t$ test on $\log_{10}$-transformed RFU values ($P = 0.0076$). Points show individual lysates (n = 3), and horizontal bars show mean ± standard deviation.

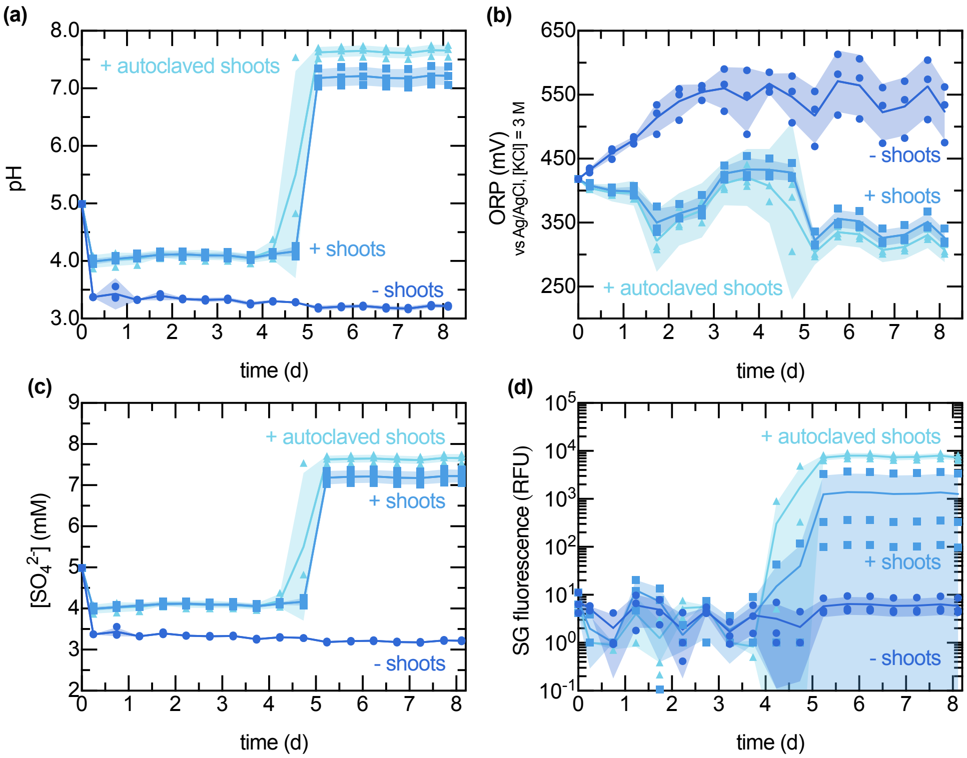


**Figure S22. Time course pH, ORP, dissolved sulfate, and SYBR Green fluorescence during water incubation of autoclaved and non-autoclaved *P. americana* shoots.** Non-REE-enriched hydroponically grown *P. americana* shoots were incubated at 1% (w/v) in sterile Milli-Q water initially adjusted to pH 5.0. No-shoot controls, non-autoclaved shoot treatments, and autoclaved shoot treatments were monitored over 8 d. Time-course measurements show (a) pH, (b) oxidation-reduction potential (ORP) measured versus an Ag/AgCl reference electrode in 3 M KCl, (c) dissolved sulfate concentration, and (d) SYBR Green fluorescence. Lines and shaded regions show mean and standard deviation, respectively, with biological replicates overlaid (n = 3).

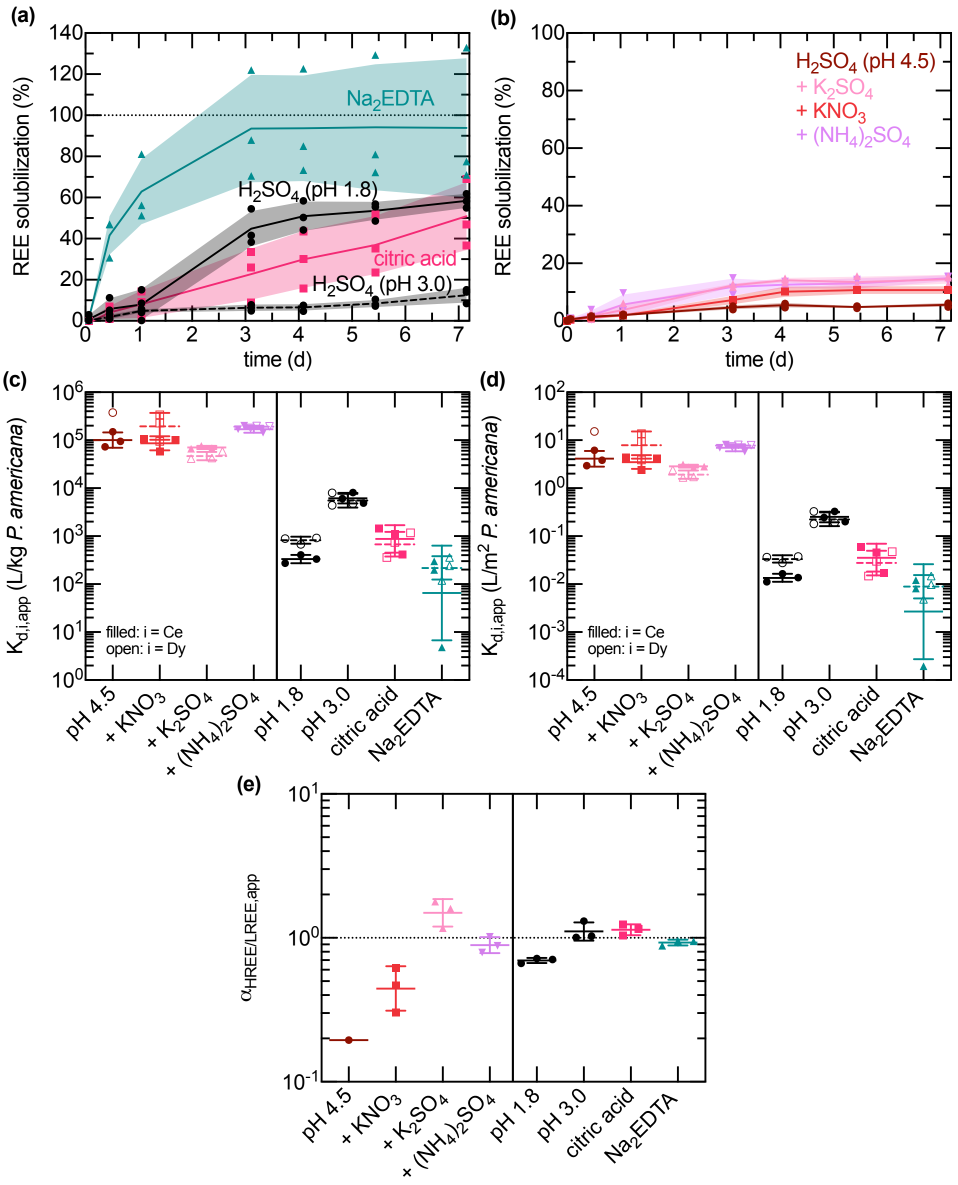


**Figure S23. REE solubilization time courses, apparent solid–liquid distribution coefficients, and HREE/LREE selectivity during acid, chelator, and electrolyte extraction of *P. americana* shoots.** Separate REE solubilization versus time data for experiments performed using the same (a) acid and chelator (0.01 M), and (b) electrolyte-supplemented (0.2 M $KNO_3$ and 0.1 M $K_2SO_4$) solid–liquid extraction conditions used in Figure 3. Lines and shaded regions show mean and standard deviation, respectively, with independent replicates overlaid as points (n = 3). Condition-specific apparent solid–liquid distribution coefficients, (c) without and (d) with surface-area-normalization, and (e) apparent selectivity of HREE over LREE, with $Dy^{3+}$ (filled symbols; solid lines) and $Ce^{3+}$ (open symbols; dashed lines) representing the HREE and LREE, respectively, calculated from extraction endpoints for conditions shown in (a-b). Wide and narrow lines indicate geometric means and standard deviations, respectively ($n = 3$). Asterisks indicate conditions which had negative $K_{d,i,app}$ which could not be shown on these plots. All conditions has 0.1 M $KNO_3$ background electrolyte to reduce the magnitude of ionic strength differences.

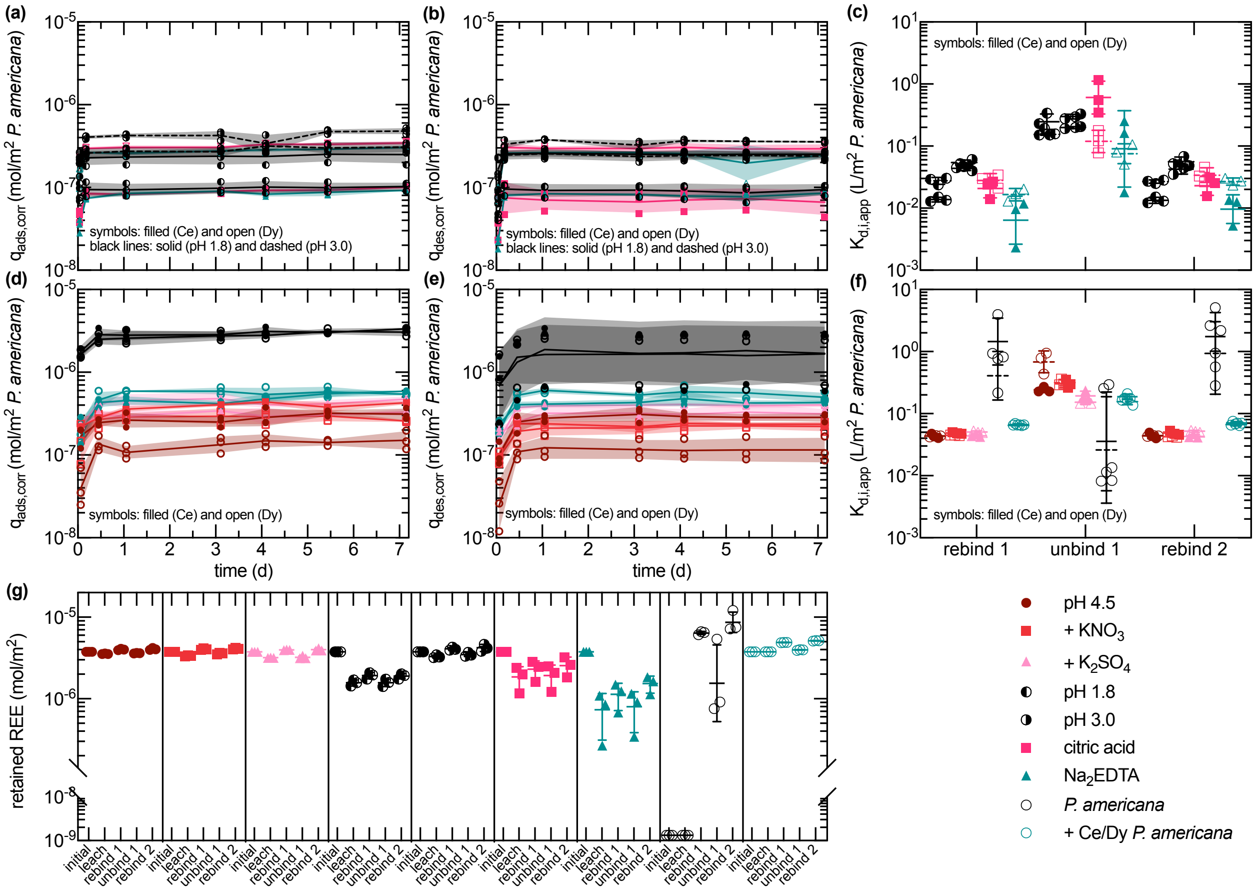


**Figure S24. Ce and Dy uptake, release, and retention during sequential rebinding, unbinding, and rebinding of previously leached *P. americana* shoot residues.** For the post leached residues generated from the experiments represented by Figure S23, apparent net Ce and Dy uptake during (a,d) first rebinding and (b,e) release during subsequent unbinding are shown for (a,b) acid- and chelator-treated residues and (d,e) electrolyte-treated residues and unleached controls. Lines and shaded regions show arithmetic mean and standard deviation, respectively, with independent replicates overlaid as points (n = 3). (c,f) Condition-specific surface-area-normalized apparent solid–liquid distribution coefficients at the endpoints of first rebinding, unbinding, and second rebinding. Wide and narrow lines indicate geometric means and standard deviations, respectively, with independent replicates overlaid as points ($n = 3$). (g) Total surface-area-normalized retained REE before leaching and after each successive stage. Wide and narrow lines indicate arithmetic means and standard deviations, respectively, with independent replicates overlaid as points ($n = 3$).

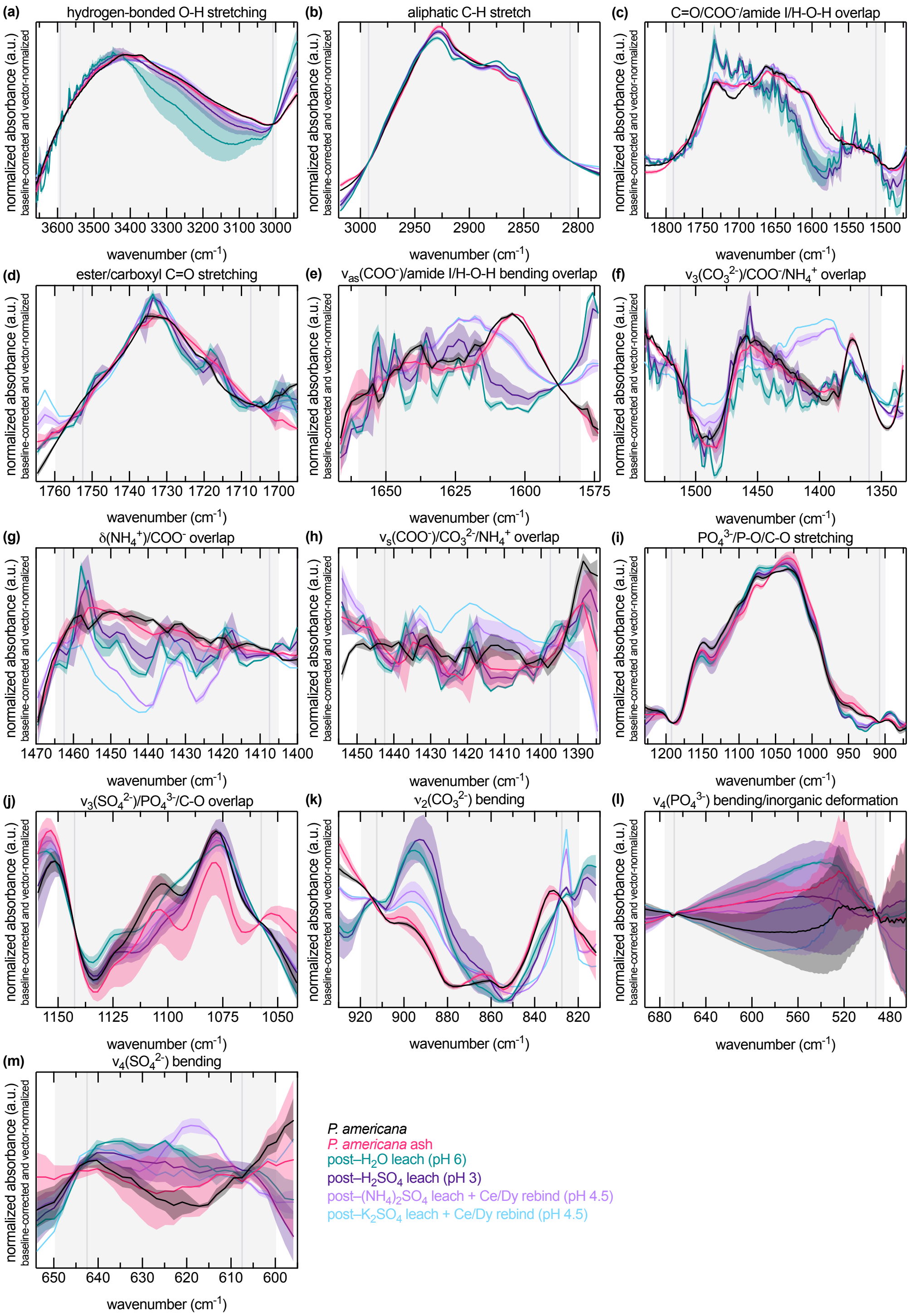


**Figure S25. Baseline-corrected and vector-normalized FTIR spectra of *P. americana* shoots before extraction, after leaching, and after Ce/Dy reloading.** Baseline-corrected and vector-normalized FTIR spectra are shown for Ce/Dy-enriched *P. americana* shoots and control biomass, post-leached residues after water extraction at pH 6, post-leached residues after $H_2SO_4$ extraction at pH 3, and post-leached residues after $(NH_4)_2SO_4$ or $K_2SO_4$ extraction followed by Ce/Dy reloading at pH 4.5. Local baseline correction between the values indicated by dark gray vertical lines improved comparison of regional band shapes and intensities across samples. Lines and shaded regions show mean ± standard deviation from biological replicates ($n = 3$).

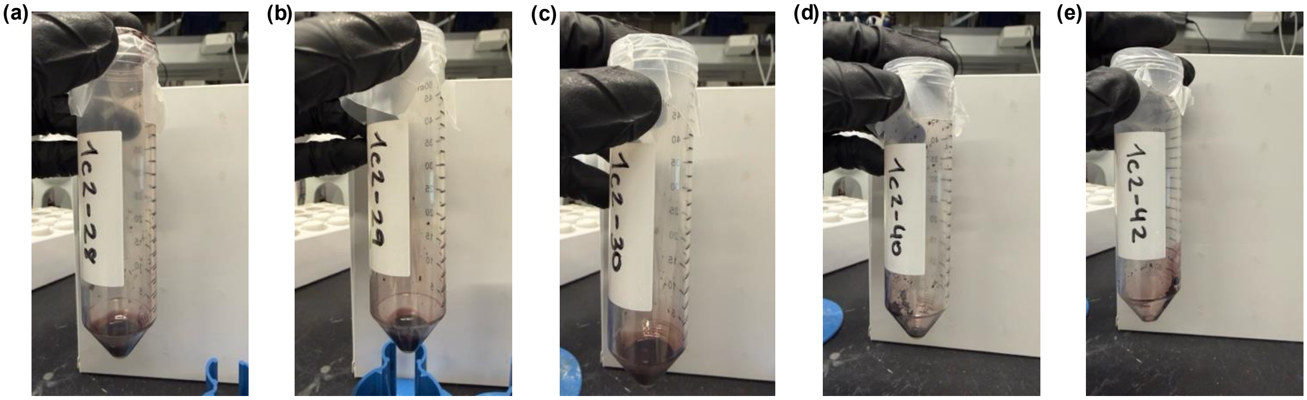


**Figure S26. Photographs of residues formed during attempted lyophilization of *P. americana* shoot leachates generated with 0.1 and 3 M $H_2SO_4$.** Representative photographs are shown for extracts generated after leaching *P. americana* shoots in (a–c) 3 M and (d,e) 0.1 M $H_2SO_4$. Attempts to lyophilize these leachates produced viscous purple residues rather than fully dried powders, consistent with concentration of acid-soluble plant-derived organic material in the leachate fraction. Images are included as qualitative documentation of the observed residue morphology and color.

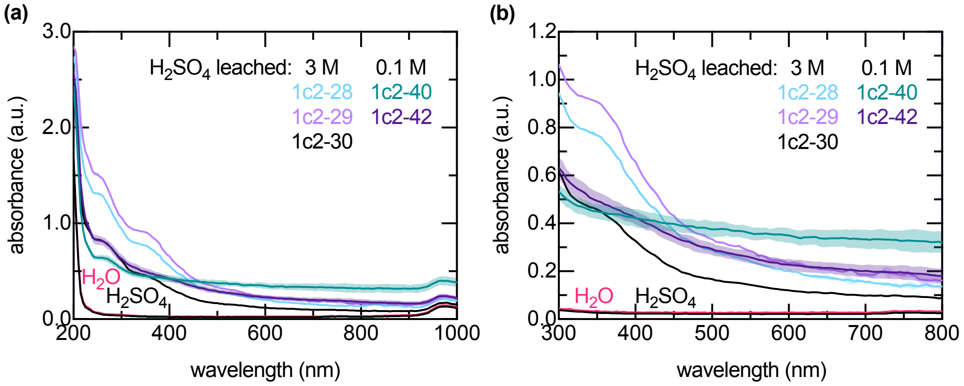


**Figure S27. UV–visible absorbance spectra of diluted residues obtained by attempted lyophilization of $H_2SO_4$ leachates from *P. americana* shoots.** Viscous purple residues formed after attempted lyophilization of *P. americana* shoot leachates generated with 0.1–3 M $H_2SO_4$ were diluted 100-fold in $H_2O$ and analyzed by UV–visible absorbance spectroscopy across the (a) 200–1,000 nm and (b) 300–800 nm ranges. The same data are shown in panels (a) and (b); panel (b) is provided for magnification purposes. Lines and shaded regions show mean ± standard deviation from three technical replicates per biological sample (n = 2–3).

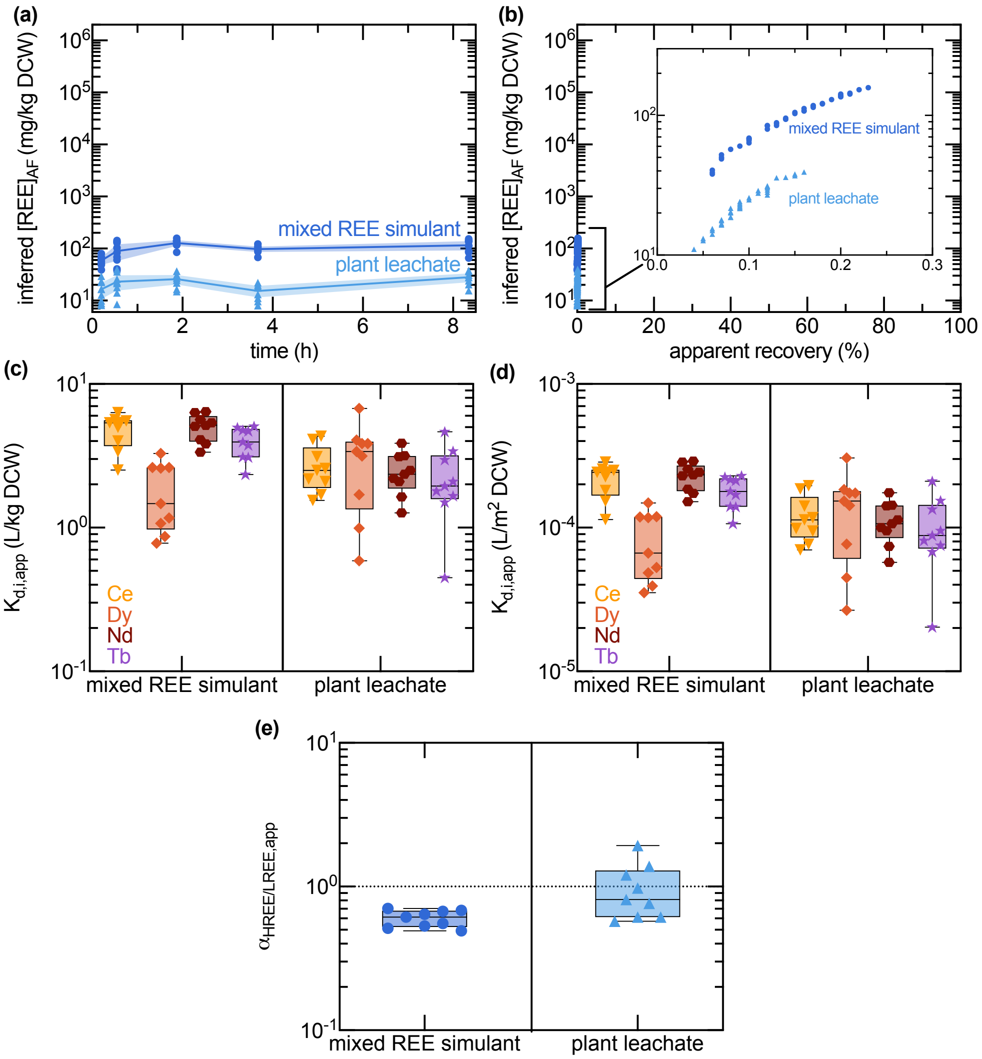


**Figure S28. Time-dependent inferred REE loading, grade–recovery relationships, and apparent partitioning by *A. ferrooxidans* in mixed-REE simulants and *P. acinosa* shoot leachate at pH 1.8.** (a) Time-dependent inferred REE loading on $OD_{600} = 1.0$ *A. ferrooxidans* biomass is shown after exposure to either an equimolar $Ce_2/Dy_2/Nd_2/Tb_2\text{-}(SO_4)_3$ simulants or isohydric *P. acinosa* shoot leachate at pH 1.8. Inferred REE loading is reported as mg REE per kg dry cell weight (DCW). Lines and shaded regions show mean ± 95% confidence intervals, with biological replicates overlaid (n = 9). (b) Apparent grade–recovery curve for the data from (a). Inset shows close-up of data for visual inspection. Condition-specific apparent solid–liquid distribution coefficients, (c) without and (d) with surface-area-normalization, for the endpoint data from (a), and the corresponding (e) apparent HREE to LREE molar selectivity. $Dy^{3+}$ and $Tb^{3+}$ represent HREEs, and $Ce^{3+}$ and $Nd^{3+}$ represent LREEs. Biological replicates are shown as symbols, with wide bars indicating the mean and whiskers the minimum–maximum range.

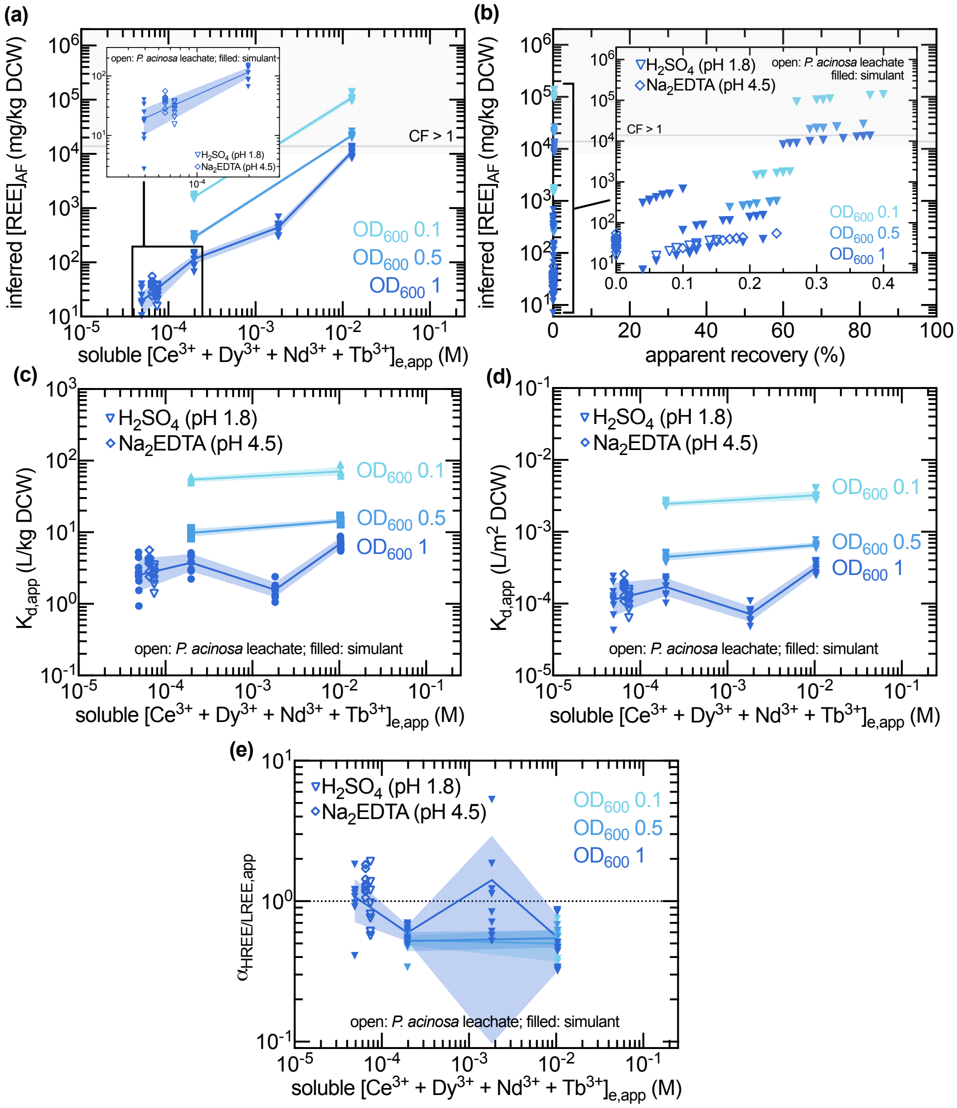


**Figure S29. Inferred REE binding, grade–recovery relationships, and apparent HREE/LREE selectivity of *A. ferrooxidans* as a function of initial mixed-REE concentrations and cell densities.** (a) Inferred REE associated with *A. ferrooxidans* as a function of the apparent mixed-REE equilibrium concentration and initial cell density, and the underlying (b) grade–recovery curve, condition-specific apparent solid–liquid distribution coefficients, (c) without and (d) with surface-area-normalization, and (e) HREE/LREE molar selectivity. $Dy^{3+}$ and $Tb^{3+}$ represent HREEs, and $Ce^{3+}$ and $Nd^{3+}$ represent LREEs. Lines and shaded band represent mean ± standard deviations for (a) and geometric means ± standard deviations for (c–e), with biological replicates overlaid as points ($n = 3 - 9$). The gray horizontal reference line indicates the concentration factor (CF) threshold of 1 relative to the corresponding reference for the REE-enriched *P. americana* shoots. Insets in (a-b) show close-ups of data for visual inspection.

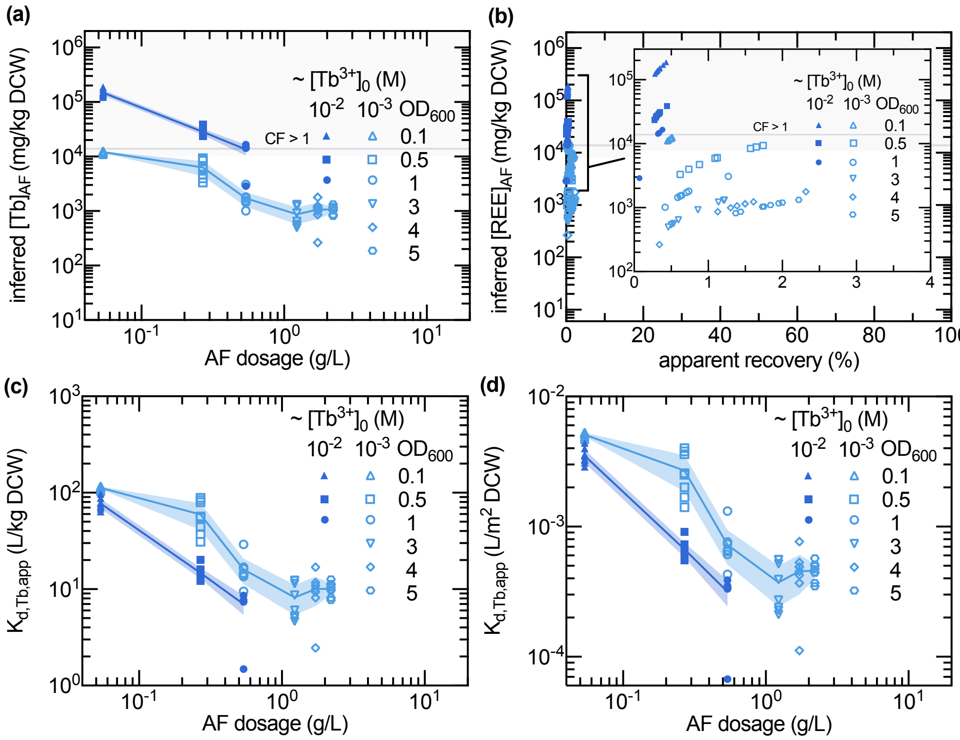


**Figure S30. Inferred Tb binding, grade–recovery relationships, and apparent solid–liquid distribution coefficients of *A. ferrooxidans* as a function of initial Tb concentrations and cell densities.** (a) Inferred Tb associated with *A. ferrooxidans* as a function of initial cell density and Tb concentration, and the underlying (b) grade–recovery curve and condition-specific apparent solid–liquid distribution coefficients, (c) without and (d) with surface-area-normalization. Lines and shaded band represent geometric mean ± standard deviations for (a,c), with biological replicates overlaid as points ($n = 3 - 9$). The gray horizontal reference line indicates the concentration factor (CF) threshold of 1 relative to the corresponding reference for the REE-enriched *P. americana* shoots. Insets in (b) show close-ups of data for visual inspection.

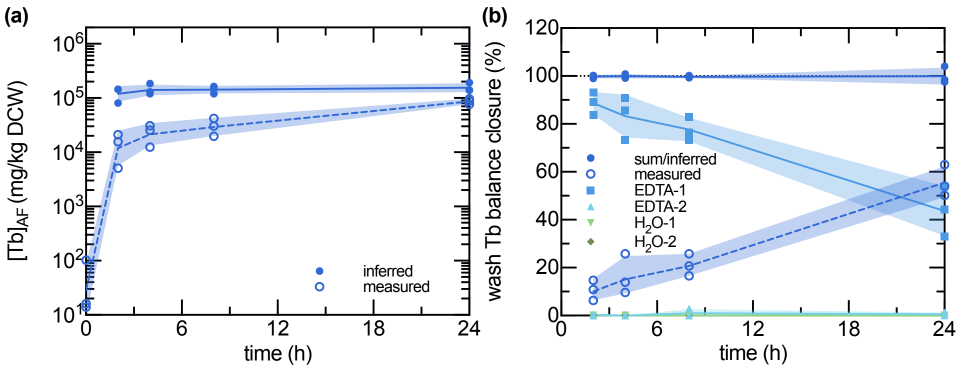


**Figure S31. Effect of binding duration on inferred Tb association with *A. ferrooxidans* and Tb distribution across sequential EDTA washes, water washes, and final pellets.** Binding tests were performed in AFM1 BS supplemented with 12.2 mM Tb from $Tb_2(SO_4)_3$ at pH 1.8 under standard conditions ($OD_{600} = 0.1$, 30 °C, 150 rpm). (a) Inferred and measured Tb associated with *A. ferrooxidans* as a function of EDTA washing duration. (b) Tb recovered in the first and second EDTA washes (EDTA-1 and EDTA-2), subsequent water washes ($H_2O$-1 and $H_2O$-2), and final pellet digest (measured), expressed as percentages of the corresponding inferred prewash Tb using Equation S55. "Sum/inferred" represents wash-stage mass-balance closure calculated using Equation S53. The dotted horizontal line indicates 100% mass balance closure. Lines and shaded bands represent mean ± standard deviations, with biological replicates overlaid as points ($n = 3$).

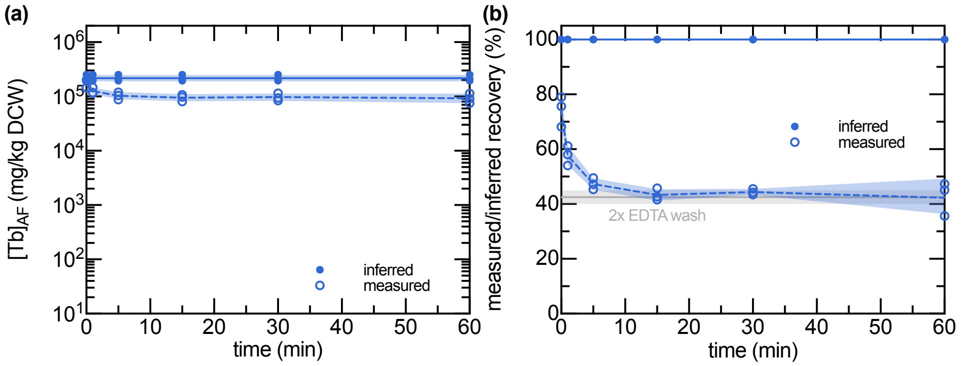


**Figure S32. Effect of EDTA washing duration and protocol on cell-associated Tb retention and recovery after 24 h binding by *A. ferrooxidans*.** Binding tests were performed in AFM1 BS supplemented with 12.2 mM Tb from $Tb_2(SO_4)_3$ at pH 1.8 under standard conditions ($OD_{600} = 0.1$, 30 °C, 150 rpm, 24 h). (a) Inferred and measured Tb associated with *A. ferrooxidans* as a function of binding duration. (b) Tb recovered in the first and second EDTA washes (EDTA-1 and EDTA-2), subsequent water washes ($H_2O$-1 and $H_2O$-2), and final pellet digest (measured), expressed as percentages of the corresponding inferred prewash Tb. "Sum/inferred" represents the summed Tb fractions recovered across the four washes and the final pellet digest divided by the inferred Tb mass bound. The dotted horizontal line indicates 100% mass balance closure. Lines and shaded bands represent mean ± standard deviations, with biological replicates overlaid as points ($n = 3$). The percentage of inferred prewash Tb retained in the final washed pellet was calculated using Equation S52.

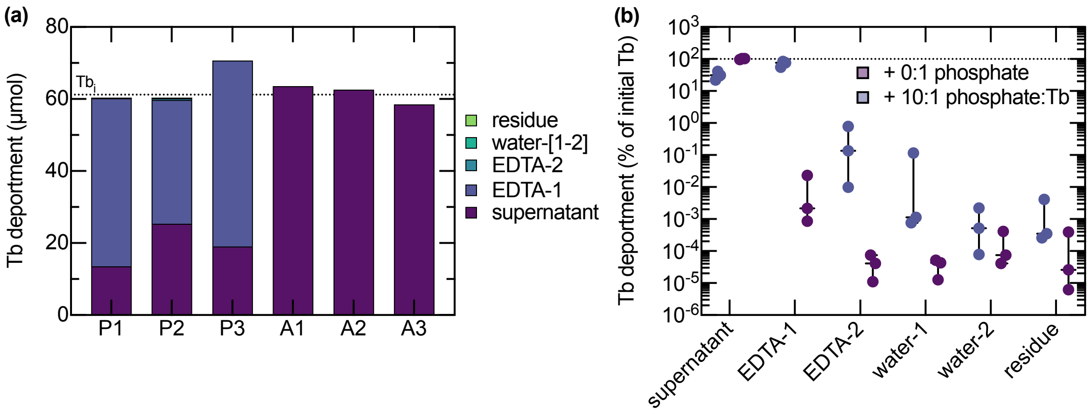


**Figure S33. Tb distribution and recovery after cell-free incubation with or without excess phosphate and subsequent sequential washing.** Binding tests were performed in AFM1 BS supplemented with 12 mM Tb from $Tb_2(SO_4)_3$ with (P1–P3) and without (A1–A3) a 10:1 molar ratio of phosphate to Tb from PBS under standard conditions (pH, 1.8 $OD_{600}$ = 0.0, 30 °C, 150 rpm, 24 h). (a) Tb deportment across post-incubation fractions for each sample, with the measured initial Tb inventory indicated by the dotted horizontal line. (b) Tb recovered in each fraction, including the separately measured water washes, expressed as a percentage of the initial Tb inventory using Equation S55, on a logarithmic scale. Box-and-whiskers indicate the minimum–maximum ranges with the median shown by the wider horizontal line, with individual replicates overlaid as points ($n$ = 3). The dotted horizontal line indicates 100% of the initial Tb inventory.

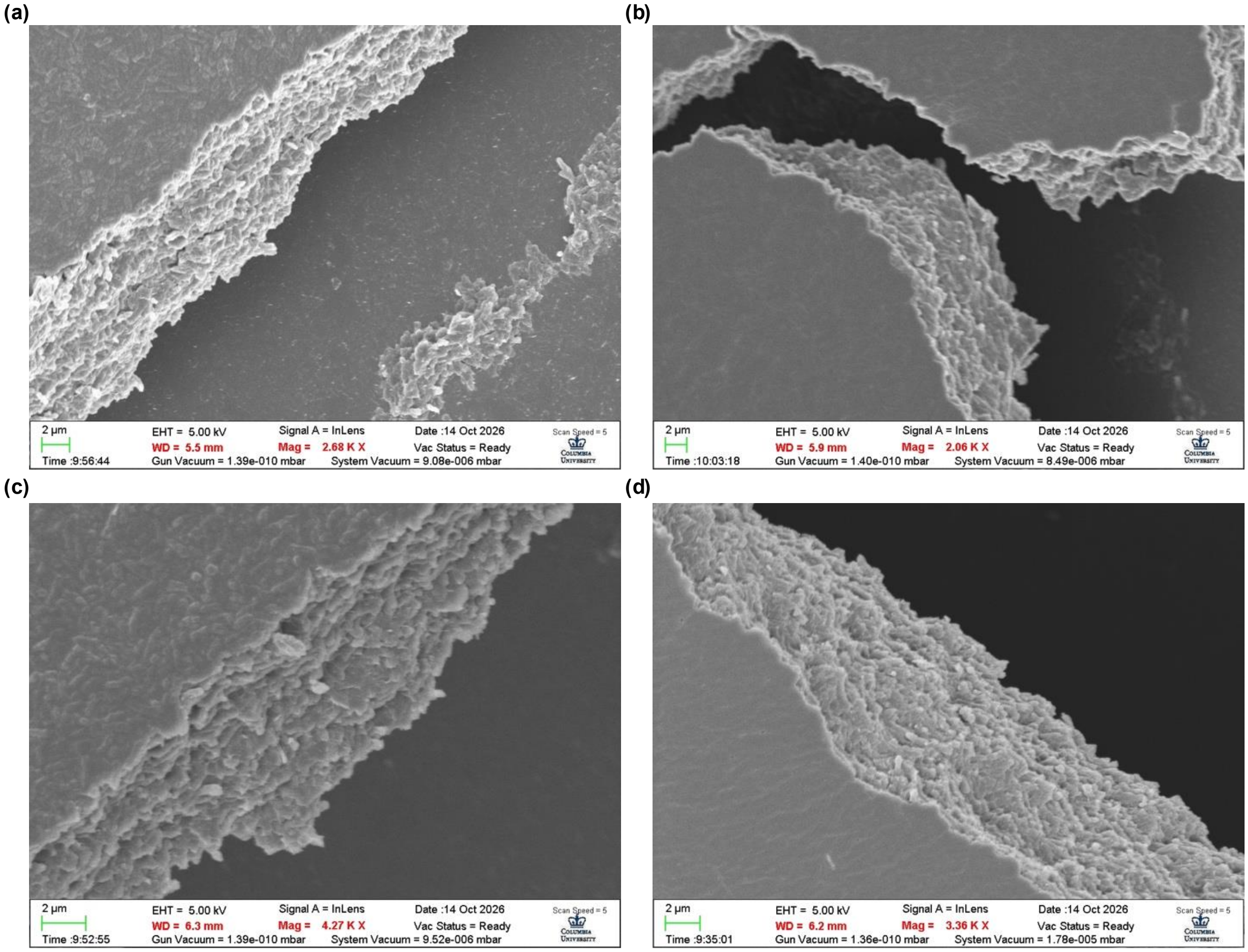


**Figure S34. SEM images of *A. ferrooxidans* preparations after 24 h incubation with or without Tb and subsequent EDTA or water-only washing.** Samples were collected after identical 24 h binding conditions as tests shown in Figure S32. Preparations were (a) without Tb exposure or EDTA washing, (b) without Tb exposure and with EDTA washing, (c) with Tb exposure and without EDTA washing, and (d) with both Tb exposure and EDTA washing. Preparations without EDTA received water-only washing. Lyophilized biomass was dispersed in water on silicon wafers, dried, and sputter-coated before imaging (Text S12). One selected field from one prepared sample per condition is shown.

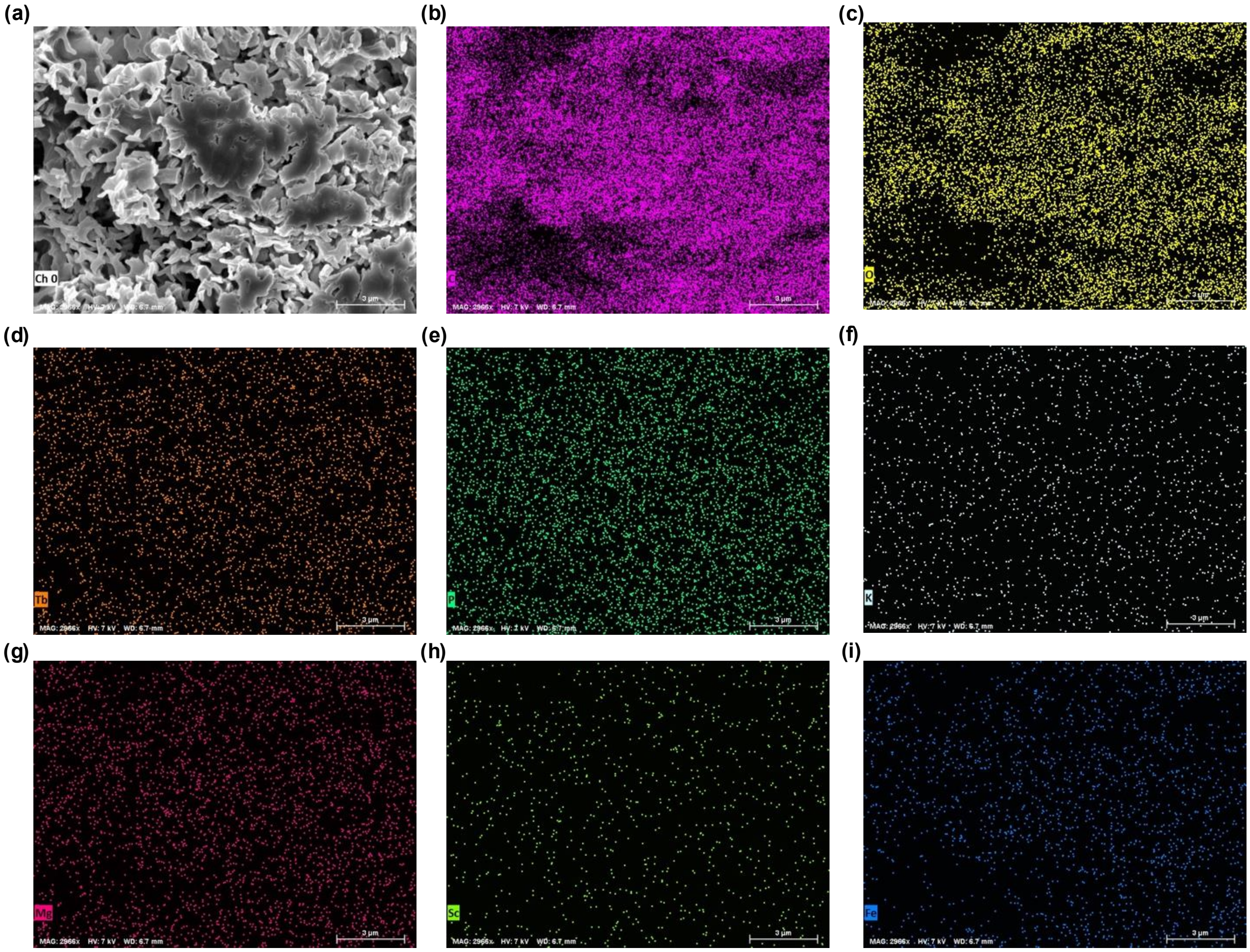

**Figure S35. SEM imaging and EDS mapping of Tb-exposed, EDTA-washed *A. ferrooxidans* biomass mounted on carbon tape.** (a) Electron image of the prepared biomass following Tb exposure and EDTA washing, lyophilization, mounting on carbon tape, and sputter-coating before analysis (Text S12). Individual EDS maps show software-assigned X-ray channels assigned to (b) C, (c) O, (d) Tb, (e) P, (f) K, (g) Mg, (h) Sc, and (i) Fe. All panels show the same field of view from one prepared sample.

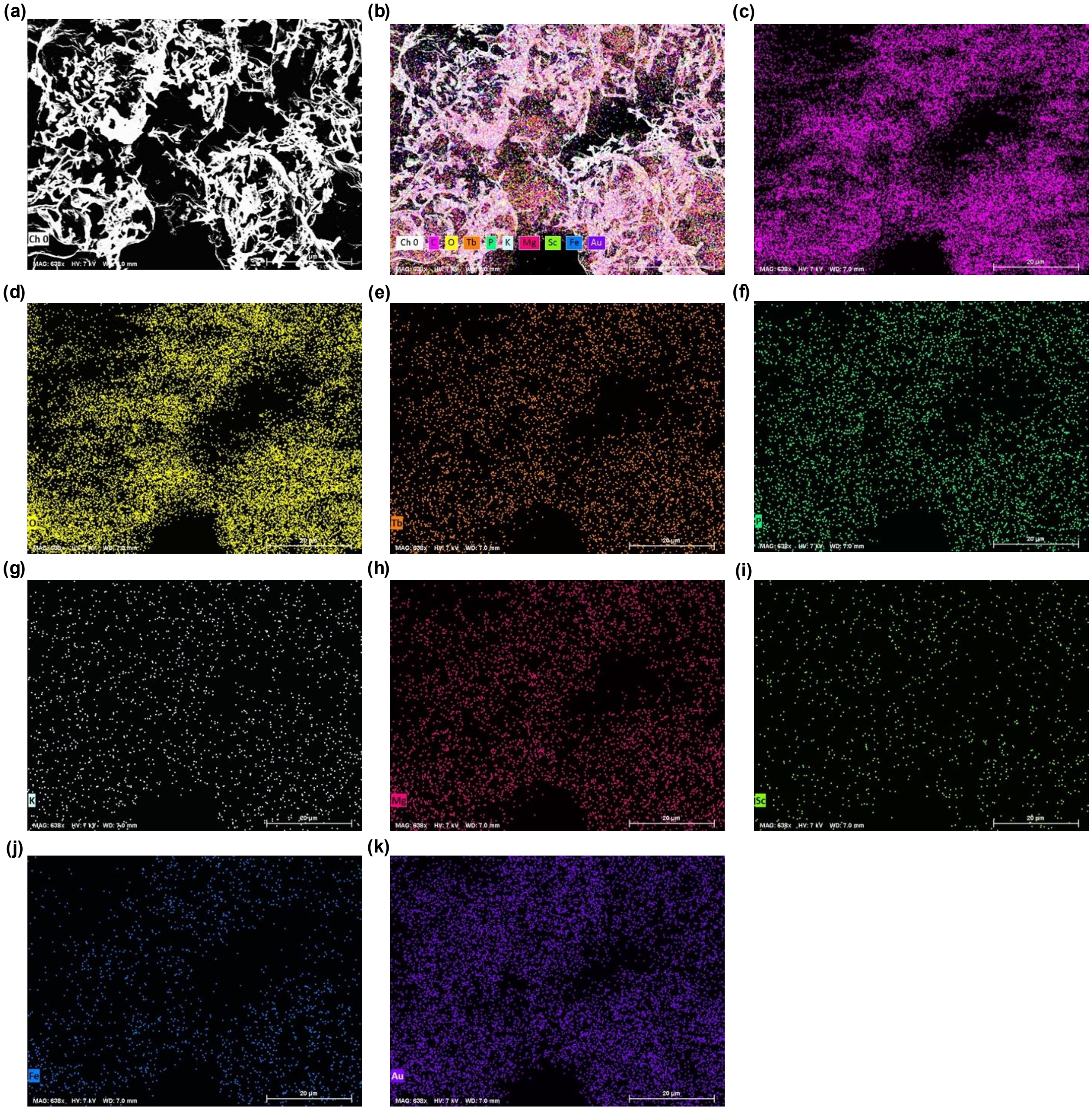


**Figure S36. SEM imaging and EDS mapping of *A. ferrooxidans* biomass collected before Tb-binding experiments and mounted on carbon tape.** Control biomass represents the cells before the Tb binding experiments. (a) Electron image and (b) composite overlay of the selected EDS channels for biomass after lyophilization, mounting on carbon tape, and sputter-coating before analysis (Text S12). Individual maps show software-assigned X-ray channels assigned to (c) C, (d) O, (e) Tb, (f) P, (g) K, (h) Mg, (i) Sc, (j) Fe, and (k) Au.

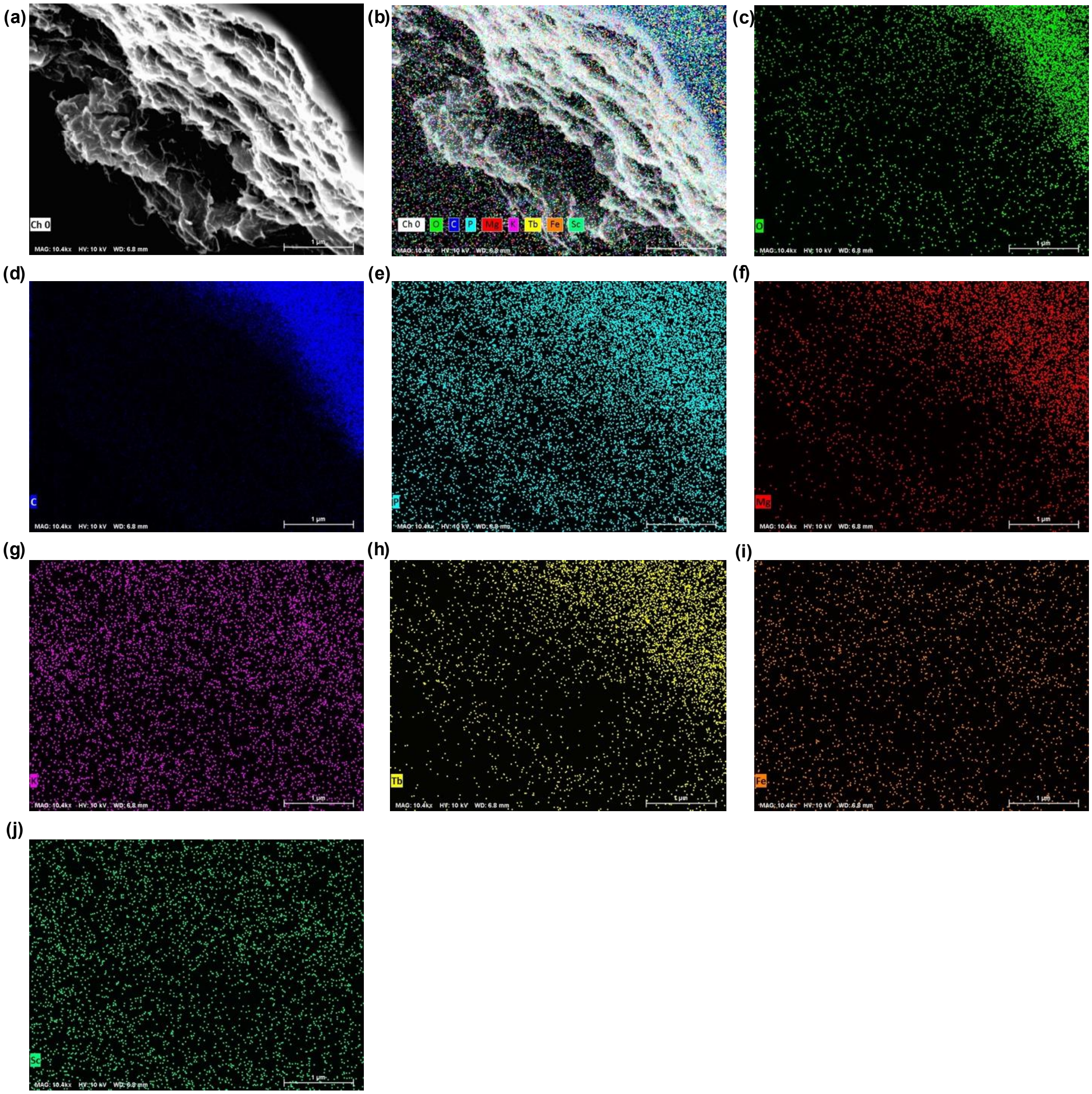


**Figure S37. SEM imaging and EDS mapping of Tb-exposed, EDTA-washed *A. ferrooxidans* biomass mounted on a silicon wafer.** (a) Electron image and (b) composite overlay of the prepared biomass following Tb exposure and EDTA washing, lyophilization, mounting on silicon wafer, and sputter-coating before analysis (Text S12). Individual EDS maps show software-assigned X-ray channels assigned to (c) O, (d) C, (e) P, (f) Mg, (g) K, (h) Tb, (i) Fe, and (j) Sc. All panels show the same field of view from one prepared sample.

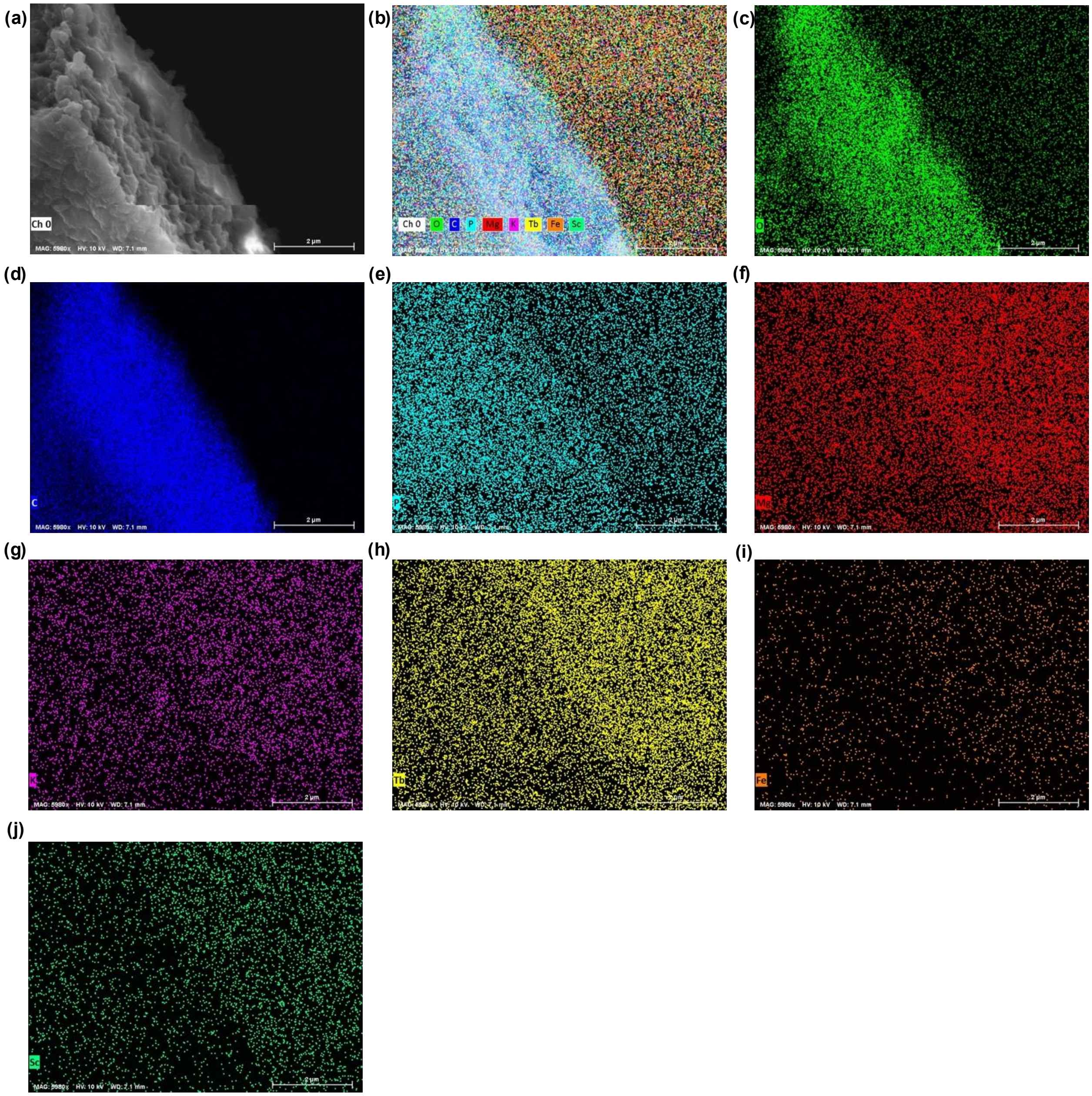


**Figure S38. SEM imaging and EDS mapping of Tb-exposed, water-washed *A. ferrooxidans* biomass mounted on a silicon wafer.** (a) Electron image and (b) composite overlay of the prepared biomass following Tb exposure and water washing, lyophilization, mounting on silicon wafer, and sputter-coating before analysis (Text S12). Individual EDS maps show software-assigned X-ray channels assigned to (c) O, (d) C, (e) P, (f) Mg, (g) K, (h) Tb, (i) Fe, and (j) Sc. All panels show the same field of view from one prepared sample.

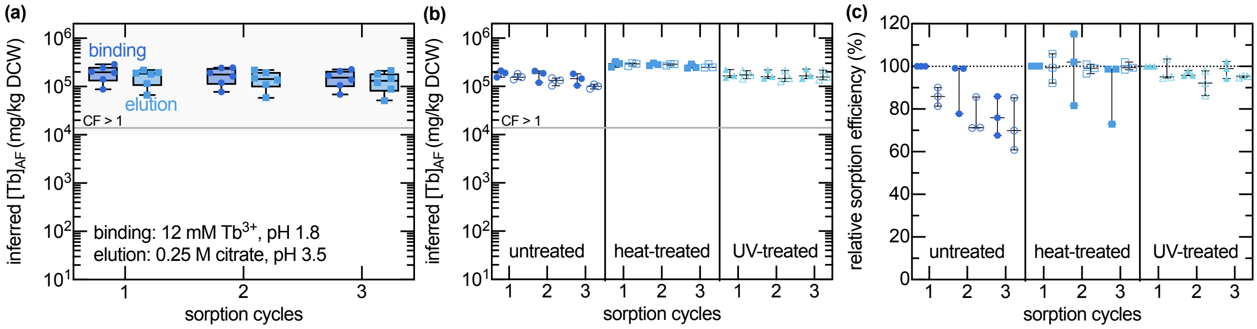


**Figure S39. Inferred Tb binding, citrate elution, and recovery across three adsorption–desorption cycles and different *A. ferrooxidans* pretreatments.** Binding tests were performed in AFM1 BS supplemented with 12.2 mM Tb from $Tb_2(SO_4)_3$ at pH 1.8 under standard conditions ($OD_{600} = 0.1$, 30 °C, 150 rpm, 8 h). (a) Inferred Tb-binding and citrate-elution capacity across three sequential sorption cycles ($n = 6$). (b) Inferred Tb-binding and citrate-elution capacity across three sequential sorption cycles for different pretreatment conditions and (c) the underlying sorption cycle efficiencies ($n = 3$). Biological replicates are shown as symbols, with wide bars indicating the mean and whiskers the minimum-maximum range. The gray horizontal reference line in (a-b) indicates the concentration factor (CF) threshold of 1 relative to the corresponding reference for the REE-enriched *P. americana* shoots. The dotted horizontal line in (c) indicates 100% mass balance closure.

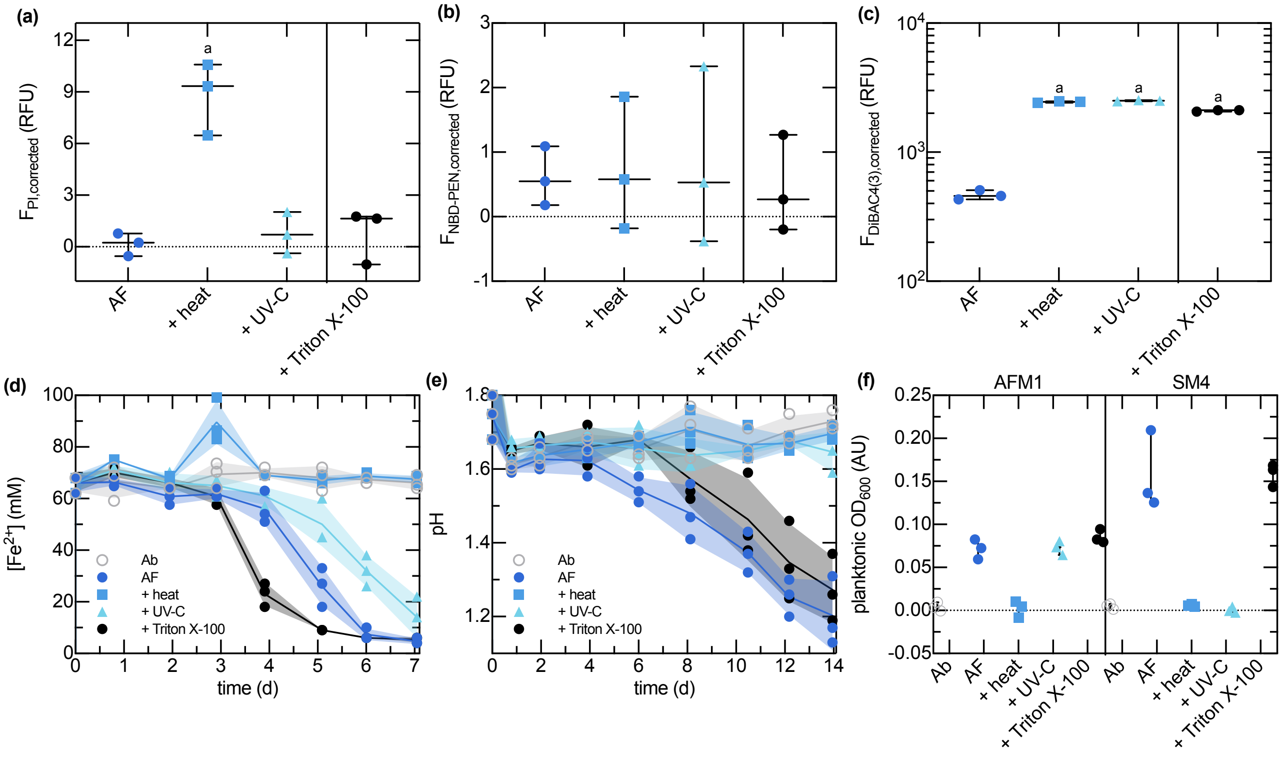


**Figure S40. Membrane-associated fluorescence, (apparent) substrate oxidation, and growth of *A. ferrooxidans* following heat, UV-C, and Triton X-100 and sonication treatments.** Conditions were untreated cells (AF), heat treatment at 90 °C, UV-C exposure at 254 nm, and 1% (v/v) Triton X-100 treatment and sonication (Text S17). Baseline-corrected fluorescence from (a) propidium iodide (PI), (b) NBD-PEN, and (c) $DiBAC_4(3)$ assays. Biological replicates are shown as symbols, with wide bars indicating the mean and whiskers the minimum–maximum range. "a" indicates a difference from the untreated *A. ferrooxidans* baseline (Table S34). (Apparent) metabolism of energy substrates in (d) $Fe^{2+}$- and (e) brown-$S^0$-containing growth media. Lines and shaded regions represent mean ± standard deviations, with biological replicates overlaid (f) Endpoint planktonic cell density measurements for the experiments plotted in (d-e). Biological replicates are shown as symbols, with wide bars indicating the mean and whiskers the minimum–maximum range.

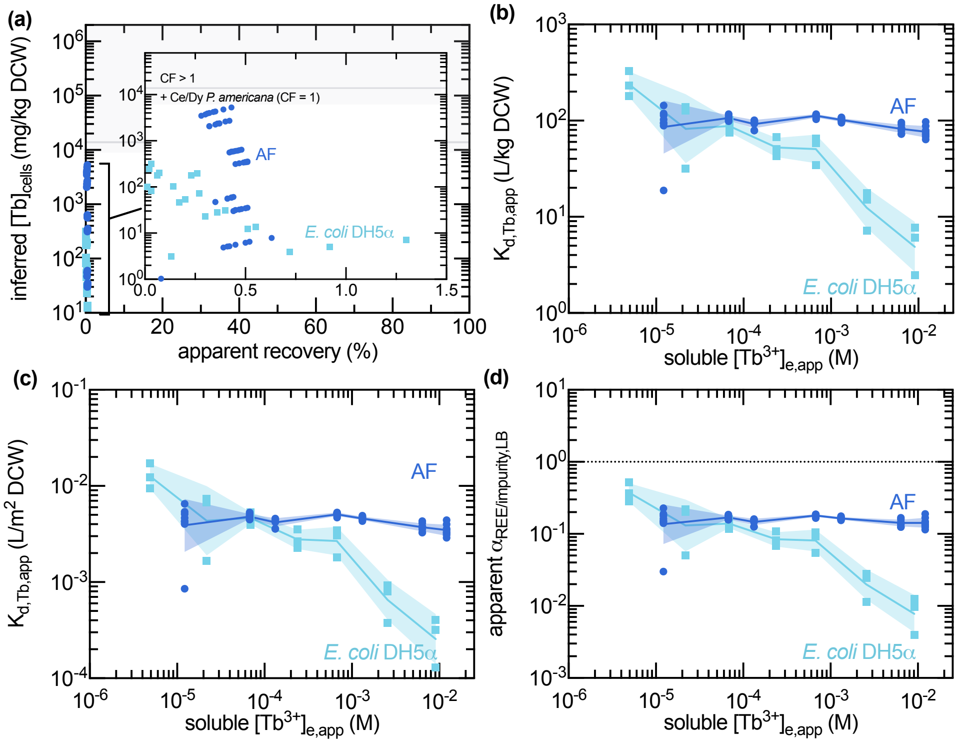


**Figure S41. Tb grade–recovery relationships, apparent partitioning, and lower-bound selectivity estimates for *A. ferrooxidans* and *E. coli* DH5$\alpha$ at pH 1.8.** The (a) grade–recovery curve, condition-specific apparent solid–liquid distribution coefficients, (b) without and (c) with surface-area-normalization, and (d) estimated lower bound of selectivity for Tb over major impurities for the data plotted in Figure 4a based on the binding media composition (Table S3). Lines and shaded band represent geometric mean ± standard deviations for (b–d), with biological replicates overlaid as points for *A. ferrooxidans* ($n = 8$) and *E. coli* DH5$\alpha$ ($n = 3$). The gray horizontal reference line indicates the concentration factor (CF) threshold of 1 relative to the corresponding reference for the REE-enriched *P. americana* shoots. Insets in (a) show close-ups of data for visual inspection.

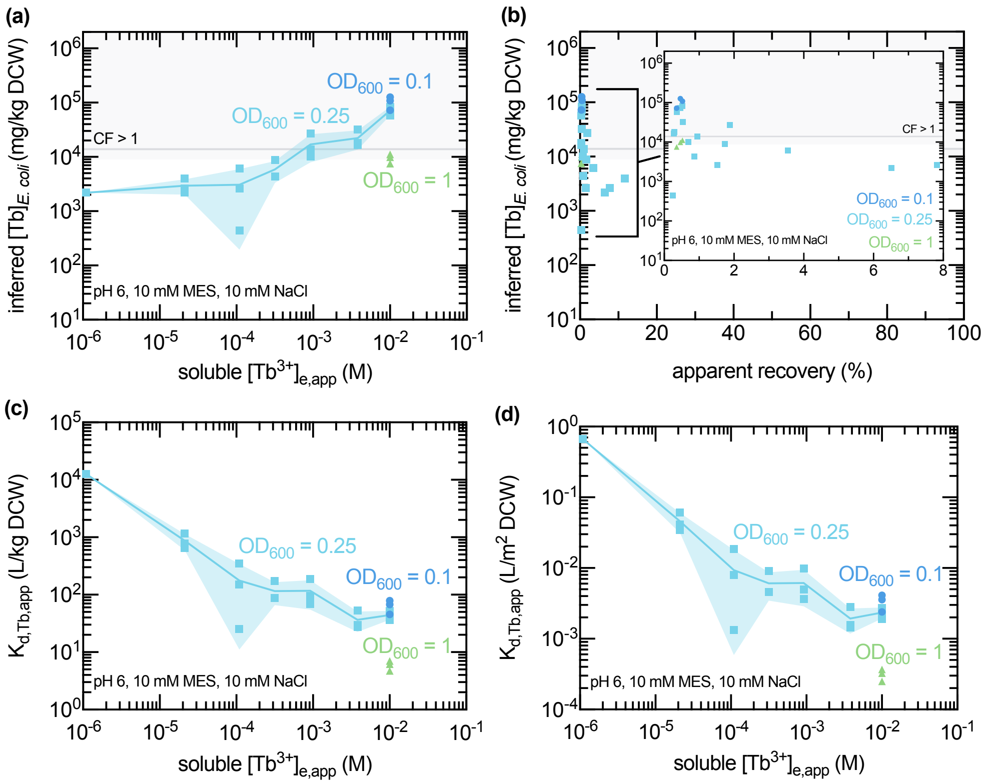


**Figure S42. Inferred Tb loading, recovery, and apparent partitioning by *E. coli* DH5$\alpha$ as a function of initial Tb concentration and cell density at pH 6.0.** Tests were performed at different cell densities ($OD_{600}$ = 0.1, 0.25, and 1.0), and inferred Tb was normalized to dry cell weight (DCW). (a) Mass-normalized inferred Tb associated with *E. coli* DH5$\alpha$ as a function of the apparent equilibrium concentration of soluble Tb and initial cell density. (b) Apparent grade–recovery curve for the data from (a). Inset shows close-up of data for visual inspection. The gray horizontal reference line indicates the concentration factor (CF) threshold of 1 relative to the corresponding reference for the REE-enriched *P. americana* shoots. Condition-specific apparent solid–liquid distribution coefficients, (c) without and (d) with surface-area-normalization. For (a,c–d), lines and shaded regions show mean ± standard deviation, with biological replicates overlaid (n = 3).

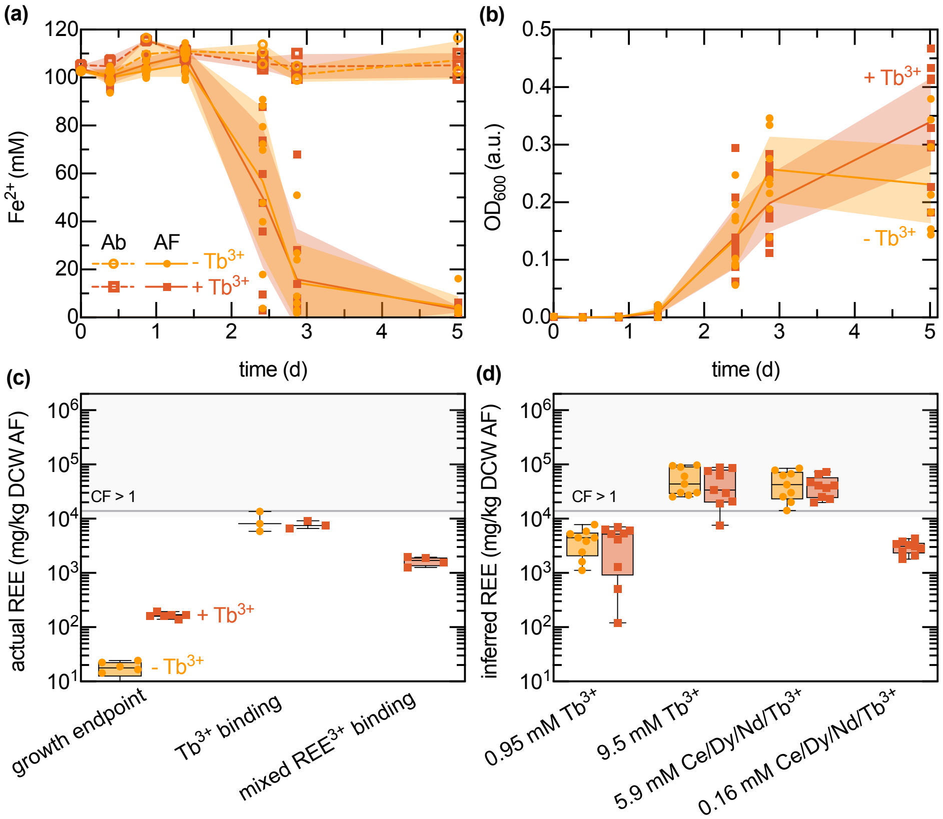


**Figure S43. $Fe^{2+}$ oxidation, growth, and subsequent REE association of *A. ferrooxidans* cultured with or without 30 $\mu$M $Tb^{3+}$ from $Tb_2(SO_4)_3$.** *A. ferrooxidans* was grown in F2S medium with or without approximately 30 $\mu$M $Tb^{3+}$ to evaluate whether this exposure altered $Fe^{2+}$ biooxidation, planktonic cell density, or subsequent REE association. (a) Time-course profiles of $Fe^{2+}$ concentrations in abiotic controls and *A. ferrooxidans* cultures. (b) Time-course profiles of planktonic cell density for *A. ferrooxidans* cultures, plotted as $OD_{600}$. Lines and shaded regions show mean ± standard deviation, with biological replicates overlaid ($n = 3 - 9$). (c) Cell-associated Tb or REE concentrations measured after growth in F2S medium with or without 30 µM $Tb^{3+}$, and after subsequent exposure to 12 mM $Tb^{3+}$ or equimolar $Ce^{3+}/Dy^{3+}/Nd^{3+}/Tb^{3+}$ solutions at pH 1.8. (d) Inferred REE-binding capacity of *A. ferrooxidans* after subsequent exposure to the indicated conditions. Simulants at pH 1.8 were used in all tests except for the 0.16 mM $Ce^{3+}/Dy^{3+}/Nd^{3+}/Tb^{3+}$ condition, which was a pH 1.0 *P. acinosa* shoot leachate. Individual biological replicates are shown as points with min-to-max box-and-whisker plots (n = 3–9). Statistical comparisons for panels (c) and (d) are provided in Table S35 and Table S36, respectively.

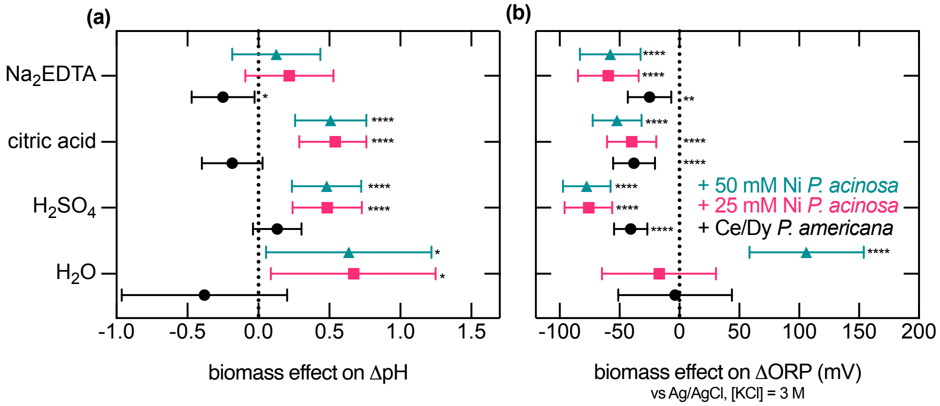


**Figure S44. Biomass-associated shifts in pH and ORP relative to matched no-shoot controls during water, acid, and EDTA extraction of *Phytolacca* shoots.** Biomass effects were calculated from 7 d leaching controls containing $H_2O$, $H_2SO_4$, citric acid, or $Na_2EDTA$. $\Delta pH$ and $\Delta ORP$ were calculated as final minus initial values. (a) Biomass effects on pH are shown as $\Delta\Delta pH = \Delta pH_{\text{biomass}} - \Delta pH_{\text{no shoots}}$, such that positive values indicate higher pH relative to the matched no-shoot control. (b) Biomass-driven ORP changes are calculated and shown analogously. Points show estimated mean differences, and horizontal bars show 95% confidence intervals. Data were analyzed by ordinary two-way ANOVA with reactant, tissue condition, and their interaction as fixed effects, followed by Dunnett-adjusted comparisons of each biomass-containing treatment against the corresponding no-shoot control within each reactant (Table S37, Table S38, Table S39). Asterisks indicate Dunnett-adjusted comparisons against the matched no-shoot control ($*P < 0.05$; $**P < 0.01$; $****P < 0.0001$).

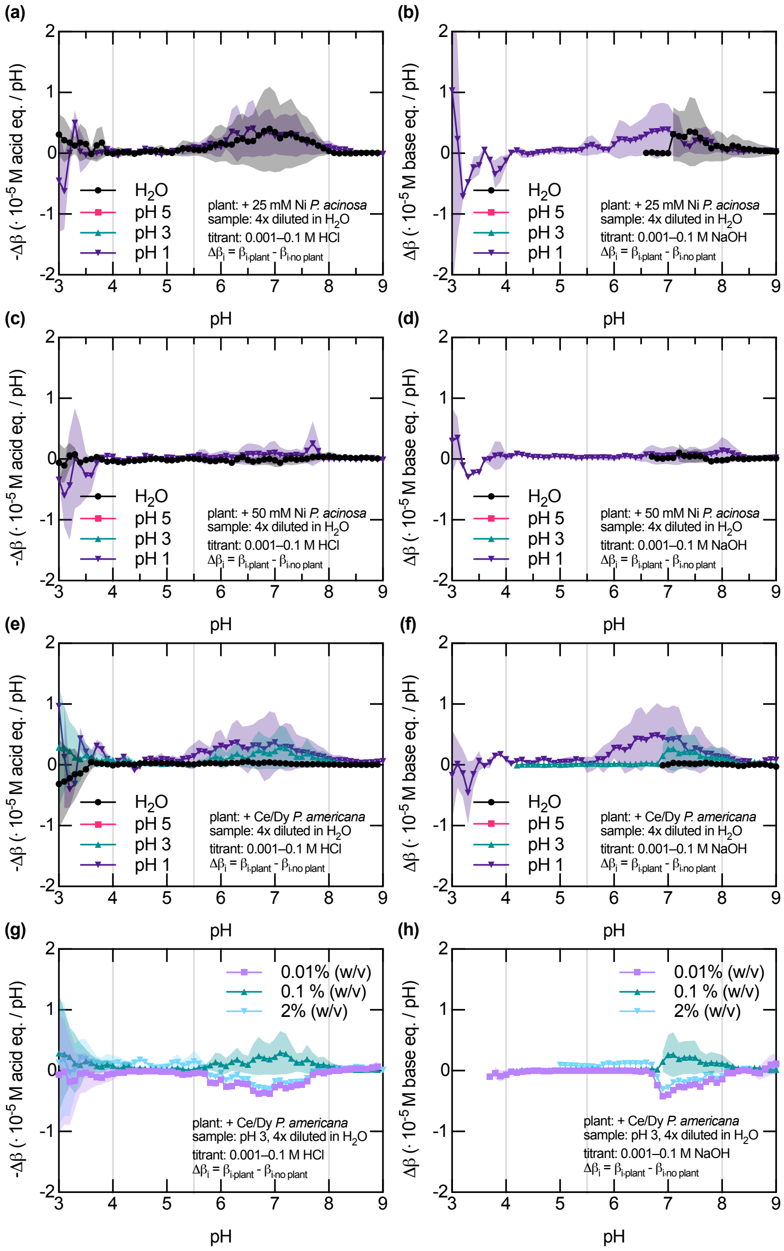


**Figure S45. Biomass-corrected acid- and base-side buffering profiles of *Phytolacca* shoot leachates as a function of initial pH and biomass loadings.** (a,b) Biomass-corrected acid- and base-side buffering profiles for +25 mM Ni-supplemented *P. acinosa* shoot leachates generated in $H_2O$ or at initial pH 5, 3, or 1. (c,d) Corresponding profiles for +50 mM Ni-supplemented *P. acinosa* shoot leachates. (e,f) Corresponding profiles for Ce/Dy-supplemented *P. americana* shoot leachates. (g,h) Biomass-loading dependence of the acid- and base-side buffering profiles for Ce/Dy-supplemented *P. americana* shoot leachates generated at initial pH 3 using 0.01, 0.1, or 2% (w/v) biomass. Acid-side buffering is shown as $-\Delta\beta$ and base-side buffering is shown as $\Delta\beta$, where $\Delta\beta_i = \beta_{i,\text{plant}} - \beta_{i,\text{no plant}}$ for the matched no-biomass leaching control. Lines and shaded regions show mean ± standard deviation, with interpolated pH values for biological replicates overlaid (n = 2).

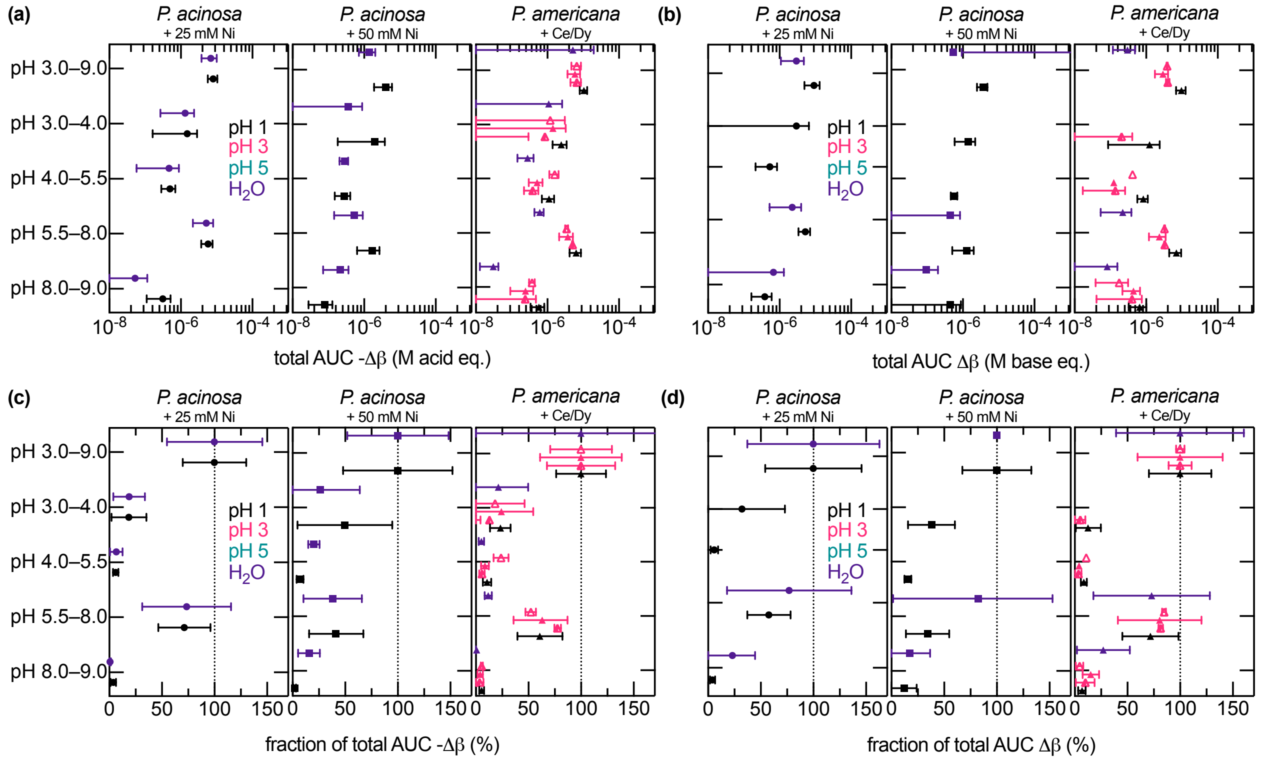


**Figure S46. Integrated buffering contributions and their distribution across pH intervals in Ni- and Ce/Dy-enriched *Phytolacca* shoot leachates.** Post-leach leachates generated from *P. acinosa* shoots supplemented with 25 or 50 mM Ni and *P. americana* shoots supplemented with Ce/Dy were titrated after extraction in $H_2O$ or acidified solutions initially adjusted to pH 5, 3, or 1 (Figure S45). (a) Total acid-side buffering contribution, calculated as the integrated area under $-\Delta\beta$ from pH 3.0–9.0. (b) Total base-side buffering contribution, calculated as the integrated area under $\Delta\beta$ from pH 3.0–9.0. (c) Fractional contribution of each pH interval to total acid-side buffering capacity. (d) Fractional contribution of each pH interval to total base-side buffering capacity. $\Delta\beta$ was calculated as the plant-containing leachate buffering capacity minus the corresponding no-plant control. Points show replicate-derived mean values, and error bars show 95% confidence intervals (n = 2).

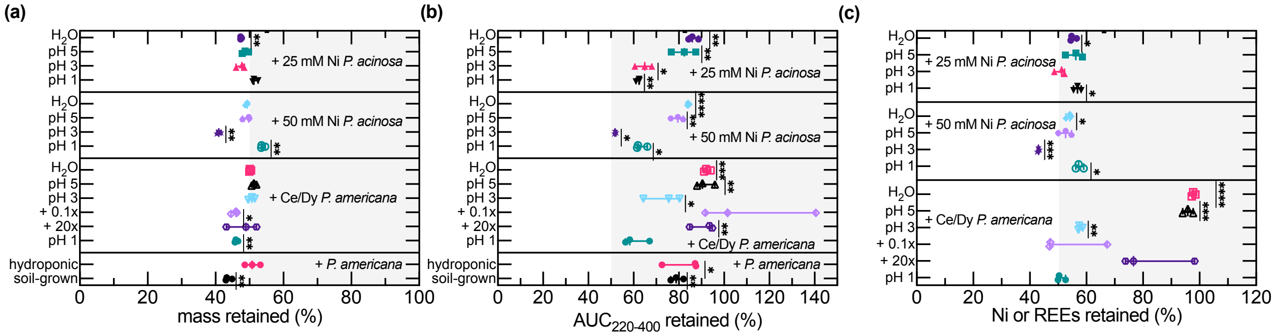


**Figure S47. Mass, UV absorbance, and Ni or REE retention during 3 kDa centrifugal ultrafiltration of some *Phytolacca* shoot leachates.** Retention metrics were calculated after the indicated extraction conditions for Ni-supplemented *P. acinosa* shoots, Ce/Dy-supplemented *P. americana* shoots, and non-enriched *P. americana* controls. (a) Percentage of mass retained above the 3 kDa cutoff after centrifugal filtration. (b) Percentage of the initial UV-absorbing material retained, calculated from integrated absorbance from 220–400 nm. (c) Percentage of the initial target metal retained in the solid fraction, shown as Ni retention for Ni-supplemented *P. acinosa* and REE retention for Ce/Dy-supplemented *P. americana*. Points show individual biological replicates, and horizontal bars show means. Asterisks indicate statistically significant differences from the corresponding reference condition or theoretical retention value, as defined in the associated statistical analyses provided in Table S40, Table S41, and Table S42, respectively.

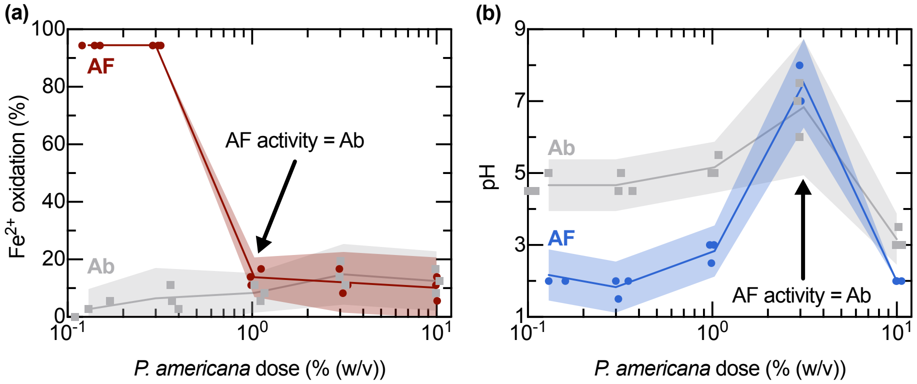


**Figure S48. Endpoint $Fe^{2+}$ oxidation and pH in *A. ferrooxidans* cultures grown with increasing *P. americana* shoot loading in $Fe^{2+}$- or $S^0$-containing media.** *A. ferrooxidans* (AF; initial $OD_{600} = 0.01$) was grown with increasing *P. americana* shoot loading in 5 mL medium in 14 mL test tubes at 30 °C and 150 rpm for 14 d. (a) In $Fe^{2+}$ medium, AF oxidized most of the initial $Fe^{2+}$ at low *P. americana* loadings, but $Fe^{2+}$ oxidation decreased sharply at $\geq$1% (w/v), approaching abiotic controls. (b) In $S^0$ medium, endpoint pH remained lower in AF cultures than abiotic controls at low *P. americana* loadings, consistent with biological acid generation, but endpoint pH increased strongly at intermediate biomass loading and converged with abiotic controls at $\geq$3% (w/v), indicating that biomass-associated buffering obscured sulfur-oxidation-dependent acidification. Lines and shaded regions show mean ± 95% confidence intervals, with biological replicates overlaid ($n = 3$).

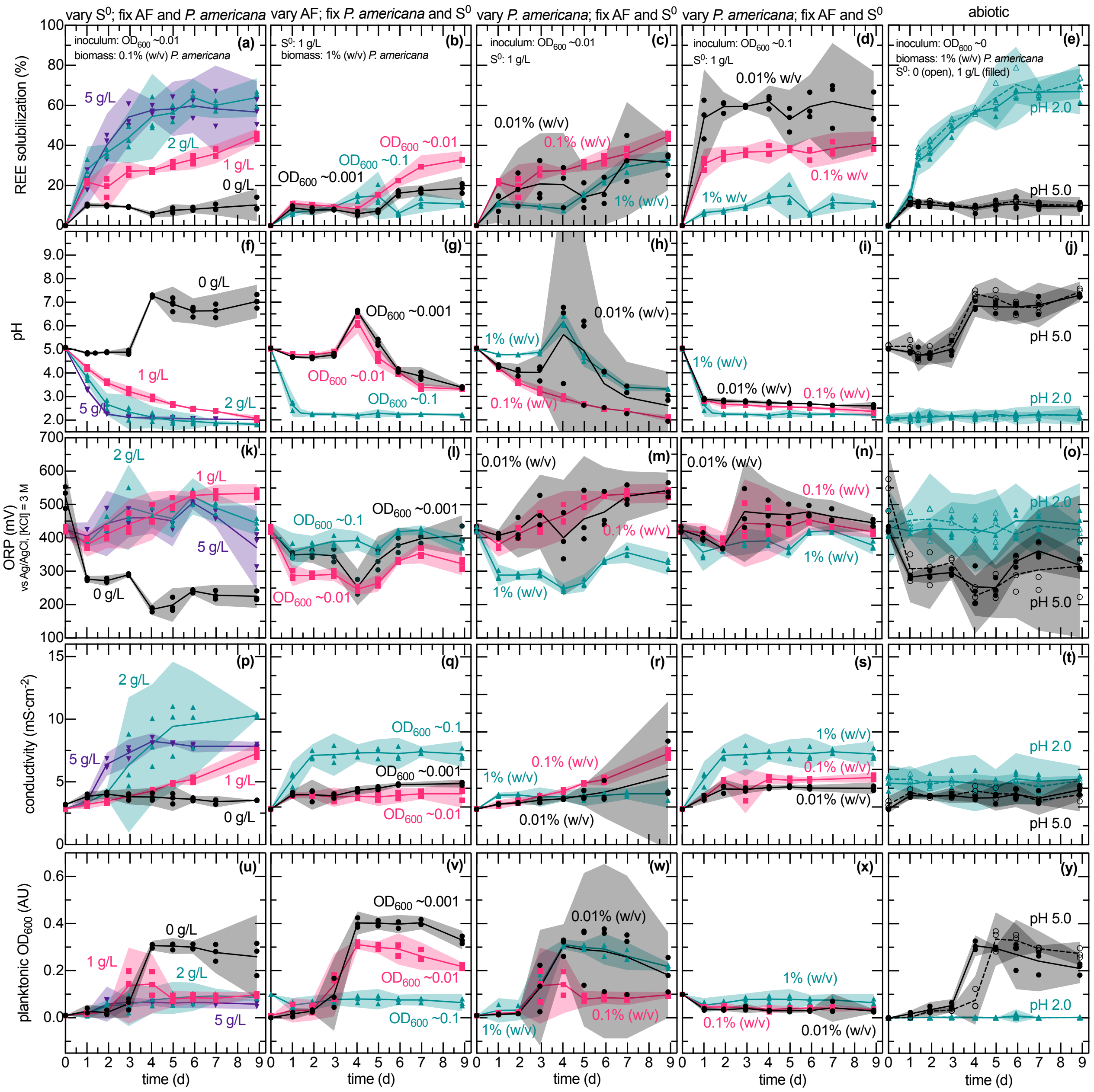


**Figure S49. Time course REE solubilization, solution chemistry, and apparent cell densities during *P. americana* bioleaching at varied shoot loadings, $S^0$ dosages, and *A. ferrooxidans* inoculum densities.** Time-resolved (a–e) REE solubilization, (f–j) pH, (k–o) solution potential (ORP; vs. Ag/AgCl, 3 M KCl), (p–t) conductivity, and (u–y) apparent planktonic cell density, plotted as $OD_{600}$, were measured during *A. ferrooxidans* cultivation with Ce/Dy-enriched *P. americana* shoot biomass to evaluate whether plant loading, sulfur availability, and inoculum density altered physicochemical conditions relevant to bioleaching. Lines and shaded regions show mean ± 95% confidence intervals, with biological replicates overlaid (n = 3). The following panels replot the same data for ease of visual comparison: 1 g/L $S^0$, 0.1% (w/v) *P. americana*, 0.01 $OD_{600}$ initial *A. ferrooxidans* density condition in columns 1 (a,f,k,p,u) and 3 (c,h,m,r,w), and the 1 g/L $S^0$, 1% (w/v) *P. americana*, 0.1 $OD_{600}$ initial *A. ferrooxidans* density condition in columns 2 (b,g,l,q,v) and 4 (d,i,n,s,x).

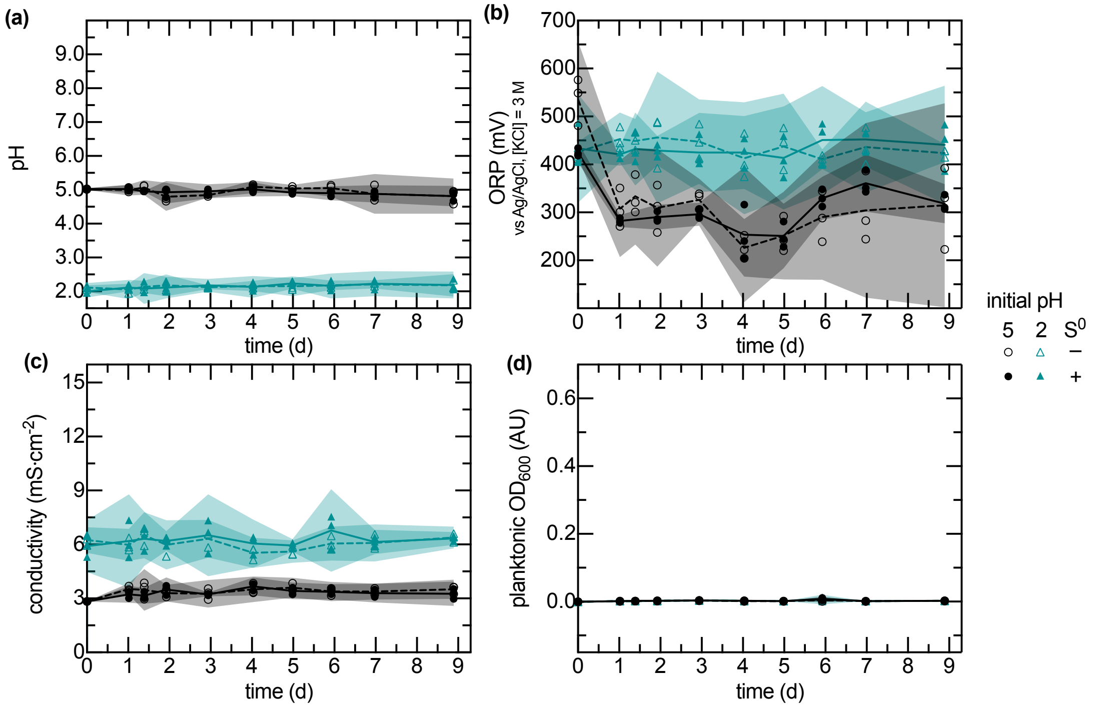


**Figure S50. Time course pH, ORP, conductivity, and apparent cell densities in bioleaching controls without *P. americana* shoots.** Time-resolved (a) pH, (b) solution potential (ORP; vs. Ag/AgCl, 3 M KCl), (c) conductivity, and (d) apparent planktonic cell density, plotted as $OD_{600}$, were measured under identical conditions to the corresponding panels in Figure S49j,o,t,y in the absence of Ce/Dy-enriched *P. americana* shoots Lines and shaded regions show mean ± 95% confidence intervals, with independent replicates overlaid (n = 3).

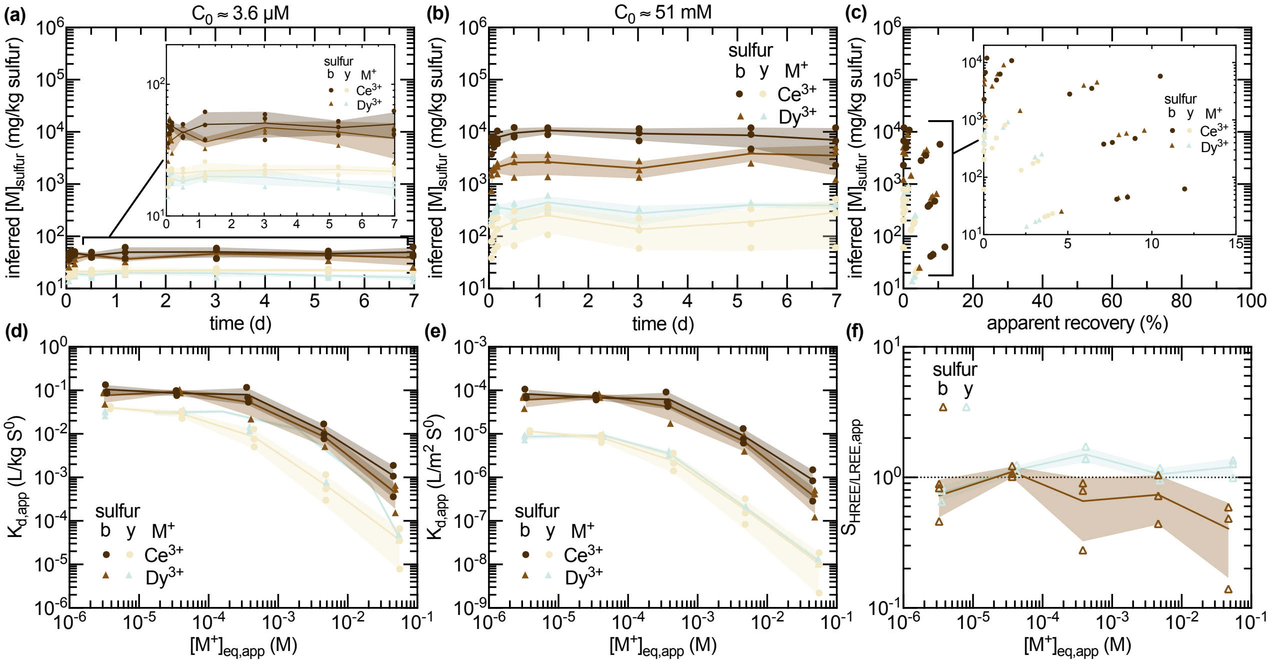


**Figure S51. Inferred Ce and Dy loading, apparent partitioning, and Dy/Ce selectivity of brown and yellow $S^0$ at pH 2.0.** Either 1 g/L brown (b) or elemental/yellow (y) $S^0$ were added to SM4 medium at pH 2.0 across the tested $Ce^{3+}$ and $Dy^{3+}$ concentrations, which were supplied as sulfate salts. Time-dependent inferred $S^0$-associated REE loading from initial soluble concentrations of about (a) 3.6 µM and (b) 51 mM, which represented the lowest and highest concentrations tested herein, respectively. (c) Inferred $S^0$-associated REE loading as a function of apparent recovery from the initial soluble concentration after 7 d across all tested concentrations, including the endpoint time-course data shown in (a) and (b). Condition-specific apparent solid–liquid distribution coefficients as a function of equilibrium soluble metal concentration (d) before and (e) after surface-area normalization. (f) Condition-specific apparent selectivity of HREE over LREE, with $Dy^{3+}$ and $Ce^{3+}$ representing the HREE and LREE, respectively, as a function of equilibrium soluble metal concentration. Lines and shaded regions show mean ± standard deviations, with independent experimental replicates overlaid ($n = 3$). Insets in (a) and (c) show close-up views of data for visual inspection.

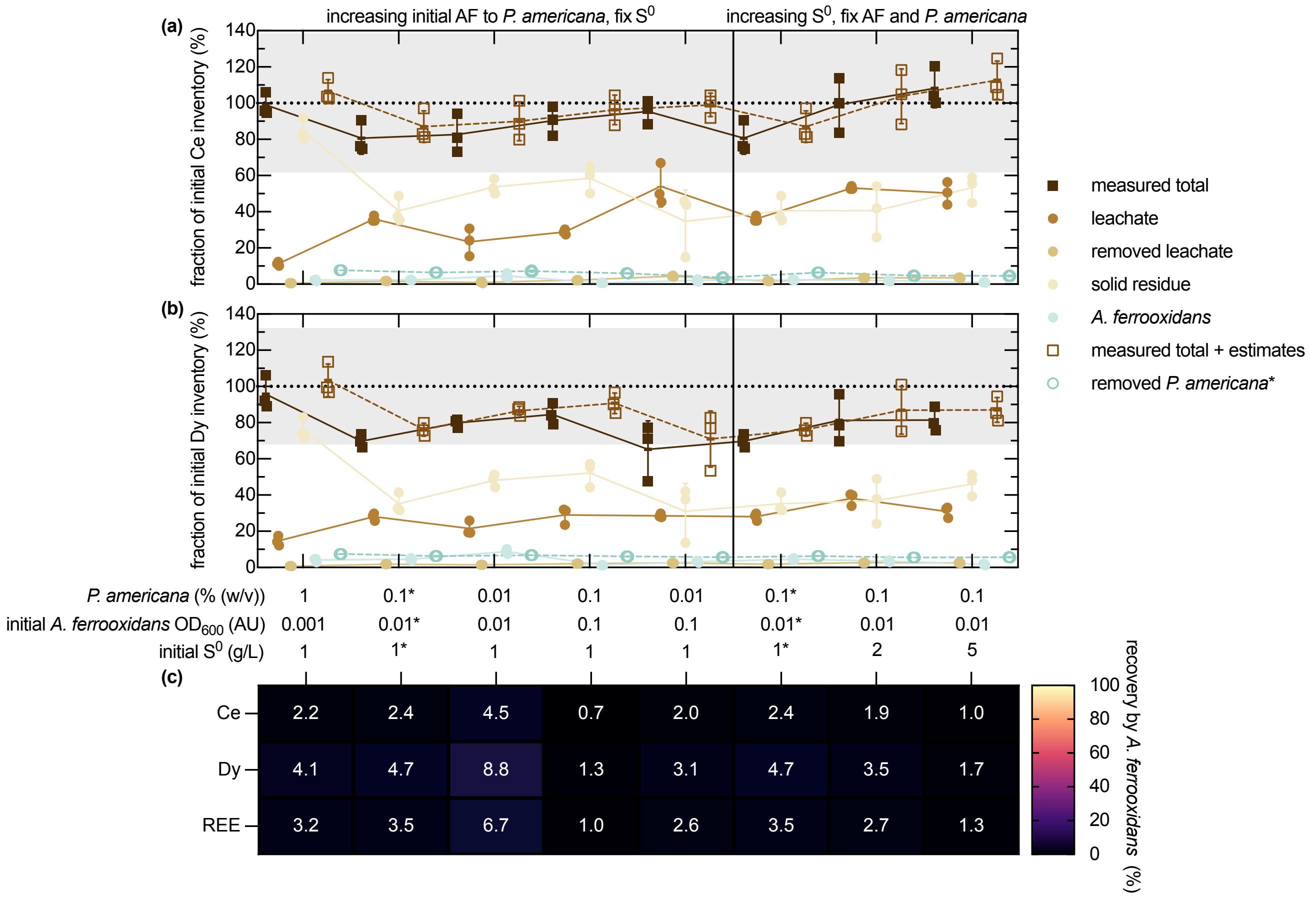


**Figure S52. Ce and Dy mass balances and cell-associated recovery in EDTA-washed *A. ferrooxidans* pellets after bioleaching of *P. americana* shoots for some experiments.** Endpoint inventories of (a) Ce and (b) Dy in the final leachate, leachate removed during serial sampling, recovered solid residue, and EDTA-washed *A. ferrooxidans* pellets are expressed as percentages of the estimated initial inventory of each element ($n = 3$). Initial inventories were calculated from the initial dry shoot mass and mean elemental concentrations of the Ce/Dy-enriched *P. americana* material (Table S4). Filled squares show the measured total obtained by summing these four fractions. Open circles show the estimated REE inventory removed with plant material during serial sampling, and open squares show the measured total plus this estimated contribution. The sampled-plant contribution assumes concomitant removal of plant solids during collection of the 1 mL aliquots; the removed plant masses were not measured. Lines and error bars show mean ± SD, with individual replicates overlaid (n = 3). The dotted horizontal lines indicate 100% of the estimated initial inventory. Gray bands show the initial-inventory variation corresponding to the mean ± standard deviations of the plant elemental concentrations, normalized to the mean. Asterisks in the condition labels identify the shared baseline of 0.1% (w/v) *P. americana*, initial *A. ferrooxidans* $OD_{600} = 0.01$, and 1 g/L $S^0$, which is replotted in both blocks for ease of comparison. (c) Heatmap of mean Ce, Dy, and combined REE recovery in the washed *A. ferrooxidans* pellets ($n = 3$). Cell annotations report the mean percentages.

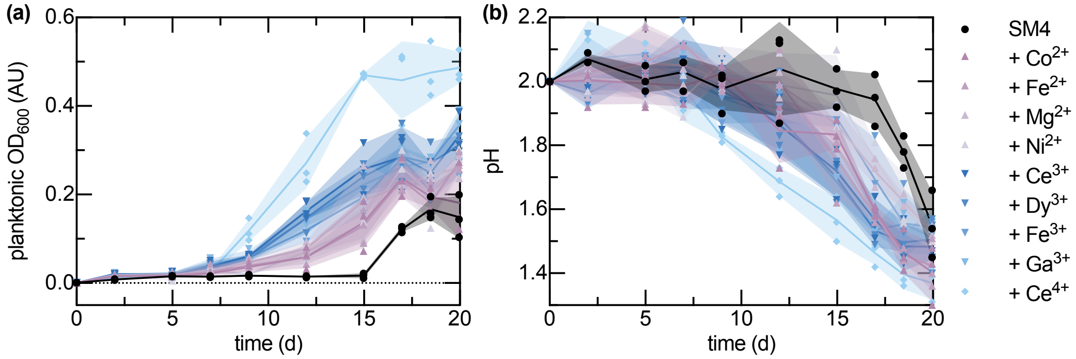


**Figure S53. Time course of planktonic *A. ferrooxidans* density and pH under 100 $\mu$M cation supplementation in the absence of plant biomass.** Cultures grown in shaker flasks with 100 µM of the indicated supplemented cation (30 °C, 150 rpm, initial $OD_{600} = 0, 0.001$). Time-resolved (a) planktonic cell density measurements, expressed as $OD_{600}$, and (b) pH measurements. Lines and shaded regions show mean ± standard deviations, with independent replicates overlaid (n = 3).

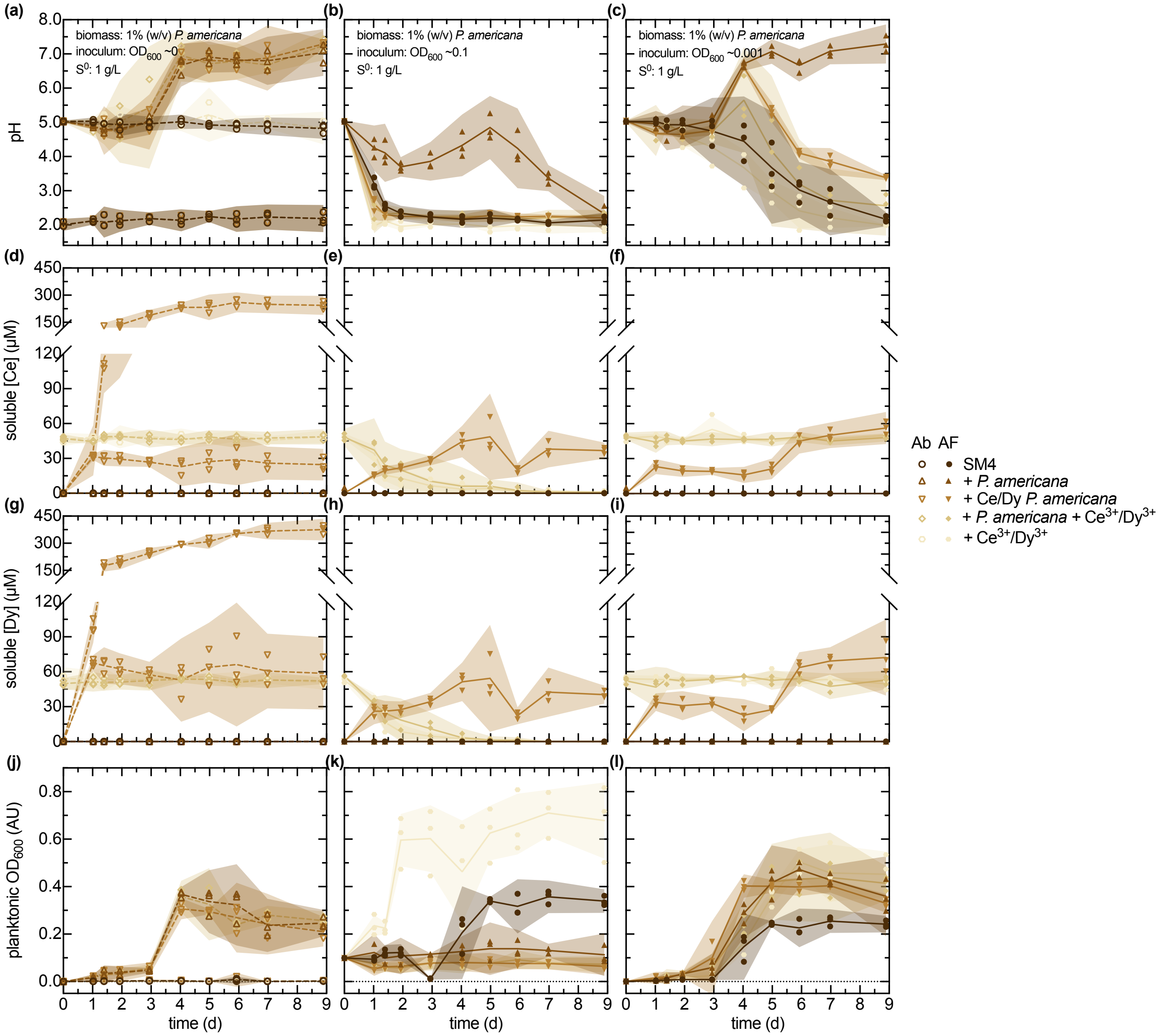


**Figure S54. Time courses of pH, soluble Ce and Dy, and planktonic cell density across *P. americana* shoot-enrichment, dissolved-REE supplementation, and *A. ferrooxidans* inoculation conditions.** Time-resolved (a–c) pH, (d–f) soluble Ce, (g–i) soluble Dy, and (j–l) planktonic cell density measurements, expressed as $OD_{600}$, collected in the presence and absence of *P. americana*, with and without prior Ce/Dy-enrichment. Additionally, some cultures were supplemented with equimolar Ce and Dy with and without the *P. americana* with no prior Ce/Dy-enrichment. Lines and shaded regions show mean ± 95% confidence intervals, with independent replicates overlaid (n = 3). Several data series replot the same data presented in Figure S49 and Figure S50 for ease of visual comparison.

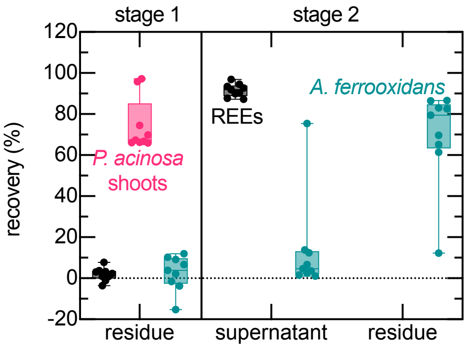


**Figure S55. Partitioning of plant solids, REEs, and *A. ferrooxidans* during two-stage centrifugation of *P. acinosa* bioleaching cultures.** Recovery of *P. acinosa* shoot solids, REEs, and *A. ferrooxidans* is shown across a two-stage centrifugation workflow. In stage 1, the residue fraction was analyzed to evaluate recovery of leached plant shoot solids relative to REEs and *A. ferrooxidans*. In stage 2, the remaining suspension was separated into supernatant and residue fractions to compare partitioning of soluble REEs and *A. ferrooxidans*. Recovery values are shown as percentages of the corresponding input to each stage. Points show individual replicate overlaid on min-to-max box-and-whisker plots (n = 9).

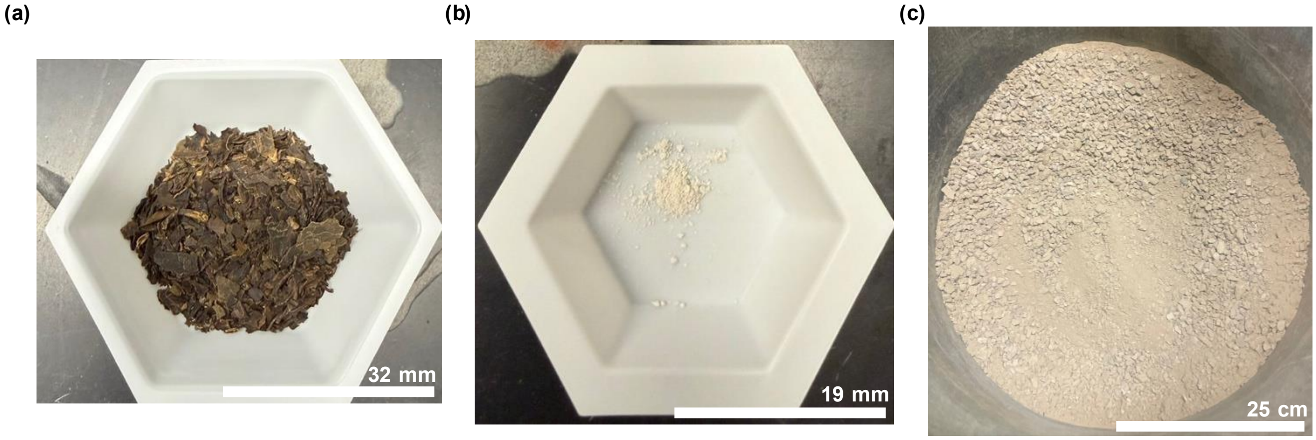


**Figure S56. Photographic comparison of dried Ce/Dy-enriched *P. americana* shoots, ash produced at 600 °C, and monazite-bearing ore.** Representative examples of the (a) dried Ce/Dy enriched *P. americana* shoots and (b) ash generated from calcination at 600 °C. (c) Mixed bastnäsite–monazite ore for visual comparison, courtesy of Steven Lycans II. Neither the plant nor the ash shown in (a–b) looked like this ore shown in (c), or reported examples of the pure mineral specimen.[63]

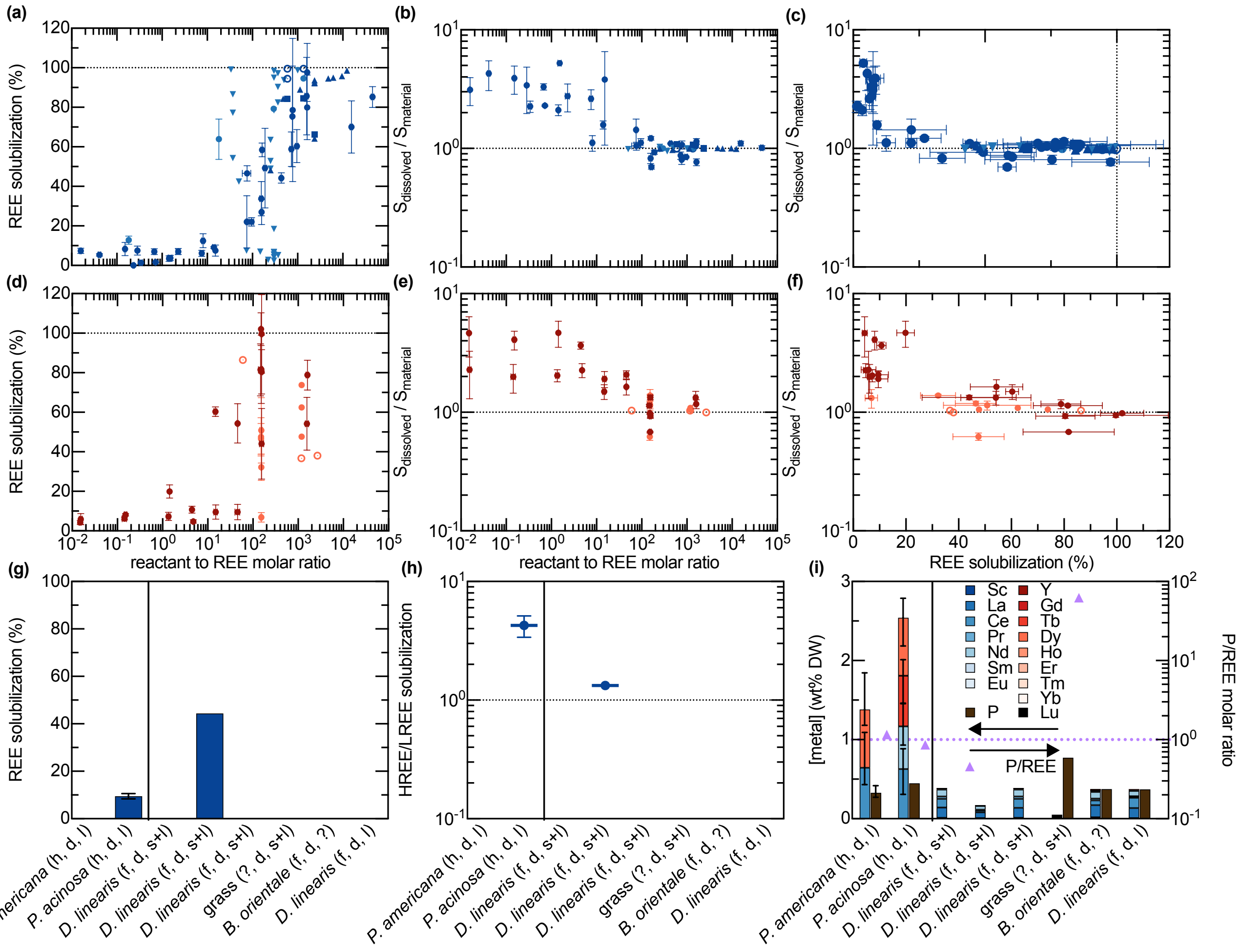


**Figure S57. Literature comparison of REE solubilization, feed-normalized HREE/LREE selectivity, and REE and P composition in plant tissues and thermally treated residues.** Standard 7 d extraction endpoints from this study are compared with published measurements from other REE-enriched plant materials and their thermally treated products.[32–37] (a—c) Acid and (d—f) chelator extraction data are expressed as (a,d) REE solubilization versus nominal reactant-to-REE molar ratio, (b,e) feed-normalized HREE/LREE selectivity versus reactant-to-REE molar ratio, and (c,f) selectivity versus REE solubilization. Feed-normalized selectivity $S_{\text{dissolved}}/S_{\text{material}}$ reflects the molar HREE/LREE ratios ($S$); values above unity indicate preferential fractional release of HREEs. (g,h) Corresponding REE solubilization and selectivity during water extraction. (i) Elemental concentrations on a dry-weight basis, shown as stacked REE bars and separate P bars on the left axis, with P/REE molar ratios shown as triangles on the right axis. REE totals and ratios include the elements quantified for each material. In (a–f), extraction reagents are distinguished by color: $H_2SO_4$, $HNO_3$, HCl, and MSA are represented by progressively lighter blue shades, while $Na_2EDTA$ and citric acid are represented by dark red and coral, respectively. Symbol shapes identify source groups: circles, this study; squares;[34] upward triangles;[32] downward triangles;[33] diamonds;[36] hexagons;[64]pentagons;[37] stars;[35] plus signs;[65] and crosses.[66] Open symbols indicate added $KNO_3$. Measurements from this study are shown as means with standard deviations. Literature values were transcribed or digitized from the reported results.[4] Axis abbreviations indicate hydroponic enrichment (h), field collection (f), dried material (d), stems (s), leaves (l), and unspecified information (?). Dotted lines indicate 100% solubilization, unity selectivity, or P/REE = 1, as applicable. Comparisons are qualitative because material preparation, extraction conditions, and analytical endpoints substantially differed among studies as summarized in Text S24.
